# AI-guided discovery of atypical protein assemblies

**DOI:** 10.64898/2026.05.03.722499

**Authors:** AmirAli Toghani, Benjamin A. Seager, Yu Sugihara, Lisa-Marie Roijen, Juan M. Azcue, Maián Garro, Maryam Sargolzaei, Ioanna Morianou, Adeline Harant, Sam Gallop, Jiorgos Kourelis, Dan MacLean, Mauricio P. Contreras, Sophien Kamoun, Daniel Lüdke

## Abstract

Artificial intelligence (AI) systems such as AlphaFold have transformed structural biology by enabling accurate prediction of protein structures. However, their capacity to uncover new classes of macromolecular assemblies remains largely untapped. We developed the Structural Novelty Index (SNI), a quantitative framework for identifying protein complexes that diverge from canonical architectures. As one implementation of SNI, we developed SNI_*NRC-Hexa*_, to identify unconventional resistosomes formed by nucleotide-binding, leucine-rich repeat immune receptors (NLRs). We used it to analyze AlphaFold 3 models of 637 non-redundant NRC proteins from 346 genomes representing 85 plant species. This analysis identified candidates with predicted architectures distinct from the canonical hexameric resistosomes of NRC proteins. Biochemical purification and negative-stain transmission electron microscopy of NRC7 orthologs from multiple species supported the SNI prediction and revealed an unexpected undecameric (11-mer) assembly. Our results establish SNI as a scalable approach for discovering atypical protein complexes.

## Main Text

Most proteins function as dynamic assemblies that underpin cellular homeostasis and physiology. AI-driven structural prediction has largely solved the protein folding problem, transforming structural biology (*1, 2*), yet its potential to discover higher-order protein assemblies remains only partly realised. In particular, tools such as AlphaFold have been underused for identifying atypical or unrecognized macromolecular architectures, leaving much of the cellular assembly landscape unexplored. Here, we present an AI-guided framework to systematically identify and prioritize unexpected protein assemblies and apply it to the discovery of atypical oligomeric assemblies of nucleotide-binding leucine-rich repeat (NLR) proteins.

NLRs are intracellular immune receptors encoded in organisms across the tree of life, from bacteria and fungi to plants and animals (*3–6*). Upon activation, NLRs undergo oligomerization into supramolecular complexes—termed inflammasomes in animals and bacteria and resistosomes in plants—that function as signaling platforms or membrane channels to initiate defence responses. In plants, NLRs represent one of the most expansive and diverse protein families, roughly comprising 1% of plant proteomes (*7*). Phylogenetic analyses revealed multiple distinct clades that assemble into resistosomes with defined stoichiometries, including tetrameric, pentameric, hexameric, and octameric architectures (*8–14*). One clade of coiled-coil NLRs (CC-NLRs) comprises the NRCs (NLR required for cell death), a deep lineage of helper NLRs that function downstream of pathogen-detecting receptors known as NRC-dependent sensors (NRC-S) (*15*). The NRC superclade groups the helper NRCs and two sister clades of NRC-S (*16–18*). This lineage emerged more than 100 million years ago and extensively diversified in asterid plants, particularly in the Solanaceae, where they form complex immune receptor networks that confer resistance against a wide range of pathogens and pests (*17–20*). Upon activation by their cognate NRC-S partners, NRC helpers oligomerize into homohexameric resistosomes that function as calcium-permeable channels (*11, 21*). Although the cryo-EM structures of three hexameric NRCs, NbNRC2a, SlNRC3, and NbNRC4c, have been experimentally resolved, the extent to which these structures define common principles of NRC immune receptor channels remains unknown.

We previously reported that AlphaFold 3 can predict with high confidence the architecture of hexameric NRC resistosomes and related plant NLR assemblies, including accurate modeling of the very N-terminal α1 helix of CC-NLRs—a structurally elusive region that has proven difficult to resolve experimentally (*11*). We reasoned that AlphaFold 3 could be deployed not only for structural prediction, but also as a discovery engine to classify NLRs into architectural categories and flag candidates with atypical assemblies (*11, 17, 22*). To avoid manual classification and provide a more scalable, less subjective approach, we developed the Structural Novelty Index (SNI), a quantitative metric that distinguishes unconventional from canonical protein complexes based on prior knowledge. Here, we tested this concept by developing and implementing a specific SNI, SNI_*NRC-Hexa*_, aimed at evaluating AlphaFold 3 models of NRC hexamers.

### SNI can distinguish canonical NRC hexamers from unconventional assemblies

To define the SNI_*NRC-Hexa*_ parameters and capture the properties of hexameric NLR resistosomes, we combined features defined by human experts with features generated by the AI system co-scientist (*23*) (**Figure S1; Data S1**). We retained 11 parameters that assess model confidences and inter-protomer interfaces (ipTM_*LCB*_, **∑**_*CONTACTS*_, BSA_*INTER-PROTO*_), resistosome ring symmetry (D_*APEX*_, σ_*θ ROT*_, S_*PROTO*_), distance between MHD and P-loop motifs (D_*MHD-P*_), CC-domain hydrophobicity (H_*ABS*_), N-terminal α1-helix length (L_*APEX*_), angle (φ_*APEX*_), and amphipathic moment (µ_*H*_) (**Figure S1; Data S1**).

We benchmarked SNI_*NRC-Hexa*_, based on the prior knowledge that NRC helper proteins assemble into hexameric resistosomes, but their phylogenetically related NRC-S fail to do so (*17, 24, 25*). To this end, we put together a benchmark dataset consisting of i) known hexameric NRCs: NbNRC2a, SlNRC3, and NbNRC4c and their orthologs from the Solanaceae (nightshade) family, and ii) NRC-S proteins selected from across the phylogenetic clade (*14, 26, 27*). We used AlphaFold 3 to model this set of 18 proteins as hexamers with 25 oleic acids as a stand-in for the plasma membrane, generating three replicates per sequence. All structural predictions of the nine helper proteins aligned with their respective cryo-EM structures with RMSDs below 1 Å (**Figure S2**), confirming the high accuracy of AlphaFold 3 in modeling NRC resistosomes.

Next, we computed the SNI_*NRC-Hexa*_ parameters for the 18 structural models and performed hierarchical clustering based on the SNI_*NRC-Hexa*_ parameters **(Figure 1A**). This resulted in two distinct clusters clearly separating the NRCs from NRC-S proteins. We conclude that SNI_*NRC-Hexa*_ can distinguish AlphaFold 3 models of NRC hexamers from unconventional assemblies.

**Figure 1.**
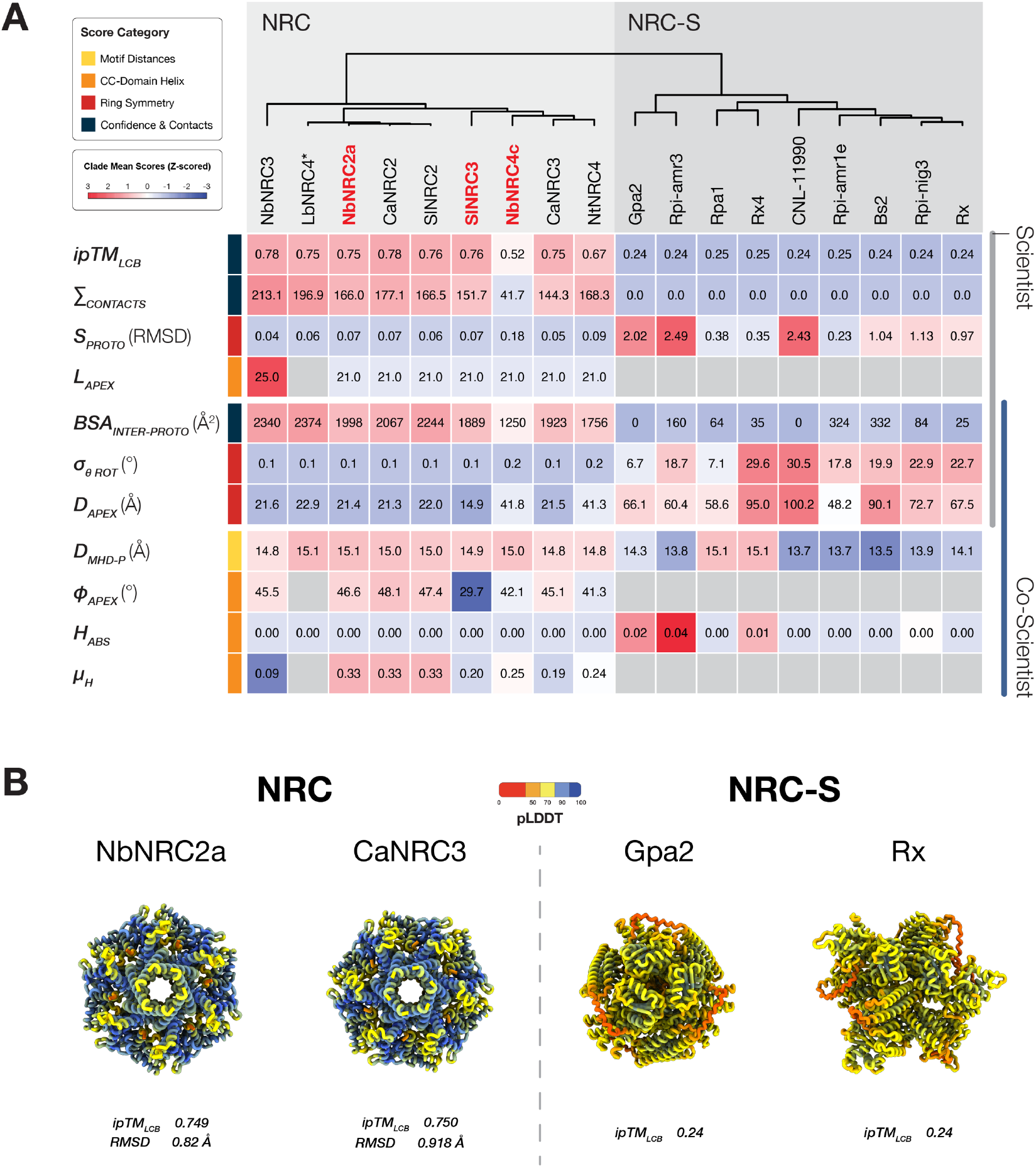
The Structural Novelty Index discriminates between NRC helper vs. sensor predictions. (**A**) Quantification of structural divergence via the tailored Structural Novelty Index (SNI_*NRC-Hexa*_) for hexameric NRC resistosomes. The parameters assess model confidences and inter-protomer interfaces (ipTM_*LCB*_, ∑_*CONTACTS*_, BSA_*INTER-PROTO*_), resistosome ring symmetry (D_*APEX*_, σ_*θ ROT*_, S_*PROTO*_), MHD-P-loop motif distance (D_*MHD-P*_), hydrophobicity of CC-domain (H_*ABS*_), and N-terminal α1-helix length (L_*APEX*_), angle (φ_*APEX*_), and amphipathic moment (µ_*H*_). Source of the parameters is indicated on the right. SNI_*NRC-Hexa*_ calculated for three NRCs with experimentally resolved resistosome structures (NRC2, NRC3, and NRC4c; highlighted red) and two orthologs from each phylogenetic clade. Nine NRC-S NLRs were used as outgroups. Hierarchical clustering of z-score normalized SNI_*NRC-Hexa*_ values for the 18 predicted hexamers show a clear distinction between NRC resistosomes and NRC-S hexamers. LbNRC4 and NRC-S, lacking a detectable MADA motif, could not be scored for α1-helix-dependent metrics. In NbNRC4c and NtNRC4, the α1 helix collapsed inward despite the NB-ARC domain forming a high-confidence resistosome ring (**Figure S3**), leading to higher D_*APEX*_ values. All proteins were modeled in three replicates with 25 oleic acids as a stand-in for the plasma membrane. Nb: *Nicotiana benthamiana*, Nt: *Nicotiana tabacum*, Sl: *Solanum lycopersicum*, Ca: *Capsicum annuum*, Lb: *Lycium barbarum*. (**B**) AlphaFold3-predicted hexameric resistosome complexes for representative helper and sensor NLRs. While helper NLR predictions showed resistosome-like ring structures, sensor NLRs were predicted with low confidence and no meaningful structure. RMSD values for helper structures are against their respective cryo-EM structures (NbNRC2a: 9FP6; SlNRC3: 9RI9). Structures were aligned with the ChimeraX matchmaker command (*28*). Pruned RMSD scores are reported (model with seed = 1 is visualized).

### SNI predicts NRC proteins with unconventional assemblies

To identify unconventional NRC resistosomes, we expanded our analysis to all available NRC protein sequences in the Solanaceae family. We reannotated 346 Solanaceae genomes from 85 species and extracted 197,834 NLR proteins from a total of 15,079,126 proteins (*29*). From this dataset, 2,658 sequences belonged to the well-defined NRC phylogenetic clade, which we further reduced to 637 non-redundant sequences spread over 18 distinct phylogenetic clades (**Data S3**). We used AlphaFold 3 to model these NRC sequences as hexamers, in three replicates and with 25 oleic acid molecules as a proxy for the plasma membrane (*30*). We calculated SNI_*NRC-Hexa*_ for all 637 predicted structures (**Figure S4; Data S4**). To minimize intra-clade variability, we computed a penalized mean score for each NRC clade (**Data S5**). The resulting matrix was normalized using Z-scores, followed by hierarchical clustering to identify divergent NRC clades based on SNI_*NRC-Hexa*_.

The NRC2, NRC3, and NRC4_other_ phylogenetic clades—each containing sequences with empirical hexameric structures—formed a core cluster encompassing 13 of the 18 NRC clades. These clades generally exhibited high mean ipTM scores and similar resistosome geometry profiles (**Figure 2A**). The remaining five clades diverged from the core cluster for several parameters. Among these, NRC4u and NRC7 showed the lowest mean ipTM scores (**Figure 2A; Data S5**) and displayed a markedly reduced number of inter-protomer contacts within these clades (**Figure 2B**). Inspection of representative models from the two most divergent clades with low ipTM scores (NRC4u and NRC7) revealed contrasting patterns. NRC4u clade sequences did not exhibit resistosome-like assemblies (**Figure 2C**). In contrast, despite similarly low ipTM scores, NRC7 clade models exhibited resistosome-like assemblies (**Figure 2C**). Notably, members of the adjacent NRC6 clade—closely related to NRC7 both phylogenetically and by hierarchical clustering— generally modeled with higher confidence and displayed canonical hexameric resistosome architectures, albeit with an inwardly collapsed α1 helix (**Figure 2B & 2C**).

**Figure 2.**
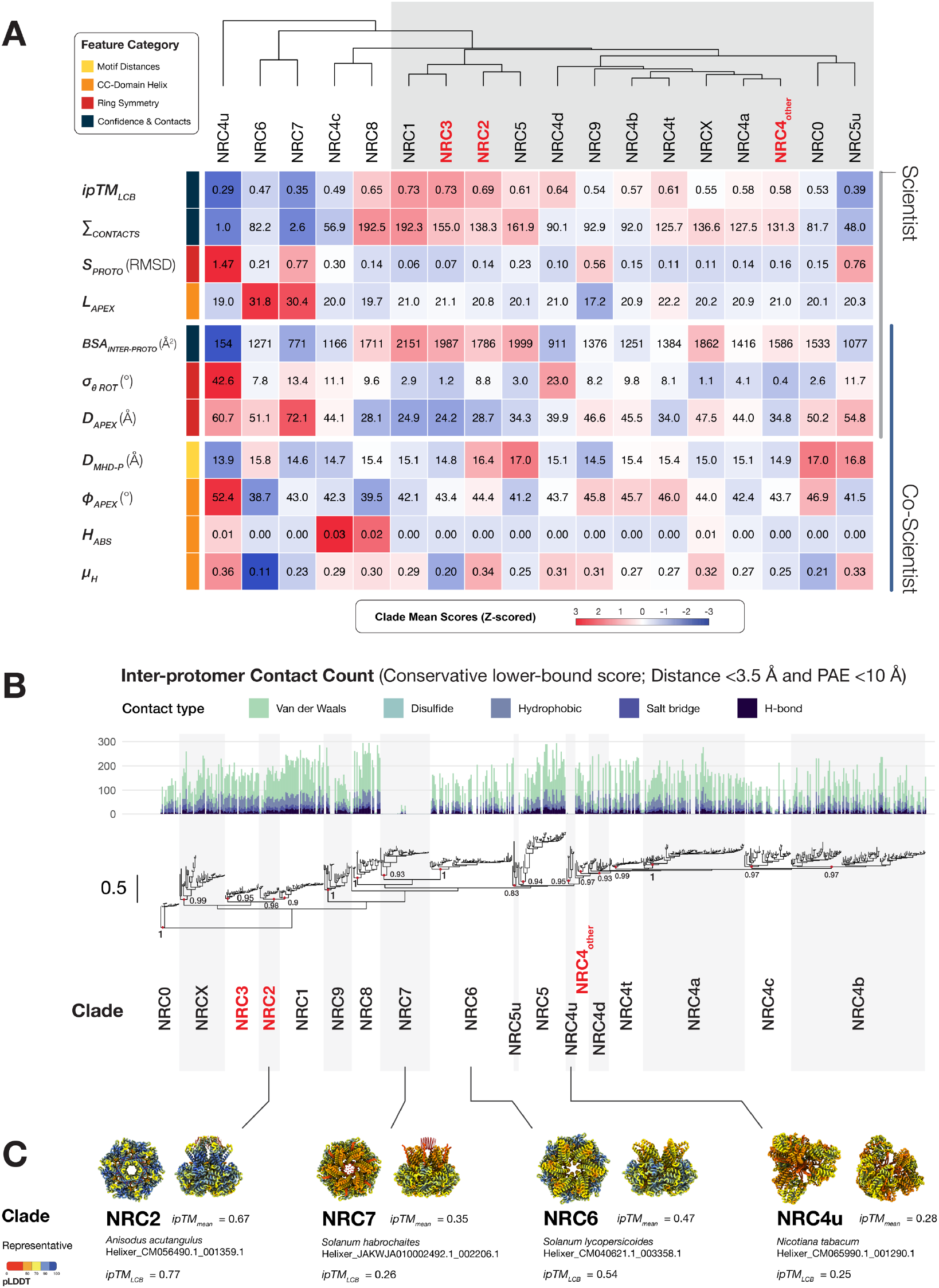
SNI predicts NRC proteins with unconventional assemblies. (**A**) A hierarchical clustering of NRC clades based on the specialized SNI_*NRC-Hexa*_. This framework uses a combination of scientist-defined and AI-generated parameters to prioritize assemblies that deviate from canonical architectures. The parameters assess model confidences and inter-protomer interfaces (ipTM_*LCB*_, ∑_*CONTACTS*_, BSA_*INTER-PROTO*_), resistosome ring symmetry (D_*APEX*_, σ_*θ ROT*_, S_*PROTO*_), MHD-P-loop motifs distance (D_*MHD-P*_), hydrophobicity of CC-domain (H_*ABS*_), and N-terminal α1-helix length (L_*APEX*_), angle (φ_*APEX*_), and amphipathic moment (µ_*H*_). Values are Z-scored by clade mean, with red indicating higher and blue indicating lower relative scores. Hierarchical clustering was applied to NRC clades. NRC clades with empirical structures form a core cluster (highlighted grey) with similar resistosome geometry and confidence profile. Five clades clustered outside of the core cluster with NRC4u and NRC7 having the lowest confidence scores (ipTM_*LCB*_). NRC clades with empirical structures are in highlighted red. (**B**) Stacked bar chart of inter-protomer contact counts for each resistosome, shown on the phylogenetic tree of NB-ARC sequences from 637 NRC sequences. Numbers next to the nodes indicate bootstrap values. Contacts classified by interaction type: hydrogen bonds, salt bridges, hydrophobic interactions, disulfide bonds, and van der Waals contacts. NRC7 and NRC4u clade NLRs show very low contact numbers compared to other NRC clades. (**C**) Representative structures from NRC2, NRC4u, NRC6, and NRC7 clades (seed = 1, with 25 oleic acids as a stand-in for the plasma membrane). AlphaFold 3 failed to predict resistosome-like structures for NRC4u clade NLRs.

### Activated NRC7 proteins form an undecameric (11-mer) resistosome

SNI_*NRC-Hexa*_ analysis identified NRC7 as the largest clade predicted to be structurally unconventional. We therefore hypothesized that NRC7 proteins assemble into a complex distinct from the canonical hexameric NRC resistosome. To test whether potato NRC7 (*Solanum tuberosum*; StNRC7) can trigger immune cell death, we introduced a point mutation in the MHD motif (D498V, hereafter StNRC7^DV^), a substitution known to confer autoactivity (*31*)(**Figure 3A and 3B**). Transient expression of StNRC7^DV^, but not wild-type StNRC7, in leaves of the model plant *Nicotiana benthamiana* induced a cell death response comparable to that triggered by the previously described autoactive SlNRC3^DV^ (**Figure 3C**) (*14*).

**Figure 3.**
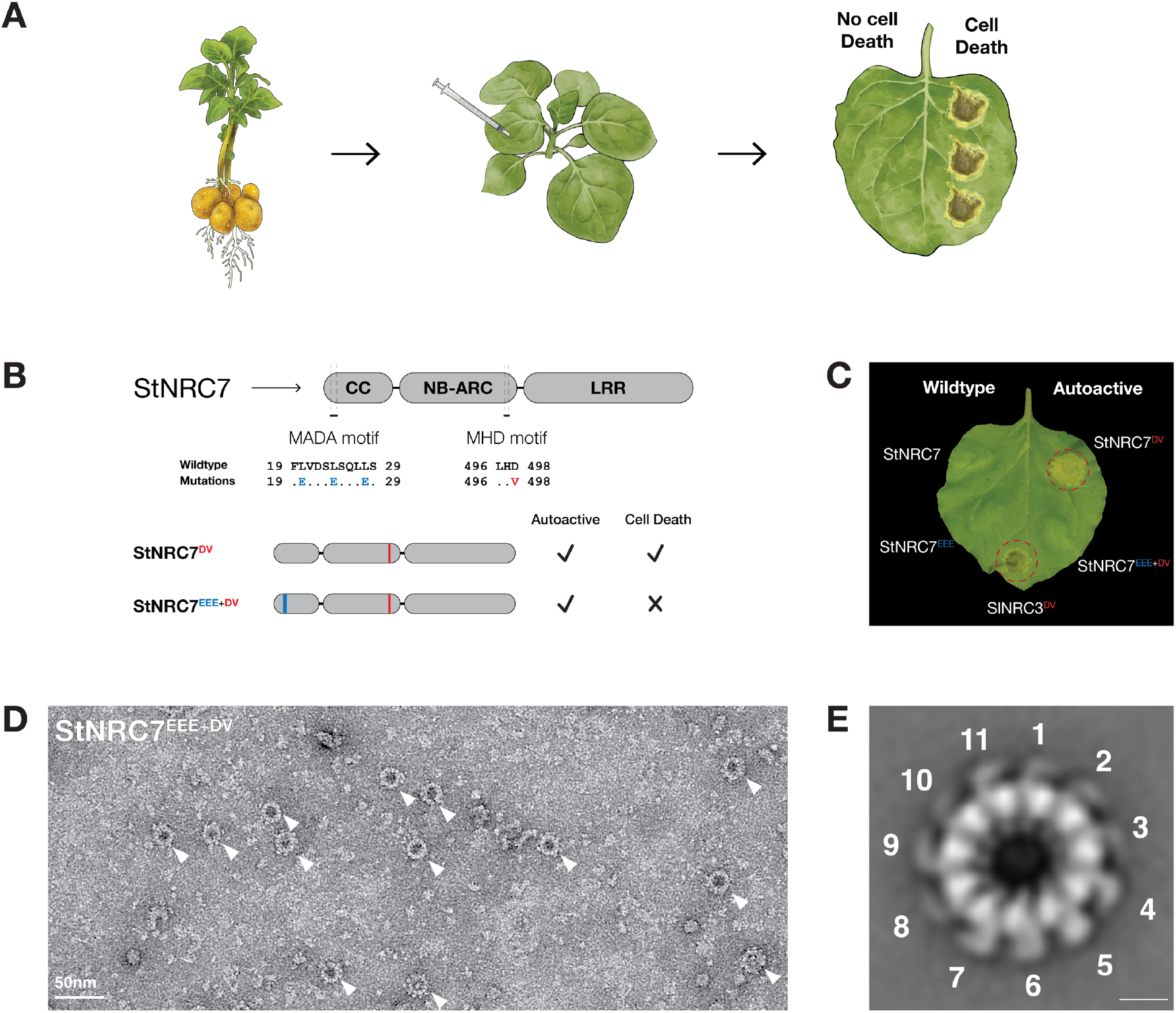
Activated NRC7 proteins form 11-mer resistosome complexes. (**A**) Schematic of the agroinfiltration assay used to test NRC7-triggered cell death. Potato NRC7 (*Solanum tuberosum*; StNRC7) was cloned into a binary expression vector and transiently expressed in *N. benthamiana* leaves by Agrobacterium-mediated transformation. (**B**) Schematic of StNRC7 showing a tripartite domain structure. The positions of the MADA and MHD motifs, and the mutant variants generated to produce an activated protein that no longer induces cell death are indicated. (**C**) Cell death induced by the autoactive StNRC7^DV^ is abolished by additional mutations in the MADA motif (StNRC7^EEE+DV^). Agrobacterium strains carrying the indicated constructs were infiltrated into leaves of 4-week-old *N. benthamiana* plants at OD600 of 0.3. SlNRC3^DV^ was used as a positive control for cell death. Spots with cell death have red circles. Photographs were taken 5 days post infiltration. (**D**) Negative-stain electron micrograph of StNRC7^EEE+DV^ showing large ring-shaped particles (white arrows). (**E**) Negative-stain 2D class average of StNRC7^EEE+DV^ revealing an 11-protomer assembly. Scale bar, 60 Å.

To enable purification of the autoactive StNRC7^DV^ mutant for structural analysis, we introduced additional mutations in the MADA motif (L20E, L24E, L28E) of the N-terminal α1 helix, which have previously been used to abolish NRC-mediated cell death without disrupting resistosome assembly (*24, 32*) (**Figure 3B and 3C**). We expressed StNRC7^EEE+DV^ protein in leaves of *N. benthamiana* and subjected it to a previously described immunoprecipitation-electron microscopy (IP-EM) workflow (*26*). We harvested leaves 2 days after infiltration, and purified StNRC7^EEE+DV^ by strep-tag affinity purification before assessment by negative-stain electron microscopy (nsEM). The nsEM micrographs revealed distinct ring-shaped particles of ∼250 Å in diameter, substantially larger than the ∼150 Å diameter expected for a canonical hexameric resistosome (**Figure 3D**). Two-dimensional (2D) classification resolved a clear ring-shaped structure with features consistent with a resistosome. Counting the peripheral densities, which likely correspond to the LRR domains, identified the complex as an atypical assembly composed of 11 protomers (**Figure 3D & 3E**).

### Additional members of the NRC7 clade form 11-mer assemblies

To determine whether the atypical 11-protomer architecture of the StNRC7 resistosome is a broader feature of the NRC7 clade, we examined NRC7 orthologs from tomato (*Solanum lycopersicum*, SlNRC7) and *N. benthamiana* (NbNRC7), which represent distinct branches of this clade (**Figure 4**). We expressed and purified MADA-mutated autoactive variants of SlNRC7 and NbNRC7 (SlNRC7^EEE+DV^ and NbNRC7^EEE+DV^, respectively) from *N. benthamiana* leaves and analyzed them by negative-stain electron microscopy. Like StNRC7, both SlNRC7 and NbNRC7 formed resistosome-like 11-mer assemblies (**Figure 4**). These findings indicate that 11-protomer resistosomes are a conserved structural feature of the NRC7 clade, in marked contrast to the canonical hexameric resistosomes formed by NbNRC2a, SlNRC3, and NbNRC4c.

**Figure 4.**
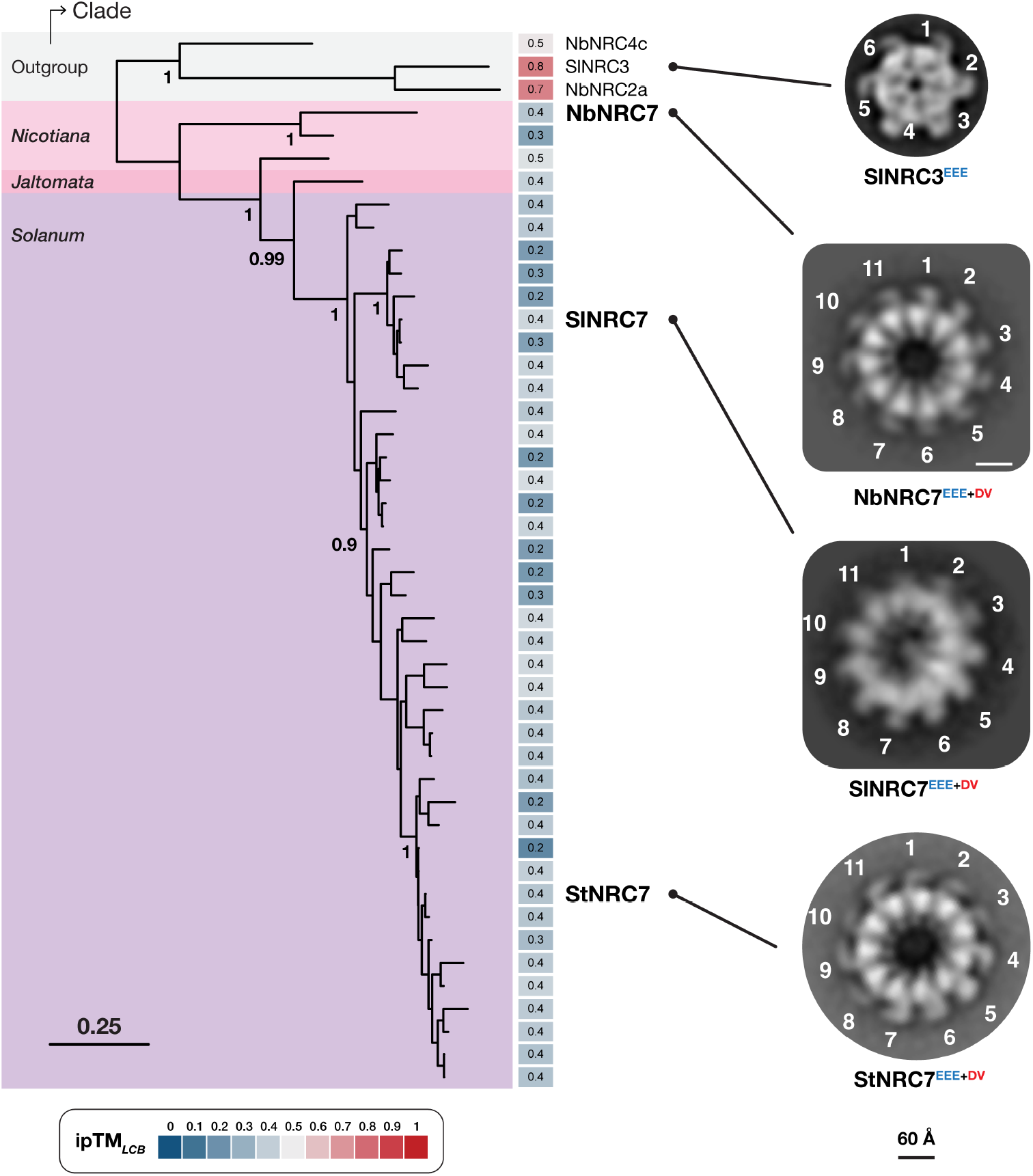
Additional NRC7 clade proteins form 11-mer resistosome assemblies. (**A**) Phylogeny of the NRC7 clade, with NRC2, NRC3 and NRC4 included as outgroups. Outgroups and genera are coloured accordingly. The scale bar represents 0.1 substitutions per site, and numbers on branches indicate bootstrap support values. The ipTM score for each AlphaFold3 hexamer model (seed = 1) is shown at the tips of the phylogeny (**B**) Negative-stain 2D class averages of StNRC7^EEE+DV^ (from **Figure 3E**), SlNRC7^EEE+DV^, and NbNRC7^EEE+DV^, showing similar 11-mer oligomeric states. The hexameric resistosome SlNRC3^EEE^, activated by the NRC-S protein Rx in response to *Potato virus X* coat protein (CP) (*14*), is included for comparison as an outgroup assembly. Scale bar, 60 Å, in all images.

### Co-scientist complements human-defined SNI parameters

The SNI framework provides an objective pipeline for discovering atypical protein assemblies and should be scalable across a broad range of protein complexes (**Figure 5**). Incorporating the AI system co-scientist (*23*) may further enhance this scalability and accelerate analysis. Human experts defined eight SNI parameters, whereas co-scientist proposed seven, four of which overlapped with the expert-defined set and three of which were unique (**Figure 1A & S1**). To assess how informative the analysis would have been using AI-generated features alone, we directly compared the human-defined and co-scientist-derived parameter sets and performed hierarchical clustering with each set independently (**Figure S5 & S6**). In the benchmark analysis, neither parameter set matched the performance of the combined SNI_*NRC-Hexa*_, which perfectly separated NRC and NRC-S models into distinct clusters (**Figure 1A & S5**). Using either parameter set alone led to one misclassification: NbNRC4c with the human-defined parameters and Rpa1 with the co-scientist-derived parameters. In the full dataset of 637 NRC proteins, both analyses identified NRC7 as atypical (**Figure S6**). However, whereas the human-defined parameters placed only three proteins outside the “hexamer” cluster containing NRC2, NRC3, and NRC4, the co-scientist-derived analysis provided greater granularity and identified four clades as unconventional (**Figure S6**). Although future experimental work will be needed to determine how many of these candidates are truly non-canonical, these results indicate that the co-scientist-derived parameters alone would have been sufficient to flag NRC7 as distinct from the core hexameric resistosome architecture.

**Figure 5.**
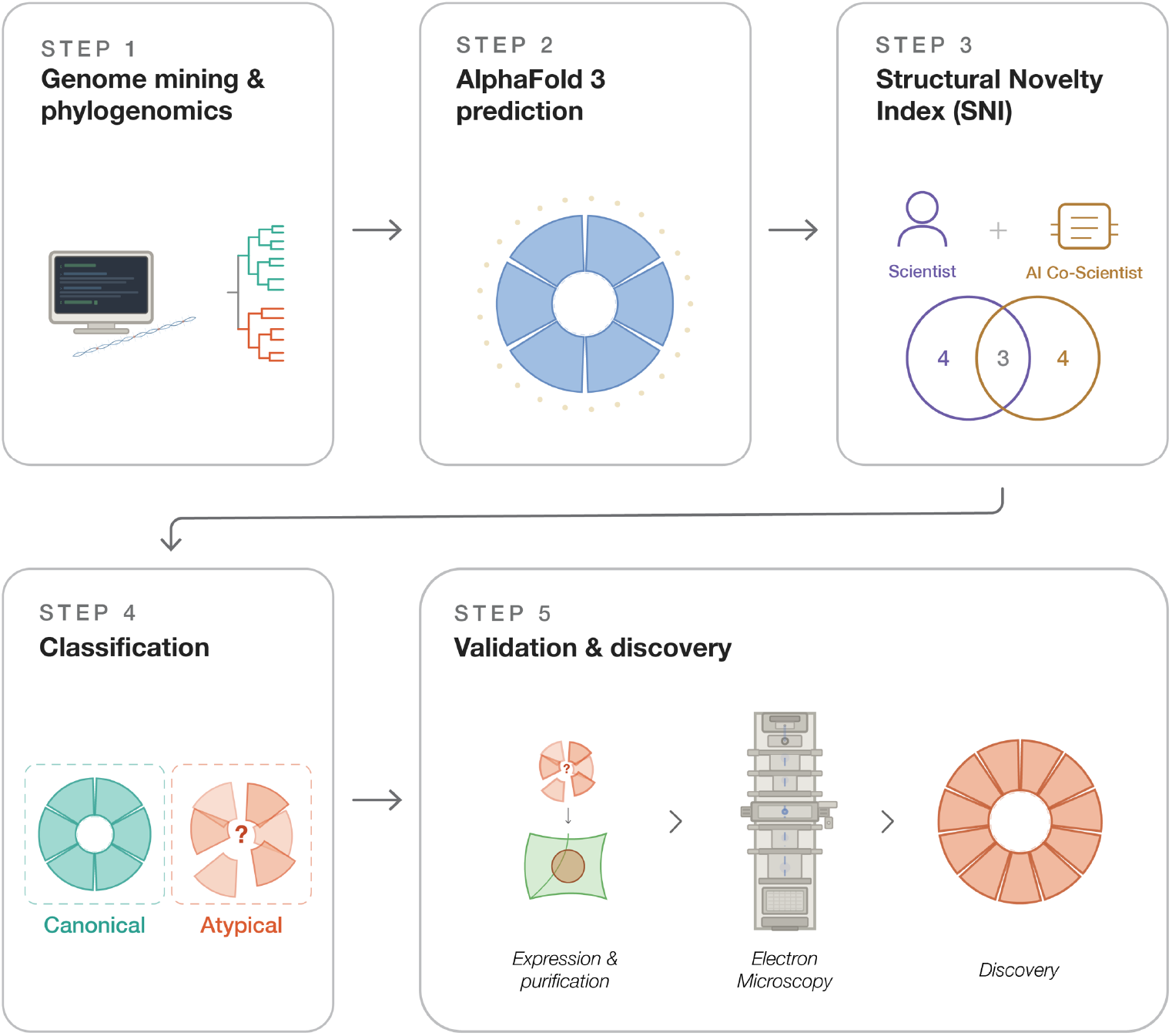
AI-guided discovery of unconventional protein assemblies. Schematic overview of the five-step pipeline for identification of atypical macromolecular architectures, applied here to the NRC family of plant immune receptors. Step 1: Large-scale genome mining and phylogenomics to extract and curate a non-redundant set of candidate protein sequences. Step 2: AlphaFold 3 modeling of all candidates in a defined reference assembly state. Step 3: A Structural Novelty Index (SNI) is constructed by integrating parameters from both human experts and the AI co-scientist system to quantitatively distinguish canonical from unconventional assemblies. Step 4: SNI-based clustering to classify candidates as canonical or atypical, flagging outliers that deviate from the expected assembly geometry. Step 5: Top candidates are subjected to experimental validation through recombinant expression, affinity purification, and electron microscopy to enable the discovery of previously unrecognized assembly architectures.

## Discussion

This work shows how integrating AI-driven structural prediction with electron microscopy can accelerate the classification of complex protein families and reveal atypical macromolecular assemblies. To avoid manual classification and establish a more scalable, less subjective approach, we developed the Structural Novelty Index (SNI), a quantitative metric that distinguishes unconventional from canonical protein complexes on the basis of prior knowledge. This AI-guided framework enables the systematic discovery of previously unrecognized assembly architectures and provides a general strategy for identifying atypical protein assemblies (**Figure 5**).

Applying a bespoke SNI to AlphaFold 3 predictions spanning the expanded phylogenetic diversity of the NRC family uncovered previously unrecognized structural variation within this well-studied class of plant immune receptors (**Figures 3 and 4**). Because cryo-EM analyses had shown that three NRC family members form hexameric resistosomes (*11, 12, 14*), it would have been natural to infer that the same architecture extends across the family. Instead, SNI highlighted candidate atypical NRCs and guided the experiments that uncovered the unexpected 11-mer resistosome stoichiometry of NRC7 proteins.

Although predictive workflows often prioritize high-confidence models for structural interpretation, our findings show that low-confidence predictions can also be informative by flagging candidates that deviate from canonical assemblies and warrant experimental investigation. This emphasis on successful predictions can create a form of survivorship bias, whereby low-confidence or failed models are overlooked even though they may carry biological meaning. NRC7 clade members consistently failed to produce confident hexameric models, and modeling across alternative oligomeric states from 5-mers to 11-mers likewise did not yield confident assemblies (**Figure S7**). Rather than treating such models as disposable failures, they can highlight outliers to prioritize candidates for experimental structural analysis, thereby countering this bias in conventional modeling workflows. More broadly, SNI extends beyond low-confidence predictions by quantitatively and objectively flagging structural models that diverge from canonical assemblies across multiple parameters.

The undecameric NRC7 resistosome was unexpected. So far, plant CC-NLR resistosomes that have been functionally characterized appear to act as calcium-permeable channels (*11, 21, 33*). Whether the NRC7 11-mer retains this activity or whether its larger assembly state channels other ions or small molecules is an open question. Future work will determine the properties of this complex and test the extent to which the structural expansion of the NRC7 resistosome is linked to functional innovation.

Co-scientist independently recovered several structural parameters identified by human experts and contributed additional features that enhanced the granularity of the analysis. Together, these findings point to a near-term future in which agentic AI systems complement expert researchers to accelerate the development of bespoke SNI frameworks for diverse protein assemblies. This complementarity could also broaden access to structural discovery by enabling teams without extensive in-house structural biology expertise to generate and benchmark useful parameter sets more efficiently.

## Author contributions

Conceptualization: D.L., S.K., A.T., B.S.

Methodology: A.T., B.S., D.L.

Data curation: A.T., B.S., Y.S., D.L.

Formal analysis: A.T., B.S., D.L.

Investigation: A.T., B.S., D.L., L.-M.R., J.M. A., M.G., I.M., M.P.C., M.S., A.H.

Resources: S.G., D.M., Y.S., M.P.C., J.K.

Writing—original draft: A.T., S.K., D.L., B.S.

Writing—review and editing: A.T., S.K., D.L., B.S., M.P.C., D.M., J.K., Y.S.

Visualization: A.T., B.S., D.L.

Supervision: S.K., D.L., B.S.

Project administration: S.K., D.L.

Funding acquisition: S.K.

## Data and materials availability

Additional supplementary material and scripts are available at https://github.com/amiralito/SolNRCH_foldome. Genome annotations and sequence data are available at https://doi.org/10.5281/zenodo.19855162 (*29*). Predicted structures are available at https://doi.org/10.5281/zenodo.19860917 (*30*).

## Acknowledgements

We thank Andrés Posbeyikian (TSL) for comments and input on the analysis, Jake Richardson for support with electron microscopy, Hsuan Pai for her plant schematic drawings in figure 3, and all TSL support staff and Horticultural Services for preparing and providing plants and media. D.L. thanks Nigel Tufnel for inspiration.

