## Supplementary Materials for "AI-guided discovery of atypical protein assemblies"

### Materials and methods

#### Phylogenetic analysis

From the NLRtracker output of 346 Solanaceae proteomes, we extracted a total of 197,834 NLR proteins (**Data S1 and S2**) (*1*)[ref for dataset]. Based on NLRtracker domain annotations, we retained 169,434 NLRs with the following domain architectures: ″CNL″, ″CNLO″, ″CN″, ″OCNL″, ″CONL″, ″NL″, ″NLO″, ″ONL″, ″BCNL″, ″BCNLO″, ″BNL″, ″BBNL″, ″BCN″, ″BCCNL″, ″BBCNL″, ″BNLO″, ″BOCNL″, ″BBCNLO″, ″RNL″, ″TN″, ″TNL″, ″TNLO″, and ″TNLJ″. We deduplicated this set using CD-HIT v4.8.1, resulting in 87,460 unique NLRs (*2*). We then filtered the dataset to retain only sequences with NB-ARC domains between 250 and 400 amino acids, yielding 73,677 NLRs. We aligned the NB-ARC domains from this set with RefPlantNLR NB-ARC domains using FAMSA v2.4.1(*3*) and constructed a phylogenetic tree using FastTree v2.1.11 (*4*).

Using the presence of reference NRCs and the branch with the highest bootstrap values, we identified the NRC clade, which contained 2,658 NLRs. To remove truncated sequences, we excluded proteins with CC domains shorter than 100 amino acids or total protein lengths shorter than 800 amino acids, leaving 2,071 sequences. From this set, we extracted the region spanning the N-terminus to the end of the NB-ARC domain (CC–NB-ARC) for AlphaFold modeling. Before modeling, we further reduced redundancy by clustering sequences at 95% identity using CD-HIT, which yielded a final set of 637 NRC sequences (**Data S2**).

Finally, we reconstructed a phylogenetic tree for this set by aligning NB-ARC domains with reference NRCs using MAFFT v7.526 (*5*, *6*) and inferring a tree with FastTree. Based on the presence of reference NRCs and well-supported divergent branches, we defined 19 NRC subclades. Phylogenetic supporting material are available at https://github.com/amiralito/SolNRCH_foldome.

#### Motif annotation

Conserved sequence motifs in NLR proteins were annotated using NLRexpress with the [--module all] option (*7*). We cross-checked the output against motif annotations from NLRtracker (*1*) and complemented missing entries from each program with the other. We assigned NA values for DMHD–P to NLRs lacking MHD or P-loop annotations in the final structural analysis output.

#### AlphaFold 3 modeling

We used a local installation of AlphaFold 3 (https://github.com/google-deepmind/alphafold3) to model CC–NB-ARC domains of NRCs as hexameric resistosomes with 25 oleic acid molecules as a stand-in for the plasma membrane (8). To efficiently model the large Solanaceae NRC dataset, we decoupled the template and MSA calculation steps from the model inference step using the AlphaFold 3 data pipeline (https://github.com/google-deepmind/alphafold3/blob/main/docs/performance.md#data-pipeline). We first generated templates and MSAs for the 637 NRC sequences and then used the AlphaFold 3 inference pipeline to model them as hexamers with seeds 1, 2, and 3. We analyzed and visualized the resulting models using a custom pipeline. All scripts are available at https://github.com/amiralito/SolNRCH_foldome.

Because A100 80GB GPUs impose a ~5,000-token limit, we used H200 141GB GPUs to model larger NRC7 assemblies. We predicted CC–NB-ARC domains of St- and SlNRC7 as 5–11-mers and visualized pTM and ipTM scores using ggplot2 in R (*9*, *10*). All models are deposited at https://doi.org/10.5281/zenodo.19860917 (*11*).

#### Co-scientist methods

We prompted AI system co-scientist to define a quantitative Structural Novelty Index capable of robustly distinguishing unconventional NRCs from canonical hexameric NRC resistosomes. Co-scientist generated 19 hypotheses, which it evaluated to produce 9 research ideas representing different approaches. After reviewing these ideas, we selected the first for implementation in our SNI pipeline. The full co-scientist prompt and output is provided in Data S6.

#### Structural Novelty Index criteria calculation

The Structural Novelty Index (SNI) aggregates a set of structural descriptors computed on each AlphaFold 3 hexameric model. All metrics were calculated per structure (per seed replicate) and then aggregated across three replicates per NLR. Full implementation details — including equations, software versions, residue-level thresholds, and the contact-classification rules — are provided in the accompanying repository (https://github.com/amiralito/SolNRCH_foldome).

Interface confidence was quantified using the AlphaFold 3 ipTM score, extracted from the summary confidence JSON. Heavy-atom contacts between ring-adjacent protomers ($\sum_{CONTACTS}$) were enumerated with a spatial index, filtered by predicted aligned error (PAE), and classified into hydrogen bonds, salt bridges, hydrophobic contacts, disulfide bonds, and van der Waals contacts;

total and per-type contact counts were reported per structure. The buried surface area of each inter-protomer interface ($BSA_{INTER\text{-}PROTO}$) was computed from SASA values with FreeSASA (*12*, *13*). Ring geometry was captured with two complementary metrics: the standard deviation of inter-protomer angles at the ring centroid ($\sigma\theta_{ROT}$), and a radial-symmetry metric describing the spread of protomer centroid-to-ring-centre distances ($S_{PROTO}$). Both return zero for a perfectly symmetric ring. The pore aperture ($D_{APEX}$) was computed as the mean pairwise distance between the N-terminal Cα atoms of protomers; when an HMM-based MADA hit was available, the reference N-terminal was corrected to the methionine at the MADA alignment start (or the closest upstream methionine) to discount spurious gene-model extensions (*14*, *15*).

Four descriptors of the CC domain were computed per protomer: the α1-helix tilt angle relative to the pore axis ($\varphi_{APEX}$), the α1-helix length ($L_{APEX}$), the mean Kyte–Doolittle hydrophobicity of the CC-domain helices ($H_{ABS}$) (*16*), and the Eisenberg amphipathic moment of α1 ($\mu_H$) (*17*). $\varphi_{APEX}$ was only reported for protomers whose α1 segment passed a linearity threshold, excluding kinked or disordered helices. A single motif-distance metric ($D_{MHD\text{-}P}$) — the Euclidean distance between the MHD and P-loop motif centroids — was used as a descriptor of NB-ARC compactness. α1-helix linearity and mean α1 pLDDT were tracked as quality indicators.

Cross-replicate aggregation penalised metrics by their reproducibility across the three seed replicates. For "higher-is-better" metrics (ipTM, contacts, BSA) the mean was reduced by a fixed multiple of the per-mean standard error; for "lower-is-better" metrics ($\sigma\theta_{ROT}$, $S_{PROTO}$, $D_{APEX}$) the mean was increased by the same quantity. With $n = 3$ replicates this penalty functions as a ranking adjustment rather than a formal confidence interval (see detailed methods). Non-penalised metrics ($\varphi_{APEX}$, $L_{APEX}$, $H_{ABS}$, $\mu_H$, $D_{MHD\text{-}P}$) were reported as mean ± standard deviation. Analyses used BioPython, SciPy, NumPy, HMMER and FreeSASA; the full stack, versioning, and command-line parameters are documented alongside the analysis scripts at https://github.com/amiralito/SolNRCH_foldome.

#### Plant growth conditions

Wild type *Nicotiana benthamiana* plants were grown in a glasshouse at 24 °C day/22 °C night, 45 – 65 % humidity, and with Levingstons F2 compost as substrate. The *N. benthamiana nrc234* mutant line was previously described (*18*).

#### Plasmid construction

Cloning was performed using the Golden Gate Modular Cloning (MoClo) (*19*) kit and the MoClo plant parts kit (*20*). Cloning design and sequence analysis were performed using Geneious Prime (v2025.0.2; https://www.geneious.com). All plasmids generated and used in this study are described in **Supplementary Data S7**.

#### Transient gene expression

Transient gene expression for protein pulldown or cell death assays was performed in 4-week-old *N. benthamiana* plants via agroinfiltration. *Agrobacterium tumefaciens* GV3101 pMP90 transformed with binary vectors were grown at 28 °C on LB plates containing appropriate antibiotics. Cells were collected from the plate, washed and resuspended in infiltration buffer (10 mM MgCl2, 10 mM MES-KOH pH 5.6, 200 µM acetosyringone). OD600 (see **Supplementary Data S7**) was adjusted after incubation of agrobacteria in the dark.

#### Protein purification from plant extracts

For protein isolation, infiltrated leaf tissue was harvested 2 days after infiltration and flash frozen in liquid nitrogen. The tissue was homogenized using mortar and pestle and 20ml of extraction buffer (FLAG purification: 50 mM Tris-HCl pH 8, 150 mM NaCl, 10% (v/v) glycerol, 10 mM DTT, 0.4% (v/v) IGEPAL and cOmplete EDTA-free protease inhibitor (1 tablet per 50mL of extraction buffer). Strep purification: 50 mM Tris-HCl pH 8, 150 mM NaCl, 10% (v/v) glycerol, 10 mM DTT, 0.4% (v/v) IGEPAL, 5% BioLock (IBA Lifesciences) and cOmplete EDTA-free protease inhibitor (Roche) (1 tablet per 50mL of extraction buffer)) per 5g of plant tissue was added. After vortexing and incubation on ice, the plant extract was centrifuged at 5000x *g* using a swing out centrifuge. The supernatant was filtered through several layers of miracloth (475855-1R), before centrifugation at 35,000x *g* in a fixed angle centrifuge. The resulting supernatant was filtered through several layers of miracloth before the addition of ~150uL of packed M2 anti-FLAG beads (SlNRC7) or Strep-Tactin XT sepharose beads (NbNRC7 and StNRC7). Plant extract and beads were incubated on a rotating wheel at 4 °C for 2h, before collecting the beads in a gravity column. The collected beads were washed with at least 10x 1ml of wash buffer (50 mM Tris-HCl pH 8, 150 mM NaCl, 3% (v/v) glycerol, 2 mM DTT, 0.1% (v/v) IGEPAL) before collecting the beads in an Eppendorf tube. 150uL of elution buffer (160 ug/mL 3x FLAG peptide in wash buffer for FLAG beads, or 50 mM biotin in wash buffer for Strep-Tactin XT sepharose beads) and incubated for 45 minutes - 1 hour at 4 °C to elute the protein. Up to five elution rounds were collected, combined, and flash frozen in liquid nitrogen. To improve purity, NbNRC7 was further purified via FLAG affinity purification. For this, elutions were pooled, incubated with 150 uL of packed M2 anti-FLAG beads and eluted as above. Elutions were pooled and concentrated on a 100K MWCO Amicon Ultra 0.5 centrifugal concentrator (Millipore) before flash-freezing in liquid nitrogen.

#### Negative stain

Proteins were thawed on ice and grids were made using undiluted sample (NbNRC7 and SlNRC7) or 5-fold diluted sample (StNRC7). Samples were applied to glow-discharged 300-mesh copper grids coated with continuous carbon (Agar Scientific) and stained with 2% (w/v) uranyl acetate in water. Grids were imaged manually on an FEI Talos F200C microscope operating at 200 KeV with a Gatan OneView camera (Gatan). Images (25-50 for each sample) were taken at a nominal magnification of 45,000 x corresponding to a pixel size of 2.87 Å. Micrographs were imported and processed in CryoSPARC (v4.7.1) (*21*). Class averages consist of 243 particles (SlNRC7), 269 particles (NbNRC7) and 426 particles (StNRC7).

#### Supplementary Data

**Data S1.** Structural Novelty Index description of parameters.

**Data S2.** $\text{SNI}_{NRC\text{-}Hexa}$ calculations for the 18 benchmark sequences.

**Data S3**. Sequence and metadata of 637 NRC helpers used for structural modeling.

**Data S4.** $\text{SNI}_{NRC\text{-}Hexa}$ calculations for the 637 NRC helper sequences.

**Data S5.** Averaged $\text{SNI}_{NRC\text{-}Hexa}$ calculations for the 18 NRC clades.

**Data S6.** AI Co-scientist output.

**Data S7.** List of plasmids used in this study.

### Supplementary Figures

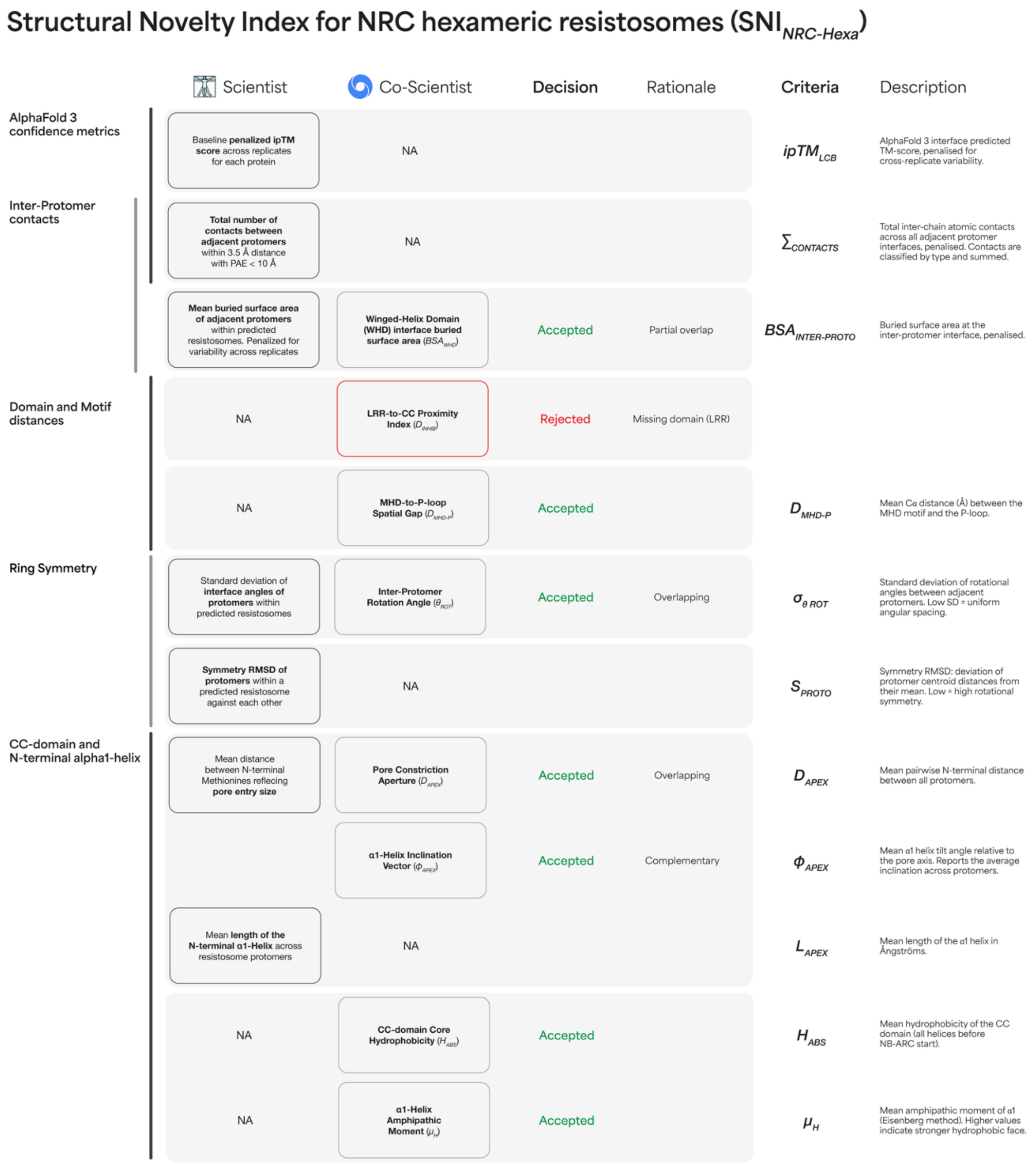


**Figure S1. Structural Novelty Index criteria.**

7 indices were drafted by scientists and 8 indices by AI co-scientist. 11 criteria were chosen based on feasibility and overlap between human-based and AI-based criteria. LRR-to-CC proximity index proposal from AI co-scientist was rejected due to missing LRR domain in resistosome predictions.

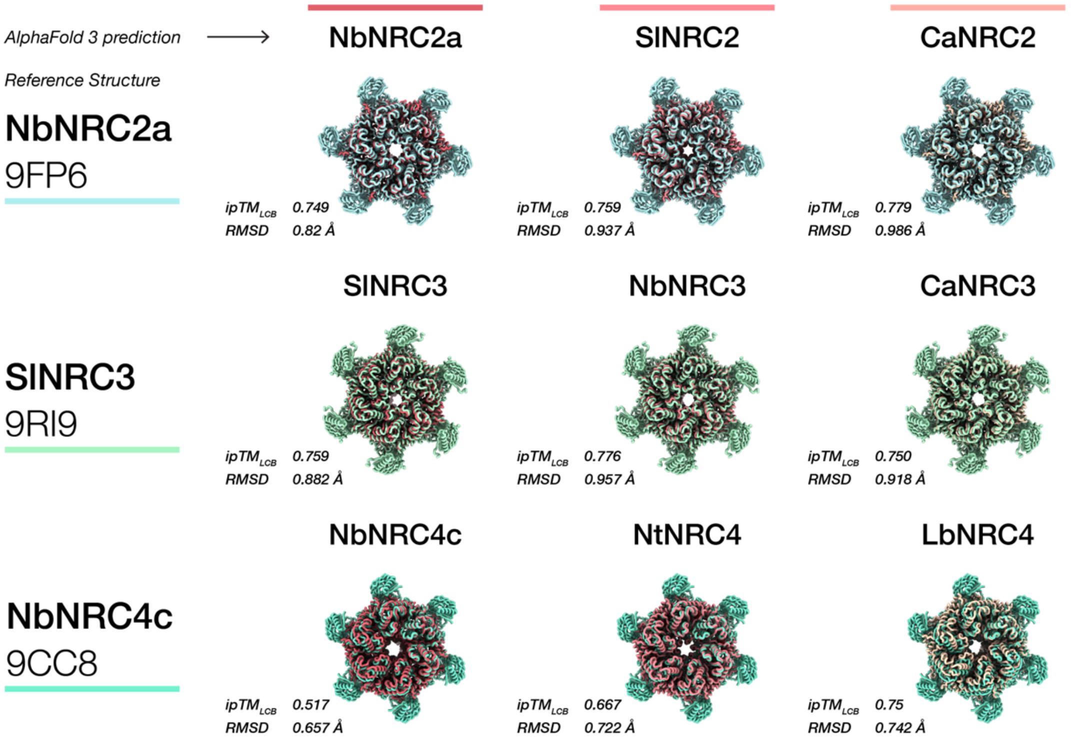


**Figure S2. AlphaFold3 recapitulates hexameric NRC resistosomes across clades.**
AlphaFold3-predicted hexameric resistosome complexes for representatives of the NRC2, NRC3, and NRC4other clades align with high fidelity to their respective experimental reference structures (NbNRC2a: PDB 9FP6; SlNRC3: PDB 9RI9, NbNRC4c: PDB 9CC8), with RMSDs below 1 Å and ipTMLCB scores consistently above 0.5. NRC proteins were modeled in three replicates with 25 oleic acids as a stand-in for the plasma membrane. Structures were aligned with the ChimeraX matchmaker command (24). Pruned RMSD scores are reported (model with seed = 1 is visualized)

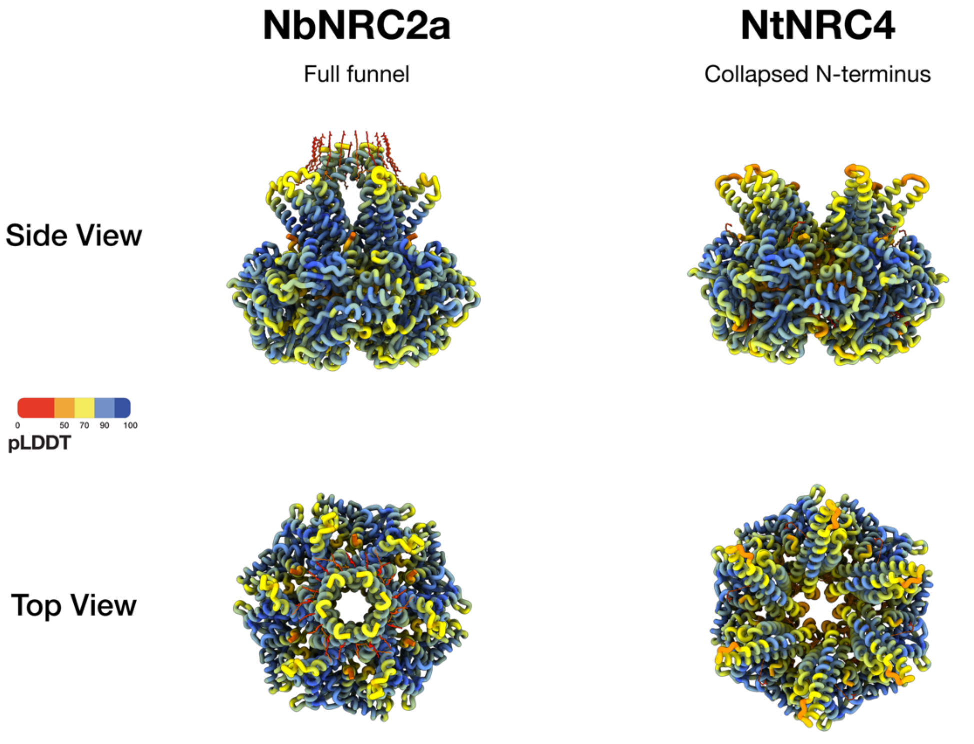


**Figure S3. Resistosome predictions can have inwards collapsed α1 helix.** AlphaFold3-predicted hexameric resistosome complexes for NbNRC2a and NtNRC4. Model with 25 oleic acids as a stand-in for the plasma membrane and seed = 1 is visualized, coloured based on pLDDT. While NbNRC2a hexamer prediction represents a full funnel, NtNRC4 resistosome has an inwards collapsed α1 helix.

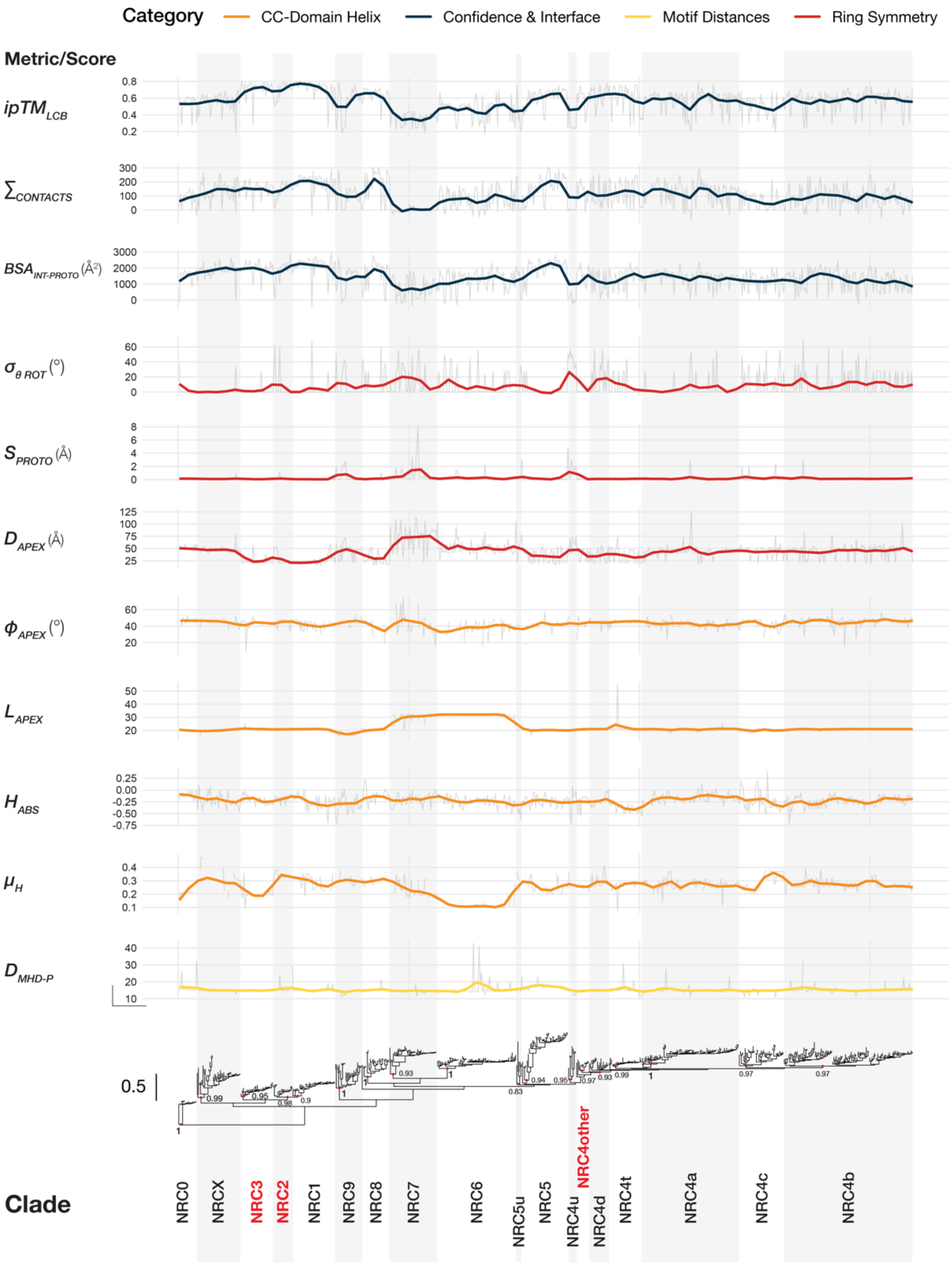


**Figure S4. SNI raw scores per sequence shown on the phylogenetic tree of NB-ARC sequences of 637 NRC sequences.**

NRC clades are annotated based on presence of reference representatives and well-supported branches dividing major clades. Numbers next to nodes indicate bootstrap values. Clades with empirical cryo-EM structures are highlighted red.

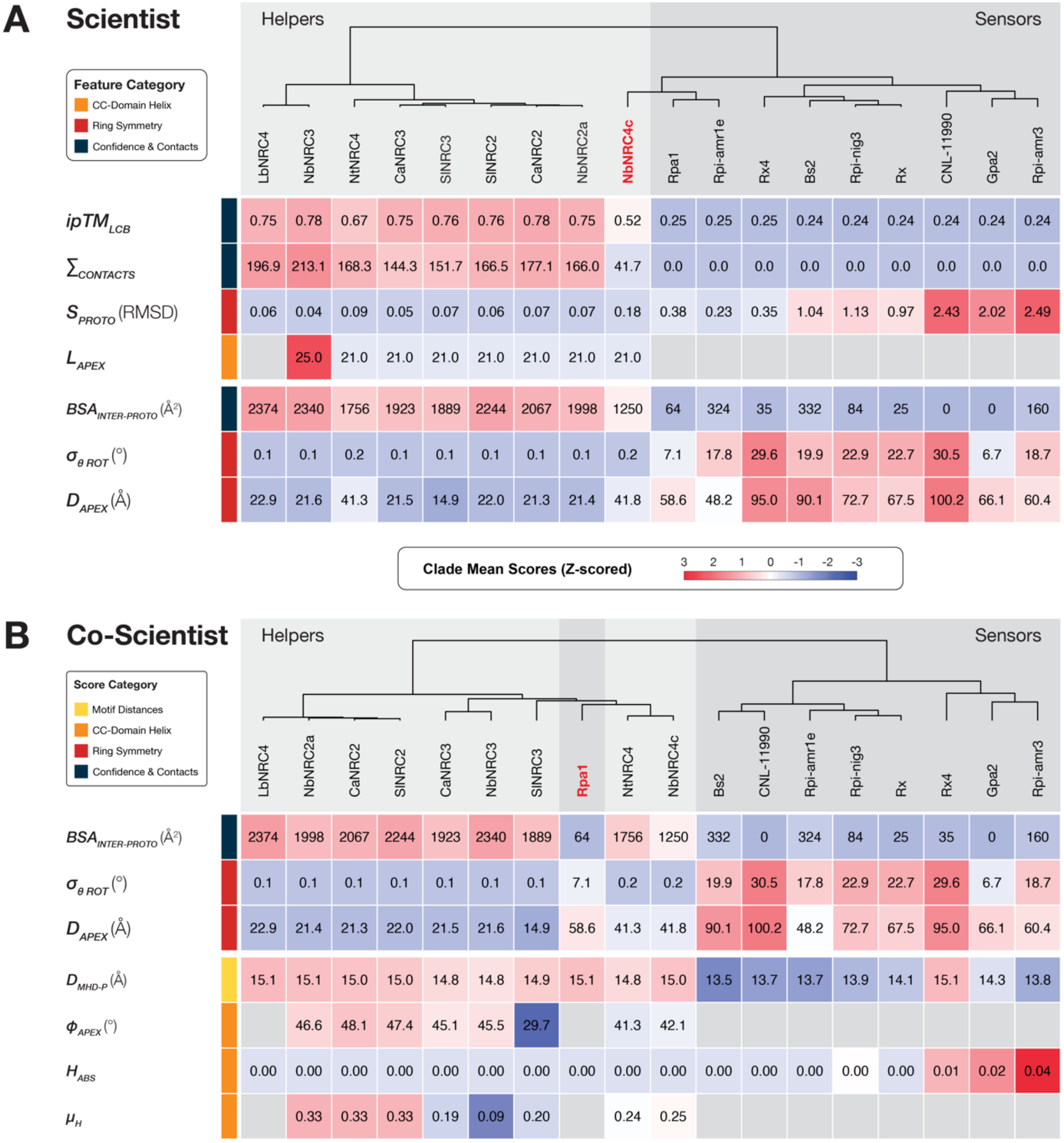


**Figure S5. Scientist and co-scientist generated parameters overlap in distinguishing hexameric predictions helper and sensor NRCs.**

Scientist-only parameters include $ipTM_{LCB}$, $\sum_{CONTACTS}$, $S_{PROTO}$, and $L_{APEX}$. Co-scientist parameters include $D_{MHD\text{-}P}$, $\varphi_{APEX}$, $H_{ABS}$, and $\mu_H$. Parameters shared between both include $BSA_{INTER\text{-}PROTO}$, $\sigma_{\theta\ ROT}$, and $D_{APEX}$. **(A)** In hierarchical clustering of Scientist-only parameters NbNRC4c is misplaced in NRC sensor cluster. **(B)** In hierarchical clustering of co-scientist parameters Rpa1 is misplaced in the middle of the NRC helper clusters.

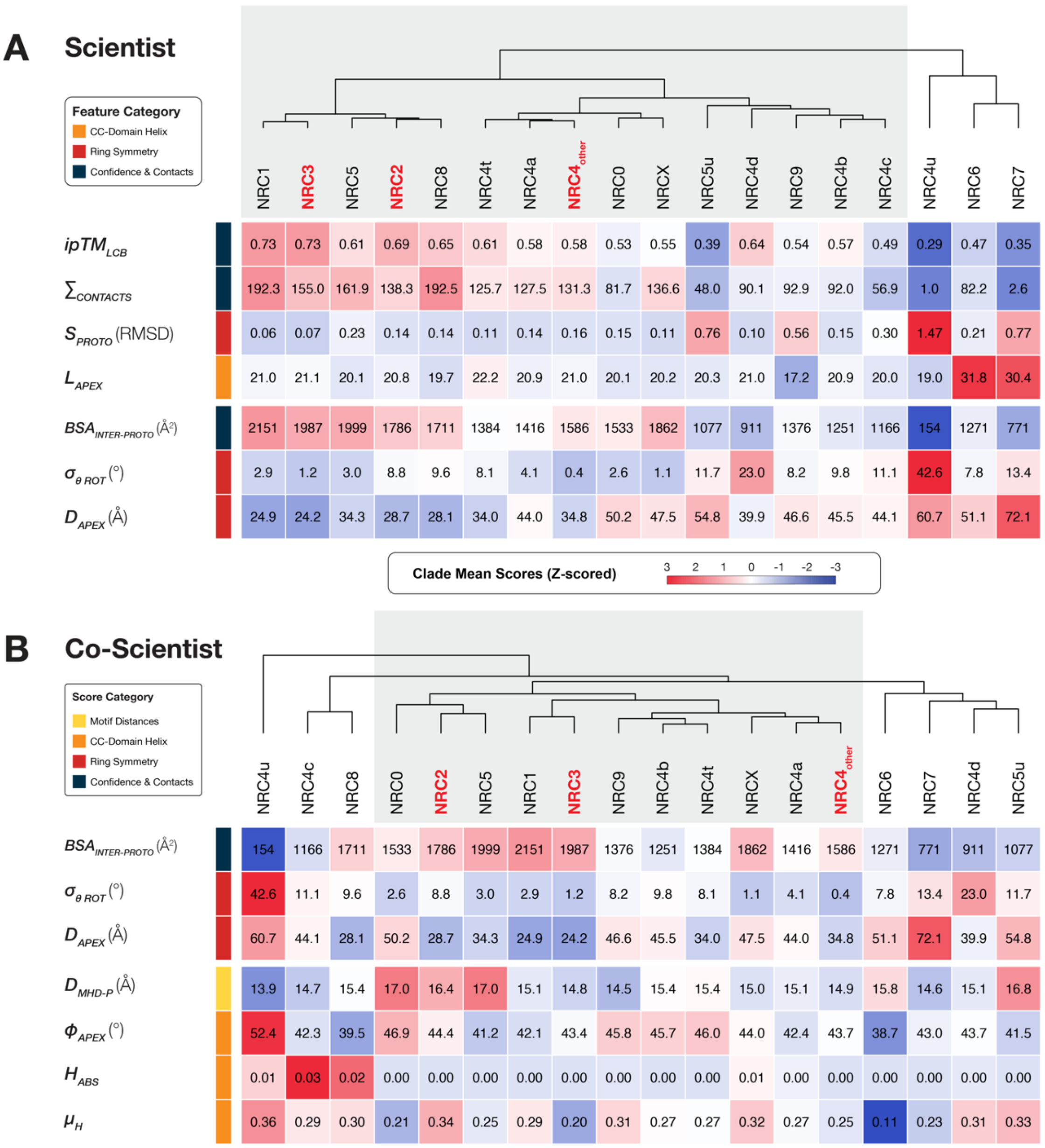


**Figure S6. Both Scientist and co-scientist generated SNI parameters detect NRC7 clade as an outlier.** Scientist-only parameters include $ipTM_{LCB}$, $\sum_{CONTACTS}$, $S_{PROTO}$, and $L_{APEX}$. Co-scientist parameters include $D_{MHD\text{-}P}$, $\varphi_{APEX}$, $H_{ABS}$, and $\mu_{H}$. Parameters shared between both include $BSA_{INTER\text{-}PROTO}$, $\sigma_{\theta\ ROT}$, and $D_{APEX}$. (**A**) In hierarchical clustering of Scientist-only parameters NRC4u, NRC6, and NRC7 clustered outside of the core NRC hexamer cluster (highlighted grey). (**B**) In hierarchical clustering of co-scientist parameters NRC8, NRC4c, NRC4d, and NRC5d are additionally placed outside of the core cluster. Both methods found NRC7 to be outside of the core cluster (highlighted grey). (**A**) & (**B**) Clades with empirical cryo-EM structures are highlighted red.

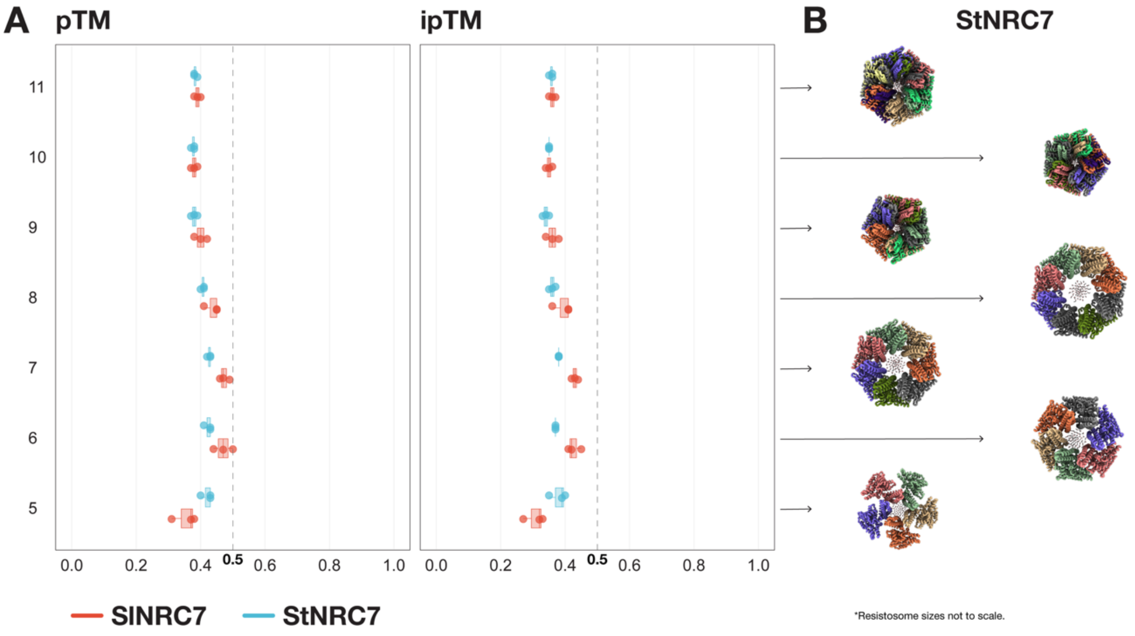


**Figure S7. NRC7 does not model well with AlphaFold 3 when modeled with 5-11 protomers.** **(A)** pTM and ipTM confidence scores for St- and SlNRC7 models predicted with 5–11 protomers. All models were generated in three independent replicates. Throughout the seven predicted stoichiometries, pTM and ipTM scores did not improve at any protomer number, including the experimentally validated 11-mer, and remained consistently low. **(B)** Representative StNRC7 predicted structures (seed = 1) are shown. Model sizes are not to scale. extending the protomer number beyond eight resulted in overlapping clashing protomers, suggesting that AlphaFold3 fails to model the higher-order architecture of the NRC7 resistosome.
