## Supplementary Data S6 for "AI-guided discovery of atypical protein assemblies": [EXTERNAL] ├░Y╠ê┬o┬1⁄4 Co-scientist - Hypothesis 1008902880708 - Quantitative SEED Index for Identifying Unconventional NRC-NLRs via Tiered AlphaFold 3 Screening - For research purposes only - Google confidential.docx

### 🧬 Co-scientist - Quantitative SEED Index for Identifying Unconventional NRC-NLRs via Tiered AlphaFold 3 Screening

**About**: *This is an experimental system for generating novel and testable hypotheses. The hypotheses are generated by a model and may be wrong. For research purposes only.*

### Proposal

$\def\mathcal#1{\mathit{#1}}\def\mathscr#1{\mathit{#1}}$

#### 1. The Eight Quantitative Parameters

The SEDI score for any sequence is the weighted sum of Z-scores calculated relative to the Reference Set (PDBs: 9RI9, 9FP6, 9CC8).

**Parameter 1: P-loop Cleft Solvent Accessibility Ratio ($R_{SASA}$)**

- **Method:** Ratio of the SASA of the Walker A Lysine in the monomeric model vs. the active hexameric model.
- **Rationale:** Canonical NRCs show a significant decrease in P-loop accessibility upon hexamerization.
- **Unconventional Signal:** A ratio near 1.0 suggests the P-loop remains exposed in the hexamer, indicating a potential failure to "lock" the nucleotide or a non-canonical activation mechanism.

**Parameter 2: LRR-NB-ARC Curvature Radius ($C_{LRR}$)**

- **Method:** Fit a circle to the $C\alpha$ coordinates of the LRR repeats and measure the radius of curvature.
- **Rationale:** The LRR maintains the "resting" state. 9RI9 (inactive) and 9CC8 (active) show distinct LRR curvatures that facilitate auto-inhibition.
- **Unconventional Signal:** A significantly tighter or flatter LRR arc indicates a different regulatory footprint or a deviation from standard auto-inhibitory control.

**Parameter 3: Protomer Interface Packing Efficiency ($\eta_{int}$)**

- **Method:** (Interface Buried Surface Area) / (Interface Frustration Energy, $E_{if}$ from FoldX).
- **Rationale:** This ratio identifies interfaces that may appear large in AI-generated models but are energetically unstable.
- **Canonical Baseline:** High efficiency (large burial, low energy) as seen in 9FP6.
- **Unconventional Signal:** Low efficiency indicates a physically implausible or unstable interface.

**Parameter 4: N-terminal Helix (MADA) Pore-Axis Displacement ($\Delta d_{pore}$)**

- **Method:** In a 3-seed AF3 hexamer run, calculate the mean distance of the MADA motif (residues 13–17) from the central symmetry axis.
- **Rationale:** Canonical NRCs (9FP6) form a tight funnel at the pore center.
- **Unconventional Signal:** High displacement or high variance across seeds suggests the N-terminus is not designed for standard pore formation, pointing to potential organelle-targeting or sensor-like functions.

**Parameter 5: CC-LRR Spatial Occlusion Index ($O_{auto}$)**

- **Method:** Distance between the CC-domain EDVID motif and the LRR-C terminal interface in the monomeric "resting" model.
- **Rationale:** Canonical NRCs use the LRR to "shield" the CC domain to prevent auto-activity.
- **Unconventional Signal:** An increased distance ($>5$ Å deviation from 9RI9) suggests the LRR does not regulate the CC domain in the standard way.

**Parameter 6: Evolutionary-Structural Interface Score ($S_{ESI}$)**

- **Method:** $\sum$ (ConSurf Score $\times$ Contact Frequency) for all residues at the protomer interface.
- **Rationale:** This distinguishes biological interfaces from potential modeling artifacts.
- **Canonical Baseline:** High conservation at the NB-ARC/NB-ARC and CC/CC interfaces.
- **Unconventional Signal:** Low $S_{ESI}$ suggests the modeled interface is not under evolutionary selection, marking the sequence as structurally novel or non-hexameric.

**Parameter 7: NB-ARC "Switch" Proximity ($P_{switch}$)**

- **Method:** Euclidean distance between the Walker A Lysine and the MHD Histidine in a single high-accuracy AF3 monomer run.
- **Rationale:** This measures if the protein can adopt the "closed" geometry required for ADP binding.
- **Unconventional Signal:** A distance $>15$ Å (canonical is ~10–12 Å) indicates a deviation from the standard molecular switch mechanism.

**Parameter 8: Interface Confidence Weight ($w_{ipTM}$)**

- **Method:** The raw AF3 ipTM (interface predicted Template Modeling score).
- **Rationale:** Acts as a quality filter. If $w_{ipTM} < 0.5$, the structural metrics are penalized to prevent false novelty signals from poor models.

#### 2. Implementation Protocol

1. **Tiered Screening (GPU Optimization):**
   - **Phase 1:** Run AF3 Monomers for all 6,000 sequences. Calculate $P_{switch}$, $C_{LRR}$, and $O_{auto}$.
   - **Phase 2:** Only sequences that sit significantly outside the reference distribution proceed to the Forced Hexamer modeling. This reduces the total AF3 workload.
   - **Phase 3:** Run 3 seeds for the hexamer to calculate $\Delta d_{pore}$ and $\eta_{int}$.
2. **Dynamic Baseline Calculation:**
   - The "Canonical Baseline" is defined as the 95% Confidence Interval of the distribution formed by the NRC2, NRC3, and NRC4 reference set.
   - A sequence is flagged as "Unconventional" if it sits $>3$ Standard Deviations outside this canonical cluster in at least 3 of the 8 parameters.

#### 3. Summary of SEDI Characteristics

- **Addressing Heterogeneity:** By using a cluster-based Z-score rather than a fixed angle, the index accommodates the natural structural differences between NRC2 and NRC4.
- **Domain Scope Expansion:** The inclusion of the LRR-NB-ARC curvature and CC-LRR occlusion identifies unconventional auto-inhibition, a key feature of sensors.
- **Modeling Noise Reduction:** By replacing the ligand-induced delta with the static $P_{switch}$ and weighting parameters by $w_{ipTM}$, the index is designed to be more robust against stochastic errors in protein modeling.
- **Computational Feasibility:** The tiered screening protocol focuses GPU resources on the most promising unconventional candidates.

### Reviews summary

$\def\mathcal#1{\mathit{#1}}\def\mathscr#1{\mathit{#1}}$

### Appendix:

**Motivation:**

$\def\mathcal#1{\mathit{#1}}\def\mathscr#1{\mathit{#1}}$

**Coherence:**

$\def\mathcal#1{\mathit{#1}}\def\mathscr#1{\mathit{#1}}$

**Deep verification:**

$\def\mathcal#1{\mathit{#1}}\def\mathscr#1{\mathit{#1}}$
