## Supplementary Data S6 for "AI-guided discovery of atypical protein assemblies": [EXTERNAL] ├░Y╠ê┬o┬1⁄4 Co-scientist - Hypothesis 62869464099376 - SNI-V2 Framework_ Integrating CC-Core Packing Density and Empirical Calibration for NRC-NLR Structural Novelty Assessment - For research purposes only - Google confidential.docx

### 🧬 Co-scientist - SNI-V2 Framework: Integrating CC-Core Packing Density and Empirical Calibration for NRC-NLR Structural Novelty Assessment

**About**: *This is an experimental system for generating novel and testable hypotheses. The hypotheses are generated by a model and may be wrong. For research purposes only.*

### Proposal

$\def\mathcal#1{\mathit{#1}}\def\mathscr#1{\mathit{#1}}$

#### The SNI-V2 Framework with CC-Core Calibration**^31^**

The SNI-V2 framework incorporates a 10th parameter focused on the CC-domain's functional core and introduces a pre-screening calibration phase to define novelty thresholds.

##### 1. The 10 Quantitative Parameters of SNI-V2**^37^**

| # | Parameter Name | Unit | Biological Significance |
| --- | --- | --- | --- |
| **1** | **Protomer Angular Deviation ($\Delta \theta_{sym}$)** | Degrees | Measures deviation from $60^\circ$ symmetry^5^. Detects $C_5$ (ZAR1-like) vs $C_6$ (NRC2-like)^24^. |
| **2** | **MADA Helix Tilt ($\Phi_{tilt}$)** | Degrees | Angle of $\alpha$1-helix to pore axis^9^. Measures if the "funnel" can reach the membrane^26^. |
| **3** | **ARC1-ARC2 Aperture ($\Psi_{ARC}$)** | Degrees | The "clamshell" opening^30^. Associated with the $10^\circ$ shift required for hexamerization^1^. |
| **4** | **Normalized Interface Area ($\bar{\rho}_{BSA}$)** | Ratio | $BSA_{query} / BSA_{ref}$ for the NB-LRR and CC-NB interfaces^33^. |
| **5** | **LRR Curvature Radius ($R_{LRR}$)** | Å | A constricted radius^32^ suggests evolution toward specific ligand perception^18^. |
| **6** | **Linker Span ($\Lambda_{link}$)** | Å | Distance from the $\alpha$4-helix C-term to the NB-domain N-term. Measures structural flexibility^22^. |
| **7** | **Hydrophobic Moment Alignment ($\Delta \mu_H$)** | Vector | Alignment of $\alpha$1-helix hydrophobic face to the pore interior^36^. |
| **8** | **MHD-Sensory Coupling ($\delta_{MHD}$)** | Å | Distance between the MHD motif and LRR internal loops (allosteric sensor-actuator link^23^). |
| **9** | **Stoichiometry Delta ($\Delta \text{ipTM}$)** | Score | $\text{ipTM}_{hexamer} - \text{ipTM}_{pentamer}$. Measures inherent oligomeric preference. |
| **10** | **CC-Core Packing Density ($\chi_{core}$)** | $\text{kcal/mol}\cdot\text{Å}^2$ | Quantifies the hydrophobic packing density of the $\alpha$2-$\alpha$4 CC barrel core^7^. Relevant for resistosome stability^20^. |

##### 2. Implementation Pipeline

**Phase 0: Empirical Threshold Calibration** Before screening the 6,000-sequence library^38^, the SNI-V2 is calculated for a **Control Set**:

- **Canonical Helpers:** NbNRC2, SlNRC3, NbNRC4, SlNRC2^10^.
- **Known Sensors:** Rx, Rpi-blb2, Bs2, Sw5-b^11^. Thresholds for "Unconventional" status^39^ are set at the point where $95\%$ of known sensors are distinguished from known helpers, replacing hypothetical thresholds with a data-driven boundary.

**Phase 1: Tiered High-Throughput Screening**

- **Tier 1 (Monomer Scan):** Model 6,000 sequences as monomers. Extract **#3 (Aperture), #8 (MHD), and #10 (CC-Core)**. Sequences where $\chi_{core}$ (CC-Core) is lower than the reference (indicating a potential degenerated pore-forming domain) are prioritized as "Unconventional Non-Pore Scaffolds."
- **Tier 2 (Multimer Scan):** Model the top 500 divergent candidates as homo-hexamers in AF3^25^ to calculate the full 10-parameter SNI.

##### 3. Design Rationale

1. **CC-Domain Analysis:** By adding **Parameter #10 ($\chi_{core}$)**, the index can distinguish between functional NRCs and those that have lost the ability to form a pore due to mutations in the hydrophobic core of the CC domain, even if the overall shape remains similar.
2. **Scoring System Calibration:** The **Control Set calibration phase** moves the framework from hypothetical Z-scores to biologically validated thresholds, reducing false-positive rates^27^ during the screening of 6,000 sequences.
3. **High-Throughput Tiering:** Using the CC-core density in Tier 1 allows for the early detection of "degenerate helpers"^19^ (NLRs that look like NRCs but cannot form a stable resistosome), allowing multimer modeling resources to be allocated to the most divergent candidates^40^.
4. **Structural Precision:** Parameter #6 specifically measures the distance between the CC $\alpha$4-helix and the NB domain, which is a known site of structural rearrangement during activation^22^.

[2]  [De Novo Design of SIK3 Inhibitors via Feedback-Driven Fine-Tuning of Seq2Seq-VAE.](https://vertexaisearch.cloud.google.com/grounding-api-redirect/AUZIYQF-5-7xOIMYE8th_hJNLML8PMH_qGzyl8UQAocR8eAk7hZvlo-VKDek7VoM6z_QLmDvInyyMoaXCy-XiaPGokGLUm8V35_U8wNX4iTXlnx894H7e7WbglB-BiptQG7I)

[3] AmirAli, Ryohei, Sophien, Yu.  [Can AI modelling of protein structures distinguish between sensor and helper NLR immune receptors?.](https://www.biorxiv.org/content/10.1101/2024.11.24.625045v1) Published 2024. [https://www.biorxiv.org/content/10.1101/2024.11.24.625045v1.](https://www.biorxiv.org/content/10.1101/2024.11.24.625045v1)

[4] AmirAli, Raoul, O., Ryohei, Sophien, Yu.  [Can AI modelling of protein structures distinguish between sensor and helper NLR immune receptors?.](https://www.biorxiv.org/content/10.1101/2024.11.24.625045v2) Published 2024. [https://www.biorxiv.org/content/10.1101/2024.11.24.625045v2.](https://www.biorxiv.org/content/10.1101/2024.11.24.625045v2)

[5] Jogi, AmirAli, P., Andres, Jake, Jiorgos, et al.  [A disease resistance protein triggers oligomerization of its NLR helper into a hexameric resistosome to mediate innate immunity.](https://www.biorxiv.org/content/10.1101/2024.06.18.599586v1) Published 2024. [https://www.biorxiv.org/content/10.1101/2024.06.18.599586v1.](https://www.biorxiv.org/content/10.1101/2024.06.18.599586v1)

[6]  [Unmasking the invaders: NLR-mal function in plant defense - PubMed Central.](https://vertexaisearch.cloud.google.com/grounding-api-redirect/AUZIYQEHFuI_Re7WAV7ECP1rAnoV3TU2jGIwqArjQAQFaWZr7wIWJPOftInkVNVvrP7H-Y2AfwCFPO29VIpzoPJjsa1dgvA8UVCQd0UF8yWwqHyxKeHJq1TjCDUR2sLGujjqcoto6u40uWz4lMnQIURK)

[7] Hung-Yu, Kim-Teng, Foong-Jing, O., Chih-Hang.  [A hydrophobic core in the coiled-coil domain essential for NRC resistosome function.](https://www.biorxiv.org/content/10.1101/2025.01.21.634219v1) Published 2025. [https://www.biorxiv.org/content/10.1101/2025.01.21.634219v1.](https://www.biorxiv.org/content/10.1101/2025.01.21.634219v1)

[8] Lida, P., Hiroaki, Jessica, Vergara, Rongrong, et al.  [Plant pathogens convergently evolved to counteract redundant nodes of an NLR immune receptor network.](https://www.biorxiv.org/content/10.1101/2021.02.03.429184v1) Published 2021. [https://www.biorxiv.org/content/10.1101/2021.02.03.429184v1.](https://www.biorxiv.org/content/10.1101/2021.02.03.429184v1)

[9] Matthew, A.  [AnglerFish: a webserver for defining the geometry of α-helices in membrane proteins.](https://www.ncbi.nlm.nih.gov/pmc/articles/PMC5860525/) Published 2016. [https://www.ncbi.nlm.nih.gov/pmc/articles/PMC5860525/.](https://www.ncbi.nlm.nih.gov/pmc/articles/PMC5860525/)

[10]  [Activation of the helper NRC4 immune receptor forms a hexameric resistosome | Request PDF - ResearchGate.](https://vertexaisearch.cloud.google.com/grounding-api-redirect/AUZIYQEzF9SZV9FSeyX8hYe6euRl_a1RD-rctlx3ge84l4W1OAfaaTLkTCBpG6KKbjdwlfxu6QnmuHDBn8T_l0D0M3Xb6hqh6LpM9MALUZMRNaL1M0OMFWN19is2n0uuDsqEzbcrguSoVFBVJ03L-FvcCZ19p8ssgVjxNZ8gIdCH2QqMgvOJyGeO5YYLZpFNB_h1lCYSN8FBw1Ejgx42wYWxDC2G9m-pw5dPdcTlxjzoZ5cuKvD7IuhA1Wz2Ow==)

[11] Chih-Hang, Sophien.  [Tomato Prf requires NLR helpers NRC2 and NRC3 to confer resistance against the bacterial speck pathogen Pseudomonas syringae pv. tomato.](https://www.biorxiv.org/content/10.1101/595744v1) Published 2019. [https://www.biorxiv.org/content/10.1101/595744v1.](https://www.biorxiv.org/content/10.1101/595744v1)

[12]  [AFsample: improving multimer prediction with AlphaFold using massive sampling.](https://vertexaisearch.cloud.google.com/grounding-api-redirect/AUZIYQH0glzQ2tjQGiOoYRMB1yn2RjWHKqziXrtMLBHEqRYQsYwrQRbImZEwPeae9BeJYocc_4cTELj2ZOKXt2Wa_VUlrn617HqNGAdBFAg5OAEdp4K4NjE93xkphg33Qno0OwwG6JHW9qT3JaUh8jHB_w66vISugG9yOMU75xMUUSIja03FqehmEFgHg4i095oaLeiMCE2wH8zlUeQMes9jKsRfPLWoPFddbhkT13WYeG7XWaJ9-TaNUES-N0A=)

[13]  [NRC Immune receptor networks show diversified hierarchical genetic architecture across plant lineages | Request PDF - ResearchGate.](https://vertexaisearch.cloud.google.com/grounding-api-redirect/AUZIYQFo5yc983u7SuQ7-F3qbfa04yC7mDENXl6U_fNHpYTaHXW2qxZvhYAobX299pyzYg7K3vcRbNXHTEJgUVRsDfwH9DdHzkY5NJEZhxnTBackqDM-qpv2tojVzfMZshusfwq3Rwyqu5S7e4BfrDnA30ow2wcDSz2gkWo7IS28lzczDhpnIS0jWaLLDv2_v04GjyF0VUJik9yQlJk1iLqOhtWV8ykaAuML1YpzTnCy0JL3edxSXarXdBdYnzELZ9yh9KNcEg9M9j_g_h6RXh8HAMTcduPLLQ==)

[14] Jaydeep, Marija, L..  [IRAA : A statistical tool for investigating a protein–protein interaction interface from multiple structures.](https://www.ncbi.nlm.nih.gov/pmc/articles/PMC9793972/) Published 2022. [https://www.ncbi.nlm.nih.gov/pmc/articles/PMC9793972/.](https://www.ncbi.nlm.nih.gov/pmc/articles/PMC9793972/)

[15] Natsumi, Hayden, J., Xiahao, Sneha, Kawsar, et al.  [Structural basis of NLR activation and innate immune signalling in plants.](https://www.ncbi.nlm.nih.gov/pmc/articles/PMC8813719/) Published 2021. [https://www.ncbi.nlm.nih.gov/pmc/articles/PMC8813719/.](https://www.ncbi.nlm.nih.gov/pmc/articles/PMC8813719/)

[16]  [A hierarchical immune receptor network in lettuce reveals contrasting patterns of evolution in sensor and helper NLRs | bioRxiv.](https://vertexaisearch.cloud.google.com/grounding-api-redirect/AUZIYQGr9SbKKMwzHBTrsPSpqYuGzuHQ_k3IccJKyy9UqOIZBGimanmwqN2-5XOOgnYeV-q693Xv9sI42mGYv8J39RMi-QYyVD36VKUQ3PLtcdEXgs7CDsa7s1QP0440tUu6WaocCwQJiB3LZyyx5-sEuYXG0NFjxb3N2Vqs6stAtNSjYQ==)

[22]  [(PDF) NLR immune receptors: structure and function in plant disease resistance.](https://vertexaisearch.cloud.google.com/grounding-api-redirect/AUZIYQGyQCrUTJFf1gouUFNuZgCJ6d7vXyNvbq1OWswH6AkVYRrA7MQSSRCqELWRyv_eCcB7vbun10_f5n9ZKw71Td-Pd4JdaTsrUn5qOAifkNA6w8C38ZT1-l-WE1I3F3zrPpzr-h9G-eutsA11X7Isa0oPwWdwV42XNsWSx8oj4kbYznumN758lER9SzksO6tzHeggsGn5qG4qnqjIEKYDWDi8rUGpNG_3T32-yCIB_ncPBlQQmpI=)

[23]  [Nucleocytoplasmic Distribution Is Required for Activation of Resistance by the Potato NB-LRR Receptor Rx1 and Is Balanced by Its Functional Domains - PubMed Central.](https://vertexaisearch.cloud.google.com/grounding-api-redirect/AUZIYQGfhSd-FxaCD4XtElN7_g571O38FFqf1XE1vexW-hjsBZ37VvGIOqhZpXhZWw32zfGy4TwAPBr_FEkD3FLPI0uzzJVYj-LdOz8ZaZLH9d6alZPtShsZBxB3XlyiGd-jLC7L8I_EEiJb0MS6Uqw=)

[24] Jogi, AmirAli, P., Andres, Jake, Jiorgos, et al.  [A disease resistance protein triggers oligomerization of its NLR helper into a hexameric resistosome to mediate innate immunity.](https://www.biorxiv.org/content/10.1101/2024.06.18.599586v1) Published 2024. [https://www.biorxiv.org/content/10.1101/2024.06.18.599586v1.](https://www.biorxiv.org/content/10.1101/2024.06.18.599586v1)

[25] Pawan, Jian, Jianlin.  [PSBench: a large-scale benchmark for estimating the accuracy of protein complex structural models.](https://arxiv.org/abs/2505.22674) Published 2025. [https://arxiv.org/abs/2505.22674.](https://arxiv.org/abs/2505.22674)

[26] P, Daniel, Hsuan, AmirAli, Sophien.  [NLR receptors in plant immunity: making sense of the alphabet soup.](https://www.ncbi.nlm.nih.gov/pmc/articles/PMC10561179/) Published 2023. [https://www.ncbi.nlm.nih.gov/pmc/articles/PMC10561179/.](https://www.ncbi.nlm.nih.gov/pmc/articles/PMC10561179/)

[27] AmirAli, Ryohei, Sophien, Yu.  [Can AI modelling of protein structures distinguish between sensor and helper NLR immune receptors?.](https://www.biorxiv.org/content/10.1101/2024.11.24.625045v1) Published 2024. [https://www.biorxiv.org/content/10.1101/2024.11.24.625045v1.](https://www.biorxiv.org/content/10.1101/2024.11.24.625045v1)

[28]  [Combining computational and experimental screening for rapid optimization of protein properties | PNAS.](https://vertexaisearch.cloud.google.com/grounding-api-redirect/AUZIYQEdptt3p48waboNKeJDDulPYYwc7hK7TPzwPX1tLw6tder_zlrHklqsiq7aWJ5ma02MDWPzCiZKfNpzagyXxY01p8SlnKRAdM-D-tCxiTdylbZdsQMwQKCFWJZf8oTzPpvsPPJ3jin9wKw=)

[40]  [AlphaFold can be used to predict the oligomeric states of proteins - bioRxiv.](https://vertexaisearch.cloud.google.com/grounding-api-redirect/AUZIYQF4ZYNk3U-o1kTEy6F2d2Ky-aW0C3POXGsuapims1DhkO8TE6krhXZPQ4T1wrvc8Zq5zdjjFng1g8MqfUV39PDJfcA_A13UZaN3nutPWzN2vK0mnKyth-24xui84HBGsLeajPVCGTLu7ZBhjSj1XvKT4_fVPM01Q1LSeDlyuMAGVA==)

### Reviews summary

$\def\mathcal#1{\mathit{#1}}\def\mathscr#1{\mathit{#1}}$

#### 1. Executive Verdict

The SNI-V2 framework proposes a 10-parameter quantitative Structural Novelty Index (SNI) designed to distinguish unconventional NRC-NLRs from canonical hexameric resistosomes through a tiered, data-driven screening pipeline. While the framework correctly identifies critical biological markers such as coiled-coil (CC) core packing and stoichiometry deltas, it relies on incorrect assumptions regarding ground-truth protein states and the observability of N-terminal motifs in experimental data. **Verdict: Proceed with Caution (Requires Fundamental Correction of Baselines).**

#### 2. Critical Flaws

- **Ground Truth Misidentification:** The hypothesis defines PDB 9RI9 (SlNRC3) as a canonical hexameric resistosome to derive baseline metrics for symmetry and aperture. However, PDB 9RI9 is established in the literature as a **resting-state homodimer**. Using a dimer to define hexameric protomer angular deviation or pore aperture is a fatal factual error that would yield mathematically invalid novelty thresholds.
- **N-terminal Baseline Observability:** Parameters #2 (MADA Helix Tilt) and #7 (Hydrophobic Moment) rely on the coordinates of residues 1–16 of the $\alpha$1-helix. These residues are disordered or missing in the experimental PDBs provided as ground truth (9FP6, 9CC8, and 9RI9). Without substituting these experimental baselines with AlphaFold 3-reconstituted models, these parameters cannot be measured or compared.

#### 3. Addressed Objections

Initially, the feasibility of the proposal was questioned due to the extreme computational cost of modeling 6,000 sequences as hexamers in AlphaFold 3. This was resolved by the framework’s **tiered screening architecture**, which utilizes a computationally inexpensive monomeric scan (Phase 1, Tier 1) to filter the library down to 500 high-priority candidates for multimeric modeling. Additionally, concerns regarding the ability of structural modeling to distinguish between closely related NLRs were countered by evidence that AlphaFold 3 confidence scores (ipTM) and the presence of stable N-terminal "funnel" structures are statistically significant discriminators between functional helpers and evolved sensors.

#### 4. Validated Risks & Limitations

- **Absence of a Unified Scoring Function:** The framework defines 10 heterogeneous metrics (measured in degrees, Å, and kcal/mol) but lacks a formalized mathematical formula to aggregate these values into a single ranking score. This complicates the identification of "novelty" when a sequence is divergent in one parameter but canonical in others.
- **Implementation Risk for Geometric Metrics:** Technical evaluations indicate that automated extraction of Buried Surface Area (BSA) and LRR Curvature Radius frequently fails or returns physically impossible values in complex NLR geometries. The hypothesis relies on these metrics without specifying specialized algorithmic solutions (e.g., Voronoi-tessellation or point-cloud fitting).
- **Calibration Sample Size:** Phase 0 relies on a limited control set of approximately 8–10 proteins. This sample size may be insufficient to establish a robust 95% confidence boundary for a 6,000-sequence dataset, increasing the risk of over-calibrating the index to a narrow set of well-characterized proteins.

#### 5. Supporting Arguments & Evidence

- **CC-Core Packing Density ($\chi_{core}$):** Parameter #10 is a significant strength, supported by 2025 research identifying a conserved hydrophobic core ($\alpha$2-$\alpha$4 helices) essential for NRC resistosome function. This provides a physics-based metric to identify "degenerate" sensors that sequence-based motifs might miss.
- **NB-ARC Aperture Logic:** The inclusion of Parameter #3 is grounded in recent structural comparisons showing that a $10^\circ$ interdomain widening in the NB-ARC module is required to accommodate hexameric stoichiometry over the pentameric ZAR1-like state.
- **Stoichiometry Delta (Parameter #9):** Utilizing the difference in predicted interface accuracy (ipTM) between $C_5$ and $C_6$ models is a validated methodology for predicting quaternary structure and identifying functional divergence in NLRs.

#### 6. Alignment & Novelty

- **Alignment:** The hypothesis is highly aligned with the research goal, providing exactly 10 quantitative parameters designed for high-throughput screening of the Solanaceae dataset using the specified PDB and AlphaFold 3 resources.
- **Novelty:** The individual biological insights (CC-core significance, stoichiometric shifts, and AF3 classification) have been previously published or preprinted in 2024–2025. The novelty of this proposal lies in the **engineering synthesis** of these findings into a formalized implementation pipeline for large-scale NLRome annotation.

#### 7. Feasibility Assessment

- **Resource Intensity:** Moderate. The tiered approach reduces the multimer modeling load to ~1,000 tasks (500 candidates $\times$ 2 stoichiometries), which is manageable on a standard laboratory GPU cluster.
- **Technical Complexity:** High. Success depends on developing custom scripts for non-standard metrics, particularly the hydrophobic moment alignment and CC-core packing density.
- **Time to Verdict:** 4–6 weeks for Phase 0 calibration and the Tier 1 monomer scan.

#### 8. Conclusion

The SNI-V2 framework is a scientifically robust engineering refinement of the current state-of-the-art in plant immunology. Its primary value is the integration of recent structural discoveries (such as the CC-core packing density) into a searchable index. However, the framework requires fundamental correction of its baseline definitions—specifically the exclusion of PDB 9RI9 from hexameric metrics and the adoption of AF3-reconstructed models for N-terminal measurements—before it can be deployed as an accurate discovery tool.

#### Research contacts

$\def\mathcal#1{\mathit{#1}}\def\mathscr#1{\mathit{#1}}$

Based on the provided research articles, here are the specific researchers who are experts in the field and would be best suited to review the Structural Novelty Index (SNI-V2) framework. They are listed in order of their expertise regarding the most specific aspects of the hypothesis (structural metrics of NRC hexamers, AlphaFold 3 modeling, and the MADA motif).

#### 1. Jogi Madhuprakash and Michael W. Webster

**Justification:** These researchers are the primary authors of the study defining the **NbNRC2 hexameric resistosome (PDB: 9FP6)**, which is the central "ground truth" for the SNI. Their work explicitly identifies several of the quantitative parameters suggested in the SNI, such as the **interdomain angle shift** and **CC-domain variations**.

- **Support from Excerpts:** In the article *"A disease resistance protein triggers oligomerization of its NLR helper into a hexameric resistosome to mediate innate immunity"* (Abstracts 4, 8, 9), they describe the **10° widening of the NB-ARC** and the **bend in the α4 helix** of NRC2 compared to ZAR1. They also performed the pore size measurements (17–19 Å) that align with the SNI's focus on pore geometry.

#### 2. AmirAli Toghani

**Justification:** Toghani is a lead researcher in using **AlphaFold 3 (AF3)** to specifically distinguish between helper and sensor NLRs using structural confidence metrics. This aligns perfectly with the SNI’s **Parameter #9 (Stoichiometry Delta)** and the use of **pTM/ipTM scores** for high-throughput screening.

- **Support from Excerpts:** In *"Can AI modeling of protein structures distinguish between sensor and helper NLR immune receptors?"* (Abstract 3), Toghani demonstrates that AF3 consistently produces higher pTM and ipTM scores for helper NLRs compared to sensors and identifies that helpers form the "funnel-shaped" structures the SNI seeks to quantify.

#### 3. Sophien Kamoun

**Justification:** Kamoun is a senior author on nearly all the provided foundational papers regarding the **NRC network**, the **MADA motif**, and the **NRC2 hexamer structure**. His expertise spans the evolutionary biology of the 6,000-sequence NRC family and the structural transition from monomers to hexamers.

- **Support from Excerpts:** His work appears in Abstract 1 (MADA motif discovery), Abstract 3 (AF3 modeling), and Abstract 9 (NRC2 hexamer structure). He provides the overarching rationale for why a quantitative baseline is needed to identify "unconventional" NLRs within the Solanaceae dataset.

#### 4. Hiroaki Adachi

**Justification:** Adachi is the lead researcher who defined the **MADA motif**, which is a core component of the SNI (Parameters #2 and #7). He specialized in identifying how this motif degenerates in sensor NLRs, providing the evolutionary logic for the "Unconventional" status the SNI aims to detect.

- **Support from Excerpts:** In *"An N-terminal motif in NLR immune receptors is functionally conserved across distantly related plant species"* (Abstract 1), Adachi defined the MADA consensus sequence ($MADAxVSFxVxKLxxLLxxEx$) and established the HMM (Hidden Markov Model) scoring system mentioned in the SNI implementation pipeline.

#### 5. Mauricio P. Contreras

**Justification:** Contreras has published extensively on the biochemical mechanisms of the **NRC family** and the structural basis of how sensor NLRs activate their NRC helpers. He is well-versed in the "death switch" mechanism of the CC domain.

- **Support from Excerpts:** He is a co-author on the NRC2 hexamer structure papers (Abstracts 4, 9) and the AF3 sensor/helper classification study (Abstract 3), making him an expert on the **protomer interface** and **CC-core packing** parameters.

#### 6. Jianlin Cheng

**Justification:** While less focused on the specific biology of NLRs, Cheng is a specialist in the **computational stoichiometry prediction** using AlphaFold 3. He would be an ideal reviewer for the technical feasibility of **Parameter #9** and the **Phase 1: Multimer Scan** part of the implementation pipeline.

- **Support from Excerpts:** In *"Accurate Prediction of Protein Complex Stoichiometry by Integrating AlphaFold3 and Template Information"* (Abstract 2), Cheng describes the **PreStoi** method, which uses AF3 scores (AF-max and AF-avg) to rank stoichiometry candidates, directly supporting the SNI’s reliance on AF3 for determining oligomeric preferences.

#### 7. Alexander Förderer and Jiorgos Kourelis

**Justification:** These researchers provide the broader structural context of **CC-NLR resistosomes** and the importance of the **EDVID motif** and **α-helical packing**.

- **Support from Excerpts:** In *"NLR immune receptors: structure and function in plant disease resistance"* (Abstract 5), they detail the hydrophobic packing of the CC barrel (relevant to **Parameter #10**) and the role of the α1 helix in membrane insertion.

### Appendix:

**All reviews:**

**Correctness:**

$\def\mathcal#1{\mathit{#1}}\def\mathscr#1{\mathit{#1}}$

#### Related Article Abstracts

1. **[14, 20, 22, 24] A hydrophobic core in the coiled-coil domain essential for NRC resistosome function (Wang et al., 2025):** These preprints identify a novel hydrophobic core ($\alpha2$-$\alpha4$ helices) in the CC domain of NRCs. This is the direct basis for Parameter #10 ($\chi_{core}$) and validates its use as a marker for helper vs. sensor differentiation.
2. **[4, 8, 10] A disease resistance protein triggers oligomerization of its NLR helper into a hexameric resistosome to mediate innate immunity (Madhuprakash et al., 2024):** Provides the cryo-EM structure of the NbNRC2 hexamer (9FP6) and details the $10^\circ$ angular shift in the NB-ARC domain compared to ZAR1. This supports Parameters #1 and #3.
3. **[1, 19] An N-terminal motif in NLR immune receptors is functionally conserved across distantly related plant species (Adachi et al., 2019):** Defines the MADA motif and its degeneracy in sensors. This is relevant for Parameters #2, #7, and #10.
4. **[3, 23, 28] Can AI modeling of protein structures distinguish between sensor and helper NLR immune receptors? (Toghani et al., 2024/2025):** Demonstrates that AF3 confidence scores (pTM/ipTM) and the presence of funnel-shaped structures differentiate helpers from sensors. Supports Parameter #9 and the Tier 2 multimer scan.
5. **[9] RCSB PDB - 9FP6: Structure of the NbNRC2 hexameric resistosome:** Ground truth for the active hexameric state of canonical helpers.
6. **[15, 21] Activation of the helper NRC4 immune receptor forms a hexameric resistosome (Liu et al., 2024):** Ground truth for 9CC8. Notes that the $\alpha1$ helix is often disordered/flexible in experimental structures, highlighting the need for AF3 models in the baseline.
7. **[2] Accurate Prediction of Protein Complex Stoichiometry by Integrating AlphaFold3 and Template Information (Liu et al., 2025):** Supports the methodology of using AF3 scores to predict stoichiometry (Parameter #9).
8. **[12] Assessment of Self-Activation and Inhibition of Wheat Coiled-Coil Domain Containing NLR Immune Receptor Yr10 CG (Wu et al., 2024):** Discusses the MHD motif's role in activation, supporting Parameter #8.
9. **[11] Diversification of the “EDVID” packing motif... (Sulkowski et al., 2025):** Discusses CC domain variation, relevant to the structural parameters proposed.
10. **[18] Protein Quaternary Structures in Solution are a Mixture of Multiple forms (Marciano et al., 2022):** Provides context on AF's ability to predict oligomeric states using ipTM.

#### Detailed Assumptions

1. **Metric Extractability:** It assumes all 10 structural parameters (e.g., MADA tilt, BSA, CC-core density) can be reliably extracted from both PDB files and AlphaFold 3 models.
2. **Ground Truth State:** It assumes the provided PDB IDs (9FP6, 9RI9, 9CC8) are consistent references for the "active hexameric resistosome" state.
3. **Hydrophobic Core Significance:** It assumes the packing density of the CC barrel core ($\alpha2$-$\alpha4$) is a primary determinant of helper functionality and that this density is measurable at the monomeric level.
4. **Stoichiometric Contrast:** It assumes that the $\Delta ipTM$ between pentameric and hexameric models is a reliable indicator of "canonical" vs "unconventional" stoichiometry.
5. **Calibration Validity:** It assumes that a 95% threshold derived from a limited Control Set (8-10 proteins) will generalize to a 6,000-sequence library.

#### Comparison with Knowledge Base

- **PDB Consistency:** The idea proposes using **9RI9** as a reference for canonical NRC resistosomes. However, the Knowledge Base explicitly states that **9RI9 is a resting-state dimer**, not an active hexamer. Using it to define "canonical" resistosome metrics like angular deviation or pore aperture is a significant factual error.
- **The "Missing Residue" Problem:** The Knowledge Base "pdb results" show that Parameter #2 (MADA Helix Tilt) failed for all NRC PDBs because the N-terminal residues (1–16) are not resolved. The idea assumes these can be measured from the PDBs without specifying the use of AF3-reconstructed models for the baseline.
- **Technical Failure of Implementation:** The Knowledge Base logs indicate that Parameter #4 (BSA) and Parameter #5 (LRR Curvature) failed or returned physically impossible values (0.00 Å²) in preliminary scripts. The idea relies on these metrics for novelty detection.
- **Protein Identification:** The Knowledge Base "uniprot results" show that common names for NRCs (e.g., "NRC3 Sl") often return non-NLR proteins (ribosomal proteins, proteinase inhibitors). The idea’s "Phase 0" depends on a Control Set that requires precise accession-based identification to avoid calibration against incorrect proteins.

#### Reasoning about Correctness for Assumptions

1. **Metric Extractability:** **Questionable.** While mathematically sound, the technical failure of BSA and LRR curvature measurements in the Knowledge Base suggests these metrics are not yet "automatable" without significant debugging of the geometric implementation.
2. **Ground Truth State:** **Incorrect.** The reliance on 9RI9 as an "active" reference is a fundamental flaw. 9RI9 will not possess the hexameric interfaces required for the SNI metrics.
3. **Hydrophobic Core Significance:** **True/High Strength.** Parameter #10 is strongly supported by recent 2025 research (Wang et al.), confirming that the $\alpha2$-$\alpha4$ core is essential for helpers and degenerated in sensors. This is the most biologically sound part of the idea.
4. **Stoichiometric Contrast:** **True.** Recent literature (Toghani et al., 2024) supports using ipTM comparisons to distinguish helper/sensor behavior.
5. **Calibration Validity:** **Questionable.** A Control Set of only 4 sensors and 4 helpers is likely too small to establish a robust 95% data-driven boundary for 6,000 sequences without over-fitting.

#### Strength of Evidence

- **Direct Evidence:**
  - Strong evidence for the **MADA motif** [1] and the **CC-hydrophobic core** [14] being functionally critical for helpers.
  - Strong evidence for NRC2/NRC4 forming hexamers [4, 15].
  - Direct evidence that AF3 confidence scores distinguish helpers from sensors [3].
- **Indirect Evidence:**
  - The use of $\Delta ipTM$ for stoichiometry is supported by CASP16 performance metrics for complex prediction [2].
  - The use of monomeric scans to filter large datasets is a standard computational strategy to manage GPU resources.

#### Suggested Improvements

1. **Correction of Ground Truth:** Explicitly exclude 9RI9 from hexameric interface calculations or use it only as a reference for the "resting-to-active" transition distance.
2. **AF3-Derived Baseline:** Instead of raw PDBs, use high-confidence AF3 models of NbNRC2, SlNRC3, and NbNRC4 as the "Ground Truth Baseline" to ensure the disordered N-terminal MADA motif is resolved and measurable for Parameters #2 and #7.
3. **Accession Rigor:** Hardcode UniProt Accession IDs (e.g., A0A290U7C4 for NbNRC2) for the Control Set to prevent the naming discrepancies found in the Knowledge Base.
4. **Geometric Fixes:** Replace the failed Buried Surface Area (BSA) implementation with a more robust Voronoi-tessellation-based interface score (e.g., VoroIF) which is more reliable in AF3 contexts [26].

#### Assessment of Goal Requirements

- **5-10 Parameters?** Yes (10 parameters).
- **Measurable from PDB/AF3?** Mostly, though the Knowledge Base identifies technical hurdles for BSA and LRR metrics.
- **Leverage 9FP6, 9RI9, 9CC8?** Yes, though it misinterprets the oligomeric state of 9RI9.
- **Distinguish from canonical NRC2/3/4?** Yes, the index is explicitly designed around this.
- **Focus on interfaces and distances?** Yes (Parameters #1, #3, #4, #6).
- **Tiered/High-throughput?** Yes, Phase 1 includes a monomeric scan.

#### Reasoning about Correctness and Recommendation

The idea is scientifically sophisticated and incorporates the most recent structural biology findings regarding the CC domain's hydrophobic core (Parameter #10). This addition significantly elevates the "Structural Novelty Index" beyond simple geometric comparisons. However, the idea suffers from a major factual error regarding the oligomeric state of PDB 9RI9, which would lead to incorrect baseline calibration for several parameters. Furthermore, the Knowledge Base indicates that several of the proposed metrics (BSA, LRR curvature) are currently failing in technical implementation.

Despite these issues, the **biological logic is highly sound**, particularly the tiered screening approach and the focus on "Executioner" domain stability. The integration of Parameter #10 is an inspired move based on very recent (2025) data. I recommend testing the idea, provided the ground truth definitions are corrected and the geometric extraction scripts are debugged.

Answer: 6

**Novelty:**

$\def\mathcal#1{\mathit{#1}}\def\mathscr#1{\mathit{#1}}$

#### Related Article Abstracts

1. **[3] Can AI modeling of protein structures distinguish between sensor and helper NLR immune receptors?**
   - **Reasoning:** Directly explores using AlphaFold 3 (AF3) confidence scores (pTM/ipTM) and structural features (funnel formation) to classify NLRs into sensors and helpers, which is the foundational goal of the SNI.
2. **[13] A disease resistance protein triggers oligomerization of its NLR helper into a hexameric resistosome to mediate innate immunity**
   - **Reasoning:** Uses AF3 to model NRCs as pentamers vs. hexamers and compares confidence scores to determine preferred stoichiometry, directly relating to Parameter #9 ($\Delta \text{ipTM}$).
3. **[14] A hydrophobic core in the coiled-coil domain essential for NRC resistosome function**
   - **Reasoning:** Identifies a conserved hydrophobic core in the $\alpha$2–$\alpha$4 CC barrel that is essential for stability and is degenerated in sensors, providing the biological basis for Parameter #10 ($\chi_{core}$).
4. **[4] A disease resistance protein triggers oligomerization of its NLR helper into a hexameric resistosome to mediate innate immunity**
   - **Reasoning:** Provides the structural basis for Parameters #1 (Angular Deviation) and #3 (ARC Aperture) by identifying the $10^\circ$ interdomain shift and protomer differences between hexameric NRC2 and pentameric ZAR1.
5. **[2] Accurate Prediction of Protein Complex Stoichiometry by Integrating AlphaFold3 and Template Information**
   - **Reasoning:** Describes the general methodology of using AF3 ranking scores across various stoichiometry candidates to predict quaternary structure, supporting the logic of Parameter #9.
6. **[10] A disease resistance protein triggers oligomerization of its NLR helper into a hexameric resistosome to mediate innate immunity**
   - **Reasoning:** Compares resting dimers (9RI9) to activated hexamers (9FP6), noting rearrangements in the NB-ARC module and order-disorder transitions in the linker, relevant to Parameters #3 and #6.
7. **[28] Can AI Modelling of Protein Structures Distinguish Between Sensor and Helper NLR Immune Receptors?**
   - **Reasoning:** A preprint emphasizing that AF3-based structural classification is more robust than sequence-based motifs (like MADA) for uncharacterized NLRs, aligning with the Rationale of the idea.
8. **[1] An N-terminal motif in NLR immune receptors is functionally conserved across distantly related plant species**
   - **Reasoning:** Defines the MADA motif ($\alpha$1-helix) and its role in membrane insertion, providing the basis for Parameters #2 (Helix Tilt) and #7 (Hydrophobic Moment).
9. **[20] A hydrophobic core in the coiled-coil domain is essential for NRC resistosome function**
   - **Reasoning:** Further validates the CC-core importance across ZAR1 and NRCs and its degeneration in sensors, reinforcing Parameter #10.
10. **[5] NLR immune receptors: structure and function in plant disease resistance**
    - **Reasoning:** Details the mechanics of the $\alpha$1-helix funnel and the hydrophobic packing of the CC barrel ($\alpha$2-$\alpha$4), relevant to Parameters #2 and #10.
11. **[15] The activated plant NRC4 immune receptor forms a hexameric resistosome**
    - **Reasoning:** Describes the dense packing of the NRC4 hexamer (9CC8) and LRR arrangements, providing data for Parameters #4 (BSA) and #5 (LRR Curvature).
12. **[42] Structural basis of NLR activation and innate immune signalling in plants**
    - **Reasoning:** Explains the NB-ARC molecular switch and MHD motif interactions, providing the context for Parameter #8 (MHD-Sensory Coupling).
13. **[23] Can AI modelling of protein structures distinguish between sensor and helper NLR immune receptors?**
    - **Reasoning:** Reinforces that helper models show statistically higher predicted interface accuracy than sensors, supporting the "Stoichiometry Delta" approach.
14. **[12] Assessment of Self-Activation and Inhibition of Wheat Coiled-Coil Domain Containing NLR Immune Receptor Yr10 CG**
    - **Reasoning:** Discusses MHD motif mutations and ADP/ATP exchange, supporting the use of Parameter #8 as a measure of activation state.
15. **[30] RCSB PDB - 9CC8: Hexameric state of the NRC4 resistosome**
    - **Reasoning:** Ground truth structure for the hexameric resistosome used to extract baseline parameters for the SNI.

#### Aspects of the idea that were already tried

- **Using AF3 Confidence Scores for Classification:** The core concept of using AF3's internal metrics (pTM, ipTM) to distinguish between helper and sensor NLRs has been explicitly explored in [3], [23], and [28]. These studies established that helpers consistently yield higher confidence scores in oligomeric models than sensors.
- **Stoichiometry Prediction via ipTM:** Using the difference in ipTM between pentameric ($C_5$) and hexameric ($C_6$) models to predict stoichiometry is described in [2] and specifically applied to the NRC family in [13] and [39].
- **Structural Metrics for Hexamerization:** The identification of the $10^\circ$ widening of the NB-ARC aperture ($\Psi_{ARC}$) and specific protomer angular distances required for hexamerization was established in the structural comparisons of NRC2 and ZAR1 in [4], [8], and [10].
- **CC-Core packing as a functional determinant:** The role of the hydrophobic core in the $\alpha$2–$\alpha$4 barrel of the CC domain as a marker for functional NRCs vs. "degenerate" sensors was discovered and detailed in [14], [20], and [24].
- **MADA Motif and Funnel Topology:** Parameters related to the $\alpha$1-helix tilt and hydrophobic moment alignment are grounded in the ZAR1 "death switch" model [1, 5, 19, 34].

#### Novel aspects of the idea

- **Integrated SNI-V2 Index:** While individual parameters (pTM, angles, motifs) have been analyzed, the integration of 10 distinct quantitative structural metrics into a single weighted "Structural Novelty Index" specifically for high-throughput screening of the 6,000-sequence NRC family is a novel methodological synthesis.
- **Phase 0 Data-Driven Calibration:** The proposal to establish novelty thresholds by calibrating the index against a control set of 95% of known sensors vs. helpers replaces hypothetical Z-scores with biologically validated boundaries.
- **Parameter #6 (Linker Span):** Using the distance from the $\alpha$4-helix C-term to the NB-domain N-term as a quantitative metric specifically targets the known site of rearrangement (order-disorder transition) during activation, as mentioned in [10], but has not been utilized as a screening parameter.
- **Tiered Screening Architecture with $\chi_{core}$:** Utilizing Parameter #10 (CC-Core Packing Density) specifically in a "Tier 1 Monomer Scan" to identify "Unconventional Non-Pore Scaffolds" before investing resources in multimer modeling is a novel computational pipeline strategy.

#### Novelty review

The idea is **not significantly novel** to experts in the field. The core hypothesis—that AlphaFold 3 structural modeling and confidence scores can distinguish between functional NRC helpers and unconventional sensors or degenerate helpers—is the primary focus of very recent high-impact papers [3, 13, 14, 28].

Specifically, Parameter #9 (Stoichiometry Delta) is the exact method used in [13]. Parameter #10 (CC-Core) is the central discovery of [14] and [20]. Parameters #1, #3, and #4 are derived directly from the comparative structural analyses of the NRC2/NRC4 hexamers [4, 15]. The idea effectively organizes the most recent 2024–2025 structural breakthroughs into an implementation pipeline. While the "Index" formulation is a logical next step for high-throughput screening, the biological and structural insights it relies on have already been explicitly explored in the context of helper/sensor classification.

#### Reasoning about novelty and recommendation

The idea is a highly competent implementation plan that applies the latest structural biology of NLRs to a larger dataset. However, its novelty is "Limited" because it relies on structural features and AF3 scoring paradigms that have been published or preprinted within the last year [3, 13, 14]. The most promising technical addition is the quantification of CC-core packing density ($\chi_{core}$) as a Tier 1 monomer filter, which could save significant computational time in a 6,000-sequence screen.

Despite the low novelty score, I **recommend testing the idea** as an internal research tool. Establishing a quantitative SNI would be extremely valuable for processing the ~350 Solanaceae species dataset and identifying "Unconventional" sequences that deviate from the NbNRC2 baseline. It is a solid engineering refinement of current state-of-the-art research, even if the conceptual core is already present in the literature.

Answer: 4

**Feasibility:**

$\def\mathcal#1{\mathit{#1}}\def\mathscr#1{\mathit{#1}}$

#### Related Article Abstracts

1. **[4] A disease resistance protein triggers oligomerization of its NLR helper into a hexameric resistosome to mediate innate immunity (Madhuprakash et al., 2024):** Provides the structural basis for Parameter #1 (angular deviation) and Parameter #3 (ARC1-ARC2 aperture) by comparing NbNRC2 (hexamer) to AtZAR1 (pentamer).
2. **[14] A hydrophobic core in the coiled-coil domain essential for NRC resistosome function (Wang et al., 2025):** Directly informs Parameter #10 ($\chi_{core}$); it identifies the conserved hydrophobic center ($\alpha$2-$\alpha$4) that distinguishes helpers from sensors.
3. **[3] Can AI modeling of protein structures distinguish between sensor and helper NLR immune receptors? (Toghani et al., 2025):** Validates the use of AF3 confidence scores (pTM/ipTM) to classify NLRs, supporting Parameter #9.
4. **[1] An N-terminal motif in NLR immune receptors is functionally conserved across distantly related plant species (Adachi et al., 2019):** Defines the MADA motif ($\alpha$1-helix), which is essential for Parameters #2 and #7.
5. **[2] Accurate Prediction of Protein Complex Stoichiometry by Integrating AlphaFold3 and Template Information (Liu et al., 2025):** Demonstrates that ranking different stoichiometries (pentamer vs. hexamer) using AF3 scores is a valid method, supporting Parameter #9.
6. **[13] A disease resistance protein triggers oligomerization... (Madhuprakash et al., 2024):** Compares AF3 models to Cryo-EM Ground Truth (9FP6, 9CC8), establishing the feasibility of using AF3 for structural metrics.
7. **[26] Assessing scoring metrics for AlphaFold2 and AlphaFold3 protein complex predictions (Genz et al., 2025):** Evaluates the reliability of ipTM and PAE at protein interfaces, relevant for Parameter #4 and #9.
8. **[15] The activated plant NRC4 immune receptor forms a hexameric resistosome (Liu et al., 2023):** Provides data on protomer packing density and LRR orientation, relevant for Parameters #1, #4, and #5.
9. **[5] NLR immune receptors: structure and function in plant disease resistance (Förderer & Kourelis, 2023):** Describes the $\alpha$1-helix "funnel" mechanism, supporting the biological significance of Parameter #2 (Tilt).
10. **[12] Assessment of Self-Activation and Inhibition... (Wu et al., 2024):** Discusses the MHD motif's role in domain reconfiguration, supporting Parameter #8.
11. **[10] A disease resistance protein triggers oligomerization... (Madhuprakash et al., 2024):** Measures specific inter-domain angles (e.g., 85° in NRC2 vs 75° in ZAR1), validating the sensitivity of Parameter #1.
12. **[18] Protein Quaternary Structures in Solution are a Mixture of Multiple forms (Marciano et al., 2022):** Discusses AF's ability to predict prevalent multimeric states, supporting the "Stoichiometry Delta" logic.
13. **[25] Deep learning facilitates precise identification of disease-resistance genes (Liu et al., 2024):** Notes the practical constraints of AF3 high-throughput tasks, influencing the feasibility of modeling 6,000 sequences.
14. **[22] A hydrophobic core in the coiled-coil domain... (Wang et al., 2025):** Mentions that A72 (part of the core) is adjacent to the EDVID motif, justifying its inclusion in structural novelty metrics.
15. **[16] A disease resistance protein triggers oligomerization... (Madhuprakash et al., 2024):** Highlights the coordination of ATP in the NB-ARC pocket, relevant for the structural precision of Parameter #3.

#### Steps to Test the Idea

1. **Go/No-Go Experiment:** Select 10 known Canonical Helpers (NRC2, 3, 4) and 10 Known Sensors (Rx, Sw5b, etc.). Run AF3 multimer (hexamer) on all 20. Calculate the 10 SNI parameters. **Requirement:** The SNI must show a statistically significant gap ($p < 0.01$) between the two groups, particularly in Parameters #1, #9, and #10.
2. **Tier 1 Monomer Generation:** Perform high-throughput AF3 monomer modeling for the 6,000 NRC sequences. Use a truncated "Core" sequence (CC-NB-ARC, ~500 residues) to stay within token limits and reduce compute time.
3. **Tier 1 Metric Extraction:** Automate a Python script using Biopython and a hydrophobic packing tool (e.g., Rosetta's PackStat or a BSA-based script) to calculate Parameters #3 (Aperture), #8 (MHD), and #10 (CC-Core).
4. **Novelty Selection:** Rank sequences by Tier 1 SNI. Identify the top 500 "Divergent" candidates (highest deviation from NbNRC2 baselines).
5. **Tier 2 Multimer Modeling:** Model the 500 candidates as both pentamers and hexamers. Extract interface-specific parameters (#1, #4, #6, #9).
6. **Hydrophobic Moment Analysis:** For Parameter #7, apply the Eisenberg scale to the $\alpha$1-helix coordinates to determine the vector alignment relative to the central pore axis.
7. **Threshold Finalization:** Use the Control Set from Step 1 to set the "Novelty Boundary" (e.g., SNI > 7.5). Apply this to the 500 candidates to identify unconventional NLRs.

#### Feasibility Reasoning

The proposed SNI-V2 framework is highly grounded in current structural biology (leveraging very recent 2024/2025 insights into the CC-hydrophobic core and hexameric symmetry).

- **Computational Scalability:** The tiered approach is essential. Modeling 6,000 hexamers directly would be **Score 5 (Resource Intensive)** due to the massive GPU demand (~12,000 AF3 tasks if testing two stoichiometries). However, by using a Tier 1 monomer scan, the multimer load is reduced to 1,000 tasks (500 seq × 2 stoichiometries). This is manageable within 1 month on a standard lab cluster.
- **Metric Measurability:** 9 out of 10 parameters rely on standard atomic coordinates (distances and angles) which are easily extractable via automated scripts. Parameter #10 (Packing Density) requires more specialized but existing bioinformatic tools. Parameter #9 ($\Delta \text{ipTM}$) utilizes standard AF3 confidence outputs.
- **Ground Truth Usage:** The index correctly utilizes the 9FP6/9CC8 hexameric data to define canonical baselines (60° symmetry, specific aperture angles). It also incorporates recent data showing that sensors have degenerated CC-cores [14], making Parameter #10 a high-confidence filter.
- **Constraints:** The main bottleneck is the development of custom scripts for non-standard metrics like "Hydrophobic Moment Alignment" and "ARC1-ARC2 Aperture," and the sheer volume of monomer modeling.

Overall, the plan is scientifically robust and avoids the "forced folding" bias of AF3 by explicitly measuring stoichiometric preference (Parameter #9).

Answer: 6

**Impact potential:**

$\def\mathcal#1{\mathit{#1}}\def\mathscr#1{\mathit{#1}}$

#### Related Article Abstracts

1. **[1] An N-terminal motif in NLR immune receptors is functionally conserved across distantly related plant species (2019):** Defines the MADA motif and its role in cell death. Relevant for Parameters #2, #7, and #10, providing the biological basis for executioner activity.
2. **[3] Can AI modeling of protein structures distinguish between sensor and helper NLR immune receptors? (2025):** Demonstrates that AF3 confidence scores (pTM/ipTM) are higher for helpers than sensors. Directly supports Parameter #9 ($\Delta ipTM$) and the tiered screening strategy.
3. **[4] A disease resistance protein triggers oligomerization of its NLR helper into a hexameric resistosome... (2024):** Compares hexameric NbNRC2 to pentameric ZAR1, noting a $10^\circ$ angular shift in the NB-ARC. Essential for Parameter #3 (ARC1-ARC2 Aperture).
4. **[9] RCSB PDB - 9FP6: Structure of the NbNRC2 hexameric resistosome (2024):** One of the ground truth PDBs. Provides the $60^\circ$ symmetry baseline for Parameter #1.
5. **[10] A disease resistance protein triggers oligomerization... (2024):** Details the outward displacement of NB domains and pore diameter differences. Supports Parameters #1 and #5.
6. **[14] A hydrophobic core in the coiled-coil domain essential for NRC resistosome function (2025):** Identifies a specific hydrophobic center in $\alpha2$-$\alpha4$ of the CC domain. Directly justifies Parameter #10 ($\chi_{core}$).
7. **[15] The activated plant NRC4 immune receptor forms a hexameric resistosome (2023):** Provides structural data for the second ground truth (9CC8), highlighting the dense packing of the NRC4 hexamer.
8. **[17] Two genetically linked Arabidopsis TIR-type NLRs are required... (2025):** Discusses the Met-His-Asp (MHD) motif as a molecular switch. Supports Parameter #8 ($MHD$ coupling).
9. **[18] Protein Quaternary Structures in Solution are a Mixture... (2022):** Benchmarks AlphaFold-Multimer accuracy for stoichiometry. Relevant for validating Parameter #9.
10. **[20] A hydrophobic core in the coiled-coil domain is essential for NRC resistosome function (2025):** Confirms that CC-core residues are conserved in helpers but degenerated in sensors. Validates the rationale for Tier 1 screening.
11. **[23] Can AI modelling... (2024):** Notes that putative helpers consistently show higher ipTM than sensors. Key evidence for using $\Delta ipTM$ as a novelty metric.
12. **[26] Assessing scoring metrics for AlphaFold2 and AlphaFold3... (2025):** benchmarks ipTM as the most reliable metric for complex accuracy. Critical for validating the SNI's reliance on AF3 scores.

#### Assumptions for Impact Potential

1. **Stoichiometric Contrast Validity:** The idea assumes that the difference in confidence scores (ipTM) between modeling a sequence as a hexamer versus a pentamer is a reliable proxy for "structural frustration" and, consequently, evolutionary novelty or functional divergence.
2. **Monomer-to-Multimer Predictivity:** It assumes that monomeric parameters (like MHD distance or CC-core density) extracted in Tier 1 are sufficiently correlated with multimeric configurations to serve as valid filters for a 6,000-sequence library.
3. **Ground Truth Completeness:** It assumes that the absence of the MADA motif coordinates in experimental PDBs (9FP6, 9CC8) can be overcome by using AF3-generated "reconstituted helpers" as the quantitative baseline for metrics like MADA tilt.
4. **Functional Degeneracy Correlation:** It assumes that structural "novelty" (deviation from NRC2/3/4) correlates directly with the functional transition from helper (pore-former) to sensor/decoy.

#### Reasoning on Feasibility and Effect

1. **AF3 Confidence as a Metric:** Recent evidence [3, 23, 26] confirms that AF3 produces significantly higher ipTM and pTM scores for functional helpers compared to sensors. Using $\Delta ipTM$ (Parameter #9) is highly feasible and likely to be one of the most robust parameters for distinguishing canonical helpers from unconventional scaffolds.
2. **The CC Hydrophobic Core:** The inclusion of Parameter #10 ($\chi_{core}$) is particularly insightful. Preprints from 2025 [14, 20] have explicitly identified this core as a requirement for resistosome formation. This parameter provides a physics-based structural metric that sequence identity alone would miss, directly addressing the goal's preference for structural biology metrics.
3. **Tiered Screening Strategy:** Modeling 6,000 hexamers is computationally intense. The Tier 1 monomer scan (Aperture, MHD, CC-Core) is a highly realistic approach to handle hardware constraints. Research [17] indicates that the NB-ARC configuration (Parameter #3) and MHD motif (Parameter #8) are discernible in monomeric models and indicative of activation potential.
4. **Control Set Calibration:** Phase 0 (using Rx, Bs2, etc.) is essential for setting empirical thresholds. Article [19] shows that sensors like Rx lack the MADA motif but still retain some N-terminal similarities. A data-driven boundary, rather than arbitrary Z-scores, increases the accuracy of identifying "Unconventional" NLRs in the 350 Solanaceae species dataset.

#### Suggested Improvements

1. **Motif-Anchor Numbering:** To ensure Parameters #6 and #8 are automatable across 6,000 diverse sequences, the implementation should explicitly use conserved motif anchors (e.g., the "P-loop" Lysine and the "MHD" Aspartate) rather than fixed sequence indices, which are prone to error due to insertions/deletions.
2. **Frustration/Clash Penalty:** To counter the "Forced Folding" bias of AF3, Parameter #9 should be weighted by a "structural clash" metric. If AF3 forces a sequence into a hexamer with high ipTM but significant van der Waals overlaps, it should be flagged as a modeling artifact rather than a "canonical helper."
3. **Domain Truncation for Multimer Scan:** For Tier 2, modeling full-length NLRs (often >1,000 amino acids) as hexamers (~6,000 residues) exceeds the token limits of most AF3 implementations. The idea should specify using truncated CC-NB-ARC modules for the multimer scan to ensure feasibility.

#### Overall Impact Potential

The SNI-V2 framework is a technically sophisticated and biologically grounded approach to a major problem in plant immunology: functional annotation of the NLRome. By moving beyond sequence homology to a multi-parameter structural index, it leverages the specific mechanical differences (like the $10^\circ$ NB-ARC aperture shift and the CC hydrophobic core density) that distinguish active immune "executors" from sensors.

The inclusion of Parameter #10 and the tiered screening approach make this idea highly feasible and computationally efficient. The shift from hypothetical thresholds to empirical calibration using known sensors/helpers [Phase 0] significantly reduces the risk of false positives. This methodology has the potential to transform how researchers identify novel immune components in crop species, moving the field toward a "structure-first" functional genomics paradigm.

**Conclusion:** The idea is highly likely to have a significant impact on the field, stimulating new directions in high-throughput structural immunology and providing a robust toolkit for breeding programs.

Answer: 8

References:

[1] [An N-terminal motif in NLR immune receptors is functionally conserved across distantly related plant species - PMC](https://pmc.ncbi.nlm.nih.gov/articles/PMC6944444/)

[2] [Accurate Prediction of Protein Complex Stoichiometry by Integrating AlphaFold3 and Template Information - PMC](https://pmc.ncbi.nlm.nih.gov/articles/PMC11761747/)

[3] [Can AI modeling of protein structures distinguish between sensor and helper NLR immune receptors? - PMC](https://pmc.ncbi.nlm.nih.gov/articles/PMC12409091/)

[4] [A disease resistance protein triggers oligomerization of its NLR helper into a hexameric resistosome to mediate innate immunity](https://www.ncbi.nlm.nih.gov/pmc/articles/PMC11540030/)

[5] [NLR immune receptors: structure and function in plant disease resistance](https://www.ncbi.nlm.nih.gov/pmc/articles/PMC10586772/)

[6] [Plant NLR immune receptor Tm-2 2 activation requires NB-ARC domain-mediated self-association of CC domain](https://www.ncbi.nlm.nih.gov/pmc/articles/PMC7205312/)

[7] [A disease resistance protein triggers oligomerization of its NLR helper into a hexameric resistosome to mediate innate immunity](https://www.biorxiv.org/content/10.1101/2024.06.18.599586v1)

[8] [A disease resistance protein triggers oligomerization of its NLR helper into a hexameric resistosome to mediate innate immunity](https://www.ncbi.nlm.nih.gov/pmc/articles/PMC11540030/)

[9] [RCSB PDB - 9FP6: Structure of the NbNRC2 hexameric resistosome](https://www.rcsb.org/structure/9fp6)

[10] [A disease resistance protein triggers oligomerization of its NLR helper into a hexameric resistosome to mediate innate immunity](https://www.biorxiv.org/content/10.1101/2024.06.18.599586v1)

[11] [Diversification of the “EDVID” packing motif underpins structural and functional variation in plant NLR coiled-coil domains](https://www.biorxiv.org/content/10.1101/2025.06.01.657260v1)

[12] [Assessment of Self-Activation and Inhibition of Wheat Coiled-Coil Domain Containing NLR Immune Receptor Yr10 CG](https://www.ncbi.nlm.nih.gov/pmc/articles/PMC11768854/)

[13] [A disease resistance protein triggers oligomerization of its NLR helper into a hexameric resistosome to mediate innate immunity](https://www.ncbi.nlm.nih.gov/pmc/articles/PMC11540030/)

[14] [A hydrophobic core in the coiled-coil domain essential for NRC resistosome function | bioRxiv](https://www.biorxiv.org/content/10.1101/2025.01.21.634219v1.full-text)

[15] [The activated plant NRC4 immune receptor forms a hexameric resistosome](https://www.biorxiv.org/content/10.1101/2023.12.18.571367v1)

[16] [A disease resistance protein triggers oligomerization of its NLR helper into a hexameric resistosome to mediate innate immunity](https://www.biorxiv.org/content/10.1101/2024.06.18.599586v1)

[17] [Two genetically linked Arabidopsis TIR-type NLRs are required for immunity and interact with NLRs encoded in a segmentally duplicated genomic region](https://www.biorxiv.org/content/10.1101/2025.04.25.650432v1)

[18] [Protein Quaternary Structures in Solution are a Mixture of Multiple forms](https://www.biorxiv.org/content/10.1101/2022.03.30.486392v1)

[19] [An N-terminal motif in NLR immune receptors is functionally conserved across distantly related plant species | eLife](https://elifesciences.org/articles/49956)

[20] [A hydrophobic core in the coiled-coil domain is essential for NRC resistosome function | bioRxiv](https://www.biorxiv.org/content/10.1101/2025.01.21.634219v3.full-text)

[21] [The activated plant NRC4 immune receptor forms a hexameric resistosome](https://www.biorxiv.org/content/10.1101/2023.12.18.571367v2)

[22] [A hydrophobic core in the coiled-coil domain is essential for NRC resistosome function](https://www.biorxiv.org/content/10.1101/2025.01.21.634219v2)

[23] [Can AI modelling of protein structures distinguish between sensor and helper NLR immune receptors?](https://www.biorxiv.org/content/10.1101/2024.11.24.625045v1)

[24] [A hydrophobic core in the coiled-coil domain is essential for NRC resistosome function](https://www.biorxiv.org/content/10.1101/2025.01.21.634219v3)

[25] [Deep learning facilitates precise identification of disease-resistance genes in plants](https://www.biorxiv.org/content/10.1101/2024.09.26.615248v1)

[26] [Assessing scoring metrics for AlphaFold2 and AlphaFold3 protein complex predictions](https://www.biorxiv.org/content/10.1101/2025.04.16.648930v1.full.pdf)

[27] [Fig. S6.Hexameric AlphaFold 3 predictions for previously reported NLR... | Download Scientific Diagram](https://www.researchgate.net/figure/Fig-S6Hexameric-AlphaFold-3-predictions-for-previously-reported-NLR-pairs-The-amino_fig7_387154533)

[28] [Can AI Modelling of Protein Structures Distinguish Between Sensor and Helper NLR Immune Receptors? - Article (Preprint v1) by AmirAli Toghani et al. | Qeios](https://www.qeios.com/read/HV8F2C)

[29] [Helper NLRs produce higher AlphaFold 3 pTM scores than their paired... | Download Scientific Diagram](https://www.researchgate.net/figure/Helper-NLRs-produce-higher-AlphaFold-3-pTM-scores-than-their-paired-sensors-a-Bar-plot_fig1_387154533)

[30] [RCSB PDB - 9CC8: Hexameric state of the NRC4 resistosome](https://www.rcsb.org/structure/9cc8)

[31] [NRC Immune receptor networks show diversified hierarchical genetic architecture across plant lineages - PMC](https://pmc.ncbi.nlm.nih.gov/articles/PMC11371147/)

[32] [NRGRank: Coarse-grained structurally-informed ultra-massive virtual screening](https://www.biorxiv.org/content/10.1101/2025.02.17.638675v1)

[33] [Insight into the structure and molecular mode of action of plant paired NLR immune receptors - PMC](https://pmc.ncbi.nlm.nih.gov/articles/PMC9528088/)

[34] [Jurassic NLR: conserved and dynamic evolutionary features of the atypically ancient immune receptor ZAR1](https://www.biorxiv.org/content/10.1101/2020.10.12.333484v3)

[35] [AlphaFold2 enables accurate deorphanization of ligands to single-pass receptors](https://www.biorxiv.org/content/10.1101/2023.03.16.531341v2)

[36] [What AlphaFold 3 struggles with | AlphaFold](https://www.ebi.ac.uk/training/online/courses/alphafold/alphafold-3-and-alphafold-server/introducing-alphafold-3/what-alphafold-3-struggles-with/)

[37] [Resurrection of plant disease resistance proteins via helper NLR bioengineering - PMC](https://pmc.ncbi.nlm.nih.gov/articles/PMC10156107/)

[38] [Functional divergence shaped the network architecture of plant immune receptors | bioRxiv](https://www.biorxiv.org/content/10.1101/2023.12.12.571219v1)

[39] [A helper NLR targets organellar membranes to trigger immunity](https://www.biorxiv.org/content/10.1101/2024.09.19.613839v1)

[40] [AlphaFold2, AlphaFold-Multimer, AlphaFold3](https://310.ai/blog/alphafold2-alphafold-multimer-alphafold3)

[41] [The nucleotide binding domain of NRC-dependent disease resistance proteins is sufficient to activate downstream helper NLR oligomerization and immune signaling | bioRxiv](https://www.biorxiv.org/content/10.1101/2023.11.30.569466v1.full-text)

[42] [Structural basis of NLR activation and innate immune signalling in plants](https://www.ncbi.nlm.nih.gov/pmc/articles/PMC8813719/)

[43] [Sensor NLR immune proteins activate oligomerization of their NRC helper](https://www.biorxiv.org/content/10.1101/2022.04.25.489342v1)

[44] [Protein-Nucleic Acid Complex Modeling with Frame Averaging Transformer](https://arxiv.org/abs/2406.09586)

[45] [How to assess the quality of AlphaFold 3 predictions | AlphaFold](https://www.ebi.ac.uk/training/online/courses/alphafold/alphafold-3-and-alphafold-server/how-to-assess-the-quality-of-alphafold-3-predictions/)

[46] [Activation and Autoinhibition Mechanisms of NLR Immune Receptor Pi36 in Rice](https://www.ncbi.nlm.nih.gov/pmc/articles/PMC11242311/)

[47] [The NRC0 gene cluster of sensor and helper NLR immune receptors is functionally conserved across asterid plants](https://www.biorxiv.org/content/10.1101/2023.10.23.563533v1)

[48] [The activated plant NRC4 immune receptor forms a hexameric resistosome](https://www.biorxiv.org/content/10.1101/2023.12.18.571367v2)

[49] [Benchmarking AlphaFold3’s protein-protein complex accuracy and machine learning prediction reliability for binding free energy changes upon mutation](https://arxiv.org/html/2406.03979v1)

[50] [The N-terminal domains of NLR immune receptors exhibit structural and functional similarities across divergent plant lineages - PMC](https://pmc.ncbi.nlm.nih.gov/articles/PMC11218826/)

[51] [NLRome dataset from 124 genomes of plants in the Solanaceae family](https://spiral.imperial.ac.uk/bitstreams/ae04bf0d-5dd3-4d04-b1ce-bbdae1d35f4a/download)

[52] [NRC immune receptor networks show diversified hierarchical genetic architecture across plant lineages | bioRxiv](https://www.biorxiv.org/content/10.1101/2023.10.25.563953v2.full-text)

**Motivation:**

$\def\mathcal#1{\mathit{#1}}\def\mathscr#1{\mathit{#1}}$

#### Motivation and Evidence

The primary motivation for the **SNI-V2 Framework** is to provide a quantitative, physics-based method to distinguish functional "helper" NRCs from "sensors" or "modulators" (like NRCX) within large datasets (6,000+ sequences). While sequence-based motifs are often used, they fail to account for structural integrity or mechanical competence.

**Key Evidence and Differentiators:**

- **CC-Core Packing Density ($\chi_{core}$):** This 10th parameter is a unique differentiator. Evidence from 2025 research identifies a conserved hydrophobic core ($\alpha$2-$\alpha$4 helices) essential for NRC resistosome function. This provides a physics-based metric to identify "degenerate" sensors that sequence-based motifs might miss.
- **Stoichiometry Delta ($\Delta \text{ipTM}$):** Utilizing the difference in predicted interface accuracy between $C_5$ and $C_6$ models is a validated methodology for predicting quaternary structure and identifying functional divergence.
- **Tiered Screening Architecture:** This makes the hypothesis special by resolving the computational bottleneck of high-throughput AlphaFold 3 (AF3) modeling, filtering the library through inexpensive monomeric scans before committing to multimeric modeling.

#### Explanatory Power

The framework provides mechanical explanations for why certain NLRs fail to trigger cell death despite having "canonical" architectures:

- **Pore Failure:** It explains why atypical members like NRCX lack cell death capacity. Observation: *"Atypical members like NRCX possess the standard tripartite architecture and a predicted MADA sequence but lack the capacity to trigger cell death and act as modulators."* The hypothesis posits that these proteins have a degenerated hydrophobic core ($\chi_{core}$) that destabilizes the CC barrel, making it structurally incompetent for pore formation.
- **MADA Mechanical Thresholds:** It quantifies the "death switch" failure. Observation: *"The N-terminal MADA motif is essential for triggering cell death... but mutations in this motif do not necessarily prevent the formation of the initial resistosome complex."* The hypothesis explains this through **Parameter #2 (MADA Helix Tilt)** and **Parameter #7 (Hydrophobic Moment Alignment)**, suggesting the switch fails because the mechanical orientation prevents the funnel from correctly interacting with the lipid bilayer.

#### Counterarguments

- **Ground Truth Misidentification:** A critical flaw identified in reviews is the use of PDB 9RI9 (SlNRC3) to define hexameric baselines. 9RI9 is established as a **resting-state homodimer**; using it to define hexameric aperture or symmetry would yield mathematically invalid novelty thresholds.
- **Observability Gap:** Parameters #2 and #7 rely on residues 1–16 of the $\alpha$1-helix. These are often disordered or missing in experimental PDBs (9FP6, 9CC8, 9RI9), meaning the framework cannot be measured against experimental ground truths without prior AF3 reconstruction.
- **Aggregation Logic:** The framework defines 10 heterogeneous metrics but lacks a formalized mathematical formula to aggregate these into a single ranking score, complicating the definition of "novelty" when metrics diverge in opposite directions.

#### Why Choose This Hypothesis?

**Choose because:** It is the only framework that specifically targets the **structural integrity of the CC-barrel core** as a discriminator, moving beyond simple MADA sequence motifs. Its tiered approach is uniquely suited for the high-throughput demands of the 6,000-sequence Solanaceae dataset.

**Avoid because:** It currently suffers from **fundamental baseline errors** regarding protein states (dimer vs. hexamer) and relies on geometric metrics (BSA, Curvature) that are notoriously difficult to automate across complex NLR geometries without specialized algorithmic solutions.

**Coherence:**

$\def\mathcal#1{\mathit{#1}}\def\mathscr#1{\mathit{#1}}$

No incoherence found.

**Deep verification:**

$\def\mathcal#1{\mathit{#1}}\def\mathscr#1{\mathit{#1}}$

The hypothesis that the SNI-V2 framework can accurately identify and categorize unconventional NLR proteins is potentially incorrect due to several biological, technical, and methodological flaws. The primary reasons are summarized below:

#### 1. Factually Incorrect Metrics and Units

- **Misapplication of Physics Units:** The framework defines **CC-Core Packing Density ($\chi_{core}$)** using the unit $\text{kcal/mol}\cdot\text{Å}^2$. This is scientifically inaccurate; packing density is a geometric or spatial ratio, whereas the proposed unit describes energy density or a spring constant (restraint). This suggests a fundamental misunderstanding of the physical parameter being measured.
- **Inaccurate Sequence Data:** The framework claims to process "exactly 6,000 sequences" in its first tier. Evidence suggests this number is inaccurate, as the actual sequence count for the target research area (NRC evolution in Solanaceae) is approximately 1,092. The 6,000 figure appears to be a misattribution of AlphaFold sampling literature rather than a precise dataset count.

#### 2. Biological and Functional Invalidity

- **Irrelevance of MHD-Sensory Coupling:** Parameter 8 (**MHD-Sensory Coupling**) is considered "extremely unlikely" to provide meaningful data for helper NLRs (NRCs). Because NRCs are activated by separate sensor proteins rather than internal ligand perception, the distance between the MHD motif and LRR loops is likely under low evolutionary pressure. Including this as a core metric would likely introduce significant statistical noise.
- **Over-simplification of Linker Flexibility:** The **Linker Span (Parameter 6)** assumes that a short physical distance between domains prevents structural rearrangement. However, the high flexibility of NLR linkers in vivo means that a static structural model may not reflect a protein's true functional capacity to transition into an active state.

#### 3. Methodological Risks in the Implementation Pipeline

- **Tier 1 Filtering Flaws:** The framework relies on a "Tier 1" monomeric scan to filter out candidates before multimeric modeling. There is a significant risk that monomeric markers (like CC-core packing or MHD coupling) are not sufficiently predictive of complex multimeric transitions. This could lead to the premature exclusion of viable "unconventional" candidates that only reveal their functional potential when modeled as multimers (hexamers/pentamers).
- **Questionable Causal Rationale:** The framework assumes that low CC-core packing density ($\chi_{core}$) definitively identifies a "degenerate" non-pore scaffold. If the underlying measurement and units for $\chi_{core}$ are fundamentally flawed (as noted in Claim 1), the causal link between this metric and the mechanical stability of the resistosome is undermined.
