## Supplementary Data S6 for "AI-guided discovery of atypical protein assemblies": [EXTERNAL] ├░Y╠ê┬o┬1⁄4 Co-scientist - Hypothesis 102303763741952 - Template-Anchored Geometric Divergence for Quantifying NRC-NLR Structural Novelty via AlphaFold 3 - For research purposes only - Google confidential.docx

### 🧬 Co-scientist - Template-Anchored Geometric Divergence for Quantifying NRC-NLR Structural Novelty via AlphaFold 3

**About**: *This is an experimental system for generating novel and testable hypotheses. The hypotheses are generated by a model and may be wrong. For research purposes only.*

### Proposal

$\def\mathcal#1{\mathit{#1}}\def\mathscr#1{\mathit{#1}}$

#### 1. The Procrustean SNI (Template-Anchored Geometric Divergence)

This approach utilizes a template-anchored alignment to quantify structural divergence. By mathematically aligning every predicted monomer to the rigid "engine" of the canonical 9FP6 hexamer, the Structural Novelty Index (SNI) quantifies the deviation of the protein's domains (CC and LRR) from this hexameric constraint. This defines a fixed coordinate system based on the highly conserved NB-ARC domain.

The following seven quantitative parameters are measured after structurally superimposing the NB-ARC domain of the AlphaFold 3 (AF3) monomer onto the NB-ARC domain of the NbNRC2 Hexamer (PDB: 9FP6, Chain A).

| # | Parameter Name | Unit | Structural Insight | Calculation Method |
| --- | --- | --- | --- | --- |
| **1** | **Hexameric Torque Angle ($\tau_{\alpha4}$)** | Degrees (°) | **The "L126 Sensor".** Measures the specific kink in the $\alpha$4 helix required to fit a 6-fold ring. | Angle between the vector of residues [100-115] and [118-135] (canonical numbering). **NRC2 Baseline:** ~$150^\circ$. **Deviation:** ~$180^\circ$ (straight helix) indicates pentameric-like or scaffold-only geometry. |
| **2** | **Radial Lever Displacement ($\delta_{CC}$)** | Angstroms (Å) | **Pore Formation Potential.** Measures if the CC "arm" can physically reach the central pore axis. | Distance between the centroid of the query's N-terminal tip (res 1-20) and the 9FP6 CC-tip centroid. High values (>15Å) suggest the protein cannot form a canonical channel. |
| **3** | **NB-ARC "Scissor" Aperture ($\Psi_{sci}$)** | Degrees (°) | **Activation State.** The rotational status of the ARC1/ARC2 subdomains relative to the nucleotide pocket. | Angle between the centroids of [NB], [ARC1], and [ARC2]. **Baseline:** Derived from 9FP6. Significant deviation indicates a "locked" or "always-on" state distinct from the canonical switch. |
| **4** | **Keystone Azimuthal Drift ($\phi_{key}$)** | Degrees (°) | **Stoichiometry Predictor.** Measures if the LRR domain fits the $60^\circ$ hexameric arc or drifts toward $72^\circ$ (pentameric) geometry. | Lateral angular deviation of the LRR centroid relative to the 9FP6 LRR centroid in the XY plane. Drifts $>8^\circ$ signal unconventional oligomerization (e.g., ZAR1-like pentamers). |
| **5** | **Switch-Groove Integrity Vector ($\Delta_{nuc}$)** | Angstroms (Å) | **Nucleotide Competency.** Measures the 3D integrity of the ATP binding pocket. | Euclidean distance between the P-loop Lysine and MHD-motif Histidine relative to the 9CC8 (NRC4) active state. Large shifts imply the NLR may be a non-enzymatic scaffold (decoy). |
| **6** | **Interface Hydrophobic Mismatch ($\chi_{hyd}$)** | Scalar (0-1) | **Assembly Glue.** Assessing if the surface "sticky patches" align with the neighbor protomer. | Dot product of the hydrophobic moment vector of the query's $\alpha$2-helix with the corresponding vector in 9FP6. Low scores (<0.5) indicate the interface is biochemically incompatible with hexamerization. |
| **7** | **Confidence-Weighted Stability ($W_{conf}$)** | Score (0-100) | **False Positive Filter.** Ensures high SNI scores aren't just artifacts of disordered predictions. | Average pLDDT score of the residues defining parameters 1, 2, and 4. The final SNI is penalized if $W_{conf} < 70$. |

#### 2. Implementation Logic

1. **Generate Monomers:** Run AlphaFold 3 on the sequences to generate monomeric structures.
2. **The Procrustean Alignment:** Rigid-body align residues 160–400 (the NB-ARC engine) of each monomer to residues 160–400 of **9FP6 Chain A**.
3. **Calculate & Index:** Extract the 7 parameters in this fixed coordinate system.
   - **Low SNI (< 5):** Canonical NRC (Like NRC2/3/4).
   - **High SNI (> 15):** Unconventional.
     - *High $\tau_{\alpha4}$ & $\phi_{key}$:* Likely Pentameric (ZAR1-like).
     - *High $\delta_{CC}$ & $\Delta_{nuc}$:* Likely Non-Channel Scaffold or Decoy.

#### 3. Connection to Ground Truth

- **Monomer Fidelity:** By aligning the NB-ARC engine to the hexamer template, the index reveals the implications of the monomer's shape (e.g., CC-arm orientation relative to the pore) without the computational requirement of modeling the full oligomer.
- **Leveraging Reference PDBs:** PDBs 9FP6 and 9RI9 serve as the structural "bed" for alignment, while 9CC8 provides the baseline for the active state "switch."
- **Mechanical Metrics:** The Torque Angle and Lever Displacement provide physical measurements to distinguish proteins that cannot function as canonical hexamers, identifying those with different gear ratios or assembly requirements.

### Reviews summary

$\def\mathcal#1{\mathit{#1}}\def\mathscr#1{\mathit{#1}}$

Positive aspects:

- **Mechanistic Structural Basis:** The index is grounded in specific, validated structural determinants of hexamerization, particularly the "L126 kink" in the $\alpha$4 helix. This parameter directly distinguishes canonical NRC hexamers from pentameric resistosomes, making the index biologically relevant rather than just a mathematical abstraction.
- **Computational Scalability:** By utilizing monomeric AlphaFold 3 predictions and rigid-body alignment to a template, the method avoids the high computational cost and memory errors associated with modeling full hexameric complexes. This makes screening 6,000 sequences feasible within a reasonable timeframe.
- **Nuanced Functional Differentiation:** Unlike simple sequence identity or RMSD checks, the SNI identifies the physical reasons for functional divergence, such as the "Keystone Azimuthal Drift," which predicts whether a monomer's geometry is compatible with a 60-degree (hexameric) or 72-degree (pentameric) arc.
- **Robust Quality Filtering:** The inclusion of confidence-weighted stability scores (pLDDT) ensures that high novelty scores are not merely the result of disordered or poorly modeled regions, effectively reducing false positives in a high-throughput pipeline.

Negative aspects:

- **Inaccurate Numerical Baselines:** The proposed baseline value for the Torque Angle ($150^\circ$) contradicts the provided ground truth data, which shows a value closer to $168^\circ$. Using the incorrect baseline would lead to systematic errors in classifying canonical versus unconventional proteins.
- **Reliance on Unresolved Structural Data:** Parameter 2 (Radial Lever Displacement) depends on the positions of residues 1–20, which are often disordered or missing in the experimental reference structures (PDB 9FP6 and 9CC8). This makes it impossible to establish a reliable, ground-truth-anchored baseline for this specific metric.
- **Conformational State Sensitivity:** The index assumes AlphaFold 3 will predict monomers in an "active" or "primed" state. Without explicitly requiring the inclusion of ligands (ATP/Mg²⁺) during modeling, the protein may default to an autoinhibited conformation that cannot be accurately compared to the active-state hexamer template.
- **Implementation Complexity for Motifs:** Automating the "Switch-Groove Integrity" calculation is difficult because mapping sequence-based motifs (like the P-loop or MHD motif) to three-dimensional coordinates across 6,000 diverse sequences is non-trivial and prone to alignment errors.

#### Research contacts

$\def\mathcal#1{\mathit{#1}}\def\mathscr#1{\mathit{#1}}$

Based on the provided research articles, the following researchers are suggested as experts to review the proposed Structural Novelty Index (SNI). They are listed in order of relevance to the specific technical aspects of the hypothesis (structural modeling, NRC hexamer/pentamer differentiation, and quantitative assessment).

#### 1. Michael W. Webster & J. Madhuprakash

- **Justification:** These researchers are the primary experts on the structure of the **NbNRC2 hexameric resistosome** (PDB ID: 9FP6), which serves as the core "Ground Truth" for your SNI. Their work directly addresses the structural basis of hexamer versus pentamer formation and the use of AlphaFold 3 to predict activated CC-NLR oligomers. They are uniquely qualified to evaluate whether the SNI parameters (such as the hexameric torque angle and radial lever displacement) accurately capture the divergence from the canonical 9FP6 template.
- **Relevant Article:** *"A disease resistance protein triggers oligomerization of its NLR helper into a hexameric resistosome to mediate innate immunity"* (Abstracts 3, 4, 5, 7, 8).

#### 2. AmirAli Toghani & Sophien Kamoun

- **Justification:** These authors have specialized in using **AlphaFold 3 to distinguish between sensor and helper NRCs** by analyzing structural metrics like pTM and ipTM. Their expertise in high-throughput structural modeling and the classification of the NRC network makes them ideal reviewers for the "scope" of your goal: applying the SNI to the ~6,000 sequences in the NRC family and distinguishing unconventional NLRs from canonical helpers.
- **Relevant Articles:** *"Can AI modeling of protein structures distinguish between sensor and helper NLR immune receptors?"* (Abstract 2); *"A hierarchical immune receptor network in lettuce reveals contrasting patterns of evolution in sensor and helper NLRs"* (Abstracts 10, 11, 12).

#### 3. Alexander Förderer

- **Justification:** Förderer focuses on the **structural variation of coiled-coil (CC) domains** and the conserved **EDVID motif**. Since your SNI includes parameters for "N-terminal CC-domain variations" and "residue conservation patterns," his expertise in how these motifs relate to oligomeric assembly and canonical CC-NLR function is critical for validating the biological relevance of your geometric parameters.
- **Relevant Article:** *"Diversification of the 'EDVID' packing motif underpins structural and functional variation in plant NLR coiled-coil domains"* (Abstract 1).

#### 4. Maya Topf

- **Justification:** The goal requires a **quantitative index (SNI)** with measurable parameters. Dr. Topf’s research focuses on benchmarking and developing **scoring metrics (e.g., DockQ, ipTM, C2Qscore) for AlphaFold 3 and protein complex predictions**. She is the ideal reviewer to assess the statistical robustness and reliability of the 5–10 quantitative parameters you have proposed for high-throughput screening.
- **Relevant Article:** *"Assessing scoring metrics for AlphaFold2 and AlphaFold3 protein complex predictions"* (Abstract 6).

### Appendix:

**All reviews:**

**Correctness:**

$\def\mathcal#1{\mathit{#1}}\def\mathscr#1{\mathit{#1}}$

This review evaluates the **Structural Novelty Index (SNI)**, a proposed metric designed to quantify the divergence of NRC-NLRs from the canonical hexameric architecture represented by NRC2/3/4.

#### 1. Related Article Abstracts

The following articles are most relevant to the evaluation of the SNI:

#### 2. Detailed Assumptions

1. **Rigid-Engine Alignment:** The NB-ARC domain (residues 160–400) is structurally conserved enough to serve as a fixed coordinate system for Procrustean alignment across 6,000 diverse NRC sequences.
2. **Active-State Monomer Fidelity:** AlphaFold 3 monomer predictions of NRCs reflect a "primed" or "active" conformation of the CC and LRR domains relative to the NB-ARC, allowing for meaningful distance/angle measurements when compared to an active hexamer template (9FP6).
3. **Stoichiometric Geometry:** The "Torque Angle" ($\tau_{\alpha4}$) and "Azimuthal Drift" ($\phi_{key}$) in a monomer are reliable predictors of the protein's eventual oligomeric stoichiometry (5-fold vs 6-fold).
4. **Residue Resolvability:** The N-terminal residues (1–20) and specific motifs (P-loop/MHD) are consistently modeled or present in reference structures to establish the SNI baseline.

#### 3. Comparison with Knowledge Base

- **Parameter 1 (Torque Angle):** The idea proposes a $150^\circ$ baseline for NRC2. However, the **Knowledge Base audit** calculated a value of **$167.71^\circ$** for 9FP6. This represents a significant deviation in the foundational baseline.
- **Parameter 2 (Radial Displacement):** The idea relies on residues 1–20. The **Knowledge Base audit** states these residues are **missing/unresolved** in ground truth PDBs 9FP6 and 9CC8, making it impossible to calculate the proposed baseline from experimental data.
- **Parameter 5 (Switch-Groove):** The audit failed to map sequence motifs to PDB entities due to "auth_asym_id" errors, suggesting that simple sequence-to-structure mapping for these motifs is non-trivial in a high-throughput context.
- **Domain Resemblance:** Madhuprakash et al. [3] confirm that individual NRC2 protomers resemble ZAR1/Sr35, supporting the use of NB-ARC as an alignment anchor.

#### 4. Reasoning about Correctness of Assumptions

- **Assumption 1 (Rigid Engine):** **True.** The NB-ARC is the most conserved part of NLRs and is known to be the "structural engine" for oligomerization.
- **Assumption 2 (Active Monomer):** **Questionable.** AF3 often predicts the "most stable" form, which may be the autoinhibited state. Without adding ATP as a ligand or using autoactive mutations (e.g., MHD->MHV), the monomer might not align correctly to the 9FP6 active-state template.
- **Assumption 3 (Stoichiometric Geometry):** **Highly Plausible.** Madhuprakash et al. [3] specifically highlight the $\alpha4$ kink as the differentiator between pentamers and hexamers.
- **Assumption 4 (Resolvability):** **False.** As noted in the KB audit, the N-terminus (res 1–20) is highly flexible and often missing in cryo-EM, making it a poor choice for a fixed geometric baseline.

#### 5. Strength of Evidence

- **Direct Evidence:** Extremely strong for Parameter 1 and 3. Literature [3, 9] specifically identifies the L126 kink and NB-ARC interdomain angles (85° vs 75°) as the defining differences between NRC hexamers and others.
- **Indirect Evidence:** Strong for the overall utility of structure-based indices (Toghani [2], Pai [10]) in distinguishing sensors from helpers.
- **Weakness:** The specific numerical baselines provided in the idea (e.g., $150^\circ$) are contradicted by the provided PDB validation logs ($167.7^\circ$).

#### 6. Suggested Improvements

1. **Recalibrate Baselines:** Update the Torque Angle baseline to $\sim 168^\circ$ for NRC2 based on 9FP6 coordinates.
2. **Redefine Parameter 2:** Instead of residues 1–20 (often missing), use the centroid of the $\alpha2-\alpha3$ helices of the CC domain, which are structurally resolved in 9FP6.
3. **Ligand-Gated Modeling:** Specify that AF3 runs must include **ATP/Mg²⁺** to ensure the monomer adopts the active-state configuration required for Procrustean alignment to the 9FP6 hexamer.
4. **Incorporate the EDVID-R Clamp:** Add a parameter measuring the distance between the EDVID motif (CC) and the R-cluster (LRR) to capture CC-LRR interface integrity.

#### 7. Goal Requirement Assessment

- **5–10 Parameters:** Yes (7 defined).
- **Measurable from PDB/AF3:** Yes.
- **Distinguish unconventional NRCs:** Yes, the parameters specifically target the 6-fold vs 5-fold and active vs decoy differences.
- **Leverage 9FP6, 9RI9, 9CC8:** Yes, though 9RI9 (dimer) is less utilized than 9FP6.
- **Quantitative/High-Throughput:** Yes, geometric calculations on monomers are automatable.

#### 8. Final Reasoning and Recommendation

The idea is **highly plausible** and scientifically grounded in the most recent structural biology of the NRC family. The identification of the $\alpha4$ kink (L126) as a primary metric is a sophisticated and direct application of the ground truth data. However, the idea suffers from **numerical inaccuracies** (baseline mismatch) and **practical data gaps** (reliance on missing N-terminal residues).

The "Procrustean" approach is much more powerful than simple sequence identity or RMSD because it isolates *why* a protein might fail to hexamerize (e.g., a "straight" $\alpha4$ helix would cause steric clashes). Despite the calibration errors, the structural logic is sound.

**Recommendation:** **Test the idea**, provided that the baselines are recalibrated using the provided PDB validation logs and the AF3 protocol is adjusted to include nucleotide ligands.

Answer: 7

**Novelty:**

$\def\mathcal#1{\mathit{#1}}\def\mathscr#1{\mathit{#1}}$

#### Related Article Abstracts

1. **[3] A disease resistance protein triggers oligomerization of its NLR helper into a hexameric resistosome to mediate innate immunity** – Detailed structural comparison between hexameric (NRC2) and pentameric (ZAR1/Sr35) resistosomes, highlighting interdomain angles and the $\alpha$4 helix kink.
2. **[2] Can AI modeling of protein structures distinguish between sensor and helper NLR immune receptors?** – Explores using AlphaFold 3 (AF3) confidence scores (pTM/ipTM) to distinguish between canonical helpers and unconventional sensors.
3. **[10] A hierarchical immune receptor network in lettuce reveals contrasting patterns of evolution in sensor and helper NLRs** – Uses AF3 metrics to show that sensor NLRs (unconventional) fail to form stable hexameric resistosome models compared to helpers.
4. **[1] Diversification of the “EDVID” packing motif underpins structural and functional variation in plant NLR coiled-coil domains** – Discusses the EDVID motif as a predictor of canonical function and structural variations in CC-domains.
5. **[13] Structure–function analyses of coiled-coil immune receptors define a hydrophobic module for improving plant virus resistance** – Analyzes CC-domain hydrophobic grooves and their role in oligomerization and activity.
6. **[15] An N-terminal motif in NLR immune receptors is functionally conserved across distantly related plant species** – Identifies the MADA motif as a "death switch" marker for canonical CC-NLRs.
7. **[14] Structure of the activated Roq1 resistosome directly recognizing the pathogen effector XopQ** – Provides a baseline for unconventional (tetrameric) stoichiometry in TNL-NLRs.
8. **[8] RCSB PDB - 9FP6: Structure of the NbNRC2 hexameric resistosome** – The primary ground truth for the canonical hexameric assembly.
9. **[6] Assessing scoring metrics for AlphaFold2 and AlphaFold3 protein complex predictions** – Discusses the utility of interface-specific scores (ipTM, pDockQ2) for evaluating predicted protein complexes.
10. **[7] A disease resistance protein triggers oligomerization of its NLR helper into a hexameric resistosome to mediate innate immunity (bioRxiv)** – Pre-print version of [3] providing additional data on AF3's ability to model NRC hexamers vs. pentamers.

#### 1. Already Explored Aspects

- **Structural differences between hexamers and pentamers:** The "kink" in the $\alpha$4 helix (centered around L126) and the 10° widening of the NB-ARC interdomain angle in NRC2 compared to ZAR1 are explicitly described in **[3], [5], and [9]**.
- **AF3 for functional classification:** Using AF3 confidence metrics (pTM, ipTM, pLDDT) to differentiate "helpers" (canonical) from "sensors" (unconventional/degenerate) has been documented in **[2]** and **[10]**.
- **Pore diameter and N-terminal displacement:** The observation that canonical NRCs have wider CC pores (~19Å) and specific N-terminal α1-helix orientations for membrane insertion is found in **[3] and [4]**.
- **Motif-based integrity:** The requirement of the MADA motif **[15]** and the EDVID/R-cluster interaction **[1]** for canonical resistosome assembly is well-established.
- **The ground truth set:** The structures 9FP6 (NRC2) and 9CC8 (NRC4) are the standard reference models for the activated state **[8, 3]**.

#### 2. Novel Aspects

- **Formalization of the Structural Novelty Index (SNI):** While the structural differences between hexamers and pentamers are known, the synthesis of these observations into a singular, multi-parametric quantitative index (7 specific metrics) for high-throughput screening of a large dataset (~6,000 sequences) is novel.
- **Procrustean NB-ARC Template Alignment:** The specific implementation of using residues 160–400 as a "rigid engine" for coordinate-system alignment to measure the "drift" of the CC and LRR domains is a distinct computational strategy not explicitly detailed in the provided literature.
- **Keystone Azimuthal Drift ($\phi_{key}$):** Quantifying the lateral angular deviation to predict stoichiometry ($60^\circ$ vs $72^\circ$) within a monomeric prediction by projecting its fit into a virtual hexameric arc is a novel geometric approach.
- **Switch-Groove Integrity Vector ($\Delta_{nuc}$):** Using the Euclidean distance between the P-loop and MHD motif relative to the active-state structure (9CC8) to identify non-enzymatic "scaffolds" (decoys) adds a layer of functional prediction beyond simple sequence identity.
- **Torque Angle ($\tau_{\alpha4}$) as a High-Throughput Metric:** Automating the measurement of the "L126 Sensor" to differentiate between straight (pentameric) and bent (hexameric) helical geometry across thousands of AlphaFold models.

#### 3. Novelty Review

The idea is **moderately novel**. The core biological insights—specifically the $\alpha$4 helix kink and NB-ARC widening—are taken directly from recently published cryo-EM comparisons (primarily **[3]**). However, the conversion of these qualitative structural observations into an automated, quantitative scoring system (the SNI) for massive sequence datasets is a significant step forward in computational structural biology.

Current methods (as seen in **[2]** and **[10]**) rely heavily on AlphaFold’s internal confidence scores (pTM/ipTM). The SNI is superior because it measures *why* a protein might be unconventional (e.g., torque angle or lever displacement) rather than just stating that AF3 is "unconfident" in its assembly. The "Keystone Azimuthal Drift" is particularly clever as it essentially predicts stoichiometry from monomeric shapes.

#### 4. Reasoning and Recommendation

Testing this idea is highly recommended. The field currently lacks a standardized mathematical way to define "deviation" from the NRC paradigm. Researchers usually rely on phylogenetic clustering, which often fails to capture structural shifts caused by small but critical mutations (like those in the $\alpha$4 helix).

The SNI would allow for the systematic identification of "structural outliers" in the 6,000-sequence NRC family. These outliers are the most likely candidates for novel immune mechanisms, such as non-pore-forming scaffolds or pentameric NRCs. The use of AlphaFold 3 as the input source makes this pipeline extremely scalable. The strict focus on 5–10 quantitative parameters ensures the index remains interpretable for structural biologists.

Answer: 6

**Feasibility:**

$\def\mathcal#1{\mathit{#1}}\def\mathscr#1{\mathit{#1}}$

#### Related Article Abstracts

1. **[3] A disease resistance protein triggers oligomerization of its NLR helper into a hexameric resistosome to mediate innate immunity** (Madhuprakash et al., 2024): This abstract identifies the 10° interdomain angle difference in NB-ARC and the α4 helix "bend" (L126) that distinguishes hexamers from pentamers, directly validating Parameters 1 and 3.
2. **[10] A hierarchical immune receptor network in lettuce reveals contrasting patterns of evolution in sensor and helper NLRs** (Pai et al., 2025): Demonstrates that AF3 confidence metrics (pTM, ipTM) effectively distinguish helper NRCs (canonical) from sensors (unconventional), supporting Parameter 7.
3. **[2] Can AI modeling of protein structures distinguish between sensor and helper NLR immune receptors?** (Toghani et al., 2025): Confirms that AF3 can capture the structural divergence of N-terminal α1 helices, which is the basis for Parameter 2 (Radial Lever Displacement).
4. **[6] Assessing scoring metrics for AlphaFold2 and AlphaFold3 protein complex predictions** (Genz et al., 2025): Validates the use of ipTM and PAE as the most reliable metrics for interface quality, providing the statistical foundation for the $W_{conf}$ weighting in Parameter 7.
5. **[1] Diversification of the “EDVID” packing motif underpins structural and functional variation...** (Sulkowski et al., 2025): Provides structural context for the interaction between CC and LRR domains, relevant for defining "deviation" in Parameter 4 and 5.
6. **[15] An N-terminal motif in NLR immune receptors is functionally conserved...** (Adachi et al., 2019): Defines the MADA motif and the funnel-shaped "death switch," which provides the biological baseline for the CC-tip centroid metrics in Parameter 2.
7. **[13] Structure–function analyses of coiled-coil immune receptors define a hydrophobic module...** (Wu et al., 2022): Discusses the hydrophobic groove in the CC domain, supporting the feasibility of Parameter 6 (Interface Hydrophobic Mismatch).
8. **[14] Structure of the activated Roq1 resistosome directly recognizing the pathogen effector XopQ** (Martin et al., 2020): Describes alternative (tetrameric) oligomerization and the role of the NB-ARC "open" conformation, helping calibrate the "Scissor" Aperture in Parameter 3.
9. **[8] RCSB PDB - 9FP6: Structure of the NbNRC2 hexameric resistosome**: The primary reference for the hexameric ground truth required for the Procrustean alignment.
10. **[11] A hierarchical immune receptor network in lettuce reveals contrasting patterns of evolution...**: Establishes the existence of the ~6,000 sequence dataset and the necessity of distinguishing helpers from sensors within it.

#### Steps to Test the Idea

1. **Calibration and Scripting:** Develop a Python script (using Biopython or PyMOL’s cmd.align) that performs a rigid-body superposition of NB-ARC residues (160–400) onto PDB 9FP6. Define functions for calculating the seven geometric parameters (angles and centroids).
2. **Go/No-Go Initial Experiment:** Run AlphaFold 3 (AF3) on a control set of 20 proteins: 10 canonical helpers (NRC2/3/4) and 10 known unconventional sensors/helpers (e.g., from Abstract [10]).
   - **Success Criterion:** The SNI score must clearly segregate the two groups (e.g., Mean SNI_canonical < 5; Mean SNI_unconventional > 10) with a p-value < 0.01.
3. **High-Throughput Monomer Generation:** Batch-process the ~6,000 NRC sequences through AF3. To optimize cost, only monomeric structures are required, as the "Procrustean" method measures potential to fit a template rather than actual multimeric folding.
4. **Batch Indexing:** Run the automated structural bioinformatic pipeline on the 6,000 models.
5. **Statistical Mapping:** Generate a distribution of SNI scores across the Solanaceae dataset. Identify clusters of "high-SNI" proteins that deviate significantly from the 9FP6/9RI9/9CC8 baseline.
6. **Validation by Functional Correlation:** Cross-reference high-SNI candidates with existing phylogenetic data. If high-SNI scores correlate with "sensor" status or known non-canonical clades, the index is validated.

#### Reasoning about Feasibility

The testing of this idea is highly feasible due to its **computational efficiency**. Unlike other resistosome screening methods that require modeling full hexamers (which are prone to AF3 memory errors and high costs), this idea uses **monomers** and aligns them to a rigid **template (9FP6)**. This reduces the computational load significantly.

1. **Scalability:** While running 6,000 AF3 monomers is resource-intensive (approx. 1-2 months on a standard GPU cluster), it is a standard bioinformatic task today. The subsequent geometric calculations take only seconds per protein.
2. **Metric Reliability:** Parameters 1, 3, and 4 are based on well-documented structural differences between hexamers and pentamers found in the Ground Truth (e.g., the L126 kink and NB-ARC interdomain angle). Parameter 7 (pLDDT) provides a built-in "trust" filter to handle the inherent flexibility of CC-domains.
3. **Measurability:** All parameters rely on atomic coordinates and confidence scores, which are standard AF3 outputs. No complex molecular dynamics or wet-lab validation is required for the initial "index" construction.

The primary bottleneck is the cost and time of the initial 6,000 AF3 runs. However, the use of a "Procrustean" alignment means the testing avoids the most difficult part of multimeric prediction (interface docking), making the index calculation straightforward once monomers are available.

Answer: 6

**Impact potential:**

$\def\mathcal#1{\mathit{#1}}\def\mathscr#1{\mathit{#1}}$

#### 1. Related Article Abstracts

The following articles are most relevant for assessing the impact and biological validity of the Structural Novelty Index (SNI):

1. **[3] A disease resistance protein triggers oligomerization of its NLR helper into a hexameric resistosome...**: Essential for Parameter 1 ($\tau_{\alpha4}$) and Parameter 4 ($\phi_{key}$); it identifies the specific L126 bend and NB-ARC interdomain angles that differentiate hexamers from pentamers.
2. **[8] RCSB PDB - 9FP6: Structure of the NbNRC2 hexameric resistosome**: Provides the primary ground-truth template for the "Procrustean" alignment and canonical baseline values for all 7 parameters.
3. **[2] Can AI modeling of protein structures distinguish between sensor and helper NLR immune receptors?**: Validates the assumption that AF3 can differentiate NLR functions (sensors vs. helpers) based on structural confidence and "funnel" formation.
4. **[15] An N-terminal motif in NLR immune receptors is functionally conserved...**: Provides the basis for Parameter 2 ($\delta_{CC}$), defining the MADA motif/α1-helix as the "death switch" that must reach the membrane.
5. **[1] Diversification of the “EDVID” packing motif...**: Relevant for Parameter 6; it suggests that the EDVID motif and its packing are predictors of canonical assembly.
6. **[6] Assessing scoring metrics for AlphaFold2 and AlphaFold3 protein complex predictions**: Supports Parameter 7 ($W_{conf}$), highlighting that ipTM and pLDDT are the most reliable metrics for evaluating AF3 interfaces.
7. **[10] A hierarchical immune receptor network in lettuce...**: Highlights the evolutionary divergence of NRC-S (sensors) and NRC-H (helpers), justifying the need for an SNI to identify unconventional sensors that have lost resistosome-forming ability.
8. **[9] A disease resistance protein triggers oligomerization of its NLR helper...**: Provides quantitative differences (e.g., pore diameters 19Å vs 12Å) used to calibrate the Radial Lever Displacement.
9. **[13] Structure–function analyses of coiled-coil immune receptors...**: Discusses the "hydrophobic groove" in the CC domain, supporting the inclusion of Parameter 6 ($\chi_{hyd}$).
10. **[14] Structure of the activated Roq1 resistosome...**: Provides a contrast to the hexameric/pentameric CC-NLRs by showing a tetrameric TIR-NLR, which helps define the "extreme" range of the SNI for non-canonical oligomers.

#### 2. Detailed Assumptions

The idea’s impact potential depends on the following:

- **Monomer-Oligomer Correlation:** A predicted monomer structure (specifically the NB-ARC-CC orientation) contains sufficient geometric information to predict its behavior in a 6-fold symmetry, even without simulating the full hexamer.
- **NB-ARC Rigidity:** The NB-ARC domain (residues 160–400) is structurally conserved enough across the ~6,000 NRC family members to act as a "rigid anchor" for alignment.
- **AF3 Conformational Accuracy:** AlphaFold 3 can accurately predict the "active state" conformation (e.g., the $\alpha$4 helix kink at L126) of a monomer in the presence of appropriate ligands (ATP/Mg²⁺) as described in recent literature [1, 10].
- **Quantifiability of Deviation:** Unconventional function (e.g., becoming a sensor or a decoy) is directly correlated with geometric "drift" from the 9FP6 template.
- **High-Throughput Scalability:** Calculating these 7 geometric parameters is computationally cheaper than running full hexameric AF3 predictions for all 6,000 sequences.

#### 3. Feasibility and Effect of Assumptions

- **Feasibility of Monomer Information:** High. Madhuprakash et al. [3, 7] show that while overall architecture differs between NRC2 (hexamer) and ZAR1 (pentamer), the individual protomers carry the specific angular instructions (e.g., a 10° difference in NB-ARC interdomain angle). This makes the SNI highly realistic.
- **Feasibility of NB-ARC Anchoring:** Very High. The NB-ARC is the defining engine of NLRs. Abstract [11] confirms the NRC superclade is monophyletic and shares this core, making it a robust coordinate system.
- **AF3 Accuracy:** Moderate to High. Abstract [2] and [10] show that AF3 distinguishes helpers from sensors with high confidence. However, if a sequence is highly unconventional, AF3 might produce a disordered prediction ($W_{conf} < 70$), which the idea wisely handles via Parameter 7 as a filter.
- **Computational Scalability:** High. This is a significant advantage. Aligning monomers to a template and calculating angles/centroids is an $O(n)$ operation once models are generated, whereas hexameric AF3 modeling is $O(n^3)$ or worse. This drastically increases the potential scope of the screening.

#### 4. Suggested Improvements

- **Ligand-Enabled Modeling:** Explicitly require AF3 to be run with ATP/Mg²⁺ for all monomers. Recent ground-truth data [5, 8] shows the active state depends on nucleotide binding; without it, the "Scissor Aperture" ($\Psi_{sci}$) may default to an inactive state, yielding false SNI scores.
- **EDVID-R Cluster Distance:** Add an 8th parameter measuring the distance between the EDVID motif (α3-helix) and the Arg-cluster in the LRR. Abstract [1] and [5] emphasize this "clamp" as a critical contact point for stabilization.
- **Interdomain Angle Metric:** Specifically incorporate the "CC-NB-ARC interdomain angle" mentioned in [3]. A 10° increase is a hallmark of hexameric vs. pentameric transition.
- **MADA HMM Integration:** Combine the structural $\delta_{CC}$ with a sequence-based MADA HMM score (from Abstract [15]) to create a hybrid "Execution Competency" sub-score.

#### 5. Overall Impact Potential

The "Procrustean SNI" idea has **high impact potential**. It addresses a major bottleneck in plant pathology: the functional annotation of the massive "dark matter" in NLRome datasets (~6,000 NRC sequences).

**Strengths:**

- **Biological Nuance:** Unlike a simple RMSD, which is a "dumb" metric, this index targets specific, validated mechanical hinges (the L126 kink, the NB-ARC scissor, the LRR azimuthal drift).
- **Feasibility:** It leverages the latest AF3 capabilities and ground-truth structures (9FP6) in a computationally efficient manner.
- **Long-term Implications:** It provides a standardized mathematical framework that can be adapted to other NLR families (e.g., ZAR1-like or TIR-type) by simply swapping the template.

The idea is likely to transform how the Solanaceae NRC network is studied, moving from phylogenetic "best guesses" to structural "functional profiling." It directly addresses the "sensor vs. helper" classification problem identified as a priority in 2024–2025 research [2, 11].

Answer: 8

References:

[1] [Diversification of the “EDVID” packing motif underpins structural and functional variation in plant NLR coiled-coil domains | bioRxiv](https://www.biorxiv.org/content/10.1101/2025.06.01.657260v1.full-text)

[2] [Can AI modeling of protein structures distinguish between sensor and helper NLR immune receptors? - PMC](https://pmc.ncbi.nlm.nih.gov/articles/PMC12409091/)

[3] [A disease resistance protein triggers oligomerization of its NLR helper into a hexameric resistosome to mediate innate immunity](https://www.ncbi.nlm.nih.gov/pmc/articles/PMC11540030/)

[4] [A disease resistance protein triggers oligomerization of its NLR helper into a hexameric resistosome to mediate innate immunity](https://www.ncbi.nlm.nih.gov/pmc/articles/PMC11540030/)

[5] [A disease resistance protein triggers oligomerization of its NLR helper into a hexameric resistosome to mediate innate immunity](https://www.ncbi.nlm.nih.gov/pmc/articles/PMC11540030/)

[6] [Assessing scoring metrics for AlphaFold2 and AlphaFold3 protein complex predictions - PMC](https://pmc.ncbi.nlm.nih.gov/articles/PMC12516916/)

[7] [A disease resistance protein triggers oligomerization of its NLR helper into a hexameric resistosome to mediate innate immunity](https://www.biorxiv.org/content/10.1101/2024.06.18.599586v1)

[8] [RCSB PDB - 9FP6: Structure of the NbNRC2 hexameric resistosome](https://www.rcsb.org/structure/9fp6)

[9] [A disease resistance protein triggers oligomerization of its NLR helper into a hexameric resistosome to mediate innate immunity](https://www.biorxiv.org/content/10.1101/2024.06.18.599586v1)

[10] [A hierarchical immune receptor network in lettuce reveals contrasting patterns of evolution in sensor and helper NLRs](https://www.biorxiv.org/content/10.1101/2025.02.25.639832v1)

[11] [A hierarchical immune receptor network in lettuce reveals contrasting patterns of evolution in sensor and helper NLRs | bioRxiv](https://www.biorxiv.org/content/10.1101/2025.02.25.639832v1.full-text)

[12] [A hierarchical immune receptor network in lettuce reveals contrasting patterns of evolution in sensor and helper NLRs](https://www.biorxiv.org/content/10.1101/2025.02.25.639832v1)

[13] [Structure–function analyses of coiled-coil immune receptors define a hydrophobic module for improving plant virus resistance - PMC](https://pmc.ncbi.nlm.nih.gov/articles/PMC10010612/)

[14] [Structure of the activated Roq1 resistosome directly recognizing the pathogen effector XopQ | bioRxiv](https://www.biorxiv.org/content/10.1101/2020.08.13.246413.full)

[15] [An N-terminal motif in NLR immune receptors is functionally conserved across distantly related plant species | eLife](https://elifesciences.org/articles/49956)

[16] [RCSB PDB - 9CC8: Hexameric state of the NRC4 resistosome](https://www.rcsb.org/structure/9cc8)

[17] [A disease resistance protein triggers oligomerization of its NLR helper into a hexameric resistosome to mediate innate immunity](https://www.ncbi.nlm.nih.gov/pmc/articles/PMC11540030/)

[18] [A helper NLR targets organellar membranes to trigger immunity](https://www.biorxiv.org/content/10.1101/2024.09.19.613839v1)

[19] [Activation of plant immunity through conversion of a helper NLR homodimer into a resistosome](https://www.ncbi.nlm.nih.gov/pmc/articles/PMC11524475/)

[20] [Subfunctionalization of NRC3 altered the genetic structure of the Nicotiana NRC network | PLOS Genetics](https://journals.plos.org/plosgenetics/article?id=10.1371/journal.pgen.1011402)

[21] [Structures of plant resistosome reveal how NLR immune receptors are activated - PMC](https://pmc.ncbi.nlm.nih.gov/articles/PMC9590527/)

[22] [(PDF) LRRpredictor—A New LRR Motif Detection Method for Irregular Motifs of Plant NLR Proteins Using an Ensemble of Classifiers](https://www.researchgate.net/publication/339789683_LRRpredictor-A_New_LRR_Motif_Detection_Method_for_Irregular_Motifs_of_Plant_NLR_Proteins_Using_an_Ensemble_of_Classifiers)

[23] [The N-terminal domains of NLR immune receptors exhibit structural and functional similarities across divergent plant lineages | Request PDF](https://www.researchgate.net/publication/379729781_The_N-terminal_domains_of_NLR_immune_receptors_exhibit_structural_and_functional_similarities_across_divergent_plant_lineages)

**Motivation:**

$\def\mathcal#1{\mathit{#1}}\def\mathscr#1{\mathit{#1}}$

**Coherence:**

$\def\mathcal#1{\mathit{#1}}\def\mathscr#1{\mathit{#1}}$

**Deep verification:**

$\def\mathcal#1{\mathit{#1}}\def\mathscr#1{\mathit{#1}}$
