## Supplementary Data S6 for "AI-guided discovery of atypical protein assemblies": [EXTERNAL] ├░Y╠ê┬o┬1⁄4 Co-scientist - Hypothesis 116914230292830 - Development of a 7-Parameter Structural Novelty Index (SNI) for Identifying Unconventional NRC-NLR Candidates - For research purposes only - Google confidential.docx

### 🧬 Co-scientist - Development of a 7-Parameter Structural Novelty Index (SNI) for Identifying Unconventional NRC-NLR Candidates

**About**: *This is an experimental system for generating novel and testable hypotheses. The hypotheses are generated by a model and may be wrong. For research purposes only.*

### Proposal

$\def\mathcal#1{\mathit{#1}}\def\mathscr#1{\mathit{#1}}$

Based on the structural analysis of the "Ground Truth" Dataset (PDB IDs: 9FP6 [NbNRC2], 9CC8 [NRC4], and 9RI9 [SlNRC3]), the Canonical NRC Resistosome is defined as a homohexameric assembly featuring a characteristic ~18 Å pore and a specific bent α4-helix configuration that enables the 6-fold symmetry.

The Structural Novelty Index (SNI) is defined below. A higher SNI score indicates a deviation from the canonical hexameric paradigm, highlighting unconventional NRC-NLR candidates.

#### **Structural Novelty Index (SNI)**

##### **1. α4-Helix Curvature (Hexamerization Determinant)**

- **Rationale:** In canonical NRCs (9FP6, 9CC8), the CC-domain α4-helix contains a critical bend (e.g., near L126 in NbNRC2) that alleviates steric clashes, allowing six protomers to fit into a ring. Canonical ZAR1-like (pentameric) NLRs have a straight α4-helix.
- **Metric:** Calculate the angle of curvature along the α4-helix backbone.
- **Canonical Baseline:** Angle > 15° (Bent).
- **SNI Contribution:** Low angle (Straight, <10°) increases SNI (suggesting pentameric or non-oligomerizing tendency).

##### **2. MADA Motif Structural Fidelity (Pore Potential)**

- **Rationale:** Canonical helpers (NRC2/3/4) possess a conserved N-terminal MADA motif that forms the α1-helix funnel (pore). Unconventional or sensor NRCs often have degenerated or missing MADA motifs (replaced by non-pore-forming sequences or extended domains).
- **Metric:** RMSD of the N-terminal 20 residues against the 9FP6 α1-helix template + pLDDT score of this region.
- **Canonical Baseline:** RMSD < 2.0 Å, High pLDDT.
- **SNI Contribution:** High RMSD or low pLDDT increases SNI.

##### **3. Hexameric Interface Steric Clash Score**

- **Rationale:** "Forcing" a predicted unconventional NRC protomer into the strict C6 symmetry of the 9FP6/9CC8 scaffold will result in physical overlaps if the protein is not evolutionarily designed for this specific hexameric packing.
- **Metric:** Total volume of atomic overlap (Å³) when 6 copies of the AF3 model are aligned to the 9FP6 symmetry axis.
- **Canonical Baseline:** < 500 Å³ (Minor surface clashes resolvable by flexibility).
- **SNI Contribution:** > 1000 Å³ increases SNI (indicates incompatibility with canonical hexamer).

##### **4. N-terminal Extension (exNT) Length**

- **Rationale:** Some unconventional clades (e.g., in Caryophyllales or specific Asterid lineages) contain extended N-terminal domains (exNT) upstream of the CC domain, unlike the canonical "MADA-first" architecture.
- **Metric:** Number of residues preceding the structural start of the CC domain (or MADA motif).
- **Canonical Baseline:** < 10 residues.
- **SNI Contribution:** > 30 residues increases SNI significantly.

##### **5. Hydrophobic Core Packing Density (CC Domain)**

- **Rationale:** Recent studies (PDB 9CC8) identified a specific hydrophobic core within the CC domain that is essential for NRC4 resistosome stability. Unconventional NLRs may rely on different stabilizing networks.
- **Metric:** Side-chain packing density (or Void Volume) of residues homologous to the NRC4 hydrophobic core.
- **Canonical Baseline:** High packing density (low void volume).
- **SNI Contribution:** Low packing density (loose core) increases SNI.

##### **6. NB-ARC Domain Orientation Angle**

- **Rationale:** The relative orientation of the Nucleotide Binding (NB) domain to the ARC1/ARC2 subdomains is fixed in the activated hexameric state. Unconventional variants may adopt distinct resting-state autoinhibitory locks.
- **Metric:** Euler angle difference of the NB-ARC domain relative to the CC domain when aligned to the 9FP6 protomer.
- **Canonical Baseline:** ±10° from 9FP6 active conformation.
- **SNI Contribution:** Large angular deviation (>20°) increases SNI.

##### **7. Predicted Pore Diameter (Geometric Estimation)**

- **Rationale:** Canonical NRC hexamers form a wide pore (~18–19 Å). Unconventional assemblies may form tighter (pentamer-like ~12 Å) or non-pore structures.
- **Metric:** Calculated minimal inner diameter (Å) of the ring formed by the α1-helices in a modeled oligomer.
- **Canonical Baseline:** 17–19 Å.
- **SNI Contribution:** Deviation (e.g., <14 Å or >22 Å) increases SNI.

#### **Reference & Ground Truth Summary**

- **Reference Set (Canonical):** NbNRC2 (9FP6), SlNRC3 (9RI9), NRC4 (9CC8).
  - *Key Features:* Hexameric, Bent α4-helix, Functional MADA pore, ~18Å diameter.
- **Target Scope:** ~6,000 NRC sequences.
- **Goal:** Filter for High SNI -> Candidates for Novel Oligomeric States (e.g., tetramers, filaments) or Non-Pore Functions (scaffolds).

### Reviews summary

$\def\mathcal#1{\mathit{#1}}\def\mathscr#1{\mathit{#1}}$

#### Research contacts

$\def\mathcal#1{\mathit{#1}}\def\mathscr#1{\mathit{#1}}$

Based on the provided research articles, the following researchers are experts in NRC-NLR structural biology, network evolution, and resistosome architecture. They are listed in order of their focus on specific aspects of your Structural Novelty Index (SNI).

#### 1. **Sophien Kamoun**

- **Justification:** Kamoun is a central figure in defining the NRC superclade and the MADA motif. His work provides the fundamental conceptual framework for understanding the "helper" versus "sensor" divergence. He would be an ideal reviewer for the validity of the SNI in biological and evolutionary terms, particularly the MADA motif fidelity and exNT parameters.
- **Supporting Articles:** *An N-terminal motif in NLR immune receptors is functionally conserved across distantly related plant species* (Abstract 13); *A disease resistance protein triggers oligomerization of its NLR helper into a hexameric resistosome to mediate innate immunity* (Abstracts 4, 6, 8, 10).

#### 2. **Chih-Hang Wu**

- **Justification:** Wu focuses on the hierarchical genetic architecture of NRC networks and the specific divergence between helper and sensor NLRs. His research into the hydrophobic core of the CC domain is directly relevant to your 5th parameter (Hydrophobic Core Packing Density) and explains the functional degeneration in sensor NLRs.
- **Supporting Articles:** *NRC Immune receptor networks show diversified hierarchical genetic architecture across plant lineages* (Abstract 1); *A hydrophobic core in the coiled-coil domain is essential for NRC resistosome function* (Abstracts 7, 9).

#### 3. **Furong Liu**

- **Justification:** Liu is a lead author on the structural characterization of the NRC4 hexameric resistosome (PDB ID 9CC8). Her expertise is critical for reviewing the validity of parameters related to hexameric assembly, LRR packing, and pore diameter (SNI Parameters 3 and 7).
- **Supporting Articles:** *The activated plant NRC4 immune receptor forms a hexameric resistosome* (Abstract 11); *Hexameric state of the NRC4 resistosome* (Abstract 14).

#### 4. **Madhuprakash Jogi**

- **Justification:** Jogi’s work provides a comparative structural analysis of NbNRC2 (PDB ID 9FP6) against pentameric models like AtZAR1. He specifically analyzes the α4-helix bend (L126) and interdomain angles, making him a perfect fit for reviewing your 1st and 6th SNI parameters.
- **Supporting Articles:** *A disease resistance protein triggers oligomerization of its NLR helper into a hexameric resistosome to mediate innate immunity* (Abstracts 4, 6, 8, 10).

#### 5. **Jiorgos Kourelis**

- **Justification:** Kourelis developed the RefPlantNLR database, a comprehensive collection of experimentally validated plant NLRs. His expertise in NLR classification and phylogenetics ensures that the SNI is grounded in a rigorous dataset and applicable to the broader ~6,000 sequence NRC family across different plant species.
- **Supporting Articles:** *RefPlantNLR is a comprehensive collection of experimentally validated plant disease resistance proteins from the NLR family* (Abstract 2); *Jurassic NLR: Conserved and dynamic evolutionary features of the atypically ancient immune receptor ZAR1* (Abstract 12).

### Appendix:

**All reviews:**

**Correctness:**

$\def\mathcal#1{\mathit{#1}}\def\mathscr#1{\mathit{#1}}$

#### Related Articles

1. **[4] A disease resistance protein triggers oligomerization of its NLR helper into a hexameric resistosome to mediate innate immunity**: Direct evidence for hexameric NRC2 (9FP6) vs. pentameric ZAR1 differences, specifically citing the 10° interdomain angle difference and the α4-helix bend (L126).
2. **[11] The activated plant NRC4 immune receptor forms a hexameric resistosome**: Confirms the hexameric state and structural organization of NRC4 (9CC8), providing the baseline for the hexameric resistosome architecture.
3. **[7] A hydrophobic core in the coiled-coil domain is essential for NRC resistosome function**: Identifies a specific hydrophobic core in the NRC4 CC domain required for stability, supporting the inclusion of Metric 5.
4. **[13] An N-terminal motif in NLR immune receptors is functionally conserved across distantly related plant species**: Establishes the importance of the MADA motif (Metric 2) and its absence/degeneration in sensor NLRs.
5. **[1] NRC Immune receptor networks show diversified hierarchical genetic architecture across plant lineages**: Discusses the evolution of noncanonical extended N-terminal domains (exNT) in specific clades, supporting Metric 4.
6. **[8] A disease resistance protein triggers oligomerization of its NLR helper into a hexameric resistosome to mediate innate immunity (Supplementary)**: Demonstrates AlphaFold 3's ability to predict oligomeric states and the N-terminal α1-helix, validating the use of AF3 for the SNI.
7. **[6] Detailed Breakdown of Panel B/C (Madhuprakash et al.)**: Provides specific pore measurements (17–19 Å) and domain orientation differences (85° vs 75°) used to derive Metrics 6 and 7.
8. **[10] A protomer-level structural comparison between the NbNRC2 hexamer and dimer**: Highlights differences in the NB-ARC region orientation between activated and resting states, justifying Metric 6.
9. **[14] RCSB PDB - 9CC8: Hexameric state of the NRC4 resistosome**: Primary data source for the hexameric baseline of NRC4.
10. **[16] Discussion - An N-terminal motif...**: Reinforces that the MADA motif "death switch" applies to helpers but has degenerated in sensors.

#### Detailed Assumptions

1. **Structural Representativeness**: It assumes that the structural features of NRC2, NRC3, and NRC4 (the "Ground Truth") are universally representative of the "canonical" state for the entire ~6,000 sequence NRC family.
2. **AlphaFold 3 Reliability**: It assumes AF3 can accurately model the activated/oligomerized state of diverse NRC sequences, even those lacking high sequence identity to existing PDB templates.
3. **Threshold Accuracy**: It assumes that the specific numerical cutoffs (e.g., <500 Å³ clash, ±10° angle, >15° curvature) are robust enough to separate helpers from sensors/unconventional variants.
4. **Scaffold Validity**: It assumes that "forcing" a protomer into a 9FP6 hexameric symmetry (Metric 3) is a valid bio-physical test for hexameric compatibility.
5. **Pore Stability**: It assumes that the α1-helix is sufficiently well-modeled by AF3 to allow for a precise "Predicted Pore Diameter" calculation (Metric 7).

#### Comparison with Knowledge Base

- **Metric 1 (α4-Helix Curvature):** Abstract [4] and the Knowledge Base confirm the bent α4-helix (specifically at L126) is a differentiator. However, the pdb results log notes that NRC2a's helix was too short to calculate curvature using the script's method, suggesting the "Canonical Baseline" needs sequence-length flexibility.
- **Metric 3 (Steric Clash):** The idea proposes a baseline of **< 500 Å³**. The pdb results log shows that canonical helpers actually produced **5,614 Å³ (NRC2a)** and **25,709 Å³ (SlNRC3)**. The idea's threshold is factually incorrect according to the ground truth analysis.
- **Metric 6 (NB-ARC Angle):** The idea proposes a baseline of **±10°**. The pdb results log shows canonical angles of **137.09°, 45.10°, and 27.16°**. This indicates the idea's proposed threshold is much too narrow and would incorrectly classify canonical proteins as "unconventional."
- **Metric 7 (Pore Diameter):** Abstract [4] confirms the 17–19 Å pore diameter for canonical NRCs, directly supporting this parameter.

#### Reasoning about Correctness

- **Can the assumptions be true?** Mostly yes, but the specific numerical baselines provided in the idea (Metrics 3 and 6) are demonstrably false based on the provided experimental/simulated logs. While the *parameters* (curvature, angles, clashes) are scientifically sound choices, the *values* used to define the "Canonical Baseline" are incorrect.
- **Oligomeric State:** The NRC4 log in the knowledge base mentions an oligomeric count of 15 (dodecamer/pentadecamer states), suggesting that even "canonical" NRCs can deviate from the homohexamer, which the idea does not fully account for.

#### Strength of Evidence

- **Direct Supporting Evidence:** Strong evidence exists for Metric 1 (α4-helix bend), Metric 2 (MADA motif), Metric 4 (exNT existence), and Metric 7 (Pore diameter) from papers [4], [11], and [13].
- **Indirect Supporting Evidence:** Evidence for Metric 5 (Hydrophobic core) is present in [7], but its applicability across the 6,000-sequence family is less certain than the MADA motif.
- **Conflict:** The direct evidence for Metrics 3 and 6 (Clash score and Angle) in the pdb results log contradicts the specific baseline values proposed in the idea.

#### Suggested Improvements

1. **Recalibrate Thresholds:** Adjust the Clash Score baseline to < 30,000 Å³ and the NB-ARC Angle to ±50° (or use a relative Z-score) based on the actual ground truth measurements.
2. **Confidence-Weighted Metrics:** Incorporate AF3 pLDDT and PAE scores into the SNI. If the MADA region has low pLDDT, the SNI should increase because a non-structural N-terminus is a sign of an unconventional/sensor NLR.
3. **Monomeric state fallback:** Since full hexamer modeling is computationally expensive for 6,000 sequences, define how these metrics (like α4 curvature) can be reliably extracted from monomeric AF3 predictions forced into an active conformation.

#### Goal Requirement Assessment

- **5-10 quantitative parameters:** Yes (7).
- **Measurable from PDB/AF3:** Yes.
- **Distinguish unconventional from canonical:** Yes, though the specific numerical values currently fail.
- **Incorporate 9FP6, 9RI9, 9CC8:** Yes.
- **Focus on interface angles/N-terminal variations:** Yes.

The idea respects every goal requirement in structure, but fails on the factual accuracy of its proposed numerical baselines.

#### Reasoning about Correctness and Recommendation

The idea is conceptually excellent and follows the structural biology of the NRC family very closely. However, it is mathematically "incorrect" because the numerical thresholds provided for Metrics 3 and 6 would cause the Ground Truth (canonical) proteins to be rejected as "unconventional." This is a major flaw for a "Structural Novelty Index" baseline. Because the parameters themselves are the correct ones to use, but the values are wrong, the idea is "Questionable/Plausible with Reservations."

I **recommend testing** the idea only after correcting the numerical baselines using the measurements found in the pdb results logs.

Answer: 5

**Novelty:**

$\def\mathcal#1{\mathit{#1}}\def\mathscr#1{\mathit{#1}}$

#### Related Article Abstracts

1. **[19] Structure of the NbNRC2 hexameric resistosome (9FP6):** Provides the foundational structural coordinates for the canonical NbNRC2 hexamer, essential for establishing the baseline for symmetry and protomer orientation.
2. **[10] A disease resistance protein triggers oligomerization of its NLR helper into a hexameric resistosome...:** Detailed protomer-level comparison between the hexamer (9FP6) and the resting-state dimer, highlighting the domain shifts (NB-ARC) required for activation.
3. **[4] A disease resistance protein triggers oligomerization... (Madhuprakash et al., 2024):** Specifically identifies the $\alpha$4-helix bend and the 10° interdomain angle shift in the NB-ARC module as the structural determinants of hexameric vs. pentameric stoichiometry.
4. **[14] 9CC8: Hexameric state of the NRC4 resistosome:** Provides the secondary reference structure for a canonical NRC hexamer (NRC4), confirming the 6-fold symmetry paradigm.
5. **[7] A hydrophobic core in the coiled-coil domain is essential for NRC resistosome function:** Identifies and characterizes the hydrophobic core residues in the CC domain of NRC4, which is a key parameter in the proposed SNI.
6. **[13] An N-terminal motif in NLR immune receptors is functionally conserved...:** Defines the MADA motif and its conservation in NRC-H (helpers) versus its degeneration in NRC-S (sensors).
7. **[11] The activated plant NRC4 immune receptor forms a hexameric resistosome:** Describes the physical dimensions of the NRC4 hexamer, including the ~180 Å diameter and the flexible $\alpha$1 helix.
8. **[18] A helper NLR targets organellar membranes to trigger immunity:** Discusses extended N-termini and different funnel lengths in CC$_R$-NLRs (NRG1/ADR1), providing a comparison for non-canonical N-terminal architectures.
9. **[1] NRC Immune receptor networks show diversified hierarchical genetic architecture...:** Details the expansion of NRC networks and the presence of non-canonical N-terminal extensions (exNT) in certain clades.
10. **[2] AlphaFold 3 predictions for paired NLR proteins:** Demonstrates the existing use of AlphaFold 3 to model pentameric and hexameric NLR structures, validating the feasibility of the proposed screening method.

#### Aspects of the idea already tried

The core structural characteristics used to define the "Canonical Baseline" in the SNI are largely derived from very recent experimental data (2024–2025):

- **$\alpha$4-Helix Curvature:** The specific bend near L126 in NbNRC2 and its role in preventing steric clashes during hexamerization was explicitly described and compared to the straight helix of ZAR1 in **[4]** and **[6]**.
- **MADA Motif Fidelity:** The identification of the MADA motif as a "death switch" and its presence in helpers vs. absence in sensors is the central theme of **[13]** and **[16]**.
- **Hydrophobic Core Packing:** The existence of a critical hydrophobic core in the CC domain of NRC4 and its necessity for function was established in **[7]** and **[9]**.
- **NB-ARC Orientation Angle:** The 10° difference in the interdomain angle of the CC-NB-ARC module between hexameric NRCs and pentameric ZAR1 was measured and reported in **[4]**, **[6]**, and **[10]**.
- **Pore Diameter:** The ~17–19 Å diameter of the NRC2/NRC4/TmSr35 pores compared to the ~12 Å ZAR1 pore was documented in **[4]** and **[6]**.
- **N-terminal Extensions (exNT):** The presence of extended N-terminal domains in certain NRC clades and sensors was noted in **[1]**, **[3]**, and **[18]**.
- **High-Throughput Modeling:** The use of AlphaFold 3 to predict oligomeric states (5-mer vs 6-mer) for large sets of NLRs is already being implemented, as shown in **[2]**, **[5]**, and **[8]**.

#### Novel aspects of the idea

The novelty of the idea lies not in the discovery of these structural features, but in their integration into a unified, **quantitative metric (SNI)** for automated screening:

- **Hexameric Interface Steric Clash Score (Metric 3):** The specific methodology of "forcing" a predicted protomer into a 9FP6/9CC8 C6-symmetry scaffold to calculate atomic overlap volume (Å³) as a proxy for oligomeric compatibility is a novel application of structural modeling in this field.
- **Euler Angle Quantification (Metric 6):** While the angle difference is known, defining it as a quantitative Euler angle deviation metric relative to a fixed CC-domain alignment for a 6,000-sequence dataset is a novel implementation of comparative structural biology.
- **Consolidated Indexing:** Combining these 7 parameters into a single "Structural Novelty Index" allows for the ranking of "unconventionality." Previous studies have focused on characterizing one or two of these features at a time or in small subsets; applying the full set to the entire ~6,000 NRC family is a significant methodological shift.

#### Novelty review

The idea is **moderately novel**.

Strictly speaking, the "baseline" and the "deviation" parameters are almost entirely composed of the findings reported in the provided Ground Truth papers (e.g., Madhuprakash et al., 2024; Liu et al., 2024). A field expert would already know that to find a "novel" NRC, one should look for a missing MADA motif, a straight $\alpha$4-helix, or a different NB-ARC orientation. Therefore, the biological criteria are not novel.

However, the **formalization** of these observations into a specific mathematical index (SNI) intended for high-throughput screening of 6,000 sequences—using specific metrics like "atomic overlap volume" and "void volume/packing density"—is a practical and useful methodological innovation. It transitions from qualitative structural description to a quantitative bioinformatics pipeline.

#### Reasoning about novelty and recommendation

The idea is highly logical and builds directly on the most recent advances in the field. It effectively leverages AlphaFold 3 capabilities to address a massive dataset (~6,000 sequences).

**Should it be tested?** Yes. While the individual parameters are known, the ability to rapidly prioritize the "weirdest" NRCs in a massive dataset using a structural (rather than just sequence-based) score is highly valuable. This could lead to the discovery of tetrameric, octameric, or non-pore-forming NRCs that would otherwise be lost in a broad sequence-similarity search. It is more of an "advanced tool development" idea than a "new biological theory," but it is precisely what the field needs to make use of the ~6,000 newly identified sequences.

**Recommendation:** The idea is solid, though its "novelty" is more about the automation and quantitative synthesis of very recent ground truth rather than a totally new concept.

Answer: 5

**Feasibility:**

$\def\mathcal#1{\mathit{#1}}\def\mathscr#1{\mathit{#1}}$

Based on the provided abstracts and the idea for a Structural Novelty Index (SNI), here is the feasibility judgment.

#### **1. Related Article Abstracts**

1. **[4] Madhuprakash et al. (2024):** Compares NbNRC2 (hexamer) to ZAR1 (pentamer). This is crucial for Parameter 1 ($\alpha$4-helix curvature) and Parameter 6 (NB-ARC orientation), as it provides the structural basis for how these angles dictate stoichiometry.
2. **[7] Wang et al. (2025):** Identifies a specific hydrophobic core in the NRC4 CC-domain. This directly supports the feasibility and biological relevance of Parameter 5 (Hydrophobic Core Packing).
3. **[8] Madhuprakash et al. (2024) [Supp]:** Demonstrates that AlphaFold 3 (AF3) can predict different oligomeric configurations (tetramer to octamer). This validates the feasibility of Parameter 7 (Pore Diameter) and Parameter 3 (Clash Score) using AF3.
4. **[10] Madhuprakash et al. (2024):** Provides a protomer-level comparison between hexamers and dimers, highlighting the "NB-ARC switch." This is essential for calibrating Parameter 6.
5. **[13] Adachi et al. (2019):** Defines the MADA motif and its conservation across ~20% of CC-NLRs. This provides the HMM and sequence baseline needed for Parameter 2.
6. **[11] Liu et al. (2023):** Describes the NRC4 hexamer and a dodecamer state. This highlights the "dense packing" of the LRR, which is relevant for the steric metrics in Parameter 3.
7. **[14] RCSB PDB 9CC8 (2024):** The official entry for the hexameric NRC4 resistosome, serving as one of the primary "Canonical Baselines" for all 7 parameters.
8. **[18] Tarhan et al. (2024):** Discusses extended N-termini in NRG1 helpers compared to canonical ones. This supports the relevance of Parameter 4 (exNT length) in identifying novel helper architectures.
9. **[17] Lin et al. (2025):** Discusses the computational limitations and Python-based analysis scripts (MDAnalysis, GROMACS) used to process AF3 outputs. This relates to the technical feasibility of automating the SNI.
10. **[3] Pai et al. (2025):** Explores NRC expansion in Asterales vs. Solanales. This provides the dataset and phylogenetic context needed to test if the SNI correctly identifies "unconventional" lineages.

#### **2. Steps to Test the Idea**

1. **Metric Automation Scripting:** Develop a Python pipeline (using Biopython and PyMOL/ROSETTA) to automate the calculation of the 7 metrics. Specifically, scripts must calculate helix curvature (Parameter 1), Euler angles between domains (Parameter 6), and atomic overlap volume (Parameter 3).
2. **Ground Truth Calibration:** Run the pipeline on PDB IDs 9FP6, 9RI9, and 9CC8. Establish the "Canonical NRC" mean and standard deviation for each metric.
3. **Outgroup Comparison:** Run the pipeline on known non-hexameric NLRs (e.g., ZAR1 6J5T, Sr35 7XE0). This will verify if the SNI successfully flags known pentamers as "Novel" (High SNI).
4. **Go/No-Go Initial Experiment:** Select 50 representative sequences from the 6,000-sequence dataset (25 known canonical helpers and 25 suspected unconventional sensors/helpers). Model these 50 as monomers using AF3. Apply Parameters 1, 2, 4, 5, and 6.
   - **Go:** If the SNI scores for the 25 sensors/unconventional NLRs are statistically higher than the 25 canonical helpers.
   - **No-Go:** If the index cannot distinguish between a canonical helper (NRC2) and a known divergent sensor (e.g., Rx).
5. **High-Throughput Screening:** Run monomeric AF3 on the ~6,000 sequences. Calculate the SNI. For the top 5% (highest SNI), perform multimer AF3 runs to validate Parameter 7 (Pore Diameter) and confirm novel oligomeric states.

#### **3. Feasibility Reasoning**

The testing of this idea is **moderately feasible to highly feasible** depending on the computational budget.

- **Measurability:** Most parameters (1, 2, 4, 5, 6) rely on monomeric models. Monomer AF3 runs for 6,000 sequences are computationally accessible for a standard lab using local clusters or cloud credits ($1,000–$2,000). The metrics themselves (curvature, angles, packing) are standard structural bioinformatics tasks.
- **Novelty of Parameter 3:** The "Hexameric Interface Steric Clash Score" is a clever, high-feasibility proxy. Instead of modeling 6,000 hexamers (which is extremely expensive and memory-intensive), one can simply align a monomer to a hexamer template and calculate overlaps. This drastically reduces the resource demand.
- **Data Availability:** The Ground Truth Dataset (9FP6, 9CC8, 9RI9) is recent and high-resolution, providing excellent baseline values.
- **Bottlenecks:** Parameter 7 (Pore Diameter) is the most difficult to test at scale because it requires "modeled oligomers." Modeling 6,000 hexamers in AF3 would take months and significant GPU resources. However, if Parameter 7 is reserved only for a "refined" subset identified by the other 6 parameters, the overall project is very streamlined.

Answer: 7

**Impact potential:**

$\def\mathcal#1{\mathit{#1}}\def\mathscr#1{\mathit{#1}}$

#### Related Article Abstracts

1. **[4] A disease resistance protein triggers oligomerization of its NLR helper into a hexameric resistosome to mediate innate immunity (Madhuprakash et al., 2024):** This is the primary source for the structural basis of hexamer formation (e.g., the $\alpha$4-helix bend at L126 and the 10° NB-ARC angle shift), which directly informs SNI Parameters 1, 6, and 7.
2. **[7] A hydrophobic core in the coiled-coil domain is essential for NRC resistosome function (Wang et al., 2025):** Identifies a specific CC-domain hydrophobic core in NRC4, providing the rationale and baseline for SNI Parameter 5.
3. **[13] An N-terminal motif in NLR immune receptors is functionally conserved across distantly related plant species (Adachi et al., 2019):** Defines the MADA motif as the "death switch" for NRCs, essential for Parameter 2's fidelity check.
4. **[11] The activated plant NRC4 immune receptor forms a hexameric resistosome (Liu et al., 2023):** Provides the structural basis for the ~180 Å diameter and specific LRR domain angles, supporting Parameter 7 and potential LRR metrics.
5. **[1] NRC Immune receptor networks show diversified hierarchical genetic architecture across plant lineages (Goh et al., 2024):** Confirms the existence of unconventional NRCs like NRCx (MADA-less) and the scale of the NRC family (thousands of sequences).
6. **[19] Structure of the NbNRC2 hexameric resistosome (PDB 9FP6):** This is the core "Ground Truth" reference for NbNRC2, essential for benchmarking all canonical metrics.
7. **[14] Hexameric state of the NRC4 resistosome (PDB 9CC8):** Provides the structural ground truth specifically for the NRC4-clade parameters.
8. **[18] A helper NLR targets organellar membranes to trigger immunity (Tarhan et al., 2024):** Discusses extended N-terminal funnels in non-canonical NLRs (NRG1), supporting Parameter 4 and 7.
9. **[3] A hierarchical immune receptor network in lettuce reveals contrasting patterns of evolution (Pai et al., 2025):** Documents the loss of specific N-terminal domains in Asterales NRC-sensors, validating the need for an index to find structural "novelty."
10. **[8] AlphaFold 3 can predict different oligomeric configurations of NbNRC2 (Madhuprakash et al., 2024 - Supplement):** Validates the feasibility of using AF3 to model the hexameric states of NRC sequences for screening.

#### Detailed Assumptions

1. **Modeling Accuracy:** It is assumed that AlphaFold 3 (AF3) can accurately predict the active, oligomeric resistosome state for thousands of sequences without experimental activation (e.g., using "activation mimic" mutations or specific seeds).
2. **Structural Conservation of the "Canonical" space:** It is assumed that the three provided PDB structures (9FP6, 9RI9, 9CC8) represent the full structural range of "canonical" hexameric helpers across all Solanaceae.
3. **Correlation of Structure and Function:** It is assumed that structural deviation (high SNI) from the hexameric model directly correlates with unconventional biological functions (e.g., non-pore signaling, sensor activity, or negative regulation).
4. **High-Throughput Extraction:** It is assumed that the 5–10 quantitative metrics (like Euler angles or void volumes) can be extracted programmatically from AF3 PDB outputs at scale (6,000+ sequences).

#### Feasibility and Reasoning

- **Feasibility of Parameter Extraction:** High. Parameters 1, 4, 6, and 7 are geometric and can be automated using standard Python libraries (MDAnalysis/Biopython). Parameter 3 (Steric Clash) is a standard protein-docking metric.
- **Feasibility of AF3 Modeling:** Moderate. While [8] shows AF3 handles NRC2 hexamers well, running 6,000 sequences as hexamers is computationally expensive. However, as [5] and [17] suggest, AF3 multimer is becoming a standard screening tool.
- **Realistic Impact of Parameters:**
  - **$\alpha$4 Curvature (Parameter 1):** Highly realistic. [4] explicitly states the "straight $\alpha$4 helix of AtZAR1 would sterically overlap... in a hexameric configuration." This is a definitive structural determinant for stoichiometry.
  - **NB-ARC Orientation (Parameter 6):** Realistic. [6] and [10] highlight a 10° widening in NRC2 vs. ZAR1 that allows for the extra protomer.
  - **Hydrophobic Core (Parameter 5):** Realistic and novel. [7] shows this core is missing in many sensors, making it a powerful discriminator.
- **Effect on Impact:** If these assumptions hold, the SNI provides a significant leap over sequence identity. Sequence identity often fails to predict the functional "switch" (e.g., NRCx looks like an NRC but lacks the MADA motif). Structural metrics capture the physical requirements of the "death switch" [16].

#### Suggested Improvements

1. **Include LRR Tangent Angle:** [11] notes that the LRR domain in NRC4 is arranged at a smaller angle relative to the tangent of the wheel compared to pentameric NLRs. This would be a more robust parameter than just NB-ARC orientation.
2. **EDVID Motif Check:** [7] and [9] emphasize the EDVID motif's role in intramolecular regulation. Adding a metric for the structural distance/interaction between the EDVID region and the LRR would enhance the "Novelty" assessment for sensors vs. helpers.
3. **pLDDT-Weighted RMSD:** Since the N-terminal $\alpha$1 is often flexible [11], the MADA fidelity metric (Parameter 2) should be weighted by the local confidence score (pLDDT) to avoid penalizing correctly predicted but flexible regions.
4. **Iterative Benchmarking:** Use the known "unconventional" helper NRCx [1] as a control to ensure the SNI correctly identifies it as a high-novelty candidate.

#### Overall Impact Potential

The idea has **high impact potential**. By moving beyond simple sequence phylogeny to a multidimensional structural index, this approach allows researchers to classify the "dark matter" of the NRC family—the 6,000+ sequences that may contain novel sensors, negative regulators, or even NLRs with different stoichiometry (tetramers or octamers).

The proposal is grounded in the most recent structural breakthroughs [4, 7, 14] and addresses a specific bottleneck in plant immunity: the functional characterization of expanded NLR networks. The use of parameters like $\alpha$4-helix curvature and NB-ARC angles is a sophisticated application of structural biology to evolutionary genomics. If implemented, this could lead to the discovery of entirely new modes of NLR signaling beyond the canonical pore-forming resistosome [18].

**Conclusion:** This idea is a well-designed, technically feasible, and biologically grounded framework that leverages state-of-the-art tools (AF3) and ground-truth data to solve a major problem in the field.

Answer: 8

References:

[1] [NRC Immune receptor networks show diversified hierarchical genetic architecture across plant lineages - PMC](https://pmc.ncbi.nlm.nih.gov/articles/PMC11371147/)

[2] [AlphaFold 3 predictions for paired NLR proteins - Obsidian Vault - Obsidian v1.8.7](https://zenodo.org/records/15552925/files/AlphaFold%203%20predictions%20for%20paired%20NLR%20proteins.pdf?download=1)

[3] [A hierarchical immune receptor network in lettuce reveals contrasting patterns of evolution in sensor and helper NLRs | bioRxiv](https://www.biorxiv.org/content/10.1101/2025.02.25.639832v1.full-text)

[4] [A disease resistance protein triggers oligomerization of its NLR helper into a hexameric resistosome to mediate innate immunity](https://www.ncbi.nlm.nih.gov/pmc/articles/PMC11540030/)

[5] [A disease resistance protein triggers oligomerization of its NLR helper into a hexameric resistosome to mediate innate immunity](https://www.ncbi.nlm.nih.gov/pmc/articles/PMC11540030/)

[6] [A disease resistance protein triggers oligomerization of its NLR helper into a hexameric resistosome to mediate innate immunity](https://www.ncbi.nlm.nih.gov/pmc/articles/PMC11540030/)

[7] [A hydrophobic core in the coiled-coil domain is essential for NRC resistosome function](https://www.biorxiv.org/content/10.1101/2025.01.21.634219v2)

[8] [A disease resistance protein triggers oligomerization of its NLR helper into a hexameric resistosome to mediate innate immunity](https://www.biorxiv.org/content/10.1101/2024.06.18.599586v1)

[9] [A hydrophobic core in the coiled-coil domain essential for NRC resistosome function](https://www.biorxiv.org/content/10.1101/2025.01.21.634219v1)

[10] [A disease resistance protein triggers oligomerization of its NLR helper into a hexameric resistosome to mediate innate immunity](https://www.biorxiv.org/content/10.1101/2024.06.18.599586v1)

[11] [The activated plant NRC4 immune receptor forms a hexameric resistosome](https://www.biorxiv.org/content/10.1101/2023.12.18.571367v1)

[12] [Jurassic NLR: Conserved and dynamic evolutionary features of the atypically ancient immune receptor ZAR1 - PMC](https://pmc.ncbi.nlm.nih.gov/articles/PMC10533333/)

[13] [An N-terminal motif in NLR immune receptors is functionally conserved across distantly related plant species | eLife](https://elifesciences.org/articles/49956)

[14] [RCSB PDB - 9CC8: Hexameric state of the NRC4 resistosome](https://www.rcsb.org/structure/9cc8)

[15] [Structures of plant resistosome reveal how NLR immune receptors are activated - PMC](https://pmc.ncbi.nlm.nih.gov/articles/PMC9590527/)

[16] [An N-terminal motif in NLR immune receptors is functionally conserved across distantly related plant species](https://www.biorxiv.org/content/10.1101/693291v1)

[17] [AlphaFold can be used to predict the oligomeric states of proteins](https://www.biorxiv.org/content/10.1101/2025.03.10.642518v1)

[18] [A helper NLR targets organellar membranes to trigger immunity](https://www.biorxiv.org/content/10.1101/2024.09.19.613839v1)

[19] [RCSB PDB - 9FP6: Structure of the NbNRC2 hexameric resistosome](https://www.rcsb.org/structure/9fp6)

[20] [A helper NLR targets organellar membranes to trigger immunity](https://www.biorxiv.org/content/10.1101/2024.09.19.613839v1)

[21] [Can AI modelling of protein structures distinguish between sensor and helper NLR immune receptors?](https://www.biorxiv.org/content/10.1101/2024.11.24.625045v1)

[22] [Activation of plant immunity through conversion of a helper NLR homodimer into a resistosome](https://www.ncbi.nlm.nih.gov/pmc/articles/PMC11524475/)

[23] [Dynamic accumulation of a helper NLR at the plant-pathogen interface underpins pathogen recognition](https://www.biorxiv.org/content/10.1101/2021.03.15.435521v1)

[24] [(PDF) Structural Transition of NRC4 to Hexameric Resistosome Activates Plant Immunity](https://www.researchgate.net/publication/386157649_Structural_Transition_of_NRC4_to_Hexameric_Resistosome_Activates_Plant_Immunity)

[25] [NRC immune receptor networks show diversified hierarchical genetic architecture across plant lineages | bioRxiv](https://www.biorxiv.org/content/10.1101/2023.10.25.563953v1.full-text)

[26] [A disease resistance protein triggers oligomerization of its NLR helper into a hexameric resistosome to mediate innate immunity](https://www.biorxiv.org/content/10.1101/2024.06.18.599586v1)

[27] [Boosting AlphaFold Protein Tertiary Structure Prediction through MSA Engineering and Extensive Model Sampling and Ranking in CASP16 - PMC](https://pmc.ncbi.nlm.nih.gov/articles/PMC12258999/)

[28] [Predicting the structure of large protein complexes using AlphaFold and Monte Carlo tree search | bioRxiv](https://www.biorxiv.org/content/10.1101/2022.03.12.484089.full)

[29] [Phylogenomics of Plant NLR Immune Receptors to Identify Functionally Conserved Sequence Motifs - PMC](https://pmc.ncbi.nlm.nih.gov/articles/PMC11237980/)

[30] [The Plant “Resistosome”: Structural Insights into Immune Signaling | Request PDF](https://www.researchgate.net/publication/335189084_The_Plant_Resistosome_Structural_Insights_into_Immune_Signaling)

[31] [Structure–function analyses of coiled-coil immune receptors define a hydrophobic module for improving plant virus resistance - PMC](https://pmc.ncbi.nlm.nih.gov/articles/PMC10010612/)

**Motivation:**

$\def\mathcal#1{\mathit{#1}}\def\mathscr#1{\mathit{#1}}$

**Coherence:**

$\def\mathcal#1{\mathit{#1}}\def\mathscr#1{\mathit{#1}}$

**Deep verification:**

$\def\mathcal#1{\mathit{#1}}\def\mathscr#1{\mathit{#1}}$
