## Supplementary Data S6 for "AI-guided discovery of atypical protein assemblies": [EXTERNAL] ├░Y╠ê┬o┬1⁄4 Co-scientist - Hypothesis 119627191265642 - A Hierarchical 8-Parameter Structural Novelty Index (SNI) for Distinguishing Unconventional NRC-NLRs via AlphaFold 3 - For research purposes only - Google confidential.docx

### 🧬 Co-scientist - A Hierarchical 8-Parameter Structural Novelty Index (SNI) for Distinguishing Unconventional NRC-NLRs via AlphaFold 3

**About**: *This is an experimental system for generating novel and testable hypotheses. The hypotheses are generated by a model and may be wrong. For research purposes only.*

### Proposal

$\def\mathcal#1{\mathit{#1}}\def\mathscr#1{\mathit{#1}}$

The Structural Novelty Index (SNI) utilizes domain-centroid geometries and interface-burial metrics, incorporating structural insights from the inactive dimer structure (9CC8) and the active hexamers (9FP6, 9RI9).

#### 1. The SNI Framework

The SNI is a hierarchical screening tool that processes sequences through three structural filters: Monomeric Autoinhibition, Dimeric Resting State, and Multimeric Resistosome Architecture.

##### 1.1. Phase 1: The Monomeric "Readiness" Index (mSNI)

*Focus: Detecting deviations in the internal "fold" of the NB-ARC and CC domains.*

1. **Parameter 1: N-terminal "MADA" Vector Tilt ($\theta_{MADA}$)**
   - **Definition:** The angle between the principal axis of the CC $\alpha1$ helix (residues 1–15) and the NBD-HD1-WHD plane.
   - **Canonical (9FP6):** $\sim 75^\circ$ (Tucked/Autoinhibited).
   - **Novelty:** Deviations $>20^\circ$ suggest a pre-exposed or non-functional executioner domain.
   - **Confidence Weight:** Multiply by the mean pLDDT of the CC $\alpha1$ helix.
2. **Parameter 2: NB-ARC Centroid Kink ($\alpha_{NB-ARC}$)**
   - **Definition:** The angle formed by the centroids of the NBD, HD1, and WHD domains.
   - **Rationale:** This captures the "bend" of the NB-ARC module regardless of loop lengths or indels.
   - **Canonical Baseline:** Mean angle derived from AF3-modeled NRC2, 3, and 4.
   - **Novelty:** Shift $>15^\circ$ indicates a fundamentally different nucleotide-binding pocket architecture.
3. **Parameter 3: LRR-WHD Surface Proximity ($D_{latch-surf}$)**
   - **Definition:** The minimum distance between the centroid of the WHD (ARC2) and the plane defined by the C$\alpha$ atoms of the central LRR repeats (LRR 4–8).
   - **Rationale:** More robust than residue-to-residue "latch" metrics.
   - **Canonical (9RI9/9CC8):** Tight proximity ($\sim 12\text{\AA}$ centroid-to-plane).
   - **Novelty:** Distance $>18\text{\AA}$ suggests a "loose" auto-regulatory state or an unconventional trigger mechanism.

##### 1.2. Phase 2: The Dimer-Interface Index (dSNI)

*Focus: Assessing the ability to form the canonical resting state (Reference: 9CC8).*

4. **Parameter 4: Inactive Dimer Burial Index ($BI_{dimer}$)**
   - **Definition:** The Ratio of the Interface Surface Area (iSASA) to the total monomer surface area when modeled as an anti-parallel homodimer (the NbNRC4 resting state).
   - **Canonical (9CC8):** High burial index ($\sim 0.15$).
   - **Novelty:** $BI < 0.05$ suggests the NLR bypasses the dimer state or uses a novel autoinhibitory interface, common in "atypical" helpers.
5. **Parameter 5: NB-NBD Interface Torsion ($\tau_{dimer}$)**
   - **Definition:** The dihedral angle between the NBD centroids of two protomers in the predicted dimer.
   - **Canonical:** Defined by the 9CC8 symmetry.
   - **Novelty:** Significant rotation suggests a different resting-state geometry, potentially affecting the speed of activation.

##### 1.3. Phase 3: The Multimeric Resistosome Index (cSNI)

*Focus: Distinguishing stoichiometry and pore architecture.*

6. **Parameter 6: Pore Radial Symmetry ($R_{pore}$)**
   - **Definition:** The distance from the centroid of the CC $\alpha1$ tips (residues 1–5) to the central symmetry axis of the multimer.
   - **Canonical (9FP6):** $\sim 15\text{\AA}$ (Standard hexameric pore).
   - **Novelty:** $>20\text{\AA}$ (Heptamer/Octamer) or $<10\text{\AA}$ (Non-pore former).
7. **Parameter 7: Protomer Interface Coherence ($PAE_{inter}$)**
   - **Definition:** The average Predicted Aligned Error (PAE) between adjacent protomers in an AF3 multimer model.
   - **Rationale:** Uses AF3's internal confidence to distinguish stable complexes from "forced" models.
   - **Canonical:** Low PAE ($<10\text{\AA}$) at the NBD-NBD interface.
   - **Novelty:** High PAE despite high monomer pLDDT suggests a non-canonical or transient stoichiometry.
8. **Parameter 8: MHD-P-loop Spatial Distance ($d_{MHD-P}$)**
   - **Definition:** Euclidean distance between the centroid of the MHD motif (activation switch) and the P-loop (nucleotide-binding site).
   - **Canonical:** Established distance in the active hexamer state.
   - **Novelty:** Deviations indicate an alternative "trigger" geometry for ATP hydrolysis or exchange.

#### 2. Implementation Strategy

##### 2.1. Dynamic Confidence-Weighted SNI ($SNI_{cw}$)

To address potential modeling artifacts, the final score for each parameter is weighted by the AF3 confidence of the involved domains: $$Value_{weighted} = Value_{observed} \times \left( \frac{\text{Mean pLDDT}_{domain1} + \text{Mean pLDDT}_{domain2}}{200} \right)$$ This ensures that "novelty" in low-confidence/disordered regions is automatically penalized.

##### 2.2. Z-Score Calibration

Before screening the 6,000 sequences, the SNI is calibrated using:

- **Positive Controls:** ZAR1 (pentamer), NRCX (non-MADA), WAI3 (octamer).
- **Negative Controls:** NbNRC2, SlNRC3, NbNRC4 (canonical). The weights ($w_i$) are adjusted via a linear classifier to maximize the distance between these two control groups.

##### 2.3. Handling Indels

By using centroids of conserved domain folds (e.g., the core 5-helix bundle of the NBD) rather than specific residue numbers, the index remains stable even if the LRR has variable repeat numbers or if the NB-ARC has elongated loops.

#### 3. Structural and Functional Rationale

- **Inclusion of 9CC8:** The index includes the dimer interface, which accounts for the resting state of canonical NRCs.
- **Robustness:** Centroid-based metrics are designed to be stable against indels compared to residue-to-residue distances.
- **Noise Reduction:** The PAE-based interface check ($PAE_{inter}$) prevents the identification of novelty in unstable multimeric models.
- **Functional Focus:** The MADA Vector Tilt and Pore Radial Symmetry relate to the protein's ability to form a functional pore, providing metrics beyond sequence similarity.

### Reviews summary

$\def\mathcal#1{\mathit{#1}}\def\mathscr#1{\mathit{#1}}$

#### Research contacts

$\def\mathcal#1{\mathit{#1}}\def\mathscr#1{\mathit{#1}}$

Based on the provided articles, the following researchers are experts in the structural biology and evolutionary networks of NRC-NLRs. They are suggested to review the hypothesis regarding the **Structural Novelty Index (SNI)**, as their work directly addresses the structural divergence, hexameric resistosome formation, and AlphaFold-based modeling of these receptors.

#### 1. Sophien Kamoun & Michael W. Webster (The Sainsbury Laboratory)

- **Justification:** These researchers are central to the structural characterization of the **NbNRC2** hexameric resistosome. Their work on **PDB ID 9FP6** provides the exact structural baseline mentioned in the goal for canonical hexameric resistosomes. They have extensively published on the transition from the dimeric resting state to the activated hexameric state, which is foundational to the "Phase 2: Dimeric Resting State" and "Phase 3: Multimeric Resistosome Architecture" components of the SNI.
- **Relevant Excerpt:** "We determined the structure of sensor-activated NbNRC2, which forms a hexameric inflammasome-like resistosome... Comparative structural analyses between the resting state homodimer and sensor-activated homohexamer revealed substantial rearrangements" (Abstract 1).

#### 2. Furong Liu & Brian J. Staskawicz (UC Berkeley)

- **Justification:** These researchers authored the primary research on the **NRC4 hexameric resistosome (PDB ID 9CC8)**. Their work is critical for reviewing the index's ability to distinguish NRC4-specific structural features. They provide deep insights into the auto-active mutations and the dodecameric state (inactive double-layer) that serves as a vital comparison for identifying "novel" or "unconventional" NRC behaviors in high-throughput screenings.
- **Relevant Excerpt:** "Our research reveals that the hNLR, known as NLR required for cell death 4 (NRC4), assembles into a hexameric resistosome... expanding our understanding of the regulation of plant immune responses" (Abstract 6).

#### 3. Chih-Hang Wu (Academia Sinica)

- **Justification:** Dr. Wu’s research focuses on the **functional diversity and conservation of the CC domain** within the NRC family. His work on identifying the "hydrophobic core" (L34, A72, I114, V118) and the subfunctionalization of NRC3 (Abstract 12, 13) provides the necessary biological context for the "N-terminal CC-domain variations" parameter in the proposed SNI. He is uniquely qualified to assess whether the SNI parameters correctly capture the evolutionary divergence of NRCs.
- **Relevant Excerpt:** "We identified a novel hydrophobic feature within the CC domain that contributes to NRC-mediated immunity... This hydrophobic core is important for NRC4 subcellular localization, oligomerization, and phospholipid association" (Abstract 7).

#### 4. AmirAli Toghani (The Sainsbury Laboratory)

- **Justification:** Toghani’s work specifically utilizes **AlphaFold 3** to distinguish between helper (NRC-H) and sensor (NRC-S) NLRs based on their capacity to form resistosome-like oligomers. Since the SNI relies heavily on AlphaFold 3 metrics (pLDDT, ipTM, PAE) for high-throughput screening, his expertise in computational structural prediction makes him an essential reviewer for the index’s feasibility and calibration.
- **Relevant Excerpt:** "Recently, we demonstrated that AlphaFold 3... can differentiate between helper and sensor NLRs within pairs by predicting resistosome-like structures for helpers but not for sensors" (Abstract 5).

#### Summary Table of Expertise

| Researcher | Key Focus Area | Relevance to SNI Goal |
| --- | --- | --- |
| **S. Kamoun / M. Webster** | NbNRC2 Cryo-EM (9FP6) | Defines the canonical hexameric resistosome baseline. |
| **F. Liu / B. Staskawicz** | NRC4 Structure (9CC8) | Provides reference for NRC4-specific hexameric packing. |
| **C. H. Wu** | CC-Domain Hydrophobic Cores | Validates CC-domain parameters and evolutionary novelty. |
| **A. Toghani** | AlphaFold 3 NLR Modeling | Reviews the high-throughput AF3 screening methodology. |

### Appendix:

**All reviews:**

**Correctness:**

$\def\mathcal#1{\mathit{#1}}\def\mathscr#1{\mathit{#1}}$

This review evaluates the proposed **Structural Novelty Index (SNI)** for identifying unconventional NRC-NLRs.

#### 1. Related Article Abstracts

1. **[1] RCSB PDB - 9FP6: Structure of the NbNRC2 hexameric resistosome**: Confirms 9FP6 is a hexameric *active* state, providing the baseline for Parameters 2, 6, and 8.
2. **[15] RCSB PDB - 9RI9: Cryo-EM structure of the tomato NRC3 hexameric resistosome**: Provides the second canonical hexameric baseline for the NRC family.
3. **[6] RCSB PDB - 9CC8: Hexameric state of the NRC4 resistosome**: Confirms 9CC8 is an *active* hexamer, directly contradicting the idea's use of it as an "inactive dimer."
4. **[3] A disease resistance protein triggers... (Abstract 3)**: Compares NbNRC2 (9FP6) and AtZAR1 (6J5T), providing specific quantitative angles ($85^\circ$ vs $75^\circ$) and pore radii ($14\text{\AA}$ vs $13\text{\AA}$).
5. **[10] A disease resistance protein triggers... (Abstract 10)**: Describes the structural transition ($180^\circ$ rotation) between the resting dimer and active hexamer.
6. **[7] A hydrophobic core in the coiled-coil domain is essential...**: Identifies key CC-domain residues (L34, A72, I114, V118) that form a hydrophobic core, relevant for Parameter 1 and 3.
7. **[5] A hierarchical immune receptor network in lettuce...**: Demonstrates that AF3 can distinguish helpers (hexameric models) from sensors (failed models), supporting Parameter 7.
8. **[4] Sensor NLR immune proteins activate oligomerization...**: Defines the MADA motif (residues 1–21), essential for the vector definitions in Parameter 1 and 6.
9. **[9] The activated plant NRC4 immune receptor forms a hexameric resistosome**: Details the unique NBD-HD1 inter-protomer interface in NRC4, relevant for Parameters 4, 5, and 7.
10. **[8] A helper NLR targets organellar membranes to trigger immunity**: Compares funnel lengths and CC-domain variations, supporting the logic for Parameter 1 (MADA tilt).

#### 2. Detailed Assumptions

1. **Activation State Accuracy:** The idea assumes PDB 9FP6 and 9CC8 represent "Tucked/Autoinhibited" and "Inactive Dimer" states, respectively.
2. **AlphaFold 3 (AF3) Modeling Reliability:** AF3 can accurately predict the stoichiometry and N-terminal $\alpha1$ helix orientations of ~6,000 diverse NRC sequences.
3. **Metric Stability:** Centroid-based geometric metrics are robust enough to handle the high sequence diversity (indels) of the ~350 Solanaceae species dataset.
4. **Dimer Universality:** All canonical NRCs (NRC2, 3, 4) utilize a conserved anti-parallel dimer interface as their primary resting state.
5. **Direct Correlation:** Deviations in static structural metrics (like MADA tilt) directly correlate with "unconventional" functional mechanisms.

#### 3. Comparison with Knowledge Base & Abstracts

- **9FP6 State:** The idea classifies 9FP6 as "Canonical (9FP6): $\sim 75^\circ$ (Tucked/Autoinhibited)." **Knowledge Base/Abstract [1] disagrees:** 9FP6 is the *activated hexameric resistosome*. In this state, the MADA motif is typically extended to form a pore, not "tucked."
- **9CC8 State:** The idea uses 9CC8 for the "Inactive Dimer Burial Index." **Knowledge Base/Abstract [6, 9, 17] disagrees:** 9CC8 is explicitly the *hexameric state* of the NRC4 resistosome. The resting dimer is a different structural configuration (e.g., PDB 8RFH in Abstract [7]).
- **Quantitative Angles:** The idea proposes a $\sim 75^\circ$ MADA tilt for NRC2. **Abstract [3] disagrees:** It explicitly labels the inter-domain angle for NRC2 as $85^\circ$, while $75^\circ$ is the value for the *AtZAR1* pentamer.
- **Pore Radius:** The idea proposes $\sim 15\text{\AA}$ for the pore. **Abstract [3] provides data:** NRC2 has a $14\text{\AA}$ pore and ZAR1 has a $13\text{\AA}$ pore.
- **Dimer Interface:** The idea cites 9CC8 for dimer torsion. Since 9CC8 is a hexamer, it cannot provide the baseline for dimer dihedral symmetry.

#### 4. Reasoning about Correctness of Assumptions

1. **Activation State Accuracy (False):** This is a major structural error. Using active resistosome coordinates to define "autoinhibited" baselines will result in the SNI failing to detect activation-related shifts.
2. **AF3 Reliability (Moderately Plausible):** Abstract [5] and [14] show AF3 is effective at modeling NRC hexamers, but Abstract [2] notes that the $\alpha1$ helix is often disordered in cryo-EM, though AF3 models it with high confidence.
3. **Metric Stability (Plausible):** Centroid-based metrics are standard in structural biology for comparing divergent proteins with indels.
4. **Dimer Universality (Plausible):** Abstract [2, 10, 16] support the dimer-to-hexamer transition for NRC2, and Abstract [7] suggests similar autoinhibitory mechanisms in NRC4.

#### 5. Strength of Evidence

- **Direct Evidence:** Abstract [3] and [10] provide direct supporting evidence for using inter-domain angles and pore dimensions to distinguish NLR architectures (hexamer vs. pentamer). Abstract [5] provides direct evidence that AF3 metrics (pTM, ipTM) can distinguish helpers.
- **Indirect Evidence:** Abstract [7] and [13] provide indirect support for "Monomeric Readiness" by identifying a hydrophobic core that regulates CC-domain transitions.

#### 6. Suggested Improvements

- **Correct PDB Baselines:** Re-assign 9FP6, 9RI9, and 9CC8 as the **Active Hexamer** baseline. Use PDB 8RFH (NRC2 dimer) for the **Inactive Dimer** baseline.
- **Adjust Angles:** Update the canonical MADA tilt baseline for NRC2 from $75^\circ$ to $85^\circ$ based on Abstract [3].
- **Refine Parameter 1:** Instead of just "Tilt," measure the **Angular Shift ($\Delta\theta$)** between the predicted monomer and the hexamer baseline to capture the "readiness" to oligomerize.
- **Clarify Parameter 4:** Since 9CC8 is a hexamer, the Burial Index should be calculated using a separate dimer model (e.g., 8RFH) or the dodecameric interface mentioned in Abstract [9].

#### 7. Goal Requirement Assessment

- **5-10 Parameters:** **Yes** (8 proposed).
- **Measurable from PDB/AF3:** **Yes**, all rely on coordinates or AF3 confidence (PAE).
- **Distinguish unconventional from canonical:** **Yes**, the framework (mSNI, dSNI, cSNI) is designed for this.
- **Leverage Ground Truth (9FP6, 9RI9, 9CC8):** **Partial failure.** It includes the IDs but misidentifies their biological states and values.
- **High-throughput/Solanaceae dataset:** **Yes**, the use of centroids and Z-scores is appropriate for 6,000 sequences.

#### 8. Reasoning about Correctness & Recommendation

The SNI framework is conceptually excellent. Dividing the index into three phases (Monomer, Dimer, Multimer) is a sophisticated approach to structural biology screening. However, the idea contains **significant factual errors** regarding the provided Ground Truth Dataset. It misidentifies the activation state of all three primary PDB references (9FP6 and 9CC8 are active hexamers, but the idea treats them as inactive/dimeric). Furthermore, the specific value for the "canonical" angle ($75^\circ$) belongs to the *AtZAR1* pentamer, not the NRC hexamer ($85^\circ$).

While the *math* and *logic* of the parameters (centroids, burial indices, PAE) are correct and valuable, the *baselines* are fundamentally wrong. A screening tool calibrated with these errors would fail to distinguish novelty from standard activation.

**Recommendation:** Do not publish in its current form. It requires a correction of the PDB reference states and baseline values.

Answer: 4

**Novelty:**

$\def\mathcal#1{\mathit{#1}}\def\mathscr#1{\mathit{#1}}$

#### 1. Related Article Abstract Titles

1. **[1] RCSB PDB - 9FP6: Structure of the NbNRC2 hexameric resistosome**: Essential for establishing the canonical hexameric baseline and protomer interface metrics.
2. **[2] A disease resistance protein triggers oligomerization of its NLR helper into a hexameric resistosome to mediate innate immunity**: Provides the structural basis for the transition from dimer to hexamer and details the NB-ARC rearrangements.
3. **[3] A disease resistance protein triggers oligomerization... (Image Description)**: Specifically mentions quantitative metrics like inter-domain angles (85° vs 75°) and pore diameters, directly relating to the proposed SNI parameters.
4. **[5] A hierarchical immune receptor network in lettuce reveals contrasting patterns of evolution...**: Demonstrates the use of AlphaFold 3 (AF3) metrics (pLDDT, pTM, ipTM) to distinguish between helper and sensor NLRs.
5. **[6] RCSB PDB - 9CC8: Hexameric state of the NRC4 resistosome**: Provides ground truth for another canonical NRC helper (NRC4).
6. **[7] A hydrophobic core in the coiled-coil domain is essential for NRC resistosome function**: Identifies specific conserved residues and internal CC-domain geometry relevant to "Phase 1" of the proposed SNI.
7. **[9] The activated plant NRC4 immune receptor forms a hexameric resistosome**: Describes the dodecameric state and specific interface interactions (NBD-HD1) that the SNI must distinguish.
8. **[10] Comparison between resting state homodimer and activated hexamer of NbNRC2...**: Details the 180° rotation of the NB-ARC modules, which is critical for the "Dimer-Interface Index" (Phase 2).
9. **[15] RCSB PDB - 9RI9: Cryo-EM structure of the tomato NRC3 hexameric resistosome**: Provides the final canonical NRC ground truth (NRC3) for the reference set.
10. **[19] An atypical NLR protein modulates the NRC immune receptor network...**: Describes NRCx, a non-MADA "atypical" NRC that serves as a perfect test case for the SNI's ability to detect unconventionality.

#### 2. Aspects of the Idea Already Explored

Several aspects of the SNI are already well-documented or utilized in recent NLR structural biology:

- **Quantitative Inter-domain Angles:** Abstract [3] and [10] explicitly measure and compare angles between domains (e.g., 10° difference in NB-ARC angles between NRC2 and ZAR1). This directly corresponds to **Parameter 2** (NB-ARC Centroid Kink).
- **Pore Radial Metrics:** Abstract [3] and [9] compare the diameter and radial distances of NRC hexamers (e.g., 180 Å diameter) versus pentamers (e.g., ZAR1 87 Å), which is the core of **Parameter 6** (Pore Radial Symmetry).
- **Use of AF3 Confidence Metrics:** Abstract [5] and [14] already use pLDDT, pTM, and ipTM to screen sequences and differentiate their ability to form resistosomes. The idea's **Parameter 7** (Protomer Interface Coherence) and the use of pLDDT for weighting are standard practices in recent NLR bioinformatics [5].
- **Transition from Dimer to Hexamer:** The transition and conformational changes between the resting dimer (9CC8) and the active hexamer are described in detail in [2] and [10], providing the biological basis for **Phase 2** (Dimer-Interface Index).
- **MADA Motif and Alpha-1 Helix Importance:** The role of the N-terminal α1 helix in forming the pore is a central theme in many abstracts [4, 8, 18], making the focus on this region (as in **Parameter 1**) a common approach.

#### 3. Novel Aspects of the Idea

The novelty of the idea lies in the formalization and technical implementation of these metrics for high-throughput screening:

- **Centroid-Centroid Geometry for Robustness:** Most existing studies [3, 10] use residue-to-residue or domain-to-domain angles. The proposal to use **centroids of conserved folds** (e.g., the core 5-helix bundle of the NBD) is a novel computational strategy to handle the high indel rate and variable loop lengths in a 6,000-sequence dataset.
- **The MADA Vector Tilt ($\theta_{MADA}$):** While the MADA motif's function is known, defining its orientation relative to the NBD-HD1-WHD plane as a specific quantitative "readiness" metric is a novel way to categorize autoinhibition state in uncharacterized sequences.
- **Hierarchical Structural Filtering:** The "Phase 1-3" approach (Monomer -> Dimer -> Multimer) provides a systematic framework for detecting novelty at every stage of the NLR lifecycle, whereas most current screenings (like [5]) focus primarily on the final multimeric state.
- **Integration of Inactive Dimer Burial Index ($BI_{dimer}$):** Utilizing the burial ratio of the antiparallel dimer (from 9CC8) to detect "atypical" helpers that might bypass this state or use different autoinhibitory interfaces is a specific and potentially powerful novel application of the ground truth data.

#### 4. Novelty Review

The idea is **moderately novel**. It successfully transitions from the qualitative "comparative structural analysis" found in recent high-impact papers [2, 9, 10] to a formalized, automated mathematical index. However, the core parameters—domain angles, pore size, burial metrics, and AF3 confidence scores—are already the standard tools of the trade for the experts currently defining this field. Abstract [3] and [5] show that researchers are already looking at precisely these quantitative differences to distinguish NRCs from ZAR1 or sensors from helpers.

The most "novel" technical contribution is the robust implementation (centroid-based vs. residue-based) and the specific hierarchical combination into an "SNI score." While useful, this is more of an engineering/methodological improvement than a conceptual breakthrough.

#### 5. Reasoning and Recommendation

The idea should be tested because it provides a necessary formalization for handling large-scale datasets (~6,000 sequences). A simple RMSD (Root-Mean-Square Deviation) check is often insufficient for NLRs because of their high sequence plasticity and variable loop regions; a centroid-based geometric index is much more likely to yield biologically meaningful "structural outliers."

However, because many of the parameters (pore size, domain angles) are already being analyzed by the authors of the ground truth papers, the "novelty" is limited to the specific formulation and scale of the index. It is a highly practical and well-designed tool for discovery, but it builds directly upon very recent (2024-2025) published methodologies.

Answer: 5

**Feasibility:**

$\def\mathcal#1{\mathit{#1}}\def\mathscr#1{\mathit{#1}}$

#### Related Article Abstracts

1. **[1] Structure of the NbNRC2 hexameric resistosome (9FP6)**: Provides the primary ground truth for a canonical activated hexamer, essential for defining the baseline values of Parameter 1 (MADA tilt) and Parameter 6 (pore symmetry).
2. **[6] Hexameric state of the NRC4 resistosome (9CC8)**: Offers the structural basis for Parameter 4 (Burial Index) and Parameter 5 (Interface Torsion) by providing the inactive dimer reference.
3. **[10] Comparison between resting state homodimer and activated hexamer of NbNRC2**: Describes the $180^\circ$ rotation and rearrangements between states, justifying the use of centroid-based angles (Parameter 2) to capture large-scale conformational changes.
4. **[5] A hierarchical immune receptor network in lettuce...**: Demonstrates that AlphaFold 3 (AF3) can distinguish between helpers and sensors using metrics like PAE and pTM, directly supporting the feasibility of Parameter 7.
5. **[7] A hydrophobic core in the coiled-coil domain is essential for NRC resistosome function**: Identifies internal CC domain folding requirements, which informs the structural validity of the Phase 1 monomeric index (mSNI).
6. **[11] Deep learning facilitates precise identification of disease-resistance genes in plants**: Discusses the practical limitations of high-throughput AF3 screening (e.g., web server limits), which is a key factor in judging the time/cost feasibility.
7. **[13] A hydrophobic core in the coiled-coil domain... (AF3 modeling)**: Shows that AF3 can successfully model NRC dimers and hexamers, validating the primary method for extracting the SNI parameters.
8. **[14] NRC Immune receptor networks show diversified hierarchical genetic architecture...**: Confirms the use of AF3 to model hexamers with oleic acid, providing a protocol for the multimeric resistosome index (cSNI).
9. **[15] Cryo-EM structure of the tomato NRC3 hexameric resistosome (9RI9)**: Provides a secondary canonical hexamer reference to ensure the SNI is not overfitted to a single species (N. benthamiana).
10. **[19] An atypical NLR protein modulates the NRC immune receptor network (NRCX)**: Highlights NRCX as a vital positive control for "novelty" since it lacks a functional MADA motif and acts as a negative regulator.

#### Steps to Test the Idea

1. **Script Development**: Write a Python tool (using Biopython and NumPy) to calculate the 8 defined geometric parameters (centroids, planes, angles, and distances) from PDB/CIF files.
2. **Initial Go/No-Go Experiment**: Apply the script to the Ground Truth structures (9FP6, 9RI9, 9CC8) and known non-canonical outliers (AtZAR1 pentamer, NRCX).
   - *Success Criterion*: The SNI must clearly separate the NRC2/3/4 group from ZAR1 and NRCX with a Z-score > 2.0. If parameters cannot distinguish these, the index must be refined.
3. **Gold Standard Subset Modeling**: Run AF3 (Multimer) on a curated subset of 100 sequences (50 canonical helpers, 50 sensors/atypical NLRs).
4. **Parameter Extraction & Weighting**: Extract the 8 parameters from the AF3 outputs and use a linear classifier (or PCA) to determine the optimal weights ($w_i$) for the calibrated SNI.
5. **Full-Scale Production**: Deploy the validated SNI on the full ~6,000 sequence dataset. This requires local installation of AF3 on a GPU cluster to bypass web server limits.
6. **Biological Validation**: Correlate high SNI scores with existing functional data (e.g., cell death assays or known "helper" vs "sensor" status) in the Solanaceae dataset.

#### Reasoning about Feasibility

The testing feasibility is high in terms of **Metric Measurability**. All 8 proposed parameters rely on standard geometric calculations (centroids, angles, SASA) or native AF3 outputs (PAE), making them highly automatable. The use of centroids rather than specific residue numbers effectively addresses the challenge of sequence indels.

However, the **Computational Scalability** presents a significant hurdle. Modeling 6,000 sequences as both dimers and hexamers (Phase 2 and 3) is a massive undertaking. A hexameric NRC model (~850 residues x 6 = ~5,100 residues) is at the upper limit of memory for standard 80GB A100 GPUs and typically requires 1–2 hours per run. Processing 6,000 sequences (12,000 total multimer runs) would require roughly 12,000–24,000 GPU hours. This falls squarely into the "Resource Intensive" category (Score 5) of the rubric, requiring 2–3 months of dedicated cluster time and significant computational budget. The initial validation (Go/No-Go and subset testing) is highly feasible within 2 weeks, but the "Goal" specifies identifying unconventional NLRs across the *entire* 6,000-sequence family.

Answer: 5

**Impact potential:**

$\def\mathcal#1{\mathit{#1}}\def\mathscr#1{\mathit{#1}}$

#### 1. Related Article Abstracts

1. **[1] RCSB PDB - 9FP6: Structure of the NbNRC2 hexameric resistosome**: Provides the primary canonical baseline for the sensor-activated hexameric state of NRC2, essential for defining "normal" parameters.
2. **[10] Comparison between resting state homodimer and activated hexamer of NbNRC2**: Details the structural modules (NB-HD1 and WHD-LRR) and the $180^\circ$ rotation required for activation, which justifies Phase 2 (dSNI) and Phase 3 (cSNI).
3. **[6] RCSB PDB - 9CC8: Hexameric state of the NRC4 resistosome**: Establishes the secondary canonical reference for NRC4, including the dodecameric state which provides a baseline for multimeric deviation.
4. **[5] A hierarchical immune receptor network in lettuce reveals contrasting patterns of evolution**: Demonstrates that AlphaFold 3 (AF3) can distinguish between helpers and sensors based on their ability to form hexameric resistosomes, supporting the feasibility of high-throughput structural screening.
5. **[7] A hydrophobic core in the coiled-coil domain is essential for NRC resistosome function**: Identifies a specific hydrophobic core (L34, A72, I114, V118) in the CC domain, providing a concrete metric for Parameter 1 and internal CC structural integrity.
6. **[19] An atypical NLR protein modulates the NRC immune receptor network**: Describes NRCX, which lacks a MADA motif and acts as a negative regulator, serving as the perfect "positive control" for an unconventional NRC to test the SNI.
7. **[8] A helper NLR targets organellar membranes to trigger immunity**: Shows that AF3 can predict diverse N-terminal funnel lengths (e.g., NRG1 vs. NRC2), supporting the validity of Parameter 6 (Pore Radial Symmetry).
8. **[15] RCSB PDB - 9RI9: Cryo-EM structure of the tomato NRC3 hexameric resistosome**: Provides the third canonical reference structure (NRC3), completing the "canonical trio" (NRC2, 3, 4) required for Z-score calibration.
9. **[13] A hydrophobic core in the coiled-coil domain is essential for NRC resistosome function**: Explains how AF3 models can reveal internal CC residues (e.g., A72) that interact with the LRR, supporting the use of centroid-based proximity metrics.
10. **[4] Sensor NLR immune proteins activate oligomerization of their NRC helper**: Highlights MADA motif conservation and its role in pore formation, justifying the use of the MADA vector tilt ($\theta_{MADA}$) as a novelty metric.

#### 2. Detailed Assumptions

1. **AF3 Structural Reliability:** The index assumes AlphaFold 3 can accurately model the multimeric states of uncharacterized NRCs with enough precision to distinguish $10^\circ–20^\circ$ shifts in domain orientations.
2. **Canonical Consistency:** It assumes that NRC2, 3, and 4 (the references) form a tight structural cluster, such that a Z-score based on their variance is a meaningful filter for the other 6,000 sequences.
3. **Centroid-Function Correlation:** It assumes that global domain-centroid geometries (angles/distances) are better predictors of "unconventional" function than specific side-chain interactions or primary sequence motifs.
4. **Resting State Universality:** The dSNI phase assumes that the anti-parallel homodimer observed in NbNRC2 [10] is the universal resting state for all canonical NRCs.
5. **Modeling Throughput:** It assumes the computational resources are available to generate high-quality AF3 multimer models (hexamers/dimers) for a significant portion of the 6,000-sequence dataset.

#### 3. Feasibility and Effect on Impact

- **AF3 Precision (High Feasibility):** Recent studies [5, 14] confirm that AF3 effectively differentiates between helper and sensor NLRs and accurately predicts the N-terminal $\alpha1$ helices that were historically difficult to resolve. This high feasibility ensures that the metrics derived (like $\theta_{MADA}$) will be based on reliable data.
- **Structural Clustering (Moderate Feasibility):** While NRC2, 3, and 4 are similar, Abstract [10] notes that NbNRC2 has a $10^\circ$ wider NB-ARC angle than AtZAR1 to accommodate a hexamer. This suggests that the "canonical" baseline itself has inherent variance. If the baseline is too broad, the SNI might fail to flag subtle but functionally important novelties.
- **Centroid Robustness (High Feasibility):** Using centroids rather than specific residues is a highly realistic way to handle the high sequence diversity and indels common in the LRR and NB-ARC domains [7, 10]. This increases the impact potential by making the tool applicable to the wide diversity of Solanaceae NRCs.
- **Resting State (Moderate Feasibility):** While 9CC8 and 9RFH show dimers, some NRCs like SlNRC2 form filaments [7]. If an unconventional NRC uses a filament or monomer as a resting state, the dSNI might return a high novelty score, which is exactly the intended impact of the tool.
- **Computational Bottleneck (Low Feasibility):** Modeling 6,000 sequences as hexamers in AF3 is a massive task. However, the proposal addresses this by positioning the SNI as a "hierarchical screening tool," implying Phase 1 (monomers) acts as a filter before moving to dimers/hexamers.

#### 4. Suggested Improvements

1. **Stoichiometry Testing:** Expand Parameter 6 ($R_{pore}$) to specifically test for pentameric (ZAR1-like) and octameric (WAI3-like) symmetries. If an AF3 run is set to $N=6$ but the protein "prefers" $N=5$, the $PAE_{inter}$ (Parameter 7) will be high, which should be explicitly coded as a novelty indicator.
2. **Membrane Interaction Metric:** Incorporate a "Hydrophobic Core exposure" metric. Since the CC-domain hydrophobic core is essential for function [7, 13], a parameter measuring the change in solvent-accessible surface area (SASA) of these core residues (L34, A72, I114, V118) between dimer and hexamer would be highly diagnostic.
3. **Lipid-Docking simulation:** Reference [5] mentions modeling with oleic acid. Adding a parameter that measures the distance or binding energy of the CC-funnel to a simulated membrane layer would enhance the "functional" aspect of the SNI.

#### 5. Overall Impact Potential

The Structural Novelty Index (SNI) is a high-impact concept. It moves the field of NLR biology beyond sequence-based phylogenetics into "structural phylogenomics."

- **Feasibility:** High. The use of PDB IDs 9FP6, 9RI9, and 9CC8 provides a ground-truth baseline that is structurally sound. The reliance on AF3 is supported by very recent literature [5, 14].
- **Scope:** Broad. It covers the entire Solanaceae NRC network, which is the most complex immune network known in plants.
- **Long-term implications:** By identifying unconventional NRCs (like NRCX [19]), this tool could lead to the discovery of entirely new modes of immune regulation, such as "decoy" helpers or "bridge" proteins.
- **Practicality:** The hierarchical approach (Monomer -> Dimer -> Hexamer) makes it computationally realistic.

This idea is likely to have a **high impact** because it provides a standardized, mathematical framework to identify the "weird" proteins in a massive dataset, which are often the keys to understanding evolutionary innovations.

Answer: 8

References:

[1] [RCSB PDB - 9FP6: Structure of the NbNRC2 hexameric resistosome](https://www.rcsb.org/structure/9fp6)

[2] [A disease resistance protein triggers oligomerization of its NLR helper into a hexameric resistosome to mediate innate immunity - PMC](https://pmc.ncbi.nlm.nih.gov/articles/PMC11540030/)

[3] [A disease resistance protein triggers oligomerization of its NLR helper into a hexameric resistosome to mediate innate immunity](https://www.ncbi.nlm.nih.gov/pmc/articles/PMC11540030/)

[4] [Sensor NLR immune proteins activate oligomerization of their NRC helper](https://www.biorxiv.org/content/10.1101/2022.04.25.489342v1)

[5] [A hierarchical immune receptor network in lettuce reveals contrasting patterns of evolution in sensor and helper NLRs](https://www.biorxiv.org/content/10.1101/2025.02.25.639832v1)

[6] [RCSB PDB - 9CC8: Hexameric state of the NRC4 resistosome](https://www.rcsb.org/structure/9cc8)

[7] [A hydrophobic core in the coiled-coil domain is essential for NRC resistosome function | bioRxiv](https://www.biorxiv.org/content/10.1101/2025.01.21.634219v3.full-text)

[8] [A helper NLR targets organellar membranes to trigger immunity](https://www.biorxiv.org/content/10.1101/2024.09.19.613839v1)

[9] [The activated plant NRC4 immune receptor forms a hexameric resistosome](https://www.biorxiv.org/content/10.1101/2023.12.18.571367.full.pdf)

[10] [A disease resistance protein triggers oligomerization of its NLR helper into a hexameric resistosome to mediate innate immunity](https://www.biorxiv.org/content/10.1101/2024.06.18.599586v1)

[11] [Deep learning facilitates precise identification of disease-resistance genes in plants](https://www.biorxiv.org/content/10.1101/2024.09.26.615248v1)

[12] [Subfunctionalization of NRC3 altered the genetic structure of the Nicotiana NRC network | PLOS Genetics](https://journals.plos.org/plosgenetics/article?id=10.1371/journal.pgen.1011402)

[13] [A hydrophobic core in the coiled-coil domain is essential for NRC resistosome function](https://www.biorxiv.org/content/10.1101/2025.01.21.634219v2)

[14] [A helper NLR targets organellar membranes to trigger immunity](https://www.biorxiv.org/content/10.1101/2024.09.19.613839v1)

[15] [RCSB PDB - 9RI9: Cryo-EM structure of the tomato NRC3 hexameric resistosome](https://www.rcsb.org/structure/9ri9)

[16] [An effector from the potato late blight pathogen bridges ENTH-domain protein TOL9a to an activated helper NLR to suppress immuni](https://www.biorxiv.org/content/biorxiv/early/2025/07/08/2025.07.06.663370.full.pdf)

[17] [(PDF) Structural Transition of NRC4 to Hexameric Resistosome Activates Plant Immunity](https://www.researchgate.net/publication/386157649_Structural_Transition_of_NRC4_to_Hexameric_Resistosome_Activates_Plant_Immunity)

[18] [NRC Immune receptor networks show diversified hierarchical genetic architecture across plant lineages - PMC](https://pmc.ncbi.nlm.nih.gov/articles/PMC11371147/)

[19] [An atypical NLR protein modulates the NRC immune receptor network in Nicotiana benthamiana - PMC](https://pmc.ncbi.nlm.nih.gov/articles/PMC9851556/)

[20] [An atypical NLR protein modulates the NRC immune receptor network in Nicotiana benthamiana | PLOS Genetics](https://journals.plos.org/plosgenetics/article?id=10.1371/journal.pgen.1010500)

[21] [An N-terminal motif in NLR immune receptors is functionally conserved across distantly related plant species](https://www.biorxiv.org/content/10.1101/693291v1)

[22] [A plant pathogen effector blocks stepwise assembly of a helper NLR resistosome](https://www.biorxiv.org/content/10.1101/2025.07.14.664264v1.full.pdf)

[23] [Helper NLR immune protein NRC3 evolved to evade inhibition by a cyst nematode virulence effector | PLOS Genetics](https://journals.plos.org/plosgenetics/article?id=10.1371/journal.pgen.1011653)

[24] [Diversification of the “EDVID” packing motif underpins structural and functional variation in plant NLR coiled-coil domains](https://www.biorxiv.org/content/10.1101/2025.06.01.657260v1)

[25] [A hydrophobic core in the coiled-coil domain is essential for NRC resistosome function](https://www.biorxiv.org/content/10.1101/2025.01.21.634219v2)

[26] [The nucleotide binding domain of NRC-dependent disease resistance proteins is sufficient to activate downstream helper NLR oligomerization and immune signaling | bioRxiv](https://www.biorxiv.org/content/10.1101/2023.11.30.569466v1.full-text)

[27] [Accurate structure prediction of biomolecular interactions with AlphaFold 3 - PMC](https://pmc.ncbi.nlm.nih.gov/articles/PMC11168924/)

**Motivation:**

$\def\mathcal#1{\mathit{#1}}\def\mathscr#1{\mathit{#1}}$

**Coherence:**

$\def\mathcal#1{\mathit{#1}}\def\mathscr#1{\mathit{#1}}$

**Deep verification:**

$\def\mathcal#1{\mathit{#1}}\def\mathscr#1{\mathit{#1}}$
