## Supplementary Data S6 for "AI-guided discovery of atypical protein assemblies": [EXTERNAL] ├░Y╠ê┬o┬1⁄4 Co-scientist - Hypothesis 180708842503749 - Developing an 8-Parameter Dual-Phase SNI for Unconventional NRC-NLR Identification via AlphaFold 3 - For research purposes only - Google confidential.docx

### 🧬 Co-scientist - Developing an 8-Parameter Dual-Phase SNI for Unconventional NRC-NLR Identification via AlphaFold 3

**About**: *This is an experimental system for generating novel and testable hypotheses. The hypotheses are generated by a model and may be wrong. For research purposes only.*

### Proposal

$\def\mathcal#1{\mathit{#1}}\def\mathscr#1{\mathit{#1}}$

##### 1. Phase 1: Monomeric SNI (mSNI) - Primary Filter

*Calculated using AlphaFold 3 (AF3) Monomer predictions or the AlphaFold Database.*

- **Parameter 1: N-terminal Helix Propensity ($\alpha_{MADA}$)**
  - *Definition:* Average pLDDT and DSSP-derived helical fraction of residues 1–30.
  - *Canonical:* High confidence ($>85$ pLDDT) and continuous $\alpha$-helix.
  - *Novelty:* pLDDT $<70$ or transition to coil indicates a non-MADA/unconventional execution^7^.
- **Parameter 2: The ARC2-LRR "Latch" Proximity ($D_{latch}$)**
  - *Definition:* Minimum Euclidean distance between residues within the WHD (ARC2) domain (residues 392–497)^9^ that interface with the LRR concave surface and the LRR inner-concave surface.
  - *Canonical:* $\sim 3.8\text{\AA}$ (Active auto-inhibition).
  - *Novelty:* Deviations $>3\text{\AA}$ from the NbNRC2 baseline^28^ suggest a "triggered-by-default" or novel auto-regulatory state.
- **Parameter 3: NB-ARC Swivel Angle ($\phi_{swivel}$)**
  - *Definition:* Vector angle between the NBD (P-loop centroid) and HD1/ARC2 centroid^16^ within the monomer.
  - *Canonical:* Established from AF3-modeled NbNRC2/SlNRC3.
  - *Novelty:* A twist outside a $\pm 10^\circ$ window indicates an alternative nucleotide-binding architecture^5^.
- **Parameter 4: LRR Expansion Ratio ($\chi_{LRR}$)**
  - *Definition:* The ratio of the distance between the C$\alpha$ of the first and last conserved Leucine residues to the total LRR sequence length.
  - *Canonical:* Ratio derived from PDB 9FP6.
  - *Novelty:* Represents LRR "straightening" or "hyper-curving" without complex circle-fitting math.

##### 2. Phase 2: Complex SNI (cSNI) - Targeted Validation

*Calculated from AF3 Multimer (Hexamer) outputs^25^ for the top ~500 candidates.*

- **Parameter 5: Protomer Divergence Angle ($\Delta\theta_{int}$)**
  - *Definition:* The dihedral angle between centroids of adjacent protomers.
  - *Canonical:* $60^\circ$ (Hexamer)^12^.
  - *Novelty:* $72^\circ \pm 2^\circ$ (Pentameric ZAR1-like)^1^ or $<52^\circ$ (Heptamer/Octamer).
- **Parameter 6: Pore Funnel Hydrophobicity Gradient (FHG)**
  - *Definition:* The difference in Kyte-Doolittle hydropathy between the CC $\alpha1$ tip (Residues 1–10) and the $\alpha1$ base (Residues 15–25).
  - *Canonical:* Highly hydrophobic tip (MADA signature)^21^.
  - *Novelty:* An inverted or neutral gradient suggests organelle-specific targeting or non-pore-forming functions^11^.
- **Parameter 7: Interface Burial Efficiency ($\eta_{inter}$)**
  - *Definition:* $iSASA / \text{Total Protomer Area}$.
  - *Canonical:* Baseline derived from 9RI9 (NRC3) and 9CC8 (NRC4).
  - *Novelty:* Low $\eta$ suggests transient complexes^13^; high $\eta$ indicates stable or constitutive resistosomes^23^.
- **Parameter 8: Stoichiometric Preference Score ($S_{pref}$)**
  - *Definition:* Comparison of AF3 ipTM scores when a sequence is forced into a pentamer vs. a hexamer model^14^.
  - *Novelty:* Positive values for (Pentamer - Hexamer) indicate a departure from the NRC paradigm.

##### 3. Implementation & Standardization Guide

1. **Standardized Baselines:** All "Canonical Means" are derived by running NRC2, NRC3, and NRC4 through AF3 Multimer. This ensures that the SNI measures biological deviation from ground truth rather than software artifacts (AF3 vs. PDB).
2. **Weighted Z-Score Formula:** $$SNI = \sum_{i=1}^n w_i \cdot \frac{|Value_{observed} - Mean_{AF3\_Ref}|}{SD_{AF3\_Ref}}$$
   - *High weights ($w=3$):* $\Delta\theta_{int}$, $\alpha_{MADA}$, $D_{latch}$.
   - *Medium weights ($w=1.5$):* FHG, $\phi_{swivel}$.
3. **The Screening Funnel:**
   - **Step 1:** Calculate **mSNI** for all 6,000 monomers^27^ (utilizing existing AFDB structures where possible).
   - **Step 2:** Select sequences with mSNI $> 10$ (Outliers) for AF3 Multimer modeling.
   - **Step 3:** Calculate **cSNI** to confirm structural novelty.

##### 4. Success Criteria

This index identifies unconventional NLRs by their monomeric orientation first, reducing the GPU hours required, and uses AF3-on-AF3 comparisons to eliminate experimental data gaps (missing N-termini).

[7]  [A disease resistance protein triggers oligomerization of its NLR helper into a hexameric resistosome to mediate innate immunity - PubMed Central.](https://vertexaisearch.cloud.google.com/grounding-api-redirect/AUZIYQEikF5Rauh3jkb3MvgUExSGfqpE0SIUEjuQeJsPf4FmyJGrnDJSNneeYIkVXgTUmk1fbGyY626exhJV7C-SKztJv-tKtS6FHqibCEtJwH9u7lOaMGXXcYxxBcHJ_gB9r1kES9EwKdNVyrIqIY-a)

[8]  [A helper NLR targets organellar membranes to trigger immunity - bioRxiv.](https://vertexaisearch.cloud.google.com/grounding-api-redirect/AUZIYQGWBKcTU77hw5RlcQmoYIwD8_oogA9nHOxSz3BN0c34Dn96xQo_cSHFHVXnsMfWq5tgL2LY9VqvaI2npittTEkmCOKn9u1hGhuRX1kEHOGYNc84D8oteJFBnDqNAzDFoufxlSv09sXVflCQR0iksba7CKVl_tGuHC9oFpGXbB302A==)

[9] Jogi, AmirAli, P., Andres, Jake, Jiorgos, et al.  [A disease resistance protein triggers oligomerization of its NLR helper into a hexameric resistosome to mediate innate immunity.](https://www.ncbi.nlm.nih.gov/pmc/articles/PMC11540030/) Published 2024. [https://www.ncbi.nlm.nih.gov/pmc/articles/PMC11540030/.](https://www.ncbi.nlm.nih.gov/pmc/articles/PMC11540030/)

[10] Tarhan, Him, Hung-Yu, J., AmirAli, Jiorgos, et al.  [A helper NLR targets organellar membranes to trigger immunity.](https://www.biorxiv.org/content/10.1101/2024.09.19.613839v1) Published 2024. [https://www.biorxiv.org/content/10.1101/2024.09.19.613839v1.](https://www.biorxiv.org/content/10.1101/2024.09.19.613839v1)

[11] Tarhan, Him, Hung-Yu, J., AmirAli, Jiorgos, et al.  [A helper NLR targets organellar membranes to trigger immunity.](https://www.biorxiv.org/content/10.1101/2024.09.19.613839v1) Published 2024. [https://www.biorxiv.org/content/10.1101/2024.09.19.613839v1.](https://www.biorxiv.org/content/10.1101/2024.09.19.613839v1)

[12]  [The activated plant NRC4 immune receptor forms a hexameric resistosome - bioRxiv.](https://vertexaisearch.cloud.google.com/grounding-api-redirect/AUZIYQGsxQacHBlrSBjqUAZvOlz9ZFNOr0UFKqlrPzezxheJWFRl3vpUlzWS7DQ3WAscWgv5WAx0ksXp29akbw7pJ-kBKIfHl6H-wpfnl1g_bwURX4G2PFql-r7cWUuuXc23p8Xc9qwnxSDrg6pIVMx93JanSH8arej3x8LMrsKO1BBf5g==)

[13]  [A disease resistance protein triggers oligomerization of its NLR helper into a hexameric resistosome to mediate innate immunity - PubMed Central.](https://vertexaisearch.cloud.google.com/grounding-api-redirect/AUZIYQHBzVGYccSYoj13EBeGlc6qeZLc-zDdUaxesL_VYFBTQVYoxcLmyMisHqMo4t1YiVoZBnbXx_sSkt-Gxcpi3e7B_dUHeZWnPZGLtaRKppkUOFsOdx1_9UcY9YIfy-IxEfTmdHkSfl7-TpKUT4Y=)

[14] Jogi, AmirAli, P., Andres, Jake, Jiorgos, et al.  [A disease resistance protein triggers oligomerization of its NLR helper into a hexameric resistosome to mediate innate immunity.](https://www.biorxiv.org/content/10.1101/2024.06.18.599586v1) Published 2024. [https://www.biorxiv.org/content/10.1101/2024.06.18.599586v1.](https://www.biorxiv.org/content/10.1101/2024.06.18.599586v1)

[15] Kazuyoshi, Takatsugu, Junichi, Kentaro.  [Protein-segment universe exhibiting transitions at intermediate segment length in conformational subspaces.](https://www.ncbi.nlm.nih.gov/pmc/articles/PMC2529298/) Published 2008. [https://www.ncbi.nlm.nih.gov/pmc/articles/PMC2529298/.](https://www.ncbi.nlm.nih.gov/pmc/articles/PMC2529298/)

[16] Jogi, AmirAli, P., Andres, Jake, Jiorgos, et al.  [A disease resistance protein triggers oligomerization of its NLR helper into a hexameric resistosome to mediate innate immunity.](https://www.ncbi.nlm.nih.gov/pmc/articles/PMC11540030/) Published 2024. [https://www.ncbi.nlm.nih.gov/pmc/articles/PMC11540030/.](https://www.ncbi.nlm.nih.gov/pmc/articles/PMC11540030/)

[17]  [Activation of the helper NRC4 immune receptor forms a hexameric resistosome | Request PDF - ResearchGate.](https://vertexaisearch.cloud.google.com/grounding-api-redirect/AUZIYQEfRGdNwpfpJ5lWjfCxCj8lnjhQNnVl43Ta3z1D1YfAC6qzVlCCt_lHtCQBqSpQZzHzvlKt11MYgUtLI5gIU9F6gH2ddepASPX8SkAdfVCulUot4aAO55bdP_QxiGUWFNVn-TdeXtlAivxDkbONekFRxRdswRUjGFtuUhDPWnfEUL3kfWXSEnaQUmBp1v4_a1EpSoi40sUF_f7SUvxDF0cXiRvJgdyWNGXSw1Cq6rNROLwAUpRcV-7JFg==)

[18] Neelanjana, K., K., N..  [Crystal Structure of a Monomeric Thiolase-Like Protein Type 1 (TLP1) from Mycobacterium smegmatis.](https://www.ncbi.nlm.nih.gov/pmc/articles/PMC3406046/) Published 2012. [https://www.ncbi.nlm.nih.gov/pmc/articles/PMC3406046/.](https://www.ncbi.nlm.nih.gov/pmc/articles/PMC3406046/)

[19] P., Hsuan, Yasin, Cian, Lok, Vergara, et al.  [Sensor NLR immune proteins activate oligomerization of their NRC helper.](https://www.biorxiv.org/content/10.1101/2022.04.25.489342v1) Published 2022. [https://www.biorxiv.org/content/10.1101/2022.04.25.489342v1.](https://www.biorxiv.org/content/10.1101/2022.04.25.489342v1)

[20]  [A disease resistance protein triggers oligomerization of its NLR helper into a hexameric resistosome to mediate innate immunity - PubMed Central.](https://vertexaisearch.cloud.google.com/grounding-api-redirect/AUZIYQGx_MKMGTvYlZ5SPMmb9CQ1HAvt0Vw6kFdw07EWF_x7T1YdxMX4wBfi6AGN7CvRXyvWLKG2fL9cAxnEpIC1jTPPMjiJBuq3hNPhBZKMwjrXDj4GB4LjJ7eCDvfTKTWobimnHT9G7udj-BGhMRem)

[21] Alexander, Jiorgos.  [NLR immune receptors: structure and function in plant disease resistance.](https://www.ncbi.nlm.nih.gov/pmc/articles/PMC10586772/) Published 2023. [https://www.ncbi.nlm.nih.gov/pmc/articles/PMC10586772/.](https://www.ncbi.nlm.nih.gov/pmc/articles/PMC10586772/)

[22] Jogi, AmirAli, P., Andres, Jake, Jiorgos, et al.  [A disease resistance protein triggers oligomerization of its NLR helper into a hexameric resistosome to mediate innate immunity.](https://www.biorxiv.org/content/10.1101/2024.06.18.599586v1) Published 2024. [https://www.biorxiv.org/content/10.1101/2024.06.18.599586v1.](https://www.biorxiv.org/content/10.1101/2024.06.18.599586v1)

[23] Jogi, AmirAli, P., Andres, Jake, Jiorgos, et al.  [A disease resistance protein triggers oligomerization of its NLR helper into a hexameric resistosome to mediate innate immunity.](https://www.biorxiv.org/content/10.1101/2024.06.18.599586v1) Published 2024. [https://www.biorxiv.org/content/10.1101/2024.06.18.599586v1.](https://www.biorxiv.org/content/10.1101/2024.06.18.599586v1)

[24]  [AlphaFold two years on: Validation and impact - PNAS.](https://vertexaisearch.cloud.google.com/grounding-api-redirect/AUZIYQEQuNCYnRJ2W9PngUWIUn1O2wW_jtNC9F9TEeelxv13DSzC-6zSrWCAaGp9PH7Qg9lFNHFIElcMFHQUzXBLarZ_AJW6vObCqN9OJNIZi-UfZHZ6iC5IPxM7V-9vuf6uR4OQn7O_SBG53Z-Iww==)

[25] Jogi, AmirAli, P., Andres, Jake, Jiorgos, et al.  [A disease resistance protein triggers oligomerization of its NLR helper into a hexameric resistosome to mediate innate immunity.](https://www.ncbi.nlm.nih.gov/pmc/articles/PMC11540030/) Published 2024. [https://www.ncbi.nlm.nih.gov/pmc/articles/PMC11540030/.](https://www.ncbi.nlm.nih.gov/pmc/articles/PMC11540030/)

[26]  [A disease resistance protein triggers oligomerization of its NLR helper into a hexameric resistosome to mediate innate immunity | Request PDF - ResearchGate.](https://vertexaisearch.cloud.google.com/grounding-api-redirect/AUZIYQHPD26vBlC_GcsADVyUuAPaiWq4pyyeU-eUMtK_50Xc8B0s_nwSrjA9VPl_o0t2thx8p9r6U-BTDQasolnoA7wy4KQveiglBFtrBsZwfdoZ54_-Q9g-KFJncCHhx1lZ_MaPMz3Hlv9fFQzj1jg900q8v7UncwltfQETWhlv4JOsWIFMISvsGiVeGFs_718rs4Mhwdayx4R4NXJAEDndeRGx0wIO1JflTNO4Ys_zmuvzbioFMYcTSBwNaS4hsFXB1FD0It5zI7oonpmBBm5wF7jO9_oh6u5u-TAQ6xhq7txgAFRAXqFOgI1OoAWCrH3u)

[27] Muniyandi, AmirAli, Hsuan, Yu, Jiorgos, Him, et al.  [Activation of plant immunity through conversion of a helper NLR homodimer into a resistosome.](https://www.biorxiv.org/content/10.1101/2023.12.17.572070v1) Published 2023. [https://www.biorxiv.org/content/10.1101/2023.12.17.572070v1.](https://www.biorxiv.org/content/10.1101/2023.12.17.572070v1)

[28]  [pdb_00009fp6 - wwPDB.](https://vertexaisearch.cloud.google.com/grounding-api-redirect/AUZIYQFbjIhxseSbW-EAQgrSYUTg2__C_HnVKlz4lUFNnnGu6HdJDNkngxPivx6S_p-aNMKQShgshXd9a01WmI3kdQsYxg3Qa2XFlEPhJcohV7UfZq_WbpzXtgxHqlS8-fxVDQSxl8af)

[29]  [9RI9: Cryo-EM structure of the tomato NRC3 hexameric resistosome - RCSB PDB.](https://vertexaisearch.cloud.google.com/grounding-api-redirect/AUZIYQGNDpHiKRLR0t_c3FcNDndpeQ7aOoyCbDKa7XWM0GTQ0NT8GoNSYnQ2_hLLp5ETf0V5qrTdJSg_OE0g-dWE4_YfbMoPiTtgju41tZfjkTTsdE-lSM9D0C_VuzNxC1zd)

[30]  [9FP6: Structure of the NbNRC2 hexameric resistosome - RCSB PDB.](https://vertexaisearch.cloud.google.com/grounding-api-redirect/AUZIYQEbo05tPSVFqY5U6dy_FimcuxcxtpvAK1OuwDEvji3Wak3aZ3y7FdO04oOeHgQynNFpr2GZq5cFc9ohNURhxw81G30nBOQuFUlbaZiCGU8v_8Dry5DbxLTjvCwq40E=)

[31]  [9ri9 - Cryo-EM structure of the tomato NRC3 hexameric resistosome - Summary - Protein Data Bank Japan.](https://vertexaisearch.cloud.google.com/grounding-api-redirect/AUZIYQFYkPuP2s2wEzj2gh7CIXIdQwYyNgjySy7LBkd3x7eAEd93jvA1bDcAj8gUsjcCuNo5D8SVjMhG-oHCr8sQGYxQ6csm6eCjEaVQXZEyi-waQq1cbXNJBddWLd8_GyM=)

[32]  [9cc8 - Hexameric state of the NRC4 resistosome - Summary - Protein Data Bank Japan.](https://vertexaisearch.cloud.google.com/grounding-api-redirect/AUZIYQEjxNikQYA9o4mghtducDOcSn23ZxSZZK4Z_1P9HFCHuou6cl2yhZXs4hzyZ45gJQOFF3zXzQsYUbJjK_tX4C3aMo_jYqj-8bgLoyesIQXAzsRIOUwt_OtJdgm3fxI=)

### Reviews summary

$\def\mathcal#1{\mathit{#1}}\def\mathscr#1{\mathit{#1}}$

#### 1. Executive Verdict

The Structural Novelty Index (SNI) is a two-phase quantitative framework designed to identify unconventional helper NLRs within the ~6,000-member NRC superfamily. It utilizes eight distinct structural parameters, derived from AlphaFold 3 (AF3) predictions, to measure deviations from the canonical hexameric resistosome architecture established by NbNRC2, SlNRC3, and NbNRC4. The phased approach—moving from high-throughput monomeric filtering (mSNI) to targeted multimeric validation (cSNI)—provides a scalable methodology for "structural phylogenetics."

**Verdict: Highly recommended for immediate implementation and testing.**

#### 2. Critical Flaws

No critical flaws found.

#### 3. Addressed Objections

The initial concerns regarding the lack of uniformity in experimental structural data have been effectively addressed. Experimental PDB structures for NRCs often feature missing N-terminal $\alpha1$ helices or inconsistent latch distances (ranging from 3.8 Å to 5.8 Å) due to protein flexibility or crystallization conditions. The SNI resolves this by employing an "AF3-on-AF3" baseline strategy. By using AF3-generated models of canonical NRCs (NRC2, NRC3, NRC4) as the reference set, the index ensures that comparisons are made between computationally consistent models, eliminating the noise inherent in comparing flexible predictions to incomplete experimental maps.

Additionally, concerns regarding the reliability of stoichiometry prediction in AlphaFold have been mitigated by recent benchmarking. Evidence confirms that AF3 can model the NRC2 hexamer with high confidence (RMSD ~0.937–1.72 Å). The proposed use of the Stoichiometric Preference Score ($S_{pref}$), which compares interface confidence (ipTM) across forced pentameric and hexameric states, is a scientifically sound application of the software’s internal scoring metrics to distinguish energetically favorable assemblies.

#### 4. Validated Risks & Limitations

- **The Monomer Assumption:** The Phase 1 screening (mSNI) assumes that the "readiness" for unconventional oligomerization is encoded within the monomeric resting geometry (e.g., the intrinsic NB-ARC swivel or LRR expansion). There remains a risk that some novel resistosome architectures only emerge during the massive conformational rearrangements of the activation process, which might not be fully captured in monomeric models, leading to potential false negatives.
- **Static Weighting and pLDDT Integration:** The current Z-score formula uses static weights ($w_i$). However, structural novelty scores should ideally be penalized in regions of low local confidence (pLDDT < 70). Without dynamic weighting that incorporates AF3's confidence scores, the SNI might interpret software modeling artifacts as biological novelty.
- **Exclusion of Resting-State Dimer Metrics:** Recent structural data (PDB: 8RFH/9CC8) confirms that canonical NRCs exist as homodimers in their inactive state. The SNI primarily focuses on the transition toward the active hexamer. By omitting the dimer interface as a parameter, the index may miss "atypical" NLRs that possess novel autoinhibitory mechanisms or bypass the dimer state entirely.

#### 5. Supporting Arguments & Evidence (Motivation)

- **Theoretical Basis:** The SNI is grounded in the molecular switch mechanism of NLR activation. The 180° rotation of the NB-HD1 module relative to the WHD-LRR module and the specific 10° shift in the NB-ARC interdomain angle are established physical determinants that distinguish hexameric NRCs from pentameric ancestors like ZAR1. The parameters (swivel, latch, divergence angle) map directly to these functional checkpoints.
- **Empirical Support:** The proposal is supported by the existence of biological outliers such as NRCX (a negative regulator lacking the MADA motif) and WAI3 (an octameric resistosome). These examples prove that the NLR family is not monolithic and that structural divergence correlates with novel immune functions. The availability of high-resolution ground truth structures (9FP6, 9RI9, 9CC8) provides the necessary coordinates to establish a robust mathematical baseline.
- **Comparative Advantage:** Traditional sequence-based screening (e.g., BLAST) often fails to identify functional homologs once sequence identity drops below 30%. The SNI provides a "functional proxy" through geometry. By focusing on the shape-dynamics of the NB-ARC and CC domains, the index can identify "structural dark matter"—proteins that have diverged in sequence but retain or evolve specific resistosome architectures.

#### 6. Alignment & Novelty

- **Alignment:** The hypothesis is perfectly aligned with the research goal to define a 5–10 parameter quantitative index that distinguishes unconventional NRC-NLRs from canonical hexamers using AF3 and ground truth PDB data.
- **Novelty:** While the individual metrics (angles, distances, and hydropathy) are derived from existing literature, the formalization of these metrics into a weighted, predictive, and automated diagnostic pipeline is a significant technical synthesis. This represents a transition from descriptive structural biology to proactive high-throughput screening, which is a novel contribution to the field of plant immunity.

#### 7. Feasibility Assessment (Go/No-Go Decision)

- **Resource Intensity:** Low. Phase 1 (mSNI) can be executed using existing structures from the AlphaFold Protein Structure Database (AFDB), requiring minimal compute. Phase 2 (cSNI) involves running approximately 500 AF3 hexamer models, a standard task for any laboratory with access to a GPU cluster (e.g., A100s).
- **Technical Complexity:** Moderate. The geometric calculations required (centroid-based angles and dihedral measurements) are standard in structural bioinformatics. Automation via Biopython or PyMOL scripts is straightforward.
- **Time to Verdict:** 2–3 weeks. A "Go/No-Go" experiment can be conducted by running the mSNI on known positive controls (NRCX and ZAR1) against the NRC2/3/4 baseline. If the index correctly identifies these known outliers, the full 6,000-sequence screen is justified.

#### 8. Conclusion

The Structural Novelty Index (SNI) is a clinically precise and well-grounded framework that leverages the accuracy of AlphaFold 3 to automate the discovery of unconventional NLRs. By formalizing the geometric "rules" of the NRC resistosome, it provides a necessary tool for navigating the vast sequence diversity of the Solanaceae immune network. While the monomeric filtering phase carries an inherent risk of missing activation-dependent novelties, the methodology's scalability and its sophisticated "AF3-on-AF3" baseline strategy make it a highly valuable study for the field of plant immunology.

#### Research contacts

$\def\mathcal#1{\mathit{#1}}\def\mathscr#1{\mathit{#1}}$

Based on the provided research articles, here are the specific researchers who are experts in the field and could review the proposed **Structural Novelty Index (SNI)**. The list is ordered by those who have looked at the most specific structural and quantitative aspects of the hypothesis (hexameric NRC structures and AlphaFold 3 metrics).

#### 1. Jogi Madhuprakash

- **Justification:** Madhuprakash is the lead researcher who determined the cryo-EM structure of the **NbNRC2 hexameric resistosome (PDB 9FP6)**, which is the primary "canonical" reference for the SNI. He is uniquely qualified to review metrics regarding protomer interface angles and the stabilization of the hexameric assembly.
- **Support:** Abstract 1/4 ("Structure of the NbNRC2 hexameric resistosome") notes that Madhuprakash determined the structure and performed comparative structural analyses between the resting state homodimer and sensor-activated homohexamer, identifying key interprotomer interactions.

#### 2. Michael W. Webster

- **Justification:** A key author of the NbNRC2 hexamer study, Webster is an expert in the structural biology of the NRC family. He deposited the ground truth structures (9FP6) that the SNI seeks to use as a baseline.
- **Support:** Abstract 1/4 lists Webster as a deposition author for the NbNRC2 hexameric resistosome and credits him with work shedding light on the "NLR structural diversity" and the assessment of AlphaFold 3.

#### 3. AmirAli Toghani

- **Justification:** Toghani is a primary expert in using **AlphaFold 3 quantitative metrics** (such as pTM, ipTM, and pLDDT) to classify NLRs. His work directly mirrors the SNI's goal of using AF3 screening to distinguish unconventional NLR configurations from standard models.
- **Support:** Abstract 5 ("Can AI modelling... distinguish between sensor and helper NLR immune receptors?") and Abstract 7 describe Toghani's use of "key structural metrics, including pLDDT, pTM, ipTM, and per-chain pTM" to differentiate between helper and sensor NLRs based on their ability to form resistosomes.

#### 4. Sophien Kamoun

- **Justification:** As a senior author on nearly all provided articles, Kamoun is the leading expert on the **NRC network** and the transition of NRCs from dimers to hexamers. He has overseen the discovery of the MADA motif and the structural characterization of NRC2, NRC3, and NRC4.
- **Support:** Kamoun is a lead author in Abstracts 1, 3, 4, 5, 6, and 7, which cover the entire scope of the NRC family, from sequence-based motifs (MADA) to hexameric structural determination and AF3 benchmarking.

#### 5. Hiroaki Adachi

- **Justification:** Adachi defined the **MADA motif** (Parameter 1 and 6 of the SNI). His expertise is essential for reviewing whether the SNI accurately captures "N-terminal CC-domain variations" and "non-MADA/unconventional execution" signatures.
- **Support:** Abstract 3 ("An N-terminal motif... is functionally conserved") describes Adachi’s work in identifying the 21-amino acid MADA consensus sequence and its role as a "death switch" in helper NLRs like NRC4.

#### 6. Ibrahim Tarhan

- **Justification:** Tarhan specializes in the **pore funnel hydrophobicity and geometry** of unconventional NLRs (specifically CC_R-NLRs like NRG1). He is a perfect fit for reviewing SNI Parameter 6 (Pore Funnel Hydrophobicity Gradient).
- **Support:** Abstract 6 ("A helper NLR targets organellar membranes") details Tarhan's work using AF3 to predict "significantly longer funnel-shaped CC regions" and distinct N-terminal sequence patterns that deviate from canonical NRCs.

#### 7. Tolga O. Bozkurt

- **Justification:** Bozkurt is an expert in the **cellular dynamics and membrane targeting** of the NRC resistosomes. He can review whether the SNI's structural parameters (like interface burial efficiency) correlate with actual biological membrane association.
- **Support:** Abstract 4 and 6 list Bozkurt as a key researcher in the structural characterization of the NRC2 hexamer and the study of how helper NLRs target membranes to trigger immunity.

#### 8. Mahinur S. Akkaya (or Nan Wu)

- **Justification:** These researchers have experience modeling the **NB-ARC domain and MHD motif** using AlphaFold (relevant to SNI Parameter 3). They have specifically looked at how single-residue changes in the NB-ARC domain (like the "swivel" or "latch" proximity) affect structural stability and auto-activation.
- **Support:** Abstract 2/8 ("Assessment of Self-Activation... of Wheat Coiled-Coil Domain Containing NLR") describes their use of AlphaFold 2 to predict structural changes in the MHD motif region and the resulting impacts on intramolecular interactions.

### Appendix:

**All reviews:**

**Correctness:**

$\def\mathcal#1{\mathit{#1}}\def\mathscr#1{\mathit{#1}}$

#### Related Articles

1. **[1] RCSB PDB - 9FP6: Structure of the NbNRC2 hexameric resistosome**: Essential for establishing the canonical hexameric baseline for NbNRC2, confirming its architecture differs from previous pentameric models.
2. **[17] RCSB PDB - 9CC8: Hexameric state of the NRC4 resistosome**: Provides the second ground-truth hexameric structure (NRC4), validating the hexameric nature of the helper NRC family.
3. **[14] A plant pathogen effector blocks stepwise assembly of a helper NLR resistosome**: References 9RI9 as the SlNRC3 hexamer structure, completing the three-member ground-truth set required by the goal.
4. **[3] An N-terminal motif in NLR immune receptors is functionally conserved across distantly related plant species**: Establishes the MADA motif (residues 1–29) as the functional executioner unit and its alpha-helical nature, supporting Parameter 1 and 6.
5. **[5] Can AI modelling of protein structures distinguish between sensor and helper NLR immune receptors?**: Validates the use of AlphaFold 3 (AF3) confidence scores (pTM/ipTM) to distinguish executioner helpers from sensors, supporting Parameter 8.
6. **[18] Accurate Prediction of Protein Complex Stoichiometry by Integrating AlphaFold3 and Template Information**: Supports the methodology of using AF3 to rank different stoichiometric candidates, validating the logic of Parameter 8 ($S_{pref}$).
7. **[12] A disease resistance protein triggers oligomerization of its NLR helper into a hexameric resistosome to mediate innate immunity**: Provides quantitative geometric comparisons between NRC2 (hexamer) and ZAR1 (pentamer), supporting the use of protomer angles (Parameter 5).
8. **[21] Can AI modelling of protein structures distinguish between sensor and helper NLR immune receptors?**: Directly supports Parameter 8 by showing that helpers consistently form stable funnels with higher confidence scores than sensors.
9. **[10] A hydrophobic core in the coiled-coil domain essential for NRC resistosome function**: Discusses the transition from resting dimer to active hexamer, providing biological context for the Swivel angle (Parameter 3).
10. **[22] A helper NLR targets organellar membranes to trigger immunity**: Shows that unconventional NLRs (like NRG1) have different N-terminal architectures, supporting the premise of the SNI to find "unconventional" receptors.
11. **[7] A hierarchical immune receptor network in lettuce reveals contrasting patterns of evolution in sensor and helper NLRs**: Confirms that NRC-sensors (NRC-S) fail to form resistosomes in AF3, justifying the "Novelty" threshold in Parameter 8.
12. **[13] An N-terminal motif in NLR immune receptors is functionally conserved across distantly related plant species - PMC**: Elaborates on the "use-it-or-lose-it" model where sensors lose the MADA motif, supporting the hydropathy and helical propensity parameters.
13. **[16] A disease resistance protein triggers oligomerization of its NLR helper into a hexameric resistosome to mediate innate immunity**: Provides the specific pore residue data (L126 for NRC2) used to evaluate the pore funnel metrics (Parameter 6).
14. **[20] Oligomerization-mediated autoinhibition and cofactor binding of a plant NLR**: Highlights the importance of the LRR inner surface and IP6 binding, supporting the "Latch" metric (Parameter 2).
15. **[4] A disease resistance protein triggers oligomerization of its NLR helper into a hexameric resistosome to mediate innate immunity - PMC**: Compares the inter-domain angles of NRC2 vs ZAR1, providing the basis for Parameter 3.

#### Detailed Assumptions

1. **Canonical Geometric Consistency:** Assumes NRC2, NRC3, and NRC4 share a conserved "Canonical Mean" for stoichiometry (60° protomer angle) and NB-ARC swivel orientation.
2. **AF3 Confidence as a Physical Proxy:** Assumes that AF3 ipTM and pTM scores accurately reflect the physical stability and stoichiometric preference of a sequence.
3. **Monomer-to-Multimer Correlation:** Assumes that specific monomeric deviations (mSNI) in the MADA helix or ARC2-LRR latch are predictive of larger multimeric architectural shifts.
4. **"AF3-on-AF3" Validity:** Assumes that the best way to avoid modeling bias is to compare AF3 models of unknown sequences against AF3 models of the ground truth (NRC2/3/4) rather than directly against experimental PDB coordinates.
5. **Domain Boundary Accuracy:** Assumes the ARC2 (WHD) domain residues 392–497 and N-terminal residues 1–30 are universal enough across the NRC family to serve as fixed anchors for distance metrics.

#### Reasoning about Correctness of Assumptions

1. **Canonical Geometry:** **Correct.** Abstract [12] and [16] confirm NbNRC2 is a hexamer with distinct inter-domain angles compared to pentamers. PDB analysis confirms 9FP6 and 9CC8 exhibit angles of ~60.0°.
2. **AF3 Confidence Score Proxy:** **Correct.** Abstracts [5], [18], and [21] provide strong evidence that AF3 ranking scores ($S_{pref}$) correctly identify the true stoichiometry and functional state of NLRs.
3. **Monomer-to-Multimer Correlation:** **Likely True.** The transition from dimer/monomer to resistosome involves massive rearrangements of the NB-ARC [4]. Monomeric "strain" or shifts in the swivel angle [12] are reasonable precursors to stoichiometric divergence.
4. **AF3-on-AF3 Approach:** **Highly Scientific.** As noted in the PDB analysis report, experimental structures often lack coordinates for the N-terminal $\alpha1$ helix (MADA motif). Using AF3 models of the canonical set as the baseline ensures Parameter 1 and 6 are calculable and that software artifacts are normalized.
5. **Domain Boundaries:** **Correct.** The Knowledge Base confirms residues 392–497 accurately represent the "latch" area in canonical NRCs.

#### Strength of Evidence

- **Direct Evidence:** Cryo-EM structures (9FP6, 9CC8, 9RI9) provide the precise 60° dihedral angle and the specific latch distances required for Phase 2 validation. Sequence data for SlNRC3 and NbNRC4 directly supports the hydrophobicity gradient (FHG) baseline [Knowledge Base].
- **Indirect Evidence:** The "use-it-or-lose-it" evolutionary model [3, 13] supports the idea that sensors/unconventional receptors will deviate in the MADA motif (Parameter 1/6). Recent studies on stoichiometry prediction [18] and sensor/helper differentiation [5, 21] provide high confidence in the ipTM contrast method (Parameter 8).

#### Suggested Improvements

1. **Expanded Reference Set:** While using 9FP6, 9RI9, and 9CC8 is great, the Weighted Z-Score calculation would benefit from a slightly larger "AF3 Ref" set (e.g., 10–15 confirmed canonical NRC orthologs from the Solanaceae dataset) to establish a more stable Standard Deviation ($SD_{AF3\_Ref}$).
2. **Explicit Motif Anchoring:** For Parameter 3 ($\phi_{swivel}$), explicitly anchor the centroids to the P-loop (GxxxxGK[T/S]) and the MHD motif rather than just "NBD" and "ARC2" to ensure robustness against length variations in the ~6,000 sequence dataset.
3. **Memory Management:** Address the hardware constraints of modeling 500 hexamers (Phase 2). Suggesting the use of truncated CC-NB-ARC models (as seen in Abstract [5]) would improve computational throughput.

#### Assessment of Goal Requirements

- **5–10 Parameters:** Met (8 parameters).
- **Measurable from AF3/PDB:** Met (All parameters use Euclidean distances, angles, or confidence scores).
- **Distinguish unconventional from canonical NRCs:** Met (Metrics cover stoichiometry, MADA executioner function, and auto-inhibition).
- **Incorporate Ground Truth (9FP6, 9RI9, 9CC8):** Met (Used to derive canonical means).
- **High-throughput design:** Met (Phased screening funnel significantly reduces GPU cost).
- **Specific metrics (Interface angles, CC variations, NB-ARC patterns):** Met (P5, P1/P6, and P3 respectively).

#### Final Reasoning and Recommendation

The Structural Novelty Index (SNI) is a highly sophisticated, technically sound proposal. It correctly identifies the primary bottleneck in high-throughput structural screening (computational cost) and provides a scientifically rigorous "Screening Funnel" to solve it. The "AF3-on-AF3" implementation strategy is particularly brilliant, as it overcomes the common problem of disordered N-termini in experimental structures. The inclusion of stoichiometric contrast modeling (Pentamer vs Hexamer fit) aligns with the latest literature on NLR classification. The parameters are grounded in specific structural biology motifs rather than simple sequence identity, fulfilling the goal's core rationale. The idea is ready for testing and likely correct.

Answer: 9

**Novelty:**

$\def\mathcal#1{\mathit{#1}}\def\mathscr#1{\mathit{#1}}$

This review evaluates the novelty of the proposed **Structural Novelty Index (SNI)** for the screening and classification of unconventional NRC-NLRs within the Solanaceae family using AlphaFold 3 (AF3).

#### Related Article Abstract Titles

1. **[1] RCSB PDB - 9FP6: Structure of the NbNRC2 hexameric resistosome:** Essential ground truth for canonical NRC2 hexameric stoichiometry and protomer arrangement.
2. **[4] A disease resistance protein triggers oligomerization of its NLR helper into a hexameric resistosome to mediate innate immunity:** Provides detailed structural analysis of NbNRC2, confirming its hexameric nature and the flexibility of the N-terminal α1 helix.
3. **[5] Can AI modelling of protein structures distinguish between sensor and helper NLR immune receptors? (Nov 2024):** Directly relates to the idea by using AF3 confidence scores (pTM/ipTM) and structural features (funnel formation) to classify NLRs in silico.
4. **[14] A plant pathogen effector blocks stepwise assembly of a helper NLR resistosome (July 2025):** Provides the structure of SlNRC3 (9RI9) and describes the structural dynamics of helper NLR activation.
5. **[17] RCSB PDB - 9CC8: Hexameric state of the NRC4 resistosome:** Essential ground truth for canonical NRC4 hexameric stoichiometry.
6. **[18] Accurate Prediction of Protein Complex Stoichiometry by Integrating AlphaFold3 and Template Information (Jan 2025):** Describes "PreStoi," a method leveraging AF3 scores to predict multimer stoichiometry, highly relevant to Parameter 8 ($S_{pref}$).
7. **[3] An N-terminal motif in NLR immune receptors is functionally conserved across distantly related plant species:** Discovered the MADA motif and its structural requirement for α1 helix formation, underpinning Parameters 1 and 6.
8. **[37] A hierarchical immune receptor network in lettuce reveals contrasting patterns of evolution in sensor and helper NLRs (Feb 2025):** Uses AF3 to distinguish NRC-H from NRC-S based on their capacity to form higher-order resistosome-like oligomers and metrics like pTM/ipTM.
9. **[20] Oligomerization-mediated autoinhibition and cofactor binding of a plant NLR:** Describes structural basis for autoinhibition in resting NRC2 dimers, relevant to Parameter 2 ($D_{latch}$).
10. **[44] Subfunctionalization of NRC3 altered the genetic structure of the Nicotiana NRC network:** Maps critical residues on NRC3 homodimer surfaces, relevant to interface burial and conformational swivel analyses.
11. **[12] Cryo-EM structure of the NbNRC2 resistosome reveals a homohexameric oligomer:** Detailed analysis of inter-protomer interactions and conformational rearrangements.
12. **[16] Comparison of NbNRC2, AtZAR1, and TmSr35 resistosomes reveals striking structural differences:** Quantitative comparison of protomer angles and pore characteristics (relevant to Parameters 3 and 5).
13. **[32] The N-terminal domains of NLR immune receptors exhibit structural and functional similarities across divergent plant lineages:** Explores the MAEPL motif as an alternative to MADA, relevant to the "unconventional" detection goal of the SNI.
14. **[26] Structural Insights into the Plant Immune Receptors PRRs and NLRs:** Review of ZAR1 resistosome assembly and pore formation logic.
15. **[31] Diversification of the “EDVID” packing motif underpins structural and functional variation in plant NLR coiled-coil domains:** Structural analysis of motifs stabilizing the CC domain during resistosome assembly.

#### Aspects of the Idea Already Explored

- **Stoichiometric Prediction via AF3:** Using AF3 to distinguish between pentameric and hexameric states (Parameter 5 and 8) is already established. Abstract **[18]** describes a method specifically integrating AF3 scores to rank stoichiometry. Abstracts **[4]** and **[12]** already performed comparative structural analysis between pentameric ZAR1 and hexameric NRC2.
- **Classification by AF3 Confidence Scores:** The core logic of "Complex SNI" (Phase 2) is nearly identical to work by Toghani et al. **[5, 11]** and Pai et al. **[37]**. These studies explicitly state that "helper NLRs consistently exhibit higher AlphaFold 3 confidence scores than sensors" and use the presence/absence of funnel-shaped structures to categorize proteins.
- **MADA Motif Propensity:** Parameter 1 ($\alpha_{MADA}$) is based on the discovery by Adachi et al. **[3]**. The use of pLDDT to verify funnel formation in AF3 is already described in **[4], [5],** and **[37]** as a reliable classification metric.
- **NB-ARC Swivel and Interface Analysis:** The conformational shift of the NB-ARC domains (Parameter 3) and inter-protomer electrostatic interactions (Parameter 7) were quantified in the ground truth research for NbNRC2 **[4, 12, 16]**, which defined the ~180° rotation and specific burial areas.
- **Autoinhibition Baselines:** The distance between subdomains (ARC2-LRR) to maintain autoinhibition (Parameter 2) is a standard metric in NLR structural biology, explored in the inactive dimers described in **[20]**.

#### Novel Aspects of the Idea

- **The SNI Formalism:** The specific integration of these disparate structural observations into a single **weighted Z-score index** ($SNI$) specifically for high-throughput outlier detection in the NRC family is not explicitly formalized in the provided articles.
- **Parameter 4 (LRR Expansion Ratio):** The simple ratio of conserved Leucine distances to sequence length ($\chi_{LRR}$) is a novel, computationally lightweight descriptor for LRR topology compared to complex circle-fitting algorithms.
- **Parameter 6 (FHG):** Defining a hydrophobicity gradient specifically across the tip vs. base of the $\alpha1$ helix to identify organelle-specific targeting vs. plasma membrane pore formation is a novel specialized application of the Kyte-Doolittle scale to AF3 models.
- **AF3-on-AF3 Standardization:** The strategy of deriving "Canonical Means" by running the reference PDB sequences (NRC2/3/4) through the AF3 multimer algorithm *before* comparison—to eliminate software bias—is a rigorous bioinformatic refinement not emphasized in the classification preprints.

#### Novelty Review

The idea is **moderately novel** in its specific implementation but **limited** in its core conceptual drive.

While the "SNI" index is a new tool, its constituent parts are largely derived from the current state-of-the-art literature published in 2024 and 2025. Specifically, the use of AF3 multimer metrics (ipTM, pTM) and funnel propensity to structurally categorize the ~6,000 NRC sequences is the current operating procedure for the field's leaders (e.g., the Kamoun and Wu labs in **[5], [11], [37]**). The SNI essentially formalizes their qualitative observations into a quantitative screening funnel. The most distinct novel element is the mathematical weighting of these parameters to detect "unconventionality" rather than just "helper-ness."

#### Reasoning and Recommendation

The formalization of structural biology insights into a multi-parametric index is a valuable technical contribution. However, the idea's core components—detecting stoichiometry shifts, funnel confidence, and interface burial—are the exact methods used in recent preprints **[5, 18, 37]** to solve the same problem (NLR classification).

**Recommendation:** The idea should be tested as a benchmarking tool for the community. Its value lies in the "Standardized Baselines" (AF3 vs AF3) and the weighted formula, which could provide more reproducibility than current qualitative AF3 model assessments. However, it does not challenge existing paradigms; it formalizes them.

**Answer: 4** (Limited Novelty: Core mechanisms and entities are currently being explored/published in this specific context by experts, though the multi-parametric Z-score tool is a new specific combination).

**Feasibility:**

$\def\mathcal#1{\mathit{#1}}\def\mathscr#1{\mathit{#1}}$

#### Related Article Abstracts

1. **[1] RCSB PDB - 9FP6: Structure of the NbNRC2 hexameric resistosome (2024):** Provides the essential ground truth coordinates for the canonical NRC2 resistosome, necessary for calculating baseline parameters.
2. **[12] A disease resistance protein triggers oligomerization of its NLR helper into a hexameric resistosome to mediate innate immunity (2024):** Compares NbNRC2 (hexamer) with AtZAR1 (pentamer) and highlights specific interdomain angular differences (e.g., a 10° shift in NB-ARC), directly supporting the feasibility of Parameter 3 and 5.
3. **[14] A plant pathogen effector blocks stepwise assembly of a helper NLR resistosome (2025):** Confirms the hexameric structure of SlNRC3 (PDB ID: 9RI9), providing the second canonical ground truth for the SNI reference set.
4. **[17] RCSB PDB - 9CC8: Hexameric state of the NRC4 resistosome (2024):** Provides the third ground truth structure (NRC4), completing the canonical reference set for Solanaceae NRCs.
5. **[13] An N-terminal motif in NLR immune receptors is functionally conserved across distantly related plant species (2019):** Defines the MADA motif and its conservation, which is the biological basis for Parameter 1 ($\alpha_{MADA}$) and Parameter 6 (FHG).
6. **[5] Can AI modelling of protein structures distinguish between sensor and helper NLR immune receptors? (2024):** Demonstrates that AlphaFold 3 (AF3) confidence scores (pTM/ipTM) and structural features (funnel formation) reliably distinguish executioner helpers from sensor NLRs, validating the utility of Parameter 8 ($S_{pref}$).
7. **[7] A hierarchical immune receptor network in lettuce reveals contrasting patterns of evolution in sensor and helper NLRs (2025):** Shows that NRC-S (sensors) fail to form funnel structures in AF3, providing a negative control population to test the SNI’s specificity.
8. **[18] Accurate Prediction of Protein Complex Stoichiometry by Integrating AlphaFold3 and Template Information (2025):** Describes methods for using AF3 to predict multimeric stoichiometry, supporting the feasibility of testing Parameter 8.
9. **[8] Assessment of Self-Activation and Inhibition of Wheat CC-NLR Immune Receptor Yr10 (2025):** Utilizes AF3 to model conformational changes in the MHD motif and N-terminal helices, mirroring the methodology required for Parameter 2 ($D_{latch}$).
10. **[20] Oligomerization-mediated autoinhibition and cofactor binding of a plant NLR (2024):** Describes the resting state dimer/tetramer interfaces of SlNRC2, essential for verifying the "Novelty" of active-state models in Parameter 7.
11. **[10] A hydrophobic core in the coiled-coil domain essential for NRC resistosome function (2025):** Identifies specific CC-domain residues critical for function, providing anchors for structural metrics in Parameter 1 and 6.
12. **[9] How to assess the quality of AlphaFold 3 predictions (2025):** Details the interpretation of pTM and ipTM scores, which are the primary data points for calculating $S_{pref}$ and weighting the SNI.
13. **[4] A disease resistance protein triggers oligomerization of its NLR helper into a hexameric resistosome... (2024):** Validates that AF3 can predict N-terminal $\alpha1$ helices with high confidence even when experimental structures cannot, which is critical for Parameter 1.
14. **[6] A helper NLR targets organellar membranes to trigger immunity (2024):** Shows that AF3 predicts significantly longer funnel structures for certain helper NLRs, justifying the inclusion of the LRR expansion and hydrophobicity metrics.
15. **[21] Can AI modelling of protein structures distinguish between sensor and helper... (2024):** Provides quantitative benchmarks for "Helper" vs "Sensor" pTM scores, useful for defining the "Novelty" threshold in the SNI formula.

#### Steps to Test the Idea

1. **Baseline Benchmarking (Canonical Reference):** Run AlphaFold 3 Multimer on NbNRC2, SlNRC3, and NbNRC4. Extract the 8 parameters for these models to calculate the *Canonical Mean* ($\mu$) and *Standard Deviation* ($\sigma$) for each metric. This eliminates discrepancies between experimental PDB coordinates and AF3-predicted coordinates.
2. **High-Throughput Monomer Screen (mSNI):** Extract or predict monomers for the ~6,000 NRC sequences (utilizing the AlphaFold Database where possible). Use a Python script (Biopython/DSSP) to calculate $\alpha_{MADA}$, $D_{latch}$, $\phi_{swivel}$, and $\chi_{LRR}$ for the entire dataset.
3. **Outlier Selection:** Apply the Weighted Z-Score formula to the monomer data. Select the top 500 sequences with the highest mSNI (indicating structural divergence).
4. **Multimer Modeling (cSNI):** Run AF3 Multimer on the 500 candidates. For each sequence, generate both a hexameric and a pentameric model (contrastive modeling).
5. **Complex Parameter Extraction:** Calculate $\Delta\theta_{int}$, FHG, $\eta_{inter}$, and $S_{pref}$ from the multimer outputs.
6. **Sensitivity/Specificity Validation (Go/No-Go):**
   - **Test Case A (Helper Outlier):** Include Arabidopsis **ZAR1** in the pipeline. *Expected result:* SNI should flag it as novel due to its pentameric stoichiometry ($72^\circ$ angle) and swivel angle deviation.
   - **Test Case B (Known Sensors):** Include known sensors (Rx, Bs2). *Expected result:* SNI should flag them as novel/degenerate due to lack of MADA helix propensity and FHG inversion.
   - **Go/No-Go Criterion:** If the SNI fails to distinguish ZAR1 from NRC2/3/4 at a significance level of $p < 0.01$, the parameters must be redefined.

#### Feasibility Reasoning

The defined Structural Novelty Index is highly feasible due to its tiered computational strategy. By using a "Monomeric Filter" (mSNI) first, the proposal avoids the massive GPU cost of modeling 6,000 hexamers (~54,000,000 residues total), which would be prohibitive.

1. **Computational Scalability:** Modeling 500 hexamers is a standard bioinformatic task. While the NRC hexamer (~5,400 residues) slightly exceeds the 5,000-token limit of the AF3 web server, it is well within the capacity of local AF3 installations on A100 GPU clusters, which are standard in modern structural biology labs.
2. **Metric Measurability:** All parameters rely on standard structural outputs. Parameters like "swivel angle" and "interface burial efficiency" can be automated using simple geometric scripts in PyMOL or Biopython. The usage of AF3-on-AF3 comparisons (Step 1) is a statistically sound way to account for the "Missing Residue" problem in PDBs 9FP6/9RI9/9CC8.
3. **Ground Truth Alignment:** The proposal leverages specific 2024 structural data. The use of the $10^\circ$ swivel window and $60^\circ$ vs $72^\circ$ inter-protomer angles matches the comparative data reported in the most recent literature [12].
4. **Potential Bottleneck:** Developing the custom script for the "ARC2-LRR Latch Proximity" ($D_{latch}$) across 6,000 sequences requires precise domain mapping (using HMMs or conserved motifs like the MHD motif) to ensure consistent residue range selection across diverse Solanaceae species. However, this is a standard bioinformatic challenge.

The idea is computationally efficient and grounded in measurable, biologically relevant structural metrics.

Answer: 7

**Impact potential:**

$\def\mathcal#1{\mathit{#1}}\def\mathscr#1{\mathit{#1}}$

This review evaluates the impact potential of the **Structural Novelty Index (SNI)**, a quantitative framework designed to screen and classify ~6,000 sequences in the NRC-NLR family to identify unconventional immune receptors.

#### 1. Related Article Abstracts

The following 11 abstracts are most relevant for assessing the impact of the SNI:

1. **[1] Structure of the NbNRC2 hexameric resistosome (9FP6):** Provides the fundamental "canonical" hexameric baseline for the NRC2 clade.
2. **[3] An N-terminal motif in NLR immune receptors is functionally conserved (MADA):** Defines the MADA motif (residues 1–29) used in Parameter 1 and 6.
3. **[4] A disease resistance protein triggers oligomerization... into a hexameric resistosome:** Assesses AlphaFold 3 (AF3) performance on NbNRC2, confirming its ability to model the N-terminal α1 helix which is often missing in PDBs.
4. **[5] Can AI modelling of protein structures distinguish between sensor and helper NLR immune receptors?:** Directly supports the feasibility of Parameter 8 ($S_{pref}$) by showing helpers have higher confidence scores in oligomeric states.
5. **[6] A helper NLR targets organellar membranes to trigger immunity:** Shows that non-canonical NLRs (NRG1/ADR1) have extended N-termini, justifying parameters looking for "unconventional" N-terminal variations.
6. **[7] A hierarchical immune receptor network in lettuce...:** Demonstrates that sensor NLRs (NRC-S) fail to form resistosome-like structures in AF3, providing a negative control for the SNI.
7. **[10] A hydrophobic core in the coiled-coil domain essential for NRC resistosome function:** Details the transition from cytosolic dimers to membrane-associated hexamers, relevant for "swivel" and "latch" metrics.
8. **[14] A plant pathogen effector blocks stepwise assembly... (9RI9):** Provides the ground-truth structure for SlNRC3 and discusses sub-hexameric intermediates.
9. **[17] RCSB PDB - 9CC8: Hexameric state of the NRC4 resistosome:** Provides the third ground-truth structure for the canonical set.
10. **[18] Accurate Prediction of Protein Complex Stoichiometry by Integrating AlphaFold3...:** Validates the methodology of using AF3 ranking scores to determine stoichiometric preference (Parameter 8).
11. **[20] Oligomerization-mediated autoinhibition and cofactor binding...:** Identifies inositol phosphate binding in the LRR concave surface, crucial for the "latch" metric (Parameter 2).

#### 2. Key Assumptions

To achieve its potential impact, the SNI depends on several critical assumptions:

1. **Modeling Accuracy:** AlphaFold 3 can accurately distinguish between genuine biological "novelty" and modeling artifacts or "forced folding" in 6,000 uncharacterized sequences.
2. **State-Function Correlation:** Monomeric structural deviations ($mSNI$) are robustly predictive of functional/stoichiometric divergence in the multimeric state ($cSNI$).
3. **Computational Feasibility:** A tiered approach (mSNI filtering before cSNI) makes the high-throughput screening of the NRC family technically possible with current hardware.
4. **Baseline Consistency:** Using AF3-on-AF3 comparisons (rather than AF3-on-PDB) successfully eliminates software-specific bias and addresses missing residues in experimental structures.

#### 3. Feasibility and Impact Analysis

- **Assumption 1 (Accuracy):** High feasibility. Recent studies [4, 12] show AF3 models the N-terminal α1 helix with high confidence, even when experimental cryo-EM fails. The inclusion of Parameter 8 ($S_{pref}$) and weighted scores helps distinguish biological signals from modeling noise.
- **Assumption 2 (Monomer-to-Multimer):** Moderate feasibility. While [10] shows massive rearrangements from resting dimers to hexamers, the "seed" of autoinhibition resides in the ARC2-LRR latch and NB-ARC orientation. Using monomeric metrics as a "primary filter" is a statistically sound way to prioritize sequences for expensive multimeric runs.
- **Assumption 3 (Scalability):** High feasibility. Modeling 6,000 hexamers would require thousands of GPU hours. The "funnel" approach (6,000 monomers $\rightarrow$ 500 multimers) is the only realistic way to process the Solanaceae dataset.
- **Assumption 4 (Baselines):** High feasibility. The "missing residue" problem in PDBs 9FP6 and 9RI9 is a known issue. By generating AF3 models of the reference set (NbNRC2, SlNRC3, NbNRC4) to set the "mean," the index ensures a mathematically fair comparison [4, 16].

#### 4. Suggested Improvements

- **Confidence Penalty:** Incorporate a "pLDDT Penalty" into the formula. If a structure appears "novel" but has an average pLDDT < 50, the SNI should be downgraded to reflect a likely modeling failure.
- **Cofactor Integration:** Since SlNRC2 requires inositol phosphate (IP) for autoinhibition [20], Parameter 2 ($D_{latch}$) should be augmented to measure the volume of the binding pocket or the conservation of IP-interacting residues.
- **Interface Shape Complementarity ($S_c$):** Add a metric for the "tightness" of the fit between protomers. If a sequence is forced into a hexamer but has high "structural frustration," it is a strong indicator of a non-hexameric preference.
- **Anchor-Based Geometry:** Ensure the Swivel Angle ($\phi_{swivel}$) uses centroids of conserved motifs (e.g., P-loop to MHD) rather than residue indices, to account for insertions/deletions in the 6,000 sequences.

#### 5. Overall Impact Potential

The proposed Structural Novelty Index has **transformative impact potential**. The NRC family represents up to 50% of the NLRome in many Solanaceae species [23], yet our understanding is limited to a few "canonical" members. By defining a robust, 8-parameter mathematical baseline that leverages the latest AI (AF3) and ground-truth structural data, this idea moves NLR biology from qualitative sequence similarity to quantitative structural topology.

The tiered screening approach makes it computationally viable to explore thousands of sequences, likely uncovering entirely new resistance mechanisms (e.g., pentameric NRCs, non-pore-forming NRCs, or organelle-targeted helpers). This would stimulate new research directions in crop engineering and plant-pathogen coevolution.

**Answer: 9**

References:

[1] [RCSB PDB - 9FP6: Structure of the NbNRC2 hexameric resistosome](https://www.rcsb.org/structure/9fp6)

[2] [Assessment of Self-Activation and Inhibition of Wheat Coiled-Coil Domain Containing NLR Immune Receptor Yr10CG - PMC](https://pmc.ncbi.nlm.nih.gov/articles/PMC11768854/)

[3] [An N-terminal motif in NLR immune receptors is functionally conserved across distantly related plant species | eLife](https://elifesciences.org/articles/49956)

[4] [A disease resistance protein triggers oligomerization of its NLR helper into a hexameric resistosome to mediate innate immunity - PMC](https://pmc.ncbi.nlm.nih.gov/articles/PMC11540030/)

[5] [Can AI modelling of protein structures distinguish between sensor and helper NLR immune receptors? | bioRxiv](https://www.biorxiv.org/content/10.1101/2024.11.24.625045v1.full-text)

[6] [A helper NLR targets organellar membranes to trigger immunity](https://www.biorxiv.org/content/10.1101/2024.09.19.613839v1)

[7] [A hierarchical immune receptor network in lettuce reveals contrasting patterns of evolution in sensor and helper NLRs](https://www.biorxiv.org/content/10.1101/2025.02.25.639832v1)

[8] [(PDF) Assessment of Self-Activation and Inhibition of Wheat Coiled-Coil Domain Containing NLR Immune Receptor Yr10CG](https://www.researchgate.net/publication/388212679_Assessment_of_Self-Activation_and_Inhibition_of_Wheat_Coiled-Coil_Domain_Containing_NLR_Immune_Receptor_Yr10CG)

[9] [How to assess the quality of AlphaFold 3 predictions | AlphaFold](https://www.ebi.ac.uk/training/online/courses/alphafold/alphafold-3-and-alphafold-server/how-to-assess-the-quality-of-alphafold-3-predictions/)

[10] [A hydrophobic core in the coiled-coil domain essential for NRC resistosome function](https://www.biorxiv.org/content/10.1101/2025.01.21.634219v1)

[11] [Can AI modelling of protein structures distinguish between sensor and helper NLR immune receptors?](https://www.biorxiv.org/content/10.1101/2024.11.24.625045v1)

[12] [A disease resistance protein triggers oligomerization of its NLR helper into a hexameric resistosome to mediate innate immunity](https://www.ncbi.nlm.nih.gov/pmc/articles/PMC11540030/)

[13] [An N-terminal motif in NLR immune receptors is functionally conserved across distantly related plant species - PMC](https://pmc.ncbi.nlm.nih.gov/articles/PMC6944444/)

[14] [A plant pathogen effector blocks stepwise assembly of a helper NLR resistosome](https://www.biorxiv.org/content/10.1101/2025.07.14.664264v1.full.pdf)

[15] [A hierarchical immune receptor network in lettuce reveals contrasting patterns of evolution in sensor and helper NLRs](https://www.biorxiv.org/content/10.1101/2025.02.25.639832v1)

[16] [A disease resistance protein triggers oligomerization of its NLR helper into a hexameric resistosome to mediate innate immunity](https://www.biorxiv.org/content/10.1101/2024.06.18.599586v1)

[17] [RCSB PDB - 9CC8: Hexameric state of the NRC4 resistosome](https://www.rcsb.org/structure/9cc8)

[18] [Accurate Prediction of Protein Complex Stoichiometry by Integrating AlphaFold3 and Template Information - PMC](https://pmc.ncbi.nlm.nih.gov/articles/PMC11761747/)

[19] [A disease resistance protein triggers oligomerization of its NLR helper into a hexameric resistosome to mediate innate immunity](https://www.ncbi.nlm.nih.gov/pmc/articles/PMC11540030/)

[20] [Oligomerization-mediated autoinhibition and cofactor binding of a plant NLR](https://pubmed.ncbi.nlm.nih.gov/38866053)

[21] [Can AI modelling of protein structures distinguish between sensor and helper NLR immune receptors?](https://www.biorxiv.org/content/10.1101/2024.11.24.625045v1)

[22] [A helper NLR targets organellar membranes to trigger immunity](https://www.biorxiv.org/content/10.1101/2024.09.19.613839v1)

[23] [Plant pathogens convergently evolved to counteract redundant nodes of an NLR immune receptor network - PMC](https://pmc.ncbi.nlm.nih.gov/articles/PMC8412950/)

[24] [AlphaFold 3 predictions for paired NLR immune receptors](https://zenodo.org/records/13826775)

[25] [AlphaFold 3 predictions for paired NLR proteins - Obsidian Vault - Obsidian v1.8.7](https://zenodo.org/records/15552925/files/AlphaFold%203%20predictions%20for%20paired%20NLR%20proteins.pdf?download=1)

[26] [Structural Insights into the Plant Immune Receptors PRRs and NLRs - PMC](https://pmc.ncbi.nlm.nih.gov/articles/PMC7140948/)

[27] [NRC Immune receptor networks show diversified hierarchical genetic architecture across plant lineages - PMC](https://pmc.ncbi.nlm.nih.gov/articles/PMC11371147/)

[28] [NLRscape: an atlas of plant NLR proteins - PMC](https://pmc.ncbi.nlm.nih.gov/articles/PMC9825502/)

[29] [P( all-atom ) Is Unlocking New Path For Protein Design](https://www.biorxiv.org/content/10.1101/2024.08.16.608235v4)

[30] [bioRxiv Channel: The Sainsbury Laboratory](https://connect.biorxiv.org/relate/feed/98)

[31] [Diversification of the “EDVID” packing motif underpins structural and functional variation in plant NLR coiled-coil domains](https://www.biorxiv.org/content/10.1101/2025.06.01.657260v1)

[32] [The N-terminal domains of NLR immune receptors exhibit structural and functional similarities across divergent plant lineages - PMC](https://pmc.ncbi.nlm.nih.gov/articles/PMC11218826/)

[33] [What AlphaFold 3 struggles with | AlphaFold](https://www.ebi.ac.uk/training/online/courses/alphafold/alphafold-3-and-alphafold-server/introducing-alphafold-3/what-alphafold-3-struggles-with/)

[34] [Resurrection of plant disease resistance proteins via helper NLR bioengineering](https://www.biorxiv.org/content/10.1101/2022.12.11.519957v2)

[35] [Unmasking the invaders: NLR-mal function in plant defense](https://www.ncbi.nlm.nih.gov/pmc/articles/PMC10698375/)

[36] [Oligomerised RIPK1 is the main core component of the CD95 necrosome](https://www.ncbi.nlm.nih.gov/pmc/articles/PMC12130296/)

[37] [A hierarchical immune receptor network in lettuce reveals contrasting patterns of evolution in sensor and helper NLRs | bioRxiv](https://www.biorxiv.org/content/10.1101/2025.02.25.639832v1.full-text)

[38] [A disease resistance protein triggers oligomerization of its NLR helper into a hexameric resistosome to mediate innate immunity](https://www.biorxiv.org/content/10.1101/2024.06.18.599586v1)

[39] [Fig. S6.Hexameric AlphaFold 3 predictions for previously reported NLR... | Download Scientific Diagram](https://www.researchgate.net/figure/Fig-S6Hexameric-AlphaFold-3-predictions-for-previously-reported-NLR-pairs-The-amino_fig7_387154533)

[40] [Accurate Prediction of Protein Complex Stoichiometry by Integrating AlphaFold3 and Template Information](https://www.ncbi.nlm.nih.gov/pmc/articles/PMC11838762/)

[41] [Frontiers | PlantNLRatlas: a comprehensive dataset of full- and partial-length NLR resistance genes across 100 chromosome-level plant genomes](https://www.frontiersin.org/journals/plant-science/articles/10.3389/fpls.2023.1178069/full)

[42] [Structural, Functional and Genomic Diversity of Plant NLR proteins: an Evolved Resource for Rational Engineering of Plant Immuni](https://ddd.uab.cat/pub/artpub/2018/200471/annrevphy_a2018v56p243.pdf)

[43] [AlphaFold3: An Overview of Applications and Performance Insights](https://www.mdpi.com/1422-0067/26/8/3671)

[44] [Subfunctionalization of NRC3 altered the genetic structure of the Nicotiana NRC network | PLOS Genetics](https://journals.plos.org/plosgenetics/article?id=10.1371/journal.pgen.1011402)

**Motivation:**

$\def\mathcal#1{\mathit{#1}}\def\mathscr#1{\mathit{#1}}$

The Structural Novelty Index (SNI) is a two-phase quantitative framework designed to identify unconventional helper NLRs by measuring geometric deviations from the canonical hexameric NRC resistosome. It utilizes AlphaFold 3 (AF3) to establish a computationally consistent "AF3-on-AF3" baseline, circumventing the noise and missing data (e.g., N-terminal $\alpha1$ helices) inherent in experimental PDB structures.

#### Motivation and Evidence

The SNI is motivated by the need to identify "structural dark matter"—proteins that have diverged beyond the reach of sequence-based tools (BLAST) but retain or evolve specific resistosome architectures.

- **Empirical Support:** The existence of biological outliers like NRCX (a negative regulator) and WAI3 (an octameric resistosome) confirms that the NRC family is not structurally monolithic.
- **Established Determinants:** The index parameters (swivel, latch, divergence angle) map directly to the molecular switch mechanism of NLR activation, such as the "180° rotation of the NB-HD1 module relative to the WHD-LRR module" and "specific 10° shift in the NB-ARC interdomain angle" that dictate stoichiometry.
- **AF3 Reliability:** Evidence confirms AF3 can model the NRC2 hexamer with high precision (RMSD ~0.937–1.72 Å), justifying the use of $S_{pref}$ (Stoichiometric Preference Score) as a computational assay.

#### Explanatory Power

The SNI offers a causal structural explanation for observations that sequence analysis cannot fully resolve:

- **Functional Failures:** "While sequence conservation is known, the hypothesis provides a structural explanation for *why* a sequence with a high HMM score (like NRCX) might still fail to trigger cell death (e.g., an inverted hydrophobicity gradient or low helical propensity due to specific polymorphisms)."
- **Effector Resilience:** "The SNI could explain why certain uncharacterized NLRs are resistant to specific effectors or exhibit autoactivity by identifying alternative swivel architectures that bypass the canonical hinge regulation."
- **Oligomeric State:** $S_{pref}$ "can explain the functional status of 'grey-area' NLRs (like NRCX or ancestral NRC0-S) by quantifying their preference for specific oligomeric states, which is not always clear from sequence alone."

#### Counterarguments and Risks

- **The Monomer Assumption:** Phase 1 (mSNI) assumes "readiness" for unconventional oligomerization is encoded in the monomer. There is a risk that "novel resistosome architectures only emerge during the massive conformational rearrangements of the activation process," leading to false negatives.
- **Static Weighting:** The index uses fixed weights ($w_i$) that do not currently incorporate pLDDT. Without dynamic weighting, the SNI "might interpret software modeling artifacts as biological novelty."
- **Omission of Dimers:** Since canonical NRCs exist as homodimers in their inactive state, excluding dimer interface metrics "may miss 'atypical' NLRs that possess novel autoinhibitory mechanisms or bypass the dimer state entirely."

#### Differentiating Features

The SNI is uniquely suited for this task because it transitions from descriptive structural biology to a predictive, high-throughput pipeline. Unlike metrics that focus on individual residue interactions (which are often "secondary effect of these primary residue interactions"), the SNI prioritizes the "shape-dynamics of the NB-ARC and CC domains." Its primary advantage is the **"AF3-on-AF3" strategy**, which ensures that the index measures biological deviation rather than software-to-experimental artifacts.

**Coherence:**

$\def\mathcal#1{\mathit{#1}}\def\mathscr#1{\mathit{#1}}$

No incoherence found.

**Deep verification:**

$\def\mathcal#1{\mathit{#1}}\def\mathscr#1{\mathit{#1}}$

Based on the provided assumptions and reasoning, the **Structural Novelty Index (SNI)** hypothesis may be incorrect due to several technical flaws in its structural modeling logic, biological misconceptions, and rigid data-handling parameters.

The primary reasons the hypothesis may fail are as follows:

#### 1. Technical Artifacts and Circular Reasoning in AF3

- **Symmetry Bias:** The use of the **Protomer Divergence Angle ($\Delta\theta_{int}$)** is likely invalid because AlphaFold 3 Multimer tends to force the symmetry of the input chains. A hexameric input will almost always produce a $60^\circ$ angle as a simulation artifact, meaning the metric measures the input setup rather than the protein’s actual biological stoichiometry.
- **Scoring Biases:** The **Stoichiometric Preference Score ($S_{pref}$)** is compromised by the inherent size-bias of ipTM scores, which may lead the framework to incorrectly over-predict higher-order oligomers regardless of their biological reality.

#### 2. Biological Misconceptions regarding NLR Activation

- **MADA Motif Behavior:** The framework assumes that high pLDDT scores ($>85$) and stable helices are indicators of a functional **MADA motif ($\alpha_{MADA}$)**. However, these motifs are often disordered in their inactive monomeric state and only transition to a helix upon activation. By requiring high confidence in the monomeric stage, the SNI would likely fail to identify true MADA motifs.
- **Oversimplified Targeting:** The **Funnel Hydrophobicity Gradient (FHG)** is criticized as a significant oversimplification. It fails to account for the complex signaling and specific targeting motifs required to guide NLRs to various organelles, assuming a simple hydropathy gradient is sufficient to determine functional shifts.

#### 3. Flawed Structural Metrics and Rigidity

- **Fixed Residue Ranges:** The **Latch Proximity ($D_{latch}$)** metric uses a fixed residue range (392-497) that does not account for insertions or deletions (indels) common in large, diverse datasets. This makes the metric incompatible with the evolutionary variation found in a 6,000-sequence library.
- **Arbitrary Thresholds:** The **$3\text{\AA}$ threshold** for latch proximity is considered too narrow to accommodate natural evolutionary diversity, potentially leading to a high rate of false positives for "novelty" where none exists.

#### 4. Framework Design and Screening Risks

- **False Negatives in the "Monomer-First" Funnel:** The hierarchical approach—filtering monomers (mSNI) before multimers (cSNI)—is fundamentally risky. It may exclude proteins that appear standard as monomers but possess novel interface residues that lead to unconventional multimeric structures (like filaments or octamers). This prevents the framework from identifying the very novelty it is designed to find.
