## Supplementary Data S6 for "AI-guided discovery of atypical protein assemblies": [EXTERNAL] ├░Y╠ê┬o┬1⁄4 Co-scientist - Hypothesis 187642863749828 - Defining a Quantitative 8-Parameter Structural Novelty Index (SNI) for Unconventional NRC-NLRs using AF3 Stoichiometric Sampling - For research purposes only - Google confidential.docx

### 🧬 Co-scientist - Defining a Quantitative 8-Parameter Structural Novelty Index (SNI) for Unconventional NRC-NLRs using AF3 Stoichiometric Sampling

**About**: *This is an experimental system for generating novel and testable hypotheses. The hypotheses are generated by a model and may be wrong. For research purposes only.*

### Proposal

$\def\mathcal#1{\mathit{#1}}\def\mathscr#1{\mathit{#1}}$

#### 1. Methodological Principles

The Structural Novelty Index (SNI) shifts from static geometry to weighted geometric fitness^25^, incorporating multi-stoichiometric sampling and confidence-weighted normalization to ensure that identified deviations represent genuine biological shifts.

- **Stoichiometric Flexibility:** Each sequence is modeled at $n=4, 6, 8$. This allows for the identification of sequences where "hexameric strain" is energetically unfavorable compared to a tetramer or octamer^15^.
- **Confidence Calibration:** Parameters are multiplied by local pLDDT scores^11^ to prevent the SNI from flagging flexible loops or poorly modeled regions as novel.
- **Functional Physics:** The index utilizes pore-walking algorithms ($E_{flux}$) and hydrophobic moment vectors ($\vec{M}_{align}$) to focus on the physics of ion conduction and membrane insertion.
- **Baseline Definition:** The SNI uses 9FP6 and 9CC8 as the active-state hexameric anchors and uses 9RI9 specifically to define the "resting-to-active" transition vector.

#### 2. Quantitative Parameters of the Structural Novelty Index (SNI)

The SNI is a composite metric ($SNI_{total} = \sum P_{n} \cdot w_{n}$) designed to quantify structural divergence. Each candidate is modeled in AlphaFold 3 across three states ($n=4, 6, 8$) to calculate stoichiometric fitness before parameters are extracted.

##### **Parameter 1: Stoichiometric Pitch Deviation ($\Delta \theta_{pitch}$)**

- **Unit:** Degrees ($^\circ$)
- **Description:** The deviation of the inter-protomer NB-ARC interface angle from the canonical $60^\circ$ (hexameric) baseline.
- **Measurement:** Calculated from the $n=6$ AF3 model. If the sequence demonstrates higher fitness at $n=4$ or $n=8$, the $n=6$ model will show significant angular strain at the interface.
- **Novelty Signal:** Values $> 5^\circ$ from the 9FP6/9CC8 baseline indicate "Stoichiometric Novelty"^6^ (suggesting potential tetrameric or octameric behavior).

##### **Parameter 2: pLDDT-Weighted Vertical Planarity ($\delta_{z, adj}$)**

- **Unit:** Angstroms ($\mathring{A}$)
- **Description:** The vertical "stair-stepping" between adjacent subunits, normalized by the interface confidence (ipTM).
- **Measurement:** $P = \text{StDev}(Z_{NB-ARC}) \times ipTM$.
- **Novelty Signal:** $\delta_{z, adj} > 4.0 \mathring{A}$ indicates a "lock-washer" or helical polymerization mechanism, whereas canonical NRCs (9FP6) are flat discs^19^ ($\delta_{z} < 1.0 \mathring{A}$).

##### **Parameter 3: NB-ARC Hinge Angle ($\alpha_{inter}$)**

- **Unit:** Degrees ($^\circ$)
- **Description:** The angle between the principal axes of the NB-ARC and the Winged Helix Domain (WHD)^20^.
- **Measurement:** Measures the internal "kink" of the domain switch rather than global RMSD.
- **Novelty Signal:** Canonical resistosomes (9CC8) show a specific activated hinge state. Deviation ($> 15^\circ$) suggests a non-canonical activation mechanism or a constitutively active sensor-like helper^4^.

##### **Parameter 4: Pore Constriction Electrostatics ($E_{flux}$)**

- **Unit:** mV or Dimensionless (Calculated via HOLE)
- **Description:** The electrostatic potential specifically at the narrowest 5 $\mathring{A}$ "throat" of the N-terminal $\alpha$1-helix bundle.
- **Measurement:** Uses a pore-walking algorithm to identify the constriction point.
- **Novelty Signal:** Canonical NRCs are cation channels (strongly electronegative at the throat). A neutral or positive $E_{flux}$ indicates an anion-selective or non-conductive scaffold NRC^12^.

##### **Parameter 5: MADA Vector Alignment ($\vec{M}_{align}$)**

- **Unit:** Degrees ($^\circ$)
- **Description:** The angle between the hydrophobic moment vector ($\mu H$) of the N-terminal $\alpha$1 helix and the central pore axis.
- **Measurement:** Calculated by projecting the hydrophobic side-chain centroids onto the radial plane.
- **Novelty Signal:** A vector pointing away from the pore center (angle $> 90^\circ$) suggests the MADA motif is sequestered or flipped, indicating the NLR does not form a traditional membrane pore.

##### **Parameter 6: Protomer Dihedral Twist ($\phi_{twist}$)**

- **Unit:** Degrees ($^\circ$)
- **Description:** The twist angle of the LRR domain relative to the NB-ARC signaling ring.
- **Measurement:** Defined by a vector from the ring centroid to the NB-ARC and a second vector to the LRR terminus.
- **Novelty Signal:** Deviations from the "straight-out" orientation of NRC2/4 indicate propensity for giant assemblies (10–14 mers) or interface occlusion.

##### **Parameter 7: Core N-terminal Radial Displacement ($\Delta R_{core}$)**

- **Unit:** Angstroms ($\mathring{A}$)
- **Description:** The distance from the pore axis to the center of the $\alpha$1 helix (residues 5-15).
- **Measurement:** Uses the geometric center of the MADA helix.
- **Novelty Signal:** A wider displacement ($> 15 \mathring{A}$) in a forced hexamer suggests the N-terminus is vestigial and cannot reach the center to form the conduction funnel seen in 9FP6.

##### **Parameter 8: Hydrophobic Packing Efficiency ($\rho_{pack}$)**

- **Unit:** Percentage (%)
- **Description:** The ratio of buried-to-total surface area in the $\alpha$1-helix bundle.
- **Measurement:** Calculated from the AF3 multimer interface.
- **Novelty Signal:** Values $< 40\%$ (with high ipTM) suggest an unstable or transient pore, potentially acting as a signal rather than a stable cytotoxic channel.

#### 3. Implementation Workflow

1. **Tier 1:** AF3 modeling at $n=6$^21^ with oleic acid for the ~6,000 NRC sequences^2^.
2. **Tier 2:** Extraction of the eight parameters. Weight all geometric results by local pLDDT and ipTM^11^.
3. **Tier 3:** High SNI candidates (Score > Threshold) are re-modeled at $n=4$ and $n=8$ to confirm stoichiometric fitness^22^.
4. **Tier 4:** Candidates deviating from 9FP6 (active) but resembling 9RI9 (resting) are flagged as inactivation-prone or decoy NRCs.

[4]  [A disease resistance protein triggers oligomerization of its NLR helper into a hexameric resistosome to mediate innate immunity - PubMed Central.](https://vertexaisearch.cloud.google.com/grounding-api-redirect/AUZIYQFVbR9sIrffCZKtK4p23yiub50KUsFhOHzgPE5KVpoQaVh3h9QwmOF5JRJS4FWqbNmd-_NJemd1YZMg-5Qg_RqBo9LJ9T0uckerLuwee_aBoR5q3Vr6qu9bJDeAayXI61ANPsVZd0TmoM1O260Z)

[5] Hung-Yu, Him, Kim-Teng, Foong-Jing, O., Chih-Hang.  [A hydrophobic core in the coiled-coil domain is essential for NRC resistosome function.](https://www.biorxiv.org/content/10.1101/2025.01.21.634219v2) Published 2025. [https://www.biorxiv.org/content/10.1101/2025.01.21.634219v2.](https://www.biorxiv.org/content/10.1101/2025.01.21.634219v2)

[6] Jogi, AmirAli, P., Andres, Jake, Jiorgos, et al.  [A disease resistance protein triggers oligomerization of its NLR helper into a hexameric resistosome to mediate innate immunity.](https://www.biorxiv.org/content/10.1101/2024.06.18.599586v1) Published 2024. [https://www.biorxiv.org/content/10.1101/2024.06.18.599586v1.](https://www.biorxiv.org/content/10.1101/2024.06.18.599586v1)

[7] Hung-Yu, Him, Kim-Teng, Foong-Jing, O., Chih-Hang.  [A hydrophobic core in the coiled-coil domain is essential for NRC resistosome function.](https://www.biorxiv.org/content/10.1101/2025.01.21.634219v3) Published 2025. [https://www.biorxiv.org/content/10.1101/2025.01.21.634219v3.](https://www.biorxiv.org/content/10.1101/2025.01.21.634219v3)

[8]  [The Pik NLR pair accumulates at the plasma membrane as a hetero-oligomeric sensor-helper immune protein complex prior to activat - bioRxiv.](https://vertexaisearch.cloud.google.com/grounding-api-redirect/AUZIYQE_XuJLf7G39KmWhU5bZLzPYRNBL93nC8oOxrYqy8q04M0iYOuQiVb8OEJ3NTml53gIiZsUB_MHxPOSAuMetoHMClJmJ3I4pOyFVoi1NN5FwDH8_gmqbpOKIQg8duZeb2pRvbBz51kvKDvAup_wFlrg-2DGT3tl5dZ4M1U5BMwkQih8QQr-SqG2yf6RZlLR)

[9] Jogi, AmirAli, P., Andres, Jake, Jiorgos, et al.  [A disease resistance protein triggers oligomerization of its NLR helper into a hexameric resistosome to mediate innate immunity.](https://www.biorxiv.org/content/10.1101/2024.06.18.599586v1) Published 2024. [https://www.biorxiv.org/content/10.1101/2024.06.18.599586v1.](https://www.biorxiv.org/content/10.1101/2024.06.18.599586v1)

[10] Hiroaki, Mauricio, Adeline, Chih-hang, Lida, Toshiyuki, et al.  [An N-terminal motif in NLR immune receptors is functionally conserved across distantly related plant species.](https://www.biorxiv.org/content/10.1101/693291v1) Published 2019. [https://www.biorxiv.org/content/10.1101/693291v1.](https://www.biorxiv.org/content/10.1101/693291v1)

[11] Baoquan, Kun, Zhenling, Alexey, Jianyi.  [Improved model building for cryo-EM maps using local attention and 3D rotary position embedding.](https://www.biorxiv.org/content/10.1101/2024.11.13.623164v4) Published 2024. [https://www.biorxiv.org/content/10.1101/2024.11.13.623164v4.](https://www.biorxiv.org/content/10.1101/2024.11.13.623164v4)

[12] Khong‐Sam, Philip.  [Taking the lead: NLR immune receptor N‐terminal domains execute plant immune responses.](https://www.ncbi.nlm.nih.gov/pmc/articles/PMC10952240/) Published 2023. [https://www.ncbi.nlm.nih.gov/pmc/articles/PMC10952240/.](https://www.ncbi.nlm.nih.gov/pmc/articles/PMC10952240/)

[13]  [A disease resistance protein triggers oligomerization of its NLR helper into a hexameric resistosome to mediate innate immunity - PubMed Central.](https://vertexaisearch.cloud.google.com/grounding-api-redirect/AUZIYQFXtKpVaWiA-fwG2XOPS8cOPgjg6H3ReanFTrVdRZbVrPLnqWn-L3S1gijGaI6qkCudHKUBOU6-YcDZh26Sc2LkfBPmSpTiCSoiX4-9Jl28FsE5OLAz3ZnXXi5cxJQGTmBSY34TSuz2c4pLXgi6)

[14] Hsuan, Toshiyuki, Andres, Raoul, Yu, P., et al.  [A hierarchical immune receptor network in lettuce reveals contrasting patterns of evolution in sensor and helper NLRs.](https://www.biorxiv.org/content/10.1101/2025.02.25.639832v1) Published 2025. [https://www.biorxiv.org/content/10.1101/2025.02.25.639832v1.](https://www.biorxiv.org/content/10.1101/2025.02.25.639832v1)

[15] Jogi, AmirAli, P., Andres, Jake, Jiorgos, et al.  [A disease resistance protein triggers oligomerization of its NLR helper into a hexameric resistosome to mediate innate immunity.](https://www.biorxiv.org/content/10.1101/2024.06.18.599586v1) Published 2024. [https://www.biorxiv.org/content/10.1101/2024.06.18.599586v1.](https://www.biorxiv.org/content/10.1101/2024.06.18.599586v1)

[16] Hsuan, Toshiyuki, Andres, Raoul, Yu, P., et al.  [A hierarchical immune receptor network in lettuce reveals contrasting patterns of evolution in sensor and helper NLRs.](https://www.biorxiv.org/content/10.1101/2025.02.25.639832v1) Published 2025. [https://www.biorxiv.org/content/10.1101/2025.02.25.639832v1.](https://www.biorxiv.org/content/10.1101/2025.02.25.639832v1)

[17]  [9FP6: Structure of the NbNRC2 hexameric resistosome - RCSB PDB.](https://vertexaisearch.cloud.google.com/grounding-api-redirect/AUZIYQEgmZwq4lzC-TNniLRmblz999h3HL8t2j9ROu93cC-r8gYYQaDs24oNdPc18MQDeBgnvlCyNZpfm4E5XRlLr-0qowRh7PfrjyYKxZd6O0w1YJLJi4-1GFGPmmmjBb0=)

[18] R., Nuren, P., William, N., N., et al.  [Pore structure controls stability and molecular flux in engineered protein cages.](https://www.biorxiv.org/content/10.1101/2021.01.27.428512v3) Published 2021. [https://www.biorxiv.org/content/10.1101/2021.01.27.428512v3.](https://www.biorxiv.org/content/10.1101/2021.01.27.428512v3)

[19] Jogi, AmirAli, P., Andres, Jake, Jiorgos, et al.  [A disease resistance protein triggers oligomerization of its NLR helper into a hexameric resistosome to mediate innate immunity.](https://www.biorxiv.org/content/10.1101/2024.06.18.599586v1) Published 2024. [https://www.biorxiv.org/content/10.1101/2024.06.18.599586v1.](https://www.biorxiv.org/content/10.1101/2024.06.18.599586v1)

[20]  [Resurrection of plant disease resistance proteins via helper NLR bioengineering - bioRxiv.](https://vertexaisearch.cloud.google.com/grounding-api-redirect/AUZIYQHyy7aBYWbV77XF2tZ1F7gN8Ty0-ymMKXhpZFWnPwkLT5t5fHrBQN_cJK5qyP57CZQEudW7wWeFLEQSguvjEY0whRoJFnK5g0E-J-izf01E2ZIhqKkqWD2G6SdhEtLAFYUTyoNBRvgGRcL0zBaeeLNJzlgv3VWeUNuidxEe1Ok=)

[21] Hsuan, Toshiyuki, Andres, Raoul, Yu, P., et al.  [A hierarchical immune receptor network in lettuce reveals contrasting patterns of evolution in sensor and helper NLRs.](https://www.biorxiv.org/content/10.1101/2025.02.25.639832v1) Published 2025. [https://www.biorxiv.org/content/10.1101/2025.02.25.639832v1.](https://www.biorxiv.org/content/10.1101/2025.02.25.639832v1)

[22]  [A disease resistance protein triggers oligomerization of its NLR helper into a hexameric resistosome to mediate innate immunity | bioRxiv.](https://vertexaisearch.cloud.google.com/grounding-api-redirect/AUZIYQGqShZFVSqDMSN4dF41LQp606jap1nZt23phiUarkiOGH6_ISaOKkl_zZV0uSBT9ZNL67aUZI7e4y-LVnQXpfC9mXIKkqi-6N-UaiPf5las3iiAvgUIBzxc_pVoDur37G2neSQN6lesnj5Ke3j9MUjA5MIqDRe0fbgU5Cm3A9TrUQ==)

[23] Yizhaq, Meytal.  [Atomic Structure of the LL-37(17-29) Human Antimicrobial Peptide Reveals Functional Helical Fibril.](https://www.biorxiv.org/content/10.1101/2020.02.04.933432v1) Published 2020. [https://www.biorxiv.org/content/10.1101/2020.02.04.933432v1.](https://www.biorxiv.org/content/10.1101/2020.02.04.933432v1)

[24]  [A disease resistance protein triggers oligomerization of its NLR helper into a hexameric resistosome to mediate innate immunity - PubMed Central.](https://vertexaisearch.cloud.google.com/grounding-api-redirect/AUZIYQElpdZYY_IVpJgGr3cXt3bBv-iG8fV7A_9aVUUStF4QWcZwnFBvN9nDuGaTGukYb0tRAfLYbZLH0O8LADU9Db2hRh9DLH-9aDcmMfpKgqC9ZOkfuuBigW666iDlmv5CvCcae2aCGLKkmgxZKX9D)

[25] Jogi, AmirAli, P., Andres, Jake, Jiorgos, et al.  [A disease resistance protein triggers oligomerization of its NLR helper into a hexameric resistosome to mediate innate immunity.](https://www.ncbi.nlm.nih.gov/pmc/articles/PMC11540030/) Published 2024. [https://www.ncbi.nlm.nih.gov/pmc/articles/PMC11540030/.](https://www.ncbi.nlm.nih.gov/pmc/articles/PMC11540030/)

### Reviews summary

$\def\mathcal#1{\mathit{#1}}\def\mathscr#1{\mathit{#1}}$

#### 1. Executive Verdict

The Structural Novelty Index (SNI) is a quantitative framework designed to identify unconventional NRC-NLRs by extracting 8 geometric and biophysical parameters from AlphaFold 3 (AF3) models across multiple stoichiometries. While the framework correctly identifies the need for confidence-weighted metrics and functional physics, it is built upon fundamental errors regarding structural baselines and methodological tool capabilities. **The hypothesis fails due to a critical misidentification of its primary structural anchor and the inclusion of mathematically invalid and technically impossible steps.**

#### 2. Critical Flaws

- **Structural Anchor Misidentification:** The hypothesis identifies PDB ID 9RI9 as a "resting-state anchor." This is an incorrect identification of the ground truth; PDB 9RI9 is the **active hexameric resistosome** of SlNRC3. Using an active state as the baseline for a "resting-to-active" transition vector invalidates the primary reference set for the index.
- **Methodological Mismatch (HOLE Software):** Parameter 4 ($E_{flux}$) relies on the HOLE program to calculate electrostatic potential in mV. HOLE is strictly a geometric radius profiler and lacks the physics engine to calculate electrostatics, which requires Poisson-Boltzmann solvers like APBS.
- **Incorrect Physical Baselines:** The hypothesis assumes a 5 Å "throat" constriction as a baseline for canonical NRC2/4 pores. Experimental data confirms that the actual physical diameter of these pores is approximately 17–19 Å, rendering the 5 Å threshold a false premise.
- **Mathematical Invalidity:** The composite formula ($SNI_{total} = \sum P_{n} \cdot w_{n}$) is mathematically invalid as it attempts to sum raw values with incommensurate units (e.g., adding degrees to angstroms). This introduces massive scale bias and makes the total score meaningless without normalization (e.g., Z-scores).
- **Computational Infeasibility:** The Tier 1 modeling of 6,000 hexamers (~900 residues per subunit) results in complexes of ~5,400 residues. This exceeds the AlphaFold 3 token limit of 5,120, making the primary high-throughput screening step impossible without significant domain truncation.

#### 3. Addressed Objections

The concern regarding **Modeling Noise vs. Thresholds** was evaluated. Critics suggested that a $5^\circ$ Pitch Deviation threshold might be lost in the stochastic noise of AF3 outputs. However, the hypothesis addresses this through **Confidence Calibration**, where geometric deviations are multiplied by local pLDDT and ipTM scores. This ensures that "novelty" is only flagged when the model demonstrates high confidence, effectively filtering out noise from flexible or poorly modeled regions. Similarly, the **Stoichiometric Sampling** approach ($n=4, 6, 8$) was validated by recent literature (e.g., the PreStoi method), which confirms that comparing confidence scores across different stoichiometry candidates is a robust way to determine the most biologically relevant assembly.

#### 4. Validated Risks & Limitations

- **Omission of Pentamers:** The framework models sequences at $n=4, 6, 8$ but ignores $n=5$. Given that pentamers are a known biological state for singleton NLRs (like ZAR1), this omission limits the index's ability to capture the full spectrum of stoichiometric evolution.
- **Universal Ligand Assumption:** The framework assumes oleic acid is the universal cofactor for all 6,000 NRCs. Forcing NRCs that require different lipids into an oleic acid-bound model may create artificial "structural strain," leading to false-positive novelty signals.
- **Lack of Statistical Calibration:** The "Novelty Signal" thresholds (e.g., $\delta_{z, adj} > 4.0 \mathring{A}$) lack calibration against a null distribution of the 350-sequence canonical set, making them arbitrary and prone to high false-positive rates.
- **Missing Residue Baselines:** Parameters focusing on the N-terminal $\alpha$1-helix (Parameters 4, 5, 7, and 8) rely on AF3-generated models of the reference set because these residues are disordered/missing in experimental PDB structures (9FP6, 9CC8). This forces the index to use a prediction as its "gold-standard" baseline.

#### 5. Supporting Arguments & Evidence (Motivation)

- **Theoretical Basis:** The shift from static geometry to functional physics (e.g., **MADA Vector Alignment** and **Pore Electrostatics**) is scientifically sound. It allows the system to distinguish between "conductive helpers" and "non-conductive scaffolds" that might appear identical under simple RMSD or sequence identity checks.
- **Empirical Support:** The use of **Hydrophobic Moment Vectors** ($\vec{M}_{align}$) is supported by established structural biology (Eisenberg moments) and aligns with findings that the orientation of the MADA motif is the primary "death switch" in these receptors.
- **Comparative Advantage:** This methodology transitions from "one protein at a time" structural biology to a systems-level approach. By integrating confidence weighting with stoichiometric fitness, it offers a more nuanced triage than current methods that rely solely on global ipTM scores.

#### 6. Alignment & Novelty

- **Alignment:** The hypothesis is highly aligned with the research goal of establishing a quantitative index (5–10 parameters) to distinguish unconventional NRCs from canonical hexamers.
- **Novelty:** The index is moderately novel. While stoichiometric sampling and inter-domain hinge analysis are established techniques, the formalization of **Vertical Planarity** ($\delta_{z, adj}$) and **Pore Constriction Electrostatics** into a unified high-throughput triage tool for the NRC family represents a novel methodological synthesis.

#### 7. Feasibility Assessment (Go/No-Go Decision)

- **Resource Intensity:** Very High. Requires significant GPU time for AF3 modeling and high-end hardware (80GB VRAM) to handle large complexes.
- **Technical Complexity:** High. Requires the integration of multiple specialized tools (HOLE, APBS) and the development of custom scripts for PCA-based geometric extraction.
- **Time to Verdict:** 2–3 months for a full-scale screen.
- **Go/No-Go Decision: No-Go.** The hypothesis cannot proceed as written due to the fundamental structural error (using active 9RI9 as a resting anchor) and the mathematical invalidity of the $SNI_{total}$ formula.

#### 8. Conclusion

The Structural Novelty Index (SNI) is a technically sophisticated proposal that correctly identifies the structural "fault lines" of the NRC family. However, it is compromised by fatal flaws: it misidentifies its resting-state reference structure, utilizes software for physics calculations it cannot perform, and employs a mathematically unsound scoring method. While the individual parameters (such as MADA alignment and vertical planarity) are valuable, the framework as a whole requires a complete revision of its baselines and statistical integration before it can be considered a viable tool for NRC discovery.

#### Research contacts

$\def\mathcal#1{\mathit{#1}}\def\mathscr#1{\mathit{#1}}$

Based on the provided research articles, here are the specific researchers who are best suited to review your Structural Novelty Index (SNI) hypothesis, listed in order of those who studied the most specific aspects of the idea first.

#### 1. Jogi Madhuprakash

**Justification:** Madhuprakash is the lead researcher on the core structural studies of NRC hexamers. He provided the exact quantitative comparisons you require, such as the $10^\circ$ difference in the interdomain angle between NRC2 and ZAR1 (85° vs 75°) and the measurement of pore diameters. **Supporting Evidence:** In the article *"A disease resistance protein triggers oligomerization of its NLR helper into a hexameric resistosome to mediate innate immunity"* (Abstract 3, 5, 7), he explicitly defines the structural parameters of the **9FP6** (NbNRC2) resistosome. His work involves measuring "angular distances between domains" and "outward displacement of the NB domains," which are the foundation of your Parameter 1 ($\Delta \theta_{pitch}$) and Parameter 7 ($\Delta R_{core}$).

#### 2. AmirAli Toghani

**Justification:** Toghani specializes in using AlphaFold 3 (AF3) to distinguish between NRC-Helpers (NRC-H) and NRC-Sensors (NRC-S) based on their ability to form hexameric oligomers. **Supporting Evidence:** In the article *"A hierarchical immune receptor network in lettuce reveals contrasting patterns of evolution in sensor and helper NLRs"* (Abstract 6), Toghani utilized AF3 to model ~6,000 sequences and analyzed metrics like **pTM, ipTM, and pLDDT** to separate canonical helpers from unconventional sensors. This directly aligns with your implementation workflow and your "Confidence Calibration" strategy.

#### 3. Sophien Kamoun

**Justification:** As a senior author on the majority of the provided NRC articles, Kamoun provides the evolutionary and functional context for the NRC network. He oversaw the transition from the MADA motif discovery to the structural determination of the hexamer. **Supporting Evidence:** His work spans the identification of the **MADA motif** (Abstract 1) to the current AF3 screening of Solanaceae species (Abstract 6, 9). He is an expert on the "activation-and-release" model and the "death switch" mechanism, which is critical for reviewing your "Functional Physics" parameters like $E_{flux}$ and $\vec{M}_{align}$.

#### 4. Hiroaki Adachi

**Justification:** Adachi is the primary expert on the N-terminal CC-domain variations and the conservation of the MADA motif. **Supporting Evidence:** In *"An N-terminal motif in NLR immune receptors is functionally conserved across distantly related plant species"* (Abstract 1, 4), he defined the MADA consensus sequence ($MADAxVSFxVxKLxxLLxxEx$). His expertise is essential for reviewing **Parameter 5 (MADA Vector Alignment)** and **Parameter 8 (Hydrophobic Packing)**, as he identified the minimal 29-amino-acid region required for cell death.

#### 5. Michael W. Webster

**Justification:** Webster is a lead structural biologist who conducted the Cryo-EM analysis of the NbNRC2 resistosome. **Supporting Evidence:** He co-authored the structural studies in Abstract 3 and 9. He is responsible for resolving the interprotomer interfaces and identifying why NRCs accommodate an additional protomer compared to pentameric NLRs (such as the "kink" in the $\alpha$4 helix at L126). He would be the ideal reviewer for **Parameter 2 (Vertical Planarity)** and **Parameter 6 (Protomer Dihedral Twist)**.

#### 6. Mauricio P. Contreras

**Justification:** Contreras has been involved in both the biochemical characterization of NRCs and the modern AF3-based modeling of their resistosomes. **Supporting Evidence:** He is a consistent contributor to the research on the NRC network (Abstract 1, 3, 6, 9). His recent work focuses on distinguishing sensors and helpers via AI modeling, making him a perfect fit to review the "Stoichiometric Flexibility" and "Tiered Implementation Workflow" of your SNI.

#### 7. Chih-hang Wu

**Justification:** Wu was instrumental in defining the NRC superclade and the original distinction between helpers and sensors in the Solanaceae dataset. **Supporting Evidence:** Mentioned in Abstract 1 and 4, his foundational work on the NRC network provides the "Baseline Definition" for your study. He can provide review on the "Scope" of your index regarding the ~350 Solanaceae species.

### Appendix:

**All reviews:**

**Correctness:**

$\def\mathcal#1{\mathit{#1}}\def\mathscr#1{\mathit{#1}}$

The following review evaluates the Structural Novelty Index (SNI) proposed for identifying unconventional NRC-NLRs.

#### 1. Related Article Abstracts

1. **[3, 7, 9, 11, 13, 14] A disease resistance protein triggers oligomerization of its NLR helper into a hexameric resistosome...**: Establishes the ground truth structure of activated NbNRC2 (9FP6) as a homohexamer and compares it to pentameric singleton resistosomes (ZAR1).
2. **[1, 4] An N-terminal motif in NLR immune receptors is functionally conserved...**: Defines the MADA motif (residues 1–21) essential for pore formation and cell death, serving as the basis for Parameters 4, 5, 7, and 8.
3. **[6, 10] A hierarchical immune receptor network in lettuce reveals contrasting patterns of evolution...**: Demonstrates that AF3 can differentiate between helper (NRC-H) and sensor (NRC-S) NLRs by their ability to form hexameric resistosomes in silico.
4. **[12] A hydrophobic core in the coiled-coil domain is essential for NRC resistosome function**: Identifies residues in the $\alpha2$ to $\alpha4$ helices critical for oligomerization, supporting the structural focus of the SNI.
5. **[15] Accurate Prediction of Protein Complex Stoichiometry by Integrating AlphaFold3...**: Validates the use of AF3 ranking scores (ipTM/pTM) to predict stoichiometry, supporting the multi-stoichiometric sampling approach.
6. **[16] Sensor NLR immune proteins activate oligomerization of their NRC helper**: Establishes the "activation-and-release" model and the resting-state dimer structure (9RI9).
7. **[8] Confidence scores in AlphaFold-Multimer**: Explains the relevance of ipTM and pTM in evaluating complex interfaces, justifying their use as weighting factors in the SNI.

#### 2. Detailed Assumptions

1. **Stoichiometric Frustration:** The sequence of a non-hexameric NLR will exhibit measurable geometric "strain" (angular pitch deviation) when forced into a hexameric model by AlphaFold 3 (AF3).
2. **Model Validity for Disordered Regions:** AF3 can generate high-confidence models of the N-terminal $\alpha1$ helix (MADA motif) that serve as a reliable baseline for pore physics, even though these residues are missing in experimental PDB structures (9FP6, 9CC8).
3. **Confidence as a Filter:** Local pLDDT and global ipTM scores are sufficient to distinguish genuine structural novelty from stochastic modeling artifacts or flexibility.
4. **Functional Redundancy of 9RI9:** The resting dimer (9RI9) provides a valid vector for the transition to activity that is applicable across the entire 6,000-sequence NRC family.
5. **Computational Scalability:** It is feasible to model 6,000 large multimeric complexes (~5,400 residues for a hexamer) using AF3 resources.

#### 3. Comparison with Knowledge Base

- **Ground Truth Consistency:** The idea correctly identifies 9FP6/9CC8 as active hexamers and 9RI9 as the resting state. This aligns with the PDB results in the knowledge base and Abstract 3.
- **The "Missing Residue" Problem:** The knowledge base warns that residues 1-17 are missing in the ground truth PDBs. The idea addresses this by basing its pore parameters ($E_{flux}$, $\vec{M}_{align}$, etc.) on AF3-generated models of the reference set (NRC2/3/4) rather than the incomplete experimental coordinates.
- **Stoichiometric Benchmarks:** Parameter 1 correctly uses $60^\circ$ as the baseline for hexamers, which is confirmed by the PDB summary ($60.00^\circ$ pitch for 9FP6).
- **Planarity:** Parameter 2 uses $\delta_z < 1.0 \text{ \AA}$ as the "flat disc" baseline, which matches the PDB summary measurement ($\approx 0.01 \text{ \AA}$).

#### 4. Reasoning about Correctness

- **Assumption 1 (Strain):** True. Abstract 6/10 provides evidence that sensors (which do not form resistosomes) show low confidence and high predicted aligned error when modeled as hexamers. Measuring "strain" via pitch deviation is a robust quantitative metric for this phenomenon.
- **Assumption 2 (AF3/MADA):** True. Abstract 3 explicitly states that AF3 allows for high-confidence modeling of the $\alpha1$ helices, a region historically difficult to resolve via cryo-EM.
- **Assumption 3 (Confidence):** True. This is standard practice in structural bioinformatics to avoid false positives in disordered loops.
- **Assumption 4 (Transition Vector):** Likely true. The NB-ARC rearrangement (Abstract 3) is a conserved mechanism across STAND proteins.
- **Assumption 5 (Scalability):** Questionable. Modeling 6,000 hexamers ($n=6$) exceeds the 5,000-residue token limit of some AF3 implementations (Abstract 15) and would require immense GPU time. However, using domain truncations (CC-NB-ARC) makes this feasible.

#### 5. Strength of Evidence

- **Direct Evidence:** Abstract 3 provides the direct structural coordinates for the canonical hexamer (9FP6) and confirms the rearrangement of the NB-ARC domains. Abstract 6/10 provides direct evidence that AF3 metrics (pTM/ipTM) can distinguish helpers from sensors in the NRC network.
- **Indirect Evidence:** The success of the MADA-motif swap (Abstract 1) between ZAR1 (pentamer) and NRC4 (hexamer) suggests that the pore-forming physics (Parameters 4, 5, 7) are the most conserved and functionally relevant aspects of structural biology in this family.

#### 6. Suggested Improvements

1. **Domain Truncation for High-Throughput:** To ensure the 6,000-sequence screen stays within AF3 token limits and reduces computational cost, use the **CC-NB-ARC** truncation (~450-500 residues per protomer) for Tier 1 and 2.
2. **NB-ARC Conservation Parameter:** The goal requires a parameter for residue conservation patterns. Add a **Parameter 9: Sequence-Structural Entropy ($S_{NB-ARC}$)**, which maps Shannon's entropy (from Abstract 10) onto the NB-ARC hinge to distinguish conserved helpers from highly variable sensors.
3. **Electrostatic Sensitivity:** Parameter 4 ($E_{flux}$) is highly sensitive to side-chain orientation. This should be cross-referenced with the **MADA hydrophobic core** (Abstract 12) to ensure the constriction is supported by stable packing.

#### 7. Goal Requirement Assessment

- **5-10 quantitative parameters:** Met (8 parameters defined).
- **Measurable from PDB/AF3:** Met.
- **Distinguish unconventional from canonical:** Met (via stoichiometric contrast and pitch deviation).
- **Incorporate 9FP6, 9RI9, 9CC8:** Met.
- **Focus on interface angles/distances:** Met (Parameters 1, 2, 3, 6, 7).
- **N-terminal CC variations:** Met (Parameters 4, 5, 7, 8).
- **NB-ARC conservation patterns:** Partially Met (hinge angles are used, but explicit sequence entropy is missing from the 8-parameter list).
- **High-throughput design:** Met (tiered approach).

#### 8. Reasoning about Correctness & Recommendation

The idea is highly sophisticated and scientifically robust. It correctly bypasses the limitations of experimental PDB structures (missing N-termini) by using AF3-modeled baselines for the canonical set. The inclusion of stoichiometric contrast ($n=4, 6, 8$) is an essential guard against "forced folding" artifacts in AI modeling. The parameters cover all major domains (CC, NB-ARC, LRR) and move beyond simple RMSD to focus on functional physics (ion flux and hydrophobic moment). While Tier 1 is computationally expensive, the tiered strategy is standard for this scale.

The idea respects the "Missing Residue" problem highlighted in the knowledge base and correctly identifies the ground truth structures. It is highly recommended for testing.

Answer: 9

**Novelty:**

$\def\mathcal#1{\mathit{#1}}\def\mathscr#1{\mathit{#1}}$

This review evaluates the novelty of the **Structural Novelty Index (SNI)**, a proposed quantitative framework to identify unconventional NRC-NLRs using 5–10 parameters derived from AlphaFold 3 (AF3) and PDB data.

#### 1. Related Article Abstract Titles

1. **[3] A disease resistance protein triggers oligomerization of its NLR helper into a hexameric resistosome...** (Confirms the hexameric ground truth for NRC2/4 and establishes inter-domain orientations).
2. **[11] A protomer-level structural comparison between the NbNRC2 hexamer and dimer...** (Directly explores inter-domain angles like the NB-ARC/WHD hinge and stoichiometry consequences).
3. **[15] Accurate Prediction of Protein Complex Stoichiometry by Integrating AlphaFold3...** (Introduces "PreStoi," which samples and ranks stoichiometries like $n=2$ to $n=9$ using AF3 ranking scores).
4. **[6] A hierarchical immune receptor network in lettuce reveals contrasting patterns of evolution...** (Estantiates the use of AF3 metrics like ipTM and pTM to distinguish sensors from helpers).
5. **[1] An N-terminal motif in NLR immune receptors is functionally conserved...** (Establishes the MADA motif as the structural executioner).
6. **[12] A hydrophobic core in the coiled-coil domain is essential for NRC resistosome function** (Identifies hydrophobic residue clusters in the CC domain relevant to SNI Parameter 8).
7. **[44] Structure of the activated Roq1 resistosome...** (Compares activated NLRs across increasing oligomeric states, identifying specific N-terminal linker conformations).
8. **[17] A helper NLR targets organellar membranes to trigger immunity** (Uses AF3 to compare CC-domain funnel lengths across different NLR clades).
9. **[48] Bridging prediction and reality: Comprehensive analysis of experimental and AlphaFold 2... structures** (Uses distances and angles between centers of mass [COM] to quantify domain organization, similar to the SNI methodology).
10. **[8] Confidence scores in AlphaFold-Multimer | AlphaFold** (Explains the use of ipTM/pTM for interface accuracy, the basis for SNI weighting).

#### 2. Aspects Already Explored

- **Stoichiometric Sampling ($n=4, 6, 8$):** The idea of modeling sequences across multiple stoichiometric states to find the most "fit" configuration is the core of the **PreStoi** method described in [15]. Abstract [3] and [11] also specifically compare hexameric NRCs to pentameric ZAR1/Sr35 to find the structural basis for stoichiometric shifts.
- **Confidence Calibration (pLDDT/ipTM weighting):** Weighting structural measurements by AF3 confidence metrics is standard practice [8]. Abstract [6] and [21] explicitly use ipTM and per-chain pTM to differentiate functional NRC-H from "degenerated" NRC-S.
- **Inter-domain Hinge Angles:** Parameter 3 ($\alpha_{inter}$) is heavily explored in [11] and [24], where the "kink" in the α4 helix and the $10^\circ$ widening of the NB-ARC relative to pentamers are identified as the drivers of NRC hexamerization.
- **Radial Displacement and Pore Width:** Abstract [11] and [13] already quantify the CC and NB pore diameters (e.g., $19\mathring{A}$ for NRC2 vs $12\mathring{A}$ for ZAR1), which directly relates to SNI Parameter 7.
- **Active-to-Resting Transition:** Using 9RI9 (dimer) and 9FP6 (hexamer) to define the "resting-to-active" transition vector is the specific focus of Abstract [11].

#### 3. Novel Aspects

- **Composite Index Formulation ($SNI_{total} = \sum P_{n} \cdot w_{n}$):** While the field currently uses individual metrics (like ipTM or RMSD) to evaluate models, the formalization of a weighted, multi-parameter index specifically for the triage of the 6,000-sequence NRC family is a novel methodological synthesis.
- **Physics-Based Functional Parameters (Parameters 4 & 5):** The introduction of **Pore Constriction Electrostatics ($E_{flux}$)** using pore-walking algorithms (HOLE) and **MADA Vector Alignment ($\vec{M}_{align}$)** via hydrophobic moment vectors moves the field from static geometry to functional channel physics. This allows for identifying "non-conductive scaffolds" or "anion-selective" NRCs that might look like canonical helpers but function differently.
- **Vertical Planarity/Lock-Washer Detection (Parameter 2):** Quantifying the vertical "stair-stepping" ($\delta_{z, adj}$) to distinguish flat disc resistosomes from helical "lock-washer" polymerization mechanisms is a specific structural descriptor not currently used in NLR high-throughput screening.
- **Protomer Dihedral Twist (Parameter 6):** Measuring the LRR terminus orientation relative to the signaled ring to predict "giant assemblies" (10–14 mers) is a novel structural triage metric.

#### 4. Novelty Review

The core idea is **moderately novel**.

The methodological framework relies heavily on "PreStoi" logic [15] (stoichiometric sampling) and the comparative structural biology already performed by the ground truth authors [11] (inter-domain angles). However, the idea adds significant value by integrating **biophysical descriptors** ($E_{flux}$, $M_{align}$) that go beyond the "interface confidence" currently used to triage NLRs in the provided literature. The SNI shifts the focus from "Is this a hexamer?" to "Does this hexamer have the physics of a cation channel?". This is a sophisticated refinement of current triage methods.

#### 5. Reasoning and Recommendation

The SNI is highly quantitative and more nuancied than a simple RMSD check. While an RMSD check might show that two proteins are different, the SNI provides a functional rationale (e.g., "MADA motif is flipped" or "pore is positive") for *why* they are unconventional. This is particularly useful for the NRC superclade, where ~80% of members lack the MADA motif or have "degenerated" [1, 24], yet some might represent functional novelties rather than mere loss-of-function.

**Recommendation:** The idea should be tested. The specific parameters (especially vertical planarity and electrostatic flux) are likely to uncover NRC "dark matter" (atypical scaffolds) that current ipTM-only screens would miss.

**Answer: 5**

**Feasibility:**

$\def\mathcal#1{\mathit{#1}}\def\mathscr#1{\mathit{#1}}$

#### Related Articles

1. **[3] A disease resistance protein triggers oligomerization of its NLR helper into a hexameric resistosome to mediate innate immunity** (Madhuprakash et al., 2024): Provides the primary ground truth for the active hexameric state (PDB 9FP6), confirming NbNRC2 assembles into a star-shaped hexamer rather than a pentamer.
2. **[16] Sensor NLR immune proteins activate oligomerization of their NRC helper** (Contreras et al., 2022): Describes the "activation-and-release" model and identifies the resting state of NbNRC2 as a homodimer (PDB 9RI9/8RFH), which is essential for the "resting-to-active" transition vector parameter.
3. **[6] A hierarchical immune receptor network in lettuce reveals contrasting patterns of evolution in sensor and helper NLRs** (Pai et al., 2025): Demonstrates that AlphaFold 3 (AF3) can distinguish between NRC helpers and sensors by successfully modeling hexameric resistosomes for helpers but not for sensors, validating the feasibility of using AF3 for this index.
4. **[15] Accurate Prediction of Protein Complex Stoichiometry by Integrating AlphaFold3 and Template Information** (Liu et al., 2025): Validates the methodological principle of using AF3 ranking scores (pTM/ipTM) across different stoichiometry candidates to determine the biologically correct assembly.
5. **[12] A hydrophobic core in the coiled-coil domain is essential for NRC resistosome function** (Wang et al., 2025): Identifies specific hydrophobic residues in the NRC CC-domain core, directly supporting the "Hydrophobic Packing Efficiency" parameter ($\rho_{pack}$).
6. **[1] An N-terminal motif in NLR immune receptors is functionally conserved across distantly related plant species** (Adachi et al., 2019): Defines the MADA motif and its role in cell death, providing the basis for the "MADA Vector Alignment" and "Radial Displacement" parameters.
7. **[8] Confidence scores in AlphaFold-Multimer** (EMBL-EBI, 2024): Defines the usage of ipTM and pLDDT scores, which the idea uses to weight geometric parameters and filter modeling noise.
8. **[5] A disease resistance protein triggers oligomerization of its NLR helper into a hexameric resistosome to mediate innate immunity** (Madhuprakash et al., 2024): Specifically compares protomer angles between hexameric NRC2 ($85^\circ$) and pentameric ZAR1 ($75^\circ$), supporting the "Stoichiometric Pitch Deviation" parameter.
9. **[13] A disease resistance protein triggers oligomerization of its NLR helper into a hexameric resistosome to mediate innate immunity** (Madhuprakash et al., 2024): Discusses the flexibility of the region connecting CC and NB-ARC, highlighting the necessity of confidence calibration (pLDDT weighting) in structural metrics.
10. **[11] A disease resistance protein triggers oligomerization of its NLR helper into a hexameric resistosome to mediate innate immunity** (Madhuprakash et al., 2024): Details the "kink" in the $\alpha$4 helix of the CC domain (residue L126) which contributes to stoichiometry, supporting the "Protomer Dihedral Twist" parameter.
11. **[2] Protein complexes in cells by AI‐assisted structural proteomics** (O'Reilly et al., 2023): Demonstrates the scalability of combining crosslinking MS/CoFrac-MS with AlphaFold-Multimer to model interactomes, proving high-throughput AI structural screening is viable.
12. **[10] A hierarchical immune receptor network in lettuce...** (Pai et al., 2025): Provides context for the Solanaceae dataset and highlights the contrast between conserved helpers and diversifying sensors.

#### Steps to Test the Idea

1. **Go/No-go Initial Experiment:** Benchmark the canonical NRCs (NRC2, NRC3, NRC4) by modeling them in AF3 at $n=4, 5, 6, 7, 8$. If the highest confidence scores (ipTM) and lowest geometric strain (Parameter 1) consistently peak at $n=6$ for all three, and clearly distinguish them from the pentameric ZAR1/Sr35 structures, proceed.
2. **Dataset Preparation:** Compile the ~6,000 NRC sequences from the ~350 Solanaceae species dataset.
3. **High-Throughput Modeling (Tier 1):** Perform AF3 multimer modeling at $n=6$ for the full dataset. *Note: Due to the token limit (~5,000 residues), sequences may need truncation to the CC-NB-ARC-WHD domains (~500 residues each) to fit 6 protomers into a single simulation.*
4. **Automated Parameter Extraction (Tier 2):** Develop Python scripts (using Biopython and MDTraj) to extract the eight quantitative parameters (Pitch, Planarity, Hinge, etc.) from the PDB/CIF outputs.
5. **Functional Physics Integration:** Integrate the HOLE algorithm to calculate the electrostatic potential and conduction "throat" characteristics ($E_{flux}$) for each model.
6. **SNI Scoring:** Calculate the composite SNI score for each sequence. Map candidates against the 9FP6/9CC8 (active) and 9RI9 (resting) anchors to calculate "deviation Z-scores."
7. **Multi-Stoichiometric Validation (Tier 3):** Re-model the top 5% highest SNI candidates at $n=4$ and $n=8$ to confirm if their structural "novelty" corresponds to a genuine preference for non-hexameric states.

#### Feasibility Reasoning

Testing this idea is **Resource Intensive**. The primary bottleneck is the computational demand of modeling 6,000 sequences as hexamers. At an average length of 900 residues, a 6-protomer complex (5,400 residues) exceeds the standard AlphaFold 3 web server token limit (5,000) and requires a local installation with high-end GPUs (e.g., A100 80GB) or a significant truncation strategy.

The proposed parameters are highly quantitative and technically sound. Metrics like **Stoichiometric Pitch Deviation** and **Vertical Planarity** are standard geometric calculations that can be automated via scripts in approximately one month. Integrating specialized tools like the **HOLE algorithm** for Parameter 4 adds a layer of complexity but is standard in structural biology.

The idea addresses "AI modeling bias" by utilizing multi-stoichiometric contrast (modeling at $n=4, 6, 8$), which ensures that deviations are not just artifacts of "forced folding" by AF3. The grounding in PDB anchors (9FP6, 9RI9) is factually accurate according to the provided literature. However, the requirement for full-scale multimeric modeling across 6,000 sequences places this in the 2–3 month timeline with high compute costs.

Answer: 5

**Impact potential:**

$\def\mathcal#1{\mathit{#1}}\def\mathscr#1{\mathit{#1}}$

#### Related Article Abstracts

1. **[3, 5, 7, 9, 11, 13, 14] A disease resistance protein triggers oligomerization of its NLR helper into a hexameric resistosome... (Madhuprakash et al., 2024):** This is the foundational ground truth for the "canonical" NRC hexamer. It establishes the hexameric stoichiometry of NbNRC2 (9FP6), the transition from dimer (9RI9) to hexamer, and the utility of AlphaFold 3 (AF3) in modeling the N-terminal funnels with lipids.
2. **[1, 4] An N-terminal motif in NLR immune receptors is functionally conserved... (Adachi et al., 2019):** Defines the MADA motif and its "death switch" role. Essential for parameters 4, 5, and 7, which focus on the $\alpha$1-helix bundle and pore physics.
3. **[6, 10] A hierarchical immune receptor network in lettuce... (Pai et al., 2025):** Demonstrates that AF3 can differentiate between helpers (pore-formers) and sensors (signaling decoys) by their ability to form stable oligomers. Validates the idea of using AF3 metrics to triage functional types.
4. **[15] Accurate Prediction of Protein Complex Stoichiometry by Integrating AlphaFold3... (Liu et al., 2025):** Confirms that AlphaFold 3 ranking scores (pTM/ipTM) across different stoichiometries can accurately predict the true oligomeric state. This supports the "Stoichiometric Flexibility" principle of the SNI.
5. **[12] A hydrophobic core in the coiled-coil domain is essential for NRC resistosome function (Wang et al., 2025):** Identifies specific hydrophobic residues in the CC domain ($\alpha$2 to $\alpha$4) that stabilize the resistosome. This provides a baseline for the "Hydrophobic Packing Efficiency" parameter (Parameter 8).
6. **[16] Sensor NLR immune proteins activate oligomerization of their NRC helper... (Contreras et al., 2022):** Establishes the activation-and-release model and the membrane-associated puncta behavior of NRCs, justifying the focus on pore physics and membrane insertion.
7. **[2] Protein complexes in cells by AI‐assisted structural proteomics (O'Reilly et al., 2023):** Validates the use of ipTM as a benchmark for high-quality structural models in complex assemblies, reinforcing the "Confidence Calibration" principle.
8. **[8] Confidence scores in AlphaFold-Multimer (EMBL-EBI):** Provides the technical definitions for pTM and ipTM used to weight the SNI parameters.

#### Assumptions for Impact Potential

1. **Computational Scalability:** The idea assumes that modeling 6,000 sequences as hexamers in AF3 is feasible. Given that AF3 models of large complexes (~5,000-6,000 residues for an NRC hexamer) are computationally expensive, this assumes access to significant GPU resources.
2. **Discriminatory Power of Geometric Strain:** It assumes that AF3's "forced folding" (attempting to fit a sequence into a hexamer even if it prefers a tetramer) will manifest as measurable geometric strain ($\Delta \theta_{pitch}$ and $\delta_{z, adj}$) rather than just a global drop in confidence.
3. **Accuracy of N-terminal Modeling:** It assumes AF3 with oleic acid correctly captures the hydrophobic moment and pore electrostatics of the MADA motif, which is often disordered in cryo-EM structures [3].
4. **Species Invariance:** It assumes that the SNI parameters defined for Solanaceae (NRC2/3/4) are robust enough to detect novelty in the diverse ~350 Solanaceae species dataset without being confounded by natural sequence drift.

#### Feasibility and Effect on Impact

- **Feasibility of Parameter Extraction:** High. The parameters are grounded in standard structural biology metrics (angles, distances, packing ratios). Articles [3] and [9] explicitly show that AF3 can model the NRC hexamer and its N-terminal funnel with high confidence, making parameters 1, 5, and 7 highly realistic to extract.
- **Stoichiometric Contrast:** Article [15] demonstrates that using AF3 scores to rank stoichiometries is a winning strategy in CASP16. This validates the "Tier 3" re-modeling step, suggesting the SNI can indeed distinguish tetramers or octamers from hexamers.
- **Biological Validity:** The inclusion of electrostatics ($E_{flux}$) and hydrophobic moments ($\vec{M}_{align}$) is highly relevant. Article [1] identifies the MADA motif as a pore-former; any deviation in its alignment or charge (Parameter 4/5) would logically indicate a shift from a "helper" to a "sensor" or a novel signal-scaffold function [6].
- **Scalability Concerns:** Modeling 6,000 hexamers is the bottleneck. However, the tiered workflow (Tier 1 modeling at $n=6$ only) mitigates this by providing a high-throughput filter before re-modeling.

#### Suggested Improvements

1. **Monomer Pre-Filtering (mSNI):** To improve scalability, introduce a Tier 0 that calculates a "Monomer SNI." If the internal NB-ARC hinge ($\alpha_{inter}$) or the MADA hydrophobic moment ($\vec{M}_{align}$) in a monomer deviates significantly from 9RI9, it could be prioritized for multimeric modeling.
2. **Motif-Anchored Measurement:** Ensure parameters like "Pitch Deviation" are measured relative to conserved motifs (e.g., P-loop to MHD motif) rather than residue numbers, to account for the diverse insertions/deletions in the 6,000-sequence set [10].
3. **Membrane-Insertion Delta:** Compare the SNI of a sequence modeled *without* oleic acid versus *with* oleic acid. A large improvement in pLDDT at the N-terminus upon adding lipids would be a strong indicator of a functional pore-forming "helper."
4. **Explicit Sensor/Helper Calibration:** Use the lettuce dataset [6] (which already identifies helpers and sensors) as a validation set to tune the SNI weights ($w_n$) before applying them to the full 6,000 sequences.

#### Overall Impact Potential

The idea has **high impact potential**. It moves beyond simple sequence similarity (which often fails to distinguish helpers from sensors in the NRC family) and focuses on the physical execution of immunity (pore formation and stoichiometric fit).

- **Feasibility:** The integration of AF3 metrics with classical physics (electrostatics/hydrophobic moments) is well-supported by recent literature [3, 9, 15].
- **Scope:** By providing a quantitative index, this allows the identification of "Dark NLRs"—proteins that look like NRCs but may form 8-mer giant pores or serve as non-conductive scaffolds.
- **Long-term Implications:** This methodology could be applied to other NLR families (e.g., TIR-NLRs or ZAR1-like singletons) to discover novel immune mechanisms across the plant kingdom. It transitions structural biology from a "one protein at a time" discipline to a "systems structural biology" approach.

While computationally intensive, the tiered implementation makes it a realistic and powerful discovery tool.

Answer: 8

References:

[1] [An N-terminal motif in NLR immune receptors is functionally conserved across distantly related plant species - PMC](https://pmc.ncbi.nlm.nih.gov/articles/PMC6944444/)

[2] [Protein complexes in cells by AI‐assisted structural proteomics - PMC](https://pmc.ncbi.nlm.nih.gov/articles/PMC10090944/)

[3] [A disease resistance protein triggers oligomerization of its NLR helper into a hexameric resistosome to mediate innate immunity - PMC](https://pmc.ncbi.nlm.nih.gov/articles/PMC11540030/)

[4] [An N-terminal motif in NLR immune receptors is functionally conserved across distantly related plant species](https://pubmed.ncbi.nlm.nih.gov/31774397/)

[5] [A disease resistance protein triggers oligomerization of its NLR helper into a hexameric resistosome to mediate innate immunity](https://www.ncbi.nlm.nih.gov/pmc/articles/PMC11540030/)

[6] [A hierarchical immune receptor network in lettuce reveals contrasting patterns of evolution in sensor and helper NLRs](https://www.biorxiv.org/content/10.1101/2025.02.25.639832v1)

[7] [A disease resistance protein triggers oligomerization of its NLR helper into a hexameric resistosome to mediate innate immunity](https://www.ncbi.nlm.nih.gov/pmc/articles/PMC11540030/)

[8] [Confidence scores in AlphaFold-Multimer | AlphaFold](https://www.ebi.ac.uk/training/online/courses/alphafold/inputs-and-outputs/evaluating-alphafolds-predicted-structures-using-confidence-scores/confidence-scores-in-alphafold-multimer/)

[9] [A disease resistance protein triggers oligomerization of its NLR helper into a hexameric resistosome to mediate innate immunity | Request PDF](https://www.researchgate.net/publication/385596452_A_disease_resistance_protein_triggers_oligomerization_of_its_NLR_helper_into_a_hexameric_resistosome_to_mediate_innate_immunity)

[10] [A hierarchical immune receptor network in lettuce reveals contrasting patterns of evolution in sensor and helper NLRs | bioRxiv](https://www.biorxiv.org/content/10.1101/2025.02.25.639832v1.full-text)

[11] [A disease resistance protein triggers oligomerization of its NLR helper into a hexameric resistosome to mediate innate immunity](https://www.biorxiv.org/content/10.1101/2024.06.18.599586v1)

[12] [A hydrophobic core in the coiled-coil domain is essential for NRC resistosome function](https://www.biorxiv.org/content/10.1101/2025.01.21.634219v3)

[13] [A disease resistance protein triggers oligomerization of its NLR helper into a hexameric resistosome to mediate innate immunity](https://www.biorxiv.org/content/10.1101/2024.06.18.599586v1)

[14] [A disease resistance protein triggers oligomerization of its NLR helper into a hexameric resistosome to mediate innate immunity](https://pubmed.ncbi.nlm.nih.gov/39504373)

[15] [Accurate Prediction of Protein Complex Stoichiometry by Integrating AlphaFold3 and Template Information - PMC](https://pmc.ncbi.nlm.nih.gov/articles/PMC11761747/)

[16] [Abstract Introduction](https://www.biorxiv.org/content/10.1101/2022.04.25.489342v1.full.pdf)

[17] [A helper NLR targets organellar membranes to trigger immunity](https://www.biorxiv.org/content/10.1101/2024.09.19.613839v1)

[18] [A disease resistance protein triggers oligomerization of its NLR helper into a hexameric resistosome to mediate innate immunity](https://www.ncbi.nlm.nih.gov/pmc/articles/PMC11540030/)

[19] [A disease resistance protein triggers oligomerization of its NLR helper into a hexameric resistosome to mediate innate immunity](https://www.ncbi.nlm.nih.gov/pmc/articles/PMC11540030/)

[20] [Improving AlphaFold2 and 3-based protein complex structure prediction with MULTICOM4 in CASP16 | bioRxiv](https://www.biorxiv.org/content/10.1101/2025.03.06.641913v1.full)

[21] [A hierarchical immune receptor network in lettuce reveals contrasting patterns of evolution in sensor and helper NLRs](https://www.biorxiv.org/content/10.1101/2025.02.25.639832v1)

[22] [A hydrophobic core in the coiled-coil domain is essential for NRC resistosome function](https://www.biorxiv.org/content/10.1101/2025.01.21.634219v2)

[23] [A disease resistance protein triggers oligomerization of its NLR helper into a hexameric resistosome to mediate innate immunity](https://www.biorxiv.org/content/10.1101/2024.06.18.599586v1)

[24] [A disease resistance protein triggers oligomerization of its NLR helper into a hexameric resistosome to mediate innate immunity](https://www.ncbi.nlm.nih.gov/pmc/articles/PMC11540030/)

[25] [Protein complex prediction with AlphaFold-Multimer | bioRxiv](https://www.biorxiv.org/content/10.1101/2021.10.04.463034.full)

[26] [Helper NLR immune protein NRC3 evolved to evade inhibition by a cyst nematode virulence effector - PMC](https://pmc.ncbi.nlm.nih.gov/articles/PMC11981194/)

[27] [A disease resistance protein triggers oligomerization of its NLR helper into a hexameric resistosome to mediate innate immunity](https://www.biorxiv.org/content/10.1101/2024.06.18.599586v1)

[28] [The MADA motif is a conserved unit at the very N-terminus of NRC4 and... | Download Scientific Diagram](https://www.researchgate.net/figure/The-MADA-motif-is-a-conserved-unit-at-the-very-N-terminus-of-NRC4-and-ZAR1-A-Schematic_fig4_334241019)

[29] [RCSB PDB - 9CC8: Hexameric state of the NRC4 resistosome](https://www.rcsb.org/structure/9cc8)

[30] [Resurrection of plant disease resistance proteins via helper NLR bioengineering - PMC](https://pmc.ncbi.nlm.nih.gov/articles/PMC10156107/)

[31] [The resistance awakens: Diversity at the DNA, RNA, and protein levels informs engineering of plant immune receptors from Arabidopsis to crops](https://www.ncbi.nlm.nih.gov/pmc/articles/PMC12118082/)

[32] [Diversification of the “EDVID” packing motif underpins structural and functional variation in plant NLR coiled-coil domains](https://www.biorxiv.org/content/10.1101/2025.06.01.657260v1)

[33] [Improving AlphaFold2 and 3-based protein complex structure prediction with MULTICOM4 in CASP16](https://www.biorxiv.org/content/10.1101/2025.03.06.641913v1)

[34] [RCSB PDB - 9FP6: Structure of the NbNRC2 hexameric resistosome](https://www.rcsb.org/structure/9fp6)

[35] [An N-terminal motif in NLR immune receptors is functionally conserved across distantly related plant species | eLife](https://elifesciences.org/articles/49956)

[36] [A disease resistance protein triggers oligomerization of its NLR helper into a hexameric resistosome to mediate innate immunity](https://www.biorxiv.org/content/10.1101/2024.06.18.599586v1)

[37] [Structural basis for heat tolerance in plant NLR immune receptors | bioRxiv](https://www.biorxiv.org/content/10.64898/2025.12.17.694812v1.full-text)

[38] [A helper NLR targets organellar membranes to trigger immunity](https://www.biorxiv.org/content/10.1101/2024.09.19.613839v1)

[39] [A helper NLR targets organellar membranes to trigger immunity](https://www.biorxiv.org/content/10.1101/2024.09.19.613839v1)

[40] [Benchmarking AlphaFold3’s protein-protein complex accuracy and machine learning prediction reliability for binding free energy changes upon mutation](https://arxiv.org/html/2406.03979v1)

[41] [NRC Immune receptor networks show diversified hierarchical genetic architecture across plant lineages - PMC](https://pmc.ncbi.nlm.nih.gov/articles/PMC11371147/)

[42] [Structural Insights into the Plant Immune Receptors PRRs and NLRs - PMC](https://pmc.ncbi.nlm.nih.gov/articles/PMC7140948/)

[43] [The MADA motif is conserved in ~20% of CC-NLRs. (A) Schematic... | Download Scientific Diagram](https://www.researchgate.net/figure/The-MADA-motif-is-conserved-in-20-of-CC-NLRs-A-Schematic-representation-of-a_fig5_334241019)

[44] [Structure of the activated Roq1 resistosome directly recognizing the pathogen effector XopQ](https://www.biorxiv.org/content/10.1101/2020.08.13.246413v1)

[45] [Ambient Proteins: Training Diffusion Models on Low Quality Structures](https://www.biorxiv.org/content/10.1101/2025.07.03.663105v1)

[46] [Structures of plant resistosome reveal how NLR immune receptors are activated - PMC](https://pmc.ncbi.nlm.nih.gov/articles/PMC9590527/)

[47] [Structural, Functional, and Genomic Diversity of Plant NLR Proteins: An Evolved Resource for Rational Engineering of Plant Immun](https://www.bmb.colostate.edu/wp-content/uploads/sites/21/2018/11/Montiero-ARPP-2018-Diversity-of-NLR-proteins.pdf)

[48] [Bridging prediction and reality: Comprehensive analysis of experimental and AlphaFold 2 full-length nuclear receptor structures](https://www.ncbi.nlm.nih.gov/pmc/articles/PMC12149446/)

[49] [An atypical NLR protein modulates the NRC immune receptor network in Nicotiana benthamiana | PLOS Genetics](https://journals.plos.org/plosgenetics/article?id=10.1371/journal.pgen.1010500)

**Motivation:**

$\def\mathcal#1{\mathit{#1}}\def\mathscr#1{\mathit{#1}}$

**Coherence:**

$\def\mathcal#1{\mathit{#1}}\def\mathscr#1{\mathit{#1}}$

No incoherence found.

**Deep verification:**

$\def\mathcal#1{\mathit{#1}}\def\mathscr#1{\mathit{#1}}$

Based on the provided assumptions, claims, and subsequent reasoning, the Structural Novelty Index (SNI) hypothesis may be incorrect due to several fundamental flaws in its biological premises, methodological execution, and data interpretation.

The reasons for its potential failure can be summarized into the following categories:

#### 1. Fundamental Misidentification of Structural Anchors

The hypothesis identifies **9RI9** as a "resting-state anchor." However, the reasoning clarifies that 9RI9 is actually a hexameric **active-state resistosome** for NRC3. In reality, the resting state of these proteins is typically a homodimer. By using an active state as a baseline for a resting state, the SNI would produce inherently flawed comparisons and fail to distinguish between activated and inactive sequences.

#### 2. Biological and Stoichiometric Inaccuracies

- **Missing Stoichiometries:** The SNI framework models sequences as tetramers, hexamers, and octamers ($n=4, 6, 8$). Scientific reasoning points out that this omits **pentamers ($n=5$)**, which are known biological states for certain NLRs, while including octamers, which lack established research support in this context.
- **Universal Ligand Assumption:** The framework assumes **oleic acid** is the universal ligand for all 6,000 NRCs. If a sequence requires a different lipid or cofactor, forcing it to model with oleic acid would create artificial "structural strain," leading to inflated SNI scores that reflect modeling errors rather than genuine biological novelty.

#### 3. Methodological and Software Errors

- **Pore Dimension Misinterpretation:** The hypothesis claims the N-terminal pore "throat" is exactly 5 Å. Reasoning indicates this is a misinterpretation of a sampling window; the actual physical diameter of canonical NRC pores is much wider (19–23 Å).
- **Misuse of Analytical Tools:** The framework attributes electrostatic potential measurements to **HOLE software**. However, HOLE is designed for geometric analysis (mapping the width of a pore), while electrostatics require different tools like APBS. This suggests the $E_{flux}$ parameter may be calculated using inappropriate methodology.

#### 4. Sensitivity to Modeling Noise

The SNI relies on a **$5^\circ$ Stoichiometric Pitch Deviation** threshold to identify non-hexameric assembly. Because AlphaFold 3 (AF3) interface coordinates can vary naturally between models, this threshold is likely too narrow. The reasoning suggests this $5^\circ$ deviation falls within the range of **modeling noise**, meaning the SNI would likely flag thousands of sequences as "novel" simply due to minor stochastic variations in the AI's output.

#### 5. Lack of Experimental Validation for Thresholds

Several key parameters—specifically the **MADA Vector Alignment** and **Vertical Planarity**—lack robust, experimentally derived "cutoff" values. Without established biological benchmarks, these parameters become subjective, making it difficult to distinguish between a functional biological shift and a non-functional structural variation.

In summary, the SNI hypothesis is undermined by a reliance on incorrect structural baselines, a narrow and potentially noisy threshold for novelty, and a failure to account for the known diversity of NLR stoichiometries and ligands.
