## Supplementary Data S6 for "AI-guided discovery of atypical protein assemblies": [EXTERNAL] ├░Y╠ê┬o┬1⁄4 Co-scientist - Hypothesis 207811288102211 - A Tiered AlphaFold 3 Screening Approach Using 8-Parameter Structural Novelty Index for Unconventional NRC-NLR Identification - For research purposes only - Google confidential.docx

### 🧬 Co-scientist - A Tiered AlphaFold 3 Screening Approach Using 8-Parameter Structural Novelty Index for Unconventional NRC-NLR Identification

**About**: *This is an experimental system for generating novel and testable hypotheses. The hypotheses are generated by a model and may be wrong. For research purposes only.*

### Proposal

$\def\mathcal#1{\mathit{#1}}\def\mathscr#1{\mathit{#1}}$

#### 1. Methodological Principles: The "Forced Hexamer" Strain Filter

To manage the computational requirements of modeling 6,000 sequences, the methodology utilizes a Tiered Screening approach:

- **Tier 1: Forced Hexamer Modeling ($n=6$):** All sequences are modeled initially as hexamers. Instead of sampling $n=4$ and $n=8$ immediately, the **Interface Tension ($\chi_{int}$)** is measured.
- **Tier 2: Tension-Gated Sampling:** Only sequences that show high "Interface Tension" (significant angular deviation or low ipTM at the $n=6$ interface) or high SNI scores are promoted to $n=4$ or $n=8$ modeling to confirm their preferred stoichiometry. This approach is designed to reduce the total AF3 GPU hours.
- **Direct Interaction Metrics:** This method uses discrete distance and conservation measurements of key functional residues identified in the Ground Truth Dataset (9FP6) in place of complex pore-walking algorithms.

#### 2. Quantitative Parameters of the Structural Novelty Index (SNI)

The SNI is a composite score ($SNI = \sum P_{n} \cdot w_{n}$) where $w_{n}$ is weighted by local pLDDT.

##### **Parameter 1: Stoichiometric Pitch Deviation ($\Delta \theta_{pitch}$)**

- **Measurement:** Angle between adjacent NB-ARC domains in the $n=6$ model.
- **Novelty Signal:** Deviation $> 5^\circ$ from the $60^\circ$ baseline (9FP6/9CC8) suggests the sequence is "squeezed" or "stretched," indicating a preference for $n=5$ or $n=7+$.

##### **Parameter 2: Vertical Planarity ($\delta_{z}$)**

- **Measurement:** The Z-axis displacement between the centroids of adjacent NB-ARC domains.
- **Novelty Signal:** $\delta_{z} > 3.5 \mathring{A}$ indicates a "lock-washer" or helical assembly rather than the flat disc seen in 9FP6.

##### **Parameter 3: NB-ARC Hinge Angle ($\alpha_{hinge}$)**

- **Measurement:** The internal angle between the NB and the WHD subdomains.
- **Novelty Signal:** Canonical NRCs (9FP6) show a "relaxed" hinge. A "tightened" hinge ($< 10^\circ$ difference from 9RI9 resting state) suggests a non-functional or decoy NRC.

##### **Parameter 4: Pore Charge Density ($\sigma_{pore}$)**

- **Measurement:** The sum of formal charges of residues 1–15 (MADA) projected onto the internal surface of the N-terminal funnel.
- **Novelty Signal:** Canonical NRCs are strongly electronegative (cation-selective). A neutral or positive $\sigma_{pore}$ identifies potential anion-selective or non-conductive scaffolds.

##### **Parameter 5: CC-Barrel Hydrophobic Core Integrity ($\Delta D_{core}$)**

- **Measurement:** The average inter-residue distance between the centroids of residues equivalent to L34, A72, I114, and V118 (from PDB 9FP6).
- **Novelty Signal:** A distance increase $> 3 \mathring{A}$ indicates a collapsed or unconventional CC-barrel, suggesting the NLR may not form a stable membrane-inserting pore.

##### **Parameter 6: Bidentate Salt Bridge Occupancy ($d_{bridge}$)**

- **Measurement:** The distance between the acidic pair (E73/D74) in the CC domain and the basic residue (R519) in the LRR of the adjacent protomer.
- **Novelty Signal:** In 9FP6, this bridge stabilizes the hexamer. A distance $> 5 \mathring{A}$ suggests a novel stabilization mechanism or a sequence that cannot form a stable hexamer.

##### **Parameter 7: Radial NB-ARC Displacement ($R_{out}$)**

- **Measurement:** The distance from the pore axis to the NB-ARC centroid.
- **Novelty Signal:** $R_{out} > 55 \mathring{A}$ (higher than 9FP6) suggests a "stretched" interface to accommodate larger stoichiometries ($n=8$).

##### **Parameter 8: MADA-Helix Hydrophobic Moment ($\vec{M}_{align}$)**

- **Measurement:** The vector sum of hydrophobic side chains in the $\alpha$1-helix relative to the pore center.
- **Novelty Signal:** An outward-pointing vector suggests the MADA motif is "flipped" or sequestered, indicating a non-pore-forming "decoy" helper.

#### 3. Implementation Workflow

1. **Phase 1 (Broad Screen):** Run AlphaFold 3 for 6,000 sequences at **$n=6$ only**, including oleic acid to stabilize the N-terminus.
2. **Phase 2 (SNI Calculation):** Extract the 8 parameters. Calculate the **Interface Tension ($\chi_{int}$)** by combining Parameter 1 and Parameter 6.
3. **Phase 3 (Targeted Refinement):**
   - **Low Tension / Low SNI:** Classify as "Canonical NRC."
   - **High Tension / High SNI:** Promote to $n=4$ and $n=8$ modeling to determine the true stoichiometric "sweet spot."
4. **Phase 4 (Functional Mapping):** Map high-SNI candidates against the Solanaceae dataset to identify clades of unconventional NRCs for wet-lab validation (cell death and BN-PAGE).

### Reviews summary

$\def\mathcal#1{\mathit{#1}}\def\mathscr#1{\mathit{#1}}$

### Appendix:

**Motivation:**

$\def\mathcal#1{\mathit{#1}}\def\mathscr#1{\mathit{#1}}$

**Coherence:**

$\def\mathcal#1{\mathit{#1}}\def\mathscr#1{\mathit{#1}}$

**Deep verification:**

$\def\mathcal#1{\mathit{#1}}\def\mathscr#1{\mathit{#1}}$
