## Supplementary Data S6 for "AI-guided discovery of atypical protein assemblies": [EXTERNAL] ├░Y╠ê┬o┬1⁄4 Co-scientist - Hypothesis 216227824734088 - Physico-Evolutionary Structural Novelty Index (PE-SNI) for High-Throughput Identification of Unconventional NRC-NLRs - For research purposes only - Google confidential.docx

#### 🧬 Co-scientist - Physico-Evolutionary Structural Novelty Index (PE-SNI) for High-Throughput Identification of Unconventional NRC-NLRs

**About**: *This is an experimental system for generating novel and testable hypotheses. The hypotheses are generated by a model and may be wrong. For research purposes only.*

#### Proposal

$\def\mathcal#1{\mathit{#1}}\def\mathscr#1{\mathit{#1}}$

### The Physico-Evolutionary Structural Novelty Index (PE-SNI)

#### 1. Executive Summary

The Physico-Evolutionary Structural Novelty Index (PE-SNI) is a high-throughput quantitative framework designed to classify the ~6,000 Solanaceae NRC sequences. The PE-SNI evaluates sequences based on their structural behavior and energetic competence relative to canonical Solanaceae resistosomes (NRC2, NRC3, NRC4).

By calibrating against a Ground Truth Dataset—Inactive NbNRC2 (PDB: 9RI9), Active NbNRC2 (PDB: 9FP6), and NRC4 (PDB: 9CC8)—the PE-SNI is designed to filter out modeling artifacts and identify unconventional NLRs by detecting specific deviations in folding dynamics, interface energetics, and switch mechanisms.

#### 2. The Seven Quantitative Parameters

The PE-SNI is a composite score derived from the following seven parameters. An NRC is classified as "Unconventional" if its aggregate Z-score deviates significantly from the canonical baseline.

##### Parameter 1: P-loop Cleft Solvation Depth ($D_{solv}$)

- **Unit:** Angstroms (Å)
- **Method:** Measurement of the Solvent Accessible Surface Area (SASA) depth relative to the P-loop Lysine $C\alpha$.
- **Rationale:** Canonical NRCs maintain a specific burial depth for the nucleotide (ADP/ATP) to regulate the "switch."
  - **Canonical Baseline:** Consistent depth (~8–10 Å) matching PDB 9RI9.
  - **Unconventional Signal:** A shallow ($<5$ Å) or hyper-buried ($>15$ Å) cleft suggests the protein may have lost nucleotide-dependent regulation.

##### Parameter 2: CC-NB-ARC Resting Tilt ($\theta_{rest}$)

- **Unit:** Degrees (°)
- **Method:** Using the ABangle framework, measure the angle between the principal axis of the N-terminal CC domain and the centroid plane of the NB-ARC domain in the monomeric AF3 model.
- **Rationale:** As seen in PDB 9CC8 and 9RI9, canonical NRCs adopt a specific tilt (~85° relative to the resistosome plane, or acute internal angles) to prevent premature oligomerization.
  - **Unconventional Signal:** A tilt deviating significantly (e.g., linear alignment or perpendicular locking) indicates the loss of the standard activation mechanism.

##### Parameter 3: Interface Frustration Energy ($E_{if}$)

- **Unit:** kcal/mol per $Å^2$ (Interface Energy / Buried Surface Area)
- **Method:** Generate a "forced hexamer" in AF3, then score the protomer-protomer interface using FoldX or Rosetta.
- **Rationale:** AF3 can force non-interacting proteins into a ring shape. This parameter evaluates chemical plausibility.
  - **Canonical Baseline:** High shape complementarity leads to favorable (negative) binding energy (calibrated to 9FP6).
  - **Unconventional Signal:** High "frustration" (positive or neutral energy) despite geometric fit indicates the protein may not physically form the canonical ring, categorizing it as an unconventional variant.

##### Parameter 4: MADA-Motif Radial Vector Variance ($\sigma^2_{RV}$)

- **Unit:** Angstrom Variance ($Å^2$)
- **Method:** Run five distinct AF3 seeds for the forced hexamer. Calculate the variance in the radial distance of the N-terminal MADA motif (L13-L17) to the central pore axis.
- **Rationale:** Variance measures structural competence.
  - **Canonical Baseline:** Low variance. The N-terminal positioning is consistent for forming the funnel (as seen in 9FP6).
  - **Unconventional Signal:** High variance. The N-termini lack a consistent structural blueprint across seeds to form a pore, pointing to non-pore-forming functions.

##### Parameter 5: Ligand-Induced Switch Delta ($\Delta D_{PM}$)

- **Unit:** Angstroms (Å)
- **Method:** Calculate the Euclidean distance between the P-loop (Walker A) and MHD-motif Histidine in two AF3 runs: one constrained with ADP and one with ATP-$\gamma$-S.
- **Rationale:** Quantifies the mechanical work potential of the NB-ARC switch.
  - **Canonical Baseline:** Significant shift ($\Delta > 4.0$ Å) reflecting the transition from Closed (9RI9) to Open (9FP6) states.
  - **Unconventional Signal:** Near-zero delta ("Dead Switch") or chaotic delta ("Unstable Switch"), indicating the protein does not utilize the canonical ATP-driven conformational change.

##### Parameter 6: Covariance-Weighted Apolar Density ($\rho_{EAD}$)

- **Unit:** Weighted Contacts / $100 Å^2$
- **Method:** Count carbon-carbon contacts at the forced interface (distance $<5$ Å). Weight each contact by the ConSurf evolutionary conservation score of the participating residues.
- **Rationale:** Distinguishes signal from noise. AF3 may pack variable hydrophobic residues together to fill space.
  - **Canonical Baseline:** High density of conserved hydrophobic contacts.
  - **Unconventional Signal:** Low weighted density. Residues are evolutionarily transient, suggesting the interface is not under selection pressure and is likely an artifact.

##### Parameter 7: Linker Ensemble Dynamics (eRMSF)

- **Unit:** Root Mean Square Fluctuation (Å)
- **Method:** Calculate the RMSF of the CC-NB-ARC linker loop residues across the five AF3 seeds.
- **Rationale:** Differentiates functional flexibility from disordered loops.
  - **Canonical Baseline:** Defined flexibility patterns (hinge-like behavior) allowing the CC domain to pivot.
  - **Unconventional Signal:** Extreme rigidity or extreme disorder (eRMSF > 5 Å) incompatible with the controlled unfolding required for resistosome assembly.

#### 3. Implementation Protocol

1. **High-Throughput Modeling:**
   - Input: ~6,000 NRC sequences.
   - Process: Run AF3 in two modes per sequence: Monomer (ADP-constrained) and Forced Hexamer (ATP-constrained). Generate five seeds per mode.
2. **Parameter Extraction:**
   - **Structural Parsing:** Use Python (Biopython/PDB module) to extract coordinates for Parameters 1, 2, 4, 5, and 7.
   - **Energetic Scoring:** Pipe "Forced Hexamer" PDBs into FoldX to calculate $E_{if}$ (Parameter 3).
   - **Evolutionary Mapping:** Map existing MSA conservation scores to interface residues for $\rho_{EAD}$ (Parameter 6).
3. **SNI Calculation & Filtering:**
   - Normalize scores against the Reference Set (NbNRC2, SlNRC3, NRC4).
   - Compute SNI Score: $\sum (Z_{param})$.
   - **Threshold:** Sequences with SNI > 2.5 are flagged as "Unconventional Candidates" for downstream validation.

#### 4. Final Answer: Methodological Advantages

This approach moves beyond looking for sequence-level differences and instead evaluates how proteins behave under physical simulation.

- By using Energy ($E_{if}$), the framework aims to reduce false positives where AF3 creates hexamers that may lack chemical stability.
- By using Variance ($\sigma^2_{RV}$), AF3's uncertainty is utilized as a data point for detecting novel N-terminal structures.
- By calibrating against PDB 9FP6/9RI9, the index measures biological deviation rather than just sequence divergence.

#### Reviews summary

$\def\mathcal#1{\mathit{#1}}\def\mathscr#1{\mathit{#1}}$

Positive aspects:

- **Mechanistic and Functional Focus:** Unlike standard sequence alignments or global RMSD checks, the PE-SNI evaluates specific biophysical mechanisms, such as the "mechanical work potential" of the NB-ARC switch and the solvation depth of the P-loop cleft. This provides deeper insight into whether a protein is a functional "engine" or a signaling decoy.
- **Innovative Use of AI Uncertainty:** The index utilizes the stochasticity of AlphaFold 3 (variance across multiple seeds) as a quantitative metric for structural competence. High variance in the N-terminal MADA motif suggests a lack of a robust structural blueprint for pore formation, effectively identifying unconventional variants.
- **Energetic Scrutiny to Filter Artifacts:** By incorporating Interface Frustration Energy ($E_{if}$) via FoldX or Rosetta, the framework can detect if AlphaFold has "forced" a non-interacting protein into a hexameric ring. This helps distinguish biologically plausible complexes from modeling artifacts.
- **Objective and Automatable Pipeline:** The parameters are anchored to strictly conserved molecular "landmarks" (e.g., P-loop Lysine, MHD Histidine). This ensures the coordinates can be extracted via script for all 6,000 sequences, making the high-throughput screening objective and reproducible.
- **Integration of Evolutionary Signal:** Parameter 6 (Covariance-Weighted Apolar Density) weights structural contacts by evolutionary conservation. This allows the index to distinguish between evolutionarily selected functional interfaces and transient hydrophobic packing, increasing the accuracy of the novelty detection.

Negative aspects:

- **Critical Factual Error in Ground Truth:** The framework misidentifies PDB 9RI9 as "Inactive NbNRC2" when it is actually the "Active SlNRC3" hexamer. This error invalidates the baseline for the "Switch Delta" and "Solvation Depth" parameters, as the canonical baseline would be comparing two active states, leading to false "dead switch" signals.
- **Prohibitive Computational Burden:** Modeling 6,000 sequences as hexamers (averaging ~5,400 residues per complex) with 5 seeds each would require an estimated 150,000 to 240,000 GPU-hours. Without a tiered screening approach (e.g., filtering via monomers first), this is unfeasible for most research environments.
- **Statistical Instability of the Index:** A reference set of only three proteins (9FP6, 9RI9, 9CC8) is insufficient to calculate stable standard deviations for Z-score normalization. This makes the entire index highly sensitive to minor coordinate noise in the ground truth models.
- **Omission of Key Domains and Confidence Metrics:** The index fails to incorporate LRR-specific parameters despite their importance in receptor orientation. Furthermore, it neglects standard AlphaFold confidence metrics (ipTM and pLDDT), risking the classification of high-confidence modeling failures as biological "novelties."
- **Double-Counting of Correlated Signals:** The simple additive Z-score model assumes the seven parameters are independent. In reality, metrics like interface energy, tilt, and switch dynamics are likely highly correlated; summing them leads to redundant signals and inflated false-positive rates.

##### Research contacts

$\def\mathcal#1{\mathit{#1}}\def\mathscr#1{\mathit{#1}}$

Based on the provided research paper excerpts, the following researchers are the most suitable candidates to review the **Physico-Evolutionary Structural Novelty Index (PE-SNI)** hypothesis. They are listed in order of their expertise regarding the specific structural and computational aspects of the NRC family and AlphaFold 3 (AF3) modeling.

##### 1. AmirAli Toghani

**Justification:** Toghani is a primary researcher involved in the most specific aspects of this hypothesis: using AlphaFold 3 to distinguish between different types of NRCs and benchmarking resistosome structures.

- **Relevance:** Toghani is a lead/key author on studies assessing AF3’s ability to predict activated CC-NLR oligomers and distinguish "Helper" (NRC-H) from "Sensor" (NRC-S) NLRs based on their ability to form hexameric resistosomes (**Abstracts 5, 9, 13**).
- **Specific Evidence:** His work specifically utilizes structural metrics like **pTM, ipTM, and pLDDT** to distinguish unconventional sequences (like NRC-S) from canonical ones (**Abstract 9**), which aligns perfectly with the goal of creating a quantitative SNI.

##### 2. Madhuprakash Jogi (or J. Madhuprakash)

**Justification:** Jogi is the lead researcher on the structural determination of the ground truth dataset mentioned in the hypothesis (specifically PDB: 9FP6).

- **Relevance:** He led the study "*Structure of the NbNRC2 hexameric resistosome*" (**Abstract 1**). His research provides the exact "Canonical Baseline" metrics requested, such as the 10° difference in interdomain angles between hexameric NRC2 and pentameric ZAR1 (**Abstract 3**).
- **Specific Evidence:** His work explicitly details the structural rearrangements between the resting and activated states of NbNRC2 and compares protomer interface packing—the core requirements for Parameters 2 and 3 of the PE-SNI (**Abstract 3, 8**).

##### 3. Sophien Kamoun

**Justification:** Kamoun is a senior author on nearly all provided articles regarding the NRC immune receptor network, its evolution, and its structural biology.

- **Relevance:** He oversaw the development of the "activation-and-release" model and the phylogenomic analysis of ~1,000 NRC sequences across the Solanaceae family (**Abstract 8, 10**).
- **Specific Evidence:** His lab’s research provided the foundational dataset for the 6,000 sequences mentioned in the rationale and established the importance of the **MADA motif** (Parameter 4) in NRC-H versus its degeneration in NRC-S (**Abstract 14**).

##### 4. Michael W. Webster

**Justification:** Webster is a structural biologist and deposition author for the NbNRC2 hexameric resistosome structure (**PDB: 9FP6**).

- **Relevance:** His expertise lies in the high-resolution cryo-EM analysis of NRCs and evaluating the accuracy of AF3 in modeling N-terminal $\alpha$1 helices (**Abstract 1**).
- **Specific Evidence:** He is uniquely qualified to review the physical plausibility of the PE-SNI's "Forced Hexamer" approach (Parameter 3) and the use of radial vector variance in AF3 seeds (Parameter 4) (**Abstract 1**).

##### 5. Chih-Hang Wu

**Justification:** Wu is an expert in the transition of NRCs from homodimers to resistosomes and the identification of conserved motifs.

- **Relevance:** He is the lead author of the study on the NbNRC2 resting state homodimer (**PDB: 8RFH/9RI9**) and the original discoverer of the **MADA motif** (**Abstract 10, 14**).
- **Specific Evidence:** His work on the dimerization interface and the "insulation" between different NRC clades provides the evolutionary and biochemical basis for Parameter 6 (Covariance-Weighted Apolar Density) and Parameter 1 (P-loop Cleft Solvation) (**Abstract 10**).

##### 6. Brian G. Pierce

**Justification:** Pierce specializes in the quantitative benchmarking of AlphaFold modeling and interface scoring.

- **Relevance:** He authored "*Benchmarking AlphaFold for protein complex modeling reveals accuracy determinants*" (**Abstract 4**).
- **Specific Evidence:** His work focuses on the exact computational metrics proposed in the PE-SNI, including **Rosetta cross-interface binding energy, shape complementarity, and interface residues** (**Abstract 4**). He could provide a critical review of the "Interface Frustration Energy" (Parameter 3) and "Apolar Density" (Parameter 6).

##### 7. Hsuan Pai

**Justification:** Pai led recent research specifically testing the "resistosome-forming capacity" of various NRC subclades using AF3.

- **Relevance:** Lead author of the study comparing lettuce NRC-H and NRC-S structures using AF3 simulations (**Abstract 9**).
- **Specific Evidence:** His work established the "pTM/ipTM < 0.5" threshold for identifying unconventional/non-functional NRCs, which is the direct precursor to the "SNI Score > 2.5" threshold proposed in the implementation protocol (**Abstract 9**).

#### Appendix:

**All reviews:**

**Correctness:**

$\def\mathcal#1{\mathit{#1}}\def\mathscr#1{\mathit{#1}}$

##### Related Article Abstracts

1. **[1] RCSB PDB - 9FP6: Structure of the NbNRC2 hexameric resistosome**: Essential for the "Active" canonical baseline. Confirms NRC2 forms hexamers and that AF3 can model the N-terminal $\alpha 1$ helices accurately.
2. **[10] Activation of plant immunity through conversion of a helper NLR homodimer into a resistosome (PDB: 8RFH/9RI9)**: Provides the "Inactive" resting state baseline. Explains that NRC2 exists as a dimer in its resting state, which is crucial for defining $D_{solv}$ and $\theta_{rest}$.
3. **[18] A hierarchical immune receptor network in lettuce reveals contrasting patterns of evolution...**: Demonstrates that AF3 can distinguish helpers (which form resistosomes) from sensors (which fail to do so), supporting the rationale for the SNI.
4. **[16] A hydrophobic core in the coiled-coil domain is essential for NRC resistosome function**: Validates the importance of CC-domain structural integrity, supporting Parameter 2 and Parameter 4.
5. **[3] A disease resistance protein triggers oligomerization... (NbNRC2 comparison)**: Details inter-domain angles and module rotations ($180^\circ$) between resting and active states, providing geometric evidence for Parameter 2 and Parameter 5.
6. **[14] An N-terminal motif in NLR immune receptors is functionally conserved...**: Establishes the MADA motif as the "death switch" signature, justifying Parameter 4 ($\sigma^2_{RV}$) and Parameter 7 (eRMSF).
7. **[2] Analysing protein complexes in plant science: insights and limitation with AlphaFold 3**: Confirms AF3’s ability to model ligands (ATP/ADP) and provide confidence metrics (ipTM), enabling Parameters 4, 5, and 7.
8. **[4] Benchmarking AlphaFold for protein complex modeling...**: Shows that Rosetta cross-interface binding energy correlates with model quality, validating Parameter 3 ($E_{if}$).
9. **[13] A disease resistance protein triggers oligomerization... (Benchmarks)**: Provides direct ipTM/pTM benchmarks for NRC2 hexamers and pentamers, essential for calibrating the SNI.
10. **[19] Structural mechanism of heavy metal-associated integrated domain engineering...**: Discusses paired NLR structural characteristics and the role of the nucleotide-binding domain as a switch, supporting Parameter 5.
11. **[15] Can AI modelling of protein structures distinguish between sensor and helper NLR immune receptors?**: Further supports the use of AF3 structural metrics (ipTM/pTM) to identify unconventional/sensor NLRs.
12. **[17] Biochemical basis of activation and inhibition of an NLR immune receptor network**: Discusses the "activation and release" model and helper oligomerization, supporting the biological context of the SNI.
13. **[8] A disease resistance protein triggers oligomerization... (Diversity)**: Discusses the diversity in MADA-CC-NLR oligomerization (hexamers vs pentamers), justifying the need for an index to detect unconventional variations.
14. **[12] A Suite of Designed Protein Cages...**: Discusses Buried Surface Area (BSA) and interface solvation energy density as filters for protein complexes, providing methodological support for Parameter 3 and Parameter 6.
15. **[11] A disease resistance protein triggers oligomerization... (N-terminal Focus)**: Focuses on N-terminal structural models and clade-specific variations, supporting Parameters 2 and 4.

##### Detailed Assumptions

1. **Computational Feasibility**: It is assumed that modeling ~6,000 sequences as both monomers and "forced hexamers" using AF3 (5 seeds each) is achievable within a reasonable timeframe/budget.
2. **Structural Frustration**: The idea assumes that unconventional NRCs, when "forced" into a hexameric ring by AF3, will exhibit higher interface energy ($E_{if}$) or structural variance ($\sigma^2_{RV}$) than canonical NRCs.
3. **Stability of Geometric Baselines**: It assumes that metrics like "Resting Tilt" ($\theta_{rest}$) and "Solvation Depth" ($D_{solv}$) are sufficiently conserved across canonical NRC2, NRC3, and NRC4 to serve as a baseline for Z-score calculation.
4. **Ligand-Modeling Accuracy**: It assumes AF3 can accurately differentiate the conformational "switch" when constrained with ADP versus ATP-$\gamma$-S.
5. **Evolutionary Signal**: It assumes that functionally relevant interfaces in helpers will show higher evolutionary conservation (Parameter 6) than artifactual interfaces produced by AI modeling of sensors or unconventional NLRs.

##### Comparison with Knowledge Base and Abstracts

- **Feasibility**: The Knowledge Base confirms that AF3 supports specify random seeds and ligands (ATP/ADP), enabling Parameters 4, 5, and 7. However, modeling 6,000 hexamers (~5,000 residues each) is a massive task. Abstract [2] notes AF3 is faster but Abstract [12] implies substantial GPU time for such volumes.
- **Resting State Baseline**: The idea correctly identifies PDB 9RI9/8RFH as the inactive baseline. Abstract [10] confirms that the resting state is a homodimer and the CC domain is often disordered in experimental density, which justifies using AF3 monomers to calibrate $\theta_{rest}$ and $D_{solv}$.
- **Geometric Variance**: The PDB audit noted a significant difference in CC-NB-ARC angles between 9FP6 and 9CC8 ($135^\circ$ vs $49^\circ$). This conflicts with the idea’s assumption of a "consistent baseline" for Parameter 2 unless the index uses modeling-derived baselines instead of experimental ones.
- **Switch Mechanism**: Parameter 5 ($\Delta D_{PM}$) measures the distance between Walker A and the MHD Histidine. Abstract [10] and [19] confirm the NB-ARC domain acts as a molecular switch, and the transition from closed to open involves substantial module rotation ($180^\circ$), validating the 4.0 Å delta assumption.

##### Reasoning about Correctness for Assumptions

1. **Computational Feasibility**: **Plausible but marginal**. 6,000 hexamers x 5 seeds = 30,000 AF3 runs. Given AF3's token limits and residue count, this requires significant HPC infrastructure.
2. **Structural Frustration**: **True**. Abstracts [4] and [18] confirm that non-functional NLRs (sensors) fail to form high-confidence AF resistosomes. Measuring energy density ($E_{if}$) is a standard structural biology approach to detect artifactual folds.
3. **Stability of Geometric Baselines**: **True (if modeled)**. While experimental angles vary (PDB Audit), the idea proposes modeling the *Reference Set* (NRC2, 3, 4) using the *same AF3 protocol*. This ensures the "Canonical Baseline" is internally consistent with the "Unconventional Candidate" data.
4. **Ligand-Induced Switch**: **True**. AF3 has demonstrated superior ability to model ligand-induced conformational changes compared to AF2.
5. **Evolutionary Signal**: **True**. Abstract [18] and [10] show that helper interfaces are conserved to maintain specific signaling pathways, while sensors diverge. Parameter 6 effectively filters out high-confidence "hallucinations" that lack evolutionary pressure.

##### Strength of Evidence

- **Direct Supporting Evidence**:
  - AF3 can distinguish sensor vs helper based on resistosome-forming capacity (Abstract [18]).
  - NRC2 and NRC4 have resolved hexameric structures (PDB 9FP6, 9CC8) and dimeric resting states (PDB 9RI9).
  - Specific motifs (MADA, Walker A, MHD) are confirmed in NRC4 (Knowledge Base).
- **Indirect Supporting Evidence**:
  - FoldX/Rosetta scoring correlates with experimental stability in other protein systems (Abstract [4]).
  - The "death switch" mechanism involving the N-terminal $\alpha 1$ helix is conserved across MADA-NLRs (Abstract [14]).

##### Suggested Improvements

1. **Tiered Screening**: To solve the feasibility issue, perform Parameter 1, 2, and 5 on *monomers* first. Only model hexamers for sequences that pass the monomeric "switch potential" check.
2. **Stoichiometric Contrast**: The guidelines suggest NRCs can be pentameric or hexameric. The PE-SNI should include a "Stoichiometry Fit" parameter, comparing ipTM scores for pentamer vs hexamer models.
3. **Anchor Parameters**: Ensure the distance measurements ($D_{solv}$, $\Delta D_{PM}$) are anchored to the *C-alpha* of strictly conserved motifs (e.g., the MHD Histidine and the P-loop Lysine) to handle sequence length variations across the ~350 species.

##### Assessment of Goal Requirements

- **5-10 parameters?** Yes (7 defined).
- **Measurable from PDB/AF3?** Yes (all coordinates and energy).
- **Distinguish unconventional vs NRC2/3/4?** Yes, specifically addresses structural and energetic divergence.
- **Incorporate 9FP6, 9RI9, 9CC8?** Yes, they are the calibration core.
- **Focus on protomer angles/domain distances?** Yes (Parameters 2, 4, 5).
- **High-throughput design?** Yes, uses automatable scripting and Z-scores.
- **Grounded in structural biology?** Yes, moves beyond sequence to energy and solvation.

##### Reasoning about Correctness

The PE-SNI is a technically robust framework. It addresses the primary weakness of AI modeling—the tendency to "force" sequences into canonical shapes—by using energetic frustration and ensemble variance (RMSF/radial distance variance) as data points. The choice of parameters reflects a deep understanding of the STAND protein switch mechanism (Walker A to MHD Histidine distance) and the execute-or-detect functional split in the NRC family (MADA variance and interface conservation). While computationally expensive, it is biologically and technically sound. It leverages the most recent 2024 experimental data (NRC2/4 hexamers) to set its baseline.

I recommend testing this idea. It is scientifically novel, quantitative, and directly fulfills the requirements for a structural biology-based screening tool.

Answer: 9

**Novelty:**

$\def\mathcal#1{\mathit{#1}}\def\mathscr#1{\mathit{#1}}$

##### Related Article Abstracts

1. **[1] Structure of the NbNRC2 hexameric resistosome (9FP6)**: Provides the primary ground truth for the active, hexameric state of a canonical NRC helper.
2. **[10] Activation of plant immunity through conversion of a helper NLR homodimer into a resistosome (8RFH)**: Defines the autoinhibited, dimeric resting state of NRC2, essential for calibrating resting tilt and switch parameters.
3. **[8] A disease resistance protein triggers oligomerization of its NLR helper into a hexameric resistosome**: Analyzes the structural basis of hexamer formation and measures interdomain angles (CC-NB-ARC), providing a precedent for SNI Parameter 2.
4. **[9] A hierarchical immune receptor network in lettuce...**: Demonstrates that AlphaFold 3 (AF3) can distinguish helpers (which form hexamers) from sensors (which fail), using pTM and ipTM.
5. **[4] Benchmarking AlphaFold for protein complex modeling...**: Evaluates alternative scoring metrics like cross-interface binding energy and shape complementarity, which are refined in the idea's "Interface Frustration Energy."
6. **[14] An N-terminal motif in NLR immune receptors is functionally conserved (MADA)**: Establishes the MADA motif's role in pore formation, supporting the focus of SNI Parameter 4.
7. **[16] A hydrophobic core in the coiled-coil domain...**: Uses AF3 to identify structural features within the CC domain necessary for function, validating the use of structural modeling for internal domain metrics.
8. **[34] In silico prediction method for plant NLR–effector interactions**: Explores the use of binding energy and affinity to classify "true" vs. "forced" interactions, which parallels the idea's "Interface Frustration Energy."
9. **[36] NLRexpress... Reveals motif stability...**: Identifies conserved motifs (EDVID, MHD, P-loop) across NLRs, providing the sequence landmarks needed for structural parameter extraction.
10. **[15] Can AI modelling... distinguish between sensor and helper NLR immune receptors?**: Direct precedent for using AF3 to classify the NRC family based on oligomeric structural quality.
11. **[21] Sensor NLR immune proteins activate oligomerization of their NRC helpers...**: Establishes the "activation-and-release" model, underpinning the rationale for SNI Parameter 5 (mechanical switch potential).
12. **[28] AlphaDesign: A de novo protein design framework based on AlphaFold**: Uses Rosetta binding energy and interface packing to validate AF-predicted complexes, providing a methodology for SNI Parameter 3.
13. **[12] A Suite of Designed Protein Cages...**: Employs shape complementarity and Buried Surface Area (BSA) as filters for AF-curated structures, validating the idea's use of energy/BSA.
14. **[31] Diversification of the “EDVID” packing motif...**: Investigates how motifs stabilize resistosome architecture, supporting the idea's focus on interdomain orientations.
15. **[25] A helper NLR targets organellar membranes to trigger immunity**: Uses AF3 to compare CC funnel lengths and structures, validating Parameter 4's focus on N-terminal geometry.

##### Aspects Already Explored

- **Using AF3 to distinguish NRC Helpers and Sensors**: Multiple studies [5, 9, 15] already demonstrate that canonical NRC helpers form high-confidence hexamers in AF3, while sensor NLRs (unconventional variants) fail to form these structures or show low ipTM scores.
- **Domain Angles and Rearrangements**: The comparison of interdomain angles (e.g., CC-NB-ARC) and the 180° rotation of modules between resting and active states is documented in detail for NbNRC2 [3, 8, 10].
- **Interface Energy Scoring**: Using Rosetta or FoldX to score the validity of AlphaFold-predicted interfaces ("true" vs. "forced") is an established benchmarking technique in computational structural biology [4, 34, 28].
- **MADA Motif and Pore Formation**: The definition of the MADA motif and its structural transition into a funnel-like pore is well-established [14, 24, 25].
- **Conserved Landmarks**: The use of motifs like the P-loop, MHD, and Walker A as landmarks for measuring conformational changes is standard practice in NLR research [36].

##### Novel Aspects

- **The Composite Z-Score Framework (SNI)**: While researchers use individual metrics (like ipTM or RMSD), the integration of seven distinct physico-evolutionary parameters into a unified "Novelty Index" for the purpose of identifying "unconventional" NRCs is novel.
- **Parameter 4: Seed-Based Variance ($\sigma^2_{RV}$)**: Utilizing the stochasticity of AF3 (the variance across different seeds) as a quantitative indicator of "structural competence" is a highly innovative use of deep-learning uncertainty. It assumes that functional proteins have a robust blueprint, whereas artifactual/novel ones will produce chaotic N-terminal funnel geometries across seeds.
- **Parameter 5: Ligand-Induced Switch Delta ($\Delta D_{PM}$)**: Running parallel AF3 simulations constrained with specific ligands (ADP vs. ATP-$\gamma$-S) to measure the "mechanical work potential" of the NB-ARC switch is a sophisticated advancement over standard folding.
- **Parameter 6: Covariance-Weighted Apolar Density ($\rho_{EAD}$)**: This parameter innovatively weights structural contacts by evolutionary conservation (ConSurf). Standard structural biology looks at contacts; standard phylogeny looks at conservation. Combining them to filter AF3 artifacts (space-filling hydrophobics vs. evolutionarily selected interfaces) is novel in this context.
- **Parameter 1: P-loop Solvation Depth**: While cleft burial is understood, its formulation as a specific high-throughput metric to detect the loss of nucleotide-dependent regulation is a specialized diagnostic tool not seen in current general screenings.

##### Novelty Review

The idea is **highly novel**. While the underlying tools (AF3, Rosetta, ConSurf) and the biological system (NRCs) are well-studied, the specific construction of the PE-SNI represents a paradigm shift from **descriptive** structural biology to **diagnostic** structural informatics. Current papers [9, 15] mostly ask, "Does it form a hexamer?" This idea asks, "How much does its physical and evolutionary behavior deviate from the canonical engine?"

Be strict: Is it just a combination of known things? Not quite. Parameter 4 (Variance) and Parameter 5 (Switch Delta) represent original ways to "interrogate" the AF3 model rather than just taking its top output. Most researchers use RMSD to compare structures; this framework replaces global RMSD with functional site metrics (Solvation depth, Switch delta, Rest tilt), which provide significantly more biological insight.

##### Reasoning and Recommendation

The PE-SNI is a robust, well-grounded methodological proposal. It effectively leverages the provided Ground Truth data (9FP6, 9RI9, 9CC8) to define a mathematical baseline for "canonical behavior."

**Recommendation:** The idea should be tested. Screening 6,000 sequences manually is impossible, and relying solely on AF3 confidence scores (pTM/ipTM) is prone to "false ring" artifacts where AF3 forces residues to pack. The PE-SNI adds layers of energetic and evolutionary scrutiny that would likely identify genuinely novel signaling mechanisms (e.g., non-pore-forming executors or ligand-independent variants) that standard sequence alignments or simple structure checks would miss.

Answer: 8

**Feasibility:**

$\def\mathcal#1{\mathit{#1}}\def\mathscr#1{\mathit{#1}}$

##### Related Article Abstracts

1. **[1] Structure of the NbNRC2 hexameric resistosome (PDB: 9FP6):** Provides the fundamental ground truth for the active, hexameric state of a canonical NRC, essential for calibrating Parameters 3, 4, 5, and 6.
2. **[10] Activation of plant immunity through conversion of a helper NLR homodimer into a resistosome (PDB: 8RFH/9RI9):** Establishes the structural basis for the resting state (dimer), which is critical for calibrating Parameters 1, 2, and 5 (the "switch" distance).
3. **[3] A disease resistance protein triggers oligomerization... (Comparative structural analysis):** Details the $180^\circ$ rotation of structural modules and $10^\circ$ interdomain angle shifts between resting and active states, providing geometric logic for Parameter 2.
4. **[2] Analysing protein complexes in plant science: insights and limitation with AlphaFold 3:** Discusses AF3’s scalability and memory constraints for large complexes (~3,000–5,000 residues), which is a key bottleneck for the proposed "forced hexamer" modeling of 6,000 sequences.
5. **[16] A hydrophobic core in the coiled-coil domain is essential for NRC resistosome function:** Identifies specific conserved hydrophobic residues in the CC domain ($\alpha 2$–$\alpha 4$) that are essential for oligomerization, supporting the biological relevance of Parameters 3, 4, and 6.
6. **[9] A hierarchical immune receptor network in lettuce... (AF3 sensor vs helper):** Demonstrates that AF3 can already distinguish helpers from sensors based on their ability to form hexamers (pTM/ipTM), providing a validation pathway for the PE-SNI.
7. **[4] Benchmarking AlphaFold for protein complex modeling reveals accuracy determinants:** Validates the use of interface binding energy (calculated via Rosetta/FoldX) as a reliable metric for model quality, supporting Parameter 3.
8. **[6] How to assess the quality of AlphaFold 3 predictions:** Explains the interpretation of ipTM, pTM, and PAE metrics, which are necessary to ensure that the structural deviations measured in the SNI represent biological signal rather than modeling noise.
9. **[14] An N-terminal motif in NLR immune receptors is functionally conserved...:** Defines the MADA motif and its "death switch" role, providing the structural anchor for Parameter 4 ($\sigma^2_{RV}$).
10. **[12] A Suite of Designed Protein Cages Using Machine Learning Algorithms...:** Provides a precedent for using shape complementarity and solvation free energy as filters for protein complexes, supporting the use of $E_{if}$ and $D_{solv}$ (Parameters 1 and 3).
11. **[13] A disease resistance protein triggers oligomerization... (Benchmarks):** Compares AF3 hexameric vs. pentameric models, reinforcing the need for stoichiometric contrast in identifying unconventional NLRs.

##### Steps to Test the Idea

1. **Phase I: Pilot Ground Truth Validation (Go/No-Go):**
   - Run the full PE-SNI protocol on the three Ground Truth sequences: NbNRC2 (Active/Inactive), SlNRC3, and NRC4.
   - Calculate the baseline values for all seven parameters.
   - **Go/No-Go Criteria:** If the PE-SNI fails to place NbNRC2 and NRC4 within the "Canonical" range ($Z < 2.5$) or if it cannot distinguish them from a known sensor (e.g., Rx or Rpi-blb2 modeled as a "forced hexamer"), the index parameters must be redefined.
2. **Phase II: Pipeline Automation:**
   - Develop Python scripts using Biopython and the PDB module to automate the calculation of $D_{solv}$, $\theta_{rest}$, $\Delta D_{PM}$, and $\sigma^2_{RV}$.
   - Integrate a FoldX wrapper to process the "Forced Hexamer" outputs for $E_{if}$.
   - Map ConSurf conservation scores onto the AF3 coordinates for Parameter 6 ($\rho_{EAD}$).
3. **Phase III: Representative Subsample Screening:**
   - Before running 6,000 sequences, model a subsample of 100 NLRs (50 helpers, 50 putative sensors) to determine the statistical variance within the family and refine the Z-score normalization.
4. **Phase IV: High-Throughput Family Screen:**
   - Model the full ~6,000 NRC sequences in monomeric (ADP) and forced hexameric (ATP) states.
   - Execute the PE-SNI pipeline and identify "Unconventional" candidates ($Z > 2.5$).
5. **Phase V: Biological Validation:**
   - Correlate the PE-SNI scores with phylogenetic data (e.g., distance from the MADA-helper clades).
   - Validate a subset of high-novelty candidates via wet-lab assays (BN-PAGE or cell death assays).

##### Reasoning About Feasibility

The testing of the PE-SNI is **feasible but resource-intensive**, primarily due to the massive computational requirements of AlphaFold 3 (AF3) for large multimeric complexes.

1. **Computational Scalability:** The proposal requires modeling 6,000 sequences as hexamers. An average NRC protomer is ~850 amino acids; a hexamer totals ~5,100 amino acids. This pushes the limits of AF3's memory handling and requires substantial GPU time (estimated ~60,000 models if using 5 seeds for both monomer and hexamer modes). This scale typically requires a high-performance A100 cluster and months of processing time.
2. **Metric Measurability:** The parameters themselves are highly elegant and rely on extractable geometric or energetic data. Using FoldX for interface energy ($E_{if}$) and variance across seeds ($\sigma^2_{RV}$) effectively addresses the risk of AF3 "forced folding" by detecting structural frustration and instability. Parameter 5 ($\Delta D_{PM}$) leverages AF3's superior ligand-handling capabilities, which is a significant advantage over AF2-based methods.
3. **Ground Truth Alignment:** The idea utilizes the provided PDB structures (9FP6, 9RI9, 9CC8) effectively, specifically citing the transition mechanisms (closed-to-open) and domain orientations. Anchoring parameters to conserved motifs (P-loop, MHD, MADA) ensures that the index is robust to the high sequence diversity (insertions/deletions) found across 350 species.
4. **Data Limitations:** The "Missing Residue" problem in experimental structures (disordered N-termini) is mitigated by using AF3-generated canonical models as the baseline, as AF3 has been shown to model these $\alpha 1$ helices with high confidence (Abstract 1).

**Summary:** The PE-SNI is a technically sound framework that uses structural biology metrics rather than simple sequence identity. The primary feasibility constraint is the cost and time of generating 6,000 high-resolution hexameric models.

Answer: 5

**Impact potential:**

$\def\mathcal#1{\mathit{#1}}\def\mathscr#1{\mathit{#1}}$

##### Related Article Abstracts

1. **[1] RCSB PDB - 9FP6: Structure of the NbNRC2 hexameric resistosome** *Relevant for providing the active state structural baseline for NbNRC2, essential for parameters involving active resistosome geometry (interface angles, pore axial distances).*
2. **[10] Activation of plant immunity through conversion of a helper NLR homodimer into a resistosome** *Contains the structural data for the inactive resting state (PDB: 9RI9/8RFH). Confirms NRC2 rests as a dimer, not a monomer, which is vital for the SNI's "resting state" baseline parameters.*
3. **[16] A hydrophobic core in the coiled-coil domain is essential for NRC resistosome function** *Identifies a critical 4-residue hydrophobic core in the CC domain. Relevant for parameters evaluating the structural integrity and "competence" of the N-terminal executioner domain.*
4. **[14] An N-terminal motif in NLR immune receptors is functionally conserved across distantly related plant species** *Defines the MADA motif (Parameter 4) and its role in pore formation, providing the biological rationale for using radial variance as a signal for unconventionality.*
5. **[2] Analysing protein complexes in plant science: insights and limitation with AlphaFold 3** *Discusses AF3’s speed, accuracy in predicting plant immune complexes, and limitations in predicting mutation effects, justifying the high-throughput implementation protocol.*
6. **[9] A hierarchical immune receptor network in lettuce reveals contrasting patterns of evolution in sensor and helper NLRs** *Demonstrates that AF3 can distinguish helpers (which form resistosomes) from sensors (which fail to model as hexamers), validating the "Forced Hexamer" concept.*
7. **[4] Benchmarking AlphaFold for protein complex modeling reveals accuracy determinants** *Identifies interface energy and shape complementarity as strong predictors of model quality, supporting Parameter 3 (Interface Frustration Energy).*
8. **[13] A disease resistance protein triggers oligomerization of its NLR helper into a hexameric resistosome to mediate innate immunity** *Provides benchmarks for NRC2 vs ZAR1 (pentamer vs hexamer) and quantitative metrics (pTM, ipTM), relevant for calibrating SNI thresholds.*
9. **[15] Can AI modelling of protein structures distinguish between sensor and helper NLR immune receptors?** *Shows that helper NLRs consistently yield higher AF3 confidence scores in hexameric states compared to sensors, supporting the use of structural metrics to identify functional divergence.*
10. **[3] A disease resistance protein triggers oligomerization of its NLR helper into a hexameric resistosome to mediate innate immunity (bioRxiv version)** *Analyzes the 180° rotation required between resting and active states, providing the rationale for Parameter 5 (Ligand-Induced Switch Delta).*
11. **[6] How to assess the quality of AlphaFold 3 predictions | AlphaFold** *Explains how pLDDT and PAE matrix values indicate structural confidence, crucial for filtering out modeling artifacts in Parameter 7.*
12. **[18] A hierarchical immune receptor network in lettuce reveals contrasting patterns of evolution in sensor and helper NLRs (full bioRxiv text)** *Details the distinct patterns of diversification in different sensor subclades, justifying the focus on Solanaceae-specific divergence.*
13. **[12] A Suite of Designed Protein Cages Using Machine Learning Algorithms and Protein Fragment-Based Protocols** *Highlights the use of shape complementarity and interface solvation free energy as filters for oligomeric competence, supporting Parameter 1 and 3.*
14. **[19] Structural mechanism of heavy metal-associated integrated domain engineering of paired nucleotide-binding and leucine-rich repeat proteins in rice** *Discusses the mechanical switch from ADP to ATP binding in the NB-ARC domain, providing the biological mechanism for Parameter 5.*
15. **[8] A disease resistance protein triggers oligomerization of its NLR helper into a hexameric resistosome to mediate innate immunity (reprint)** *Further structural comparisons of NB domains being pushed outward in NbNRC2 relative to AtZAR1, relevant for Parameter 2.*

##### Detailed Assumptions

1. **Modeling Scalability:** The protocol assumes that modeling ~6,000 sequences as both monomers and "forced hexamers" (60,000 total models across 5 seeds) is computationally feasible within a reasonable research timeframe.
2. **AF3 Determinism/Signal:** It assumes that AlphaFold 3’s stochastic uncertainty (variance across seeds) reflects biological structural competence rather than random noise or hardware variability.
3. **Forced Assembly Signal:** It assumes that "forcing" unconventional proteins into a hexameric ring in AF3 will yield a detectable "stress" signal (high energy frustration or geometric distortion) rather than AF3 successfully "fixing" the structure into a plausible but incorrect hexamer.
4. **Residue-Specific Dynamics:** It assumes that measuring distance changes between specific motifs (e.g., P-loop to MHD) across different ligand-constrained runs (ADP vs. ATP) accurately captures the protein's mechanical switch potential.
5. **Motif-Based Anchoring:** The idea assumes that motifs like the P-loop, MHD, and MADA are identifiable across all 6,000 sequences to serve as anchors for geometric measurements.

##### Feasibility and Reasoning

1. **Scalability:** Modeling 6,000 hexamers is extremely intensive. Article [2] notes AF3 is faster and uses less memory than AF2, but modeling ~5,000 amino acids (6 x ~850aa) per run remains at the upper limit of current hardware. A tiered screening (monomer first) as suggested in the Review Guidelines would be more realistic.
2. **Structural Frustration:** Using FoldX or Rosetta to score AF3 interfaces (Parameter 3) is highly feasible and well-supported by Article [4], which shows interface binding energy correlates strongly with model quality. This is a robust way to detect proteins that "do not belong" in a hexamer.
3. **Variance as Signal:** Using seed variance (Parameter 4 and 7) is an innovative and feasible approach. Article [9] confirms that NRC sensors yield "low confidence with high predicted aligned errors" when modeled as hexamers. Quantifying this as variance effectively operationalizes AF3's "uncertainty."
4. **Nucleotide Switch:** Parameter 5 (Switch Delta) depends on ligand-constrained modeling. AF3 allows explicit ligand input (ADP/ATP) [2, 6]. Measuring the Euclidean shift between the Walker A and MHD histidine is a standard way to quantify STAND protein mechanical work, making this parameter biologically sound.
5. **MADA Disorder:** A potential feasibility issue exists for Parameters 4 and 7 regarding the MADA motif. Experimental structures (9FP6, 9CC8) often lack N-terminal coordinates due to disorder [1, 10]. However, AF3 can model these helices with high confidence [1], so using AF3 models of the canonical set as the "ground truth" baseline solves this data gap.

##### Suggested Improvements

1. **Incorporate AF3 Confidence Metrics:** The SNI should explicitly include **ipTM** and **pTM** as weights. A high SNI (novelty) associated with low ipTM should be penalized, as it likely indicates a modeling failure rather than genuine biological novelty [Guideline 4].
2. **Add a "CC-Hydrophobic Core" Metric:** Article [16] highlights a newly discovered hydrophobic core (L34, A72, I114, V118 in NRC4) essential for resistosome function. Adding a parameter for the "Planarity/Compactness of the CC-Core" would enhance the detection of unconventional executioners.
3. **Tiered Screening Approach:** To address computational costs, implement a "Monomer-SNI" (mSNI) using Parameters 1, 2, 5, and 7 to filter the 6,000 sequences. Only the top ~10% should proceed to the "Complex-SNI" (cSNI) involving forced hexamers and interface energetics [Guideline 2].
4. **Stoichiometric Contrast:** Instead of only forcing a hexamer, model a subset of candidates as pentamers (like AtZAR1 [3]) and compare the SNI. This would help identify if an unconventional NLR is simply a pentameric variant.

##### Overall Impact Potential

The **Physico-Evolutionary Structural Novelty Index (PE-SNI)** has **high impact potential**. It moves the field of NLR biology beyond simple sequence-based phylogeny into **functional structural informatics**.

- **Feasibility:** The use of AF3, FoldX, and Python-based structural parsing is technically sound and reproducible.
- **Scope:** The application to 6,000 sequences across the Solanaceae family addresses a major "dark matter" problem in plant immunity—identifying which NLRs are signaling decoys (sensors) vs. executors (helpers).
- **nuance:** Unlike a simple RMSD check, the PE-SNI evaluates specific mechanistic features like nucleotide cleft solvation, resting tilt, and interface "frustration." This provides deep biological insight into *why* a protein is unconventional.
- **Long-term Implications:** This framework could be adapted to other protein families (e.g., mammalian NLRs or bacterial TIRs), establishing a new standard for defining "structural novelty" in the era of AI-driven structural biology.

The primary limitation is the extreme computational requirement for 6,000 hexamers, but this can be mitigated by the suggested tiered screening. Overall, the idea is a statistically rigorous, biologically grounded, and innovative approach to high-throughput structural discovery.

Answer: 8

References:

[1] [RCSB PDB - 9FP6: Structure of the NbNRC2 hexameric resistosome](https://www.rcsb.org/structure/9fp6)

[2] [Analysing protein complexes in plant science: insights and limitation with AlphaFold 3 - PMC](https://pmc.ncbi.nlm.nih.gov/articles/PMC12098255/)

[3] [A disease resistance protein triggers oligomerization of its NLR helper into a hexameric resistosome to mediate innate immunity](https://www.biorxiv.org/content/10.1101/2024.06.18.599586v1)

[4] [Benchmarking AlphaFold for protein complex modeling reveals accuracy determinants](https://www.biorxiv.org/content/10.1101/2021.10.23.465575v1)

[5] [Can AI modelling of protein structures distinguish between sensor and helper NLR immune receptors?](https://www.biorxiv.org/content/10.1101/2024.11.24.625045v1)

[6] [How to assess the quality of AlphaFold 3 predictions | AlphaFold](https://www.ebi.ac.uk/training/online/courses/alphafold/alphafold-3-and-alphafold-server/how-to-assess-the-quality-of-alphafold-3-predictions/)

[7] [In silico prediction method for plant Nucleotide‐binding leucine‐rich repeat‐ and pathogen effector interactions](https://www.ncbi.nlm.nih.gov/pmc/articles/PMC12042882/)

[8] [A disease resistance protein triggers oligomerization of its NLR helper into a hexameric resistosome to mediate innate immunity](https://www.ncbi.nlm.nih.gov/pmc/articles/PMC11540030/)

[9] [A hierarchical immune receptor network in lettuce reveals contrasting patterns of evolution in sensor and helper NLRs](https://www.biorxiv.org/content/10.1101/2025.02.25.639832v1)

[10] [Activation of plant immunity through conversion of a helper NLR homodimer into a resistosome - PMC](https://pmc.ncbi.nlm.nih.gov/articles/PMC11524475/)

[11] [A disease resistance protein triggers oligomerization of its NLR helper into a hexameric resistosome to mediate innate immunity](https://www.ncbi.nlm.nih.gov/pmc/articles/PMC11540030/)

[12] [A Suite of Designed Protein Cages Using Machine Learning Algorithms and Protein Fragment-Based Protocols](https://www.biorxiv.org/content/10.1101/2023.10.09.561468v1)

[13] [A disease resistance protein triggers oligomerization of its NLR helper into a hexameric resistosome to mediate innate immunity](https://www.biorxiv.org/content/10.1101/2024.06.18.599586v1)

[14] [An N-terminal motif in NLR immune receptors is functionally conserved across distantly related plant species | eLife](https://elifesciences.org/articles/49956)

[15] [Can AI modelling of protein structures distinguish between sensor and helper NLR immune receptors?](https://www.biorxiv.org/content/10.1101/2024.11.24.625045v1)

[16] [A hydrophobic core in the coiled-coil domain is essential for NRC resistosome function | bioRxiv](https://www.biorxiv.org/content/10.1101/2025.01.21.634219v3.full-text)

[17] [Biochemical basis of activation and inhibition of an NLR immune receptor network](https://ueaeprints.uea.ac.uk/id/eprint/93477/1/2023ContrerasMPhD.pdf)

[18] [A hierarchical immune receptor network in lettuce reveals contrasting patterns of evolution in sensor and helper NLRs | bioRxiv](https://www.biorxiv.org/content/10.1101/2025.02.25.639832v1.full-text)

[19] [Structural mechanism of heavy metal-associated integrated domain engineering of paired nucleotide-binding and leucine-rich repeat proteins in rice - PMC](https://pmc.ncbi.nlm.nih.gov/articles/PMC10338059/)

[20] [(PDF) A hierarchical immune receptor network in lettuce reveals contrasting patterns of evolution in sensor and helper NLRs](https://www.researchgate.net/publication/389403429_A_hierarchical_immune_receptor_network_in_lettuce_reveals_contrasting_patterns_of_evolution_in_sensor_and_helper_NLRs)

[21] [Sensor NLR immune proteins activate oligomerization of their NRC helpers in response to plant pathogens - PMC](https://pmc.ncbi.nlm.nih.gov/articles/PMC9975940/)

[22] [Assessing ATP binding and hydrolysis by NLR proteins - PMC](https://pmc.ncbi.nlm.nih.gov/articles/PMC4009365/)

[23] [Mechanism of Action of Plant Resistosome NRC4 - Lifeasible](https://www.lifeasible.com/blog/mechanism-of-action-of-plant-resistosome-nrc4/)

[24] [Structural basis of NLR activation and innate immune signalling in plants](https://www.ncbi.nlm.nih.gov/pmc/articles/PMC8813719/)

[25] [A helper NLR targets organellar membranes to trigger immunity](https://www.biorxiv.org/content/10.1101/2024.09.19.613839v1)

[26] [Structural basis of NLR activation and innate immune signalling in plants - PMC](https://pmc.ncbi.nlm.nih.gov/articles/PMC8813719/)

[27] [A root-specific NLR network mediates immune signaling of resistance genes against plant parasitic nematodes - PMC](https://pmc.ncbi.nlm.nih.gov/articles/PMC12236159/)

[28] [AlphaDesign: A de novo protein design framework based on AlphaFold](https://www.biorxiv.org/content/10.1101/2021.10.11.463937v1)

[29] [A helper NLR targets organellar membranes to trigger immunity](https://www.biorxiv.org/content/10.1101/2024.09.19.613839v1)

[30] [Diversification of the “EDVID” packing motif underpins structural and functional variation in plant NLR coiled-coil domains](https://www.biorxiv.org/content/10.1101/2025.06.01.657260v1)

[31] [Diversification of the “EDVID” packing motif underpins structural and functional variation in plant NLR coiled-coil domains](https://www.biorxiv.org/content/10.1101/2025.06.01.657260v1)

[32] [p K a Calculations of GPCRs: Understanding Protonation States in Receptor Activation](https://www.ncbi.nlm.nih.gov/pmc/articles/PMC11388449/)

[33] [An atypical NLR protein modulates the NRC immune receptor network in Nicotiana benthamiana](https://www.ncbi.nlm.nih.gov/pmc/articles/PMC9851556/)

[34] [In silico prediction method for plant Nucleotide‐binding leucine‐rich repeat‐ and pathogen effector interactions - PMC](https://pmc.ncbi.nlm.nih.gov/articles/PMC12042882/)

[35] [(PDF) Analysing protein complexes in plant science: insights and limitation with AlphaFold 3](https://www.researchgate.net/publication/391993456_Analysing_protein_complexes_in_plant_science_insights_and_limitation_with_AlphaFold_3)

[36] [Frontiers | NLRexpress—A bundle of machine learning motif predictors—Reveals motif stability underlying plant Nod-like receptors diversity](https://www.frontiersin.org/journals/plant-science/articles/10.3389/fpls.2022.975888/full)

**Motivation:**

$\def\mathcal#1{\mathit{#1}}\def\mathscr#1{\mathit{#1}}$

**Coherence:**

$\def\mathcal#1{\mathit{#1}}\def\mathscr#1{\mathit{#1}}$

**Deep verification:**

$\def\mathcal#1{\mathit{#1}}\def\mathscr#1{\mathit{#1}}$
