## Supplementary Data S6 for "AI-guided discovery of atypical protein assemblies": [EXTERNAL] ├░Y╠ê┬o┬1⁄4 Co-scientist - Hypothesis 230450355145970 - Defining a Structural Novelty Index (SNI) for Quantifying Divergence in Unconventional NRC-NLR Hexameric Resitosomes - For research purposes only - Google confidential.docx

### 🧬 Co-scientist - Defining a Structural Novelty Index (SNI) for Quantifying Divergence in Unconventional NRC-NLR Hexameric Resitosomes

**About**: *This is an experimental system for generating novel and testable hypotheses. The hypotheses are generated by a model and may be wrong. For research purposes only.*

### Proposal

$\def\mathcal#1{\mathit{#1}}\def\mathscr#1{\mathit{#1}}$

#### 1. The 8-Parameter Structural Novelty Index (SNI)

The Structural Novelty Index (SNI) is a multi-parametric framework that quantifies the architectural distance of any NRC sequence from the canonical hexameric baseline established by NbNRC2 (PDB ID: 9FP6) and SlNRC3 (PDB ID: 9RI9). The SNI is calculated by modeling a query sequence as a homohexamer in AlphaFold 3 and extracting eight specific geometric and energetic parameters. A high SNI score indicates a "Symmetry Breaker" or an unconventional architecture.

#### 2. The 8 Quantitative Parameters of the SNI

| # | Parameter Name | Unit | Structural Significance |
| --- | --- | --- | --- |
| **1** | **Azimuthal Protomer Rotation ($\theta_{Az}$)** | Degrees ($^\circ$) | Measures the angle between the centroids of adjacent NB-ARC domains. Canonical = $60^\circ$. Deviation indicates a shift to pentameric ($72^\circ$) or heptameric ($51^\circ$) states. |
| **2** | **Axial Stair-Step Displacement ($\Delta z$)** | Angstroms ($\AA$) | The vertical offset between adjacent protomers along the central axis. Canonical $\approx 0 \AA$ (flat ring). High values indicate a "spiral" or "open-ended" filament rather than a closed resistosome. |
| **3** | **MADA-Helix Tilt Angle ($\alpha_{M}$)** | Degrees ($^\circ$) | The angle of the N-terminal $\alpha$1-helix relative to the membrane normal. Canonical $\approx 15\text{--}25^\circ$. Deviations suggest a shielded or non-pore-forming N-terminus. |
| **4** | **NB-ARC "Flip" Vector ($\vec{V}_{flip}$)** | Normalized Ratio | Measures the distance between the NBD and WHD domains in the AF3 model compared to the closed (inactive) vs open (9FP6) state. Detects non-flipping activation mechanisms. |
| **5** | **Interface Shape Complementarity ($S_c$)** | 0 to 1 (Index) | Quantifies how well the surfaces of adjacent NB-ARC domains fit together. High $S_c$ in a hexamer model suggests a canonical fit; low $S_c$ suggests the sequence is physically incompatible with hexamerization. |
| **6** | **Pore Hydrophobic Density ($\rho_{H}$)** | $kcal/mol/\AA^2$ | The ratio of hydrophobic surface area to total surface area within the central 10$\AA$ of the pore. Distinguishes water-conducting pores from plugged or signaling-only variants. |
| **7** | **LRR-to-Core Radial Pitch ($R_{LRR}$)** | Angstroms ($\AA$) | The distance from the central pore axis to the center of mass of the LRR domain. Canonical NRCs have a tight crown; unconventional variants may have an expanded sensor ring. |
| **8** | **MHD-Motif Spatial Variance ($\delta_{MHD}$)** | Angstroms ($\AA$) | The coordinate deviation of the conserved MHD motif relative to the ATP-binding pocket. Identifies variants that have the sequence but lack the structural geometry for nucleotide exchange. |

#### 3. The Extraction Process (AF3 Workflow)

1. **Multimer Prediction:** Force the query NRC sequence into a homohexameric assembly in AlphaFold 3.
2. **Coordinate Extraction:** Use a Python script (utilizing Biopython or MDTraj) to define the centroids of the four sub-domains (CC, NBD, HD1, WHD).
3. **Vector Calculation:** Calculate the angles and distances defined above. For example, $\theta_{Az}$ is the angle between vectors $\vec{C1}$ and $\vec{C2}$ (where $C$ is the centroid of the NB-ARC domain of adjacent protomers).
4. **Z-Score Normalization:** Compare the results against the mean values of the Reference Set (PDB IDs: 9FP6, 9RI9, 9CC8).
5. **SNI Aggregation:** The final SNI is the Euclidean distance of the 8-parameter vector from the "Canonical Origin" (the mean of the ground truth dataset).

#### 4. Identification of Unconventional NLRs

The SNI allows for the identification of several classes of unconventional NLRs:

- **The Symmetry Breaker:** A sequence that yields a high Axial Stair-Step ($\Delta z$) but low Interface Complementarity ($S_c$) is likely a protein that cannot form a ring, potentially acting as a monomeric decoy or a dimeric regulator.
- **The Non-Pore Helper:** A sequence with a canonical NB-ARC structure but a high MADA Tilt Angle ($\alpha_{M}$) and low Pore Hydrophobic Density ($\rho_{H}$) is flagged as a helper that signals via protein-protein interaction rather than membrane lysis.
- **The Stoichiometry Shifter:** If the Azimuthal Rotation ($\theta_{Az}$) consistently clusters at $72^\circ$ across multiple AF3 seeds, the protein is identified as a potential pentameric NRC, representing a structural link between ZAR1 and NRC2.

By applying this SNI to the ~6,000 Solanaceae sequences, the study transitions from observing sequence data to predicting function based on structural geometry, prioritizing structurally divergent candidates for further research in plant immunity.

### Reviews summary

$\def\mathcal#1{\mathit{#1}}\def\mathscr#1{\mathit{#1}}$

Positive aspects:

- **Methodological Formalization of Structural Topology:** The idea moves beyond qualitative visual inspection or simple sequence homology to a rigorous, multi-parametric mathematical framework. By quantifying the "Euclidean distance" from a canonical structural origin, it allows for the objective identification of unconventional NLRs that fall into a "structural gray zone."
- **Biologically Validated Parameters:** The chosen metrics, such as the Azimuthal Protomer Rotation ($\theta_{Az}$) and NB-ARC "Flip" Vector, directly address known structural differences between different oligomeric states (e.g., hexamers vs. pentamers). Literature confirms that specific angular shifts in these domains are the primary drivers of stoichiometry and function.
- **Diagnostic Use of AlphaFold 3:** The "Force-Hexamer" approach is a novel and clever implementation of structural prediction. By forcing sequences into a specific assembly, the resulting structural strain (measured via axial displacement or interface complementarity) acts as a diagnostic tool to determine if a protein is physically incompatible with the canonical resistosome architecture.
- **Multidomain Functional Coverage:** Unlike simple RMSD checks, the SNI parameters span the entire protein architecture, including the pore-forming N-terminus (MADA tilt), the activation-governing NB-ARC domain (MHD variance), and the sensor-regulating LRR ring (radial pitch). This ensures the index captures functional nuance rather than just global shape.
- **High-Throughput Automation Potential:** The parameters are derived from standard atomic coordinates, making them easily extractable via Python-based bioinformatics scripts. This allows the workflow to scale to thousands of sequences, providing a systematic way to triage the massive NRC family dataset.

Negative aspects:

- **Prohibitive Computational Requirements:** Running AlphaFold 3 predictions for 6,000 sequences as homohexamers represents a massive computational bottleneck. Such a task requires immense GPU resources and time, making the goal of screening the entire Solanaceae dataset difficult to achieve without significant institutional infrastructure.
- **Omission of Model Confidence Metrics:** The index lacks the integration of AF3-specific confidence scores like ipTM (interface predicted Template Modeling score). Without weighting parameters by prediction confidence, the index risks flagging "structural novelty" that is actually just an artifact of a low-confidence or failed AF3 model.
- **Sensitivity to Structural Noise in Flexible Regions:** Several parameters, such as the MADA-Helix Tilt and Pore Hydrophobic Density, rely on the N-terminal $\alpha$1-helix. This region is notoriously flexible and dynamic; even with AF3, the inherent noise in modeling these "death switches" could lead to inconsistent SNI scores.
- **Lack of Environmental Context in Modeling:** The current parameters do not mandate the inclusion of essential ligands like ATP/ADP or membrane lipids in the AF3 input. Recent studies indicate these molecules are critical for stabilizing the active resistosome, and their absence could result in misleading structural geometries.
- **Absence of Resting State Comparison:** The SNI focuses exclusively on the activated hexamer. It does not account for the stability of the resting homodimer interface, which is a critical checkpoint in NRC activation. A sequence might appear unconventional in a hexamer simply because its primary biological deviation occurs at the dimerization stage.

#### Research contacts

$\def\mathcal#1{\mathit{#1}}\def\mathscr#1{\mathit{#1}}$

Based on the provided research articles, the following researchers are suggested to review the hypothesis regarding the Structural Novelty Index (SNI). They are experts in NRC resistosome structures, the application of AlphaFold 3 (AF3) for NLR modeling, and the evolution of the NRC network.

#### 1. Jogi Madhuprakash

- **Justification:** Madhuprakash is a key researcher in solving the structures of NRC resistosomes and refining them using computational tools. His work directly involves the "Reference Set" mentioned in the goal (NbNRC2, PDB ID: 9FP6) and involves detailed refinement protocols that align with the SNI's "Extraction Process."
- **Relevant Articles:** *Structure of the NbNRC2 hexameric resistosome* (Abstract 1); *A plant pathogen effector blocks stepwise assembly of a helper NLR resistosome* (Abstract 8).

#### 2. AmirAli Toghani

- **Justification:** Toghani has published on the use of AlphaFold 3 to differentiate between helper and sensor NLRs by predicting resistosome-like structures. His expertise in leveraging AF3 metrics to distinguish between different oligomeric states (hexamers vs. pentamers) makes him an ideal reviewer for the "Symmetry Breaker" and "Stoichiometry Shifter" aspects of the proposed SNI.
- **Relevant Articles:** *A hierarchical immune receptor network in lettuce reveals contrasting patterns of evolution in sensor and helper NLRs* (Abstract 3/6); *Activation of plant immunity through conversion of a helper NLR homodimer into a resistosome* (Abstract 2).

#### 3. Sophien Kamoun

- **Justification:** As a senior author across multiple provided abstracts (1, 7, 8, 11, 13), Kamoun has led the field in establishing the NRC network paradigm. His research provides the foundational biological context for distinguishing unconventional NRC-NLRs from canonical ones, specifically regarding helper NLR (NRC-H) and sensor NLR (NRC-S) diversification.
- **Relevant Articles:** *Structure of the NbNRC2 hexameric resistosome* (Abstract 1); *A plant pathogen effector blocks stepwise assembly of a helper NLR resistosome* (Abstract 7); *An atypical NLR protein modulates the NRC immune receptor network in Nicotiana benthamiana* (Abstract 10).

#### 4. Chih-Hang Wu

- **Justification:** Wu focuses on the structural and sequence-based divergence between NRC helpers and sensors, specifically identifying motifs (like the hydrophobic core) that are essential for function. His work is highly relevant to the "N-terminal CC-domain variations" and "NB-ARC domain conservation" parameters of the SNI.
- **Relevant Articles:** *A hydrophobic core in the coiled-coil domain is essential for NRC resistosome function* (Abstract 9); *Activation of plant immunity through conversion of a helper NLR homodimer into a resistosome* (Abstract 2).

#### 5. B.J. Staskawicz

- **Justification:** Staskawicz is an author on the study identifying the hexameric state of the NRC4 resistosome (PDB ID: 9CC8). Since NRC4 is one of the essential components of the "Reference Set" for the SNI, his expertise in the structural biology of NRC4 is vital for validating the parameters used to define canonical versus unconventional architectures.
- **Relevant Article:** *Hexameric state of the NRC4 resistosome* (Abstract 4).

### Appendix:

**All reviews:**

**Correctness:**

$\def\mathcal#1{\mathit{#1}}\def\mathscr#1{\mathit{#1}}$

This review evaluates the proposed **8-Parameter Structural Novelty Index (SNI)** based on the provided ground truth structures and current structural biology research on the NRC immune receptor network.

#### 1. Related Article Abstracts

1. **[1] RCSB PDB - 9FP6: Structure of the NbNRC2 hexameric resistosome:** Essential for establishing the canonical hexameric baseline for NRC2.
2. **[4] RCSB PDB - 9CC8: Hexameric state of the NRC4 resistosome:** Provides the structural coordinates for the NRC4 canonical baseline.
3. **[8] A disease resistance protein triggers oligomerization... (Madhuprakash et al. 2024):** Specifically compares NRC2 (hexamer) to ZAR1 (pentamer) using angular measurements and pore diameters, validating the use of geometric metrics.
4. **[11] AlphaFold 3 prediction of CC-NLR resistosomes... (Madhuprakash et al. 2024):** Demonstrates that AF3 can accurately predict NRC hexamer stoichiometry and N-terminal funnels, supporting the AF3-based extraction workflow.
5. **[2] Activation of plant immunity through conversion of a helper NLR homodimer into a resistosome (Selvaraj et al. 2024):** Details the resting state (homodimer) and interface residues, relevant for the "NB-ARC Flip" and "Interface Shape Complementarity" parameters.
6. **[15] An N-terminal motif in NLR immune receptors is functionally conserved... (Adachi et al. 2019):** Defines the MADA motif and notes its absence in NRC-S, supporting "MADA-Helix Tilt" as a key parameter.
7. **[10] An atypical NLR protein modulates the NRC immune receptor network... (Adachi et al. 2023):** Describes NRCX, an "unconventional" NRC that lacks a functional MADA motif, justifying the need for an SNI to identify such variants.
8. **[9] A hydrophobic core in the coiled-coil domain is essential for NRC resistosome function (Wang et al. 2025):** Identifies specific hydrophobic residues in the CC domain that are essential for function but diverged in sensors.
9. **[3] A hierarchical immune receptor network in lettuce... (Pai et al. 2025):** Confirms that AF3 can differentiate between helpers and sensors by failing to form resistosomes for the latter, supporting the use of $S_c$ and oligomerization metrics.
10. **[14] A disease resistance protein triggers oligomerization... (Madhuprakash et al. 2024 - bioRxiv):** Details the 10° interdomain angle difference between NRC2 and ZAR1, providing direct evidence for the "NB-ARC Flip/Orientation" metrics.

#### 2. Detailed Assumptions

1. **Hexameric Compatibility:** Forcing a sequence into a homohexamer in AF3 will yield measurable structural strain or geometric deviation if the sequence is naturally monomeric, dimeric, or pentameric.
2. **Geometric Representative Power:** Geometric parameters like $\theta_{Az}$ and $\Delta z$ are sufficient to capture the "stoichiometry shift" or "filamentous" nature of unconventional NLRs.
3. **Accuracy of AF3 N-termini:** AF3 can model the flexible N-terminal $\alpha$1-helix with enough precision to calculate a meaningful "Tilt Angle" ($\alpha_{M}$).
4. **Baseline Stability:** The "Reference Set" (NRC2, 3, 4) exhibits sufficiently low variance across these 8 parameters to establish a tight "Canonical Origin" for Z-score normalization.
5. **Functional Correlation:** Structural "divergence" (high SNI) correlates with functional "unconventionality" (e.g., decoy, regulator, or non-pore-forming helper).

#### 3. Comparison with Knowledge Base & Reasoning

- **Azimuthal Rotation ($\theta_{Az}$):** **Correct.** Abstract [8] and [14] explicitly use angular comparisons between NRC2 and ZAR1 to explain why one is a hexamer and the other a pentamer. A shift from $60^\circ$ to $72^\circ$ is a mathematically sound way to detect pentameric tendencies.
- **MADA-Helix Tilt ($\alpha_{M}$):** **Correct.** Abstract [15] and [11] emphasize the importance of the $\alpha$1-helix funnel. Abstract [13] notes that including lipids (oleic acid) in AF3 allows high-confidence modeling of these funnels. This parameter captures the "death switch" capability.
- **Interface Shape Complementarity ($S_c$):** **Correct.** Abstract [3] shows that sensor NLRs (NRC-S) fail to form resistosome-like structures in AF3. Low $S_c$ would be the quantitative manifestation of this failure.
- **MHD-Motif Spatial Variance ($\delta_{MHD}$):** **Correct.** The MHD motif is a known molecular switch [2, 12]. If the sequence exists but the spatial orientation is off, the nucleotide exchange mechanism is likely broken.
- **NB-ARC "Flip" Vector ($\vec{V}_{flip}$):** **Correct.** Abstract [14] mentions a 10° difference in interdomain angles between NRC2 and ZAR1. This confirms that the internal arrangement of NB-ARC subdomains is a key differentiator of stoichiometry.

#### 4. Strength of Evidence

- **Direct Evidence:** Extremely high. The idea uses the exact PDB IDs requested (9FP6, 9RI9, 9CC8). Recent literature [8, 11, 14] already uses "interdomain angles," "pore diameter," and "AF3 modeling confidence" to classify these proteins. The idea formalizes these observations into a unified index.
- **Indirect Evidence:** High. The existence of "atypical" NRCs like NRCX [10] and the specialization of sensor vs. helper clades [6, 15] provides a clear biological rationale for why an SNI is needed to sort the 6,000 sequences.

#### 5. Suggested Improvements

1. **Include AF3 Confidence Metrics:** Integrate **ipTM** (interface predicted Template Modeling score) as a 9th parameter. Abstract [5, 11] states that ipTM > 0.8 indicates a confident interaction. A low ipTM in a forced hexamer is a primary indicator of structural novelty (non-hexamerization).
2. **Membrane Context:** Specify that AF3 runs should include **oleic acid or phospholipids** (as suggested in [3, 11, 13]) to stabilize the N-terminal funnel for accurate $\alpha_{M}$ calculation.
3. **Define Centroids:** Specifically define the NB-ARC centroid as the center of mass of the NBD, HD1, and WHD sub-domains collectively to ensure $\theta_{Az}$ is robust against small local fluctuations.

#### 6. Goal Requirements Assessment

- **5-10 Parameters:** **Yes** (8).
- **Measurable from PDB/AF3:** **Yes**, all are geometric or energetic.
- **Distinguish unconventional from canonical NRC2/3/4:** **Yes**, by using them as the "Canonical Origin."
- **Incorporate Ground Truth (9FP6, 9RI9, 9CC8):** **Yes**, used for Z-score normalization.
- **Focus on Protomer Angles/CC variations:** **Yes** (Parameters 1, 3, 4, 7).
- **Applicable to 6,000 sequences:** **Yes**, the AF3-to-Python workflow is highly automatable.

#### 7. Reasoning about Correctness

The idea is structurally and biologically sound. It moves beyond simple sequence homology (which fails to distinguish NRC-H from NRC-S effectively in large datasets) to **topological homology**. By "forcing" the hexameric state, the index essentially tests the **potential energy surface** and **geometric compatibility** of a sequence with the known active state of the family. The parameters chosen (rotation, tilt, complementarity) are standard in structural bioinformatics and directly address the differences identified in recent cryo-EM papers (e.g., NRC2's wider NB-ARC angle compared to ZAR1).

#### Conclusion and Recommendation

The idea is excellent, mathematically grounded, and directly addresses the goal using the latest structural biology tools (AF3). It provides a scalable solution to triage thousands of NLRs.

**Recommendation:** **Test the idea.** The workflow can be immediately implemented using the AlphaFold 3 API and Biopython.

Answer: 9

**Novelty:**

$\def\mathcal#1{\mathit{#1}}\def\mathscr#1{\mathit{#1}}$

#### 1. Related Article Abstract Titles

1. **[1] Structure of the NbNRC2 hexameric resistosome (9FP6)**: Establishes the primary canonical structural baseline for the SNI.
2. **[8/14] A disease resistance protein triggers oligomerization of its NLR helper into a hexameric resistosome to mediate innate immunity**: Provides specific geometric comparisons (angles/pore sizes) between hexameric (NRC2) and pentameric (ZAR1) states.
3. **[11/13] AlphaFold 3 prediction of CC-NLR resistosomes reveals a diversity of N-terminal pore-like structures**: Demonstrates the existing workflow of using AF3 to predict and classify NLR oligomers.
4. **[3/6] A hierarchical immune receptor network in lettuce reveals contrasting patterns of evolution in sensor and helper NLRs**: Uses AF3 metrics (ipTM, pLDDT) to distinguish NRC helpers from sensors based on their ability to form resistosomes.
5. **[2] Activation of plant immunity through conversion of a helper NLR homodimer into a resistosome**: Identifies specific interface residues and dimerization mechanics that define the resting vs. active state.
6. **[15] An N-terminal motif in NLR immune receptors is functionally conserved across distantly related plant species**: Defines the MADA motif (α1-helix) which is central to the SNI parameters for pore formation and tilt.
7. **[9] A hydrophobic core in the coiled-coil domain is essential for NRC resistosome function**: Highlights the importance of CC-domain hydrophobic residues, supporting SNI parameters like Pore Hydrophobic Density.
8. **[10] An atypical NLR protein modulates the NRC immune receptor network in Nicotiana benthamiana**: Describes NRCX, an "atypical" NRC that would be the primary target for identification via a high SNI score.
9. **[4] Hexameric state of the NRC4 resistosome (9CC8)**: Provides the third ground truth structure for the canonical reference set.
10. **[5] How to assess the quality of AlphaFold 3 predictions**: Details the standard metrics (ipTM, pTM, PAE) that the SNI seeks to augment with more specific structural geometry.

#### 2. Aspects of the idea that were already tried

- **AlphaFold 3 for NLR stoichiometry prediction**: The use of AF3 to model pentameric vs. hexameric resistosomes is well-established [11, 13]. Experts already compare ipTM and pTM values for different oligomeric states to determine the most likely configuration.
- **Geometric comparisons of resistosomes**: Measuring the "Azimuthal Protomer Rotation" ($\theta_{Az}$) and "Interdomain Angles" is explicitly described in [8] and [14]. Specifically, the 10° difference in interdomain angles between NbNRC2 (hexamer) and AtZAR1 (pentamer) and the resulting change in pore diameter (12 Å vs. 17–19 Å) have been measured and published.
- **MADA-helix analysis**: The identification and functional validation of the MADA motif (Parameter 3 and 6) are established [15]. The "funnel-shaped structure" and its insertion into the membrane are standard concepts in the resistosome model [17].
- **Distinguishing Helpers from Sensors via AF3**: Recent work in lettuce [3, 6] already uses AF3 modeling of hexamers to triage sequences into "helpers" (which form resistosomes) and "sensors" (which do not).
- **Interface Residue Mapping**: Using structural models to identify residues at the NB-LRR interface and their conservation patterns is a core component of recent NRC research [2].

#### 3. Aspects of the idea which are novel

- **The Structural Novelty Index (SNI) Framework**: While structural metrics (angles, distances) are used in individual papers, the formalization of these into a single, multi-parametric, quantitative index (a weighted 8-parameter vector) is novel.
- **High-Throughput Geometric Triage**: Current workflows rely heavily on AF3 confidence scores (ipTM/pTM) or visual inspection [3, 11]. The SNI proposes a transition to automated "structural topology" screening, where the *physical deviation* from the canonical origin is the primary classifier, rather than just prediction confidence.
- **The "Force-Hexamer" Diagnostic**: The methodology of deliberately forcing 6,000 sequences into a homohexameric assembly to observe the resulting "strain" or "symmetry breaking" in the azimuthal rotation ($\theta_{Az}$) or axial displacement ($\Delta z$) as a diagnostic tool for stoichiometry is a novel implementation of AF3.
- **Specific Parameters**: Parameters like **Axial Stair-Step Displacement ($\Delta z$)** to detect spiral filaments and **MHD-Motif Spatial Variance ($\delta_{MHD}$)** as a structural (rather than sequence) metric for activation readiness represent more nuanced metrics than currently reported.

#### 4. Novelty review

The idea is **moderately novel** but leans heavily on established structural biology practices.

The core of the idea—using specific angles and distances to describe NLR activation—is exactly how the "hexamer vs. pentamer" discovery was made in the first place [8, 14]. Experts in the field are already using AlphaFold 3 to predict NRC structures and are aware that the stoichiometry is linked to inter-domain angles [13].

However, the idea's merit lies in its **methodological formalization**. Most current research uses AF3 as a *confirmation* tool or a *qualitative* triage tool (e.g., "it forms a ring" vs. "it doesn't"). Proposing a standardized, mathematical "Euclidean distance from Canonical Origin" allows for the identification of sequences that fall into a "structural gray zone"—atypical NLRs that might form dimers, filaments, or non-pore-forming hexamers. This is a logical and useful evolution of the field's current toolkit.

#### 5. Reasoning about novelty and recommendation

The idea is not "groundbreaking" because it uses known parameters (angles, tilt, hydrophobicity) and existing tools (AF3). However, it is **highly practical** and significantly advances the efficiency of screening the vast NRC family (~6,000 sequences).

The recommendation is to **test the idea**. The field is currently flooded with sequence data, and simple pLDDT/ipTM metrics from AF3 often fail to capture the functional nuances of "atypical" NLRs like NRCX [10]. A formal SNI would allow researchers to systematically identify "Symmetry Breakers" and "Non-Pore Helpers" that would otherwise be missed or dismissed as "poor AF3 models."

The parameters selected are biologically relevant and leverage the ground truth structures (9FP6, 9RI9, 9CC8) effectively. Specifically, using $\Delta z$ and $\theta_{Az}$ to characterize non-closed assemblies is a robust way to find unconventional immune regulators.

Answer: 5

**Feasibility:**

$\def\mathcal#1{\mathit{#1}}\def\mathscr#1{\mathit{#1}}$

#### Related Article Abstracts

1. **[1] Structure of the NbNRC2 hexameric resistosome (PDB: 9FP6)**: Provides the primary "canonical" ground truth for the NB-ARC interface and hexameric geometry used to define the SNI baseline.
2. **[2] Activation of plant immunity through conversion of a helper NLR homodimer into a resistosome**: Details the transition from a resting homodimer to an active hexamer, crucial for measuring the "NB-ARC Flip" and "Interface Shape Complementarity" parameters.
3. **[3] A hierarchical immune receptor network in lettuce reveals contrasting patterns of evolution in sensor and helper NLRs**: Confirms that AlphaFold 3 (AF3) can distinguish helpers from sensors by whether they successfully form resistosomes, validating the core modeling strategy.
4. **[8/14] A disease resistance protein triggers oligomerization... hexameric resistosome (Madhuprakash et al. 2024)**: Directly compares pentameric (ZAR1) vs. hexameric (NRC2) geometry, providing specific angle/pore width values used to define "Azimuthal Rotation" and "Pore Hydrophobic Density."
5. **[11] AlphaFold 3 prediction of CC-NLR resistosomes reveals a diversity of N-terminal pore-like structures**: Demonstrates that forcing hexameric configurations in AF3 yields high-confidence models for triage, supporting the feasibility of the extraction workflow.
6. **[15] An N-terminal motif in NLR immune receptors is functionally conserved... (MADA motif)**: Justifies the use of "MADA-Helix Tilt Angle" and "Pore Hydrophobic Density" as functional indicators of pore formation.
7. **[9] A hydrophobic core in the coiled-coil domain is essential for NRC resistosome function**: Highlights internal structural requirements in the CC domain, relevant for the "MADA-Helix Tilt" and "Radial Pitch" parameters.
8. **[5] How to assess the quality of AlphaFold 3 predictions**: Details PAE and ipTM metrics, which are essential for filtering out noise from the SNI parameters when processing low-confidence models.
9. **[4] Hexameric state of the NRC4 resistosome (PDB: 9CC8)**: Provides a secondary ground truth dataset for NRC4, ensuring the SNI is not overfitted to NRC2/3.
10. **[13] A disease resistance protein triggers oligomerization... (Madhuprakash et al. 2024)**: Discusses the use of AF3 as a platform for hypothesis generation and NLR classification, reinforcing the feasibility of structural triage.

#### Steps to Test the Idea

1. **Pilot Validation (Go/No-Go):** Select a "Gold Standard" subset: 20 canonical helpers (NRC2/3/4), 20 sensors (NRC-S), and 5 known atypical members (e.g., NRCX). Run AF3 to model these as homohexamers.
   - *Go Criteria:* The SNI parameters must clearly distinguish canonical helpers (low SNI) from sensors/atypical members (high SNI).
   - *No-Go Criteria:* Parameters show high variance within the ground truth set or fail to separate sensors from helpers.
2. **Script Development:** Develop a Python-based geometric extraction pipeline using Biopython (for centroids and angles), MDTraj (for distances), and FreeSASA (for hydrophobic surface area/density).
3. **Reference Calibration:** Use PDB IDs 9FP6, 9RI9, and 9CC8 to define the "Canonical Origin." Calculate the mean and standard deviation for each of the 8 parameters to allow for Z-score normalization.
4. **High-Throughput Modeling:** Run AF3 for the full ~6,000 sequence Solanaceae dataset. Given the computational load, this would likely be performed in batches on a high-performance computing (HPC) cluster or cloud environment (e.g., A100 GPUs).
5. **SNI Calculation & Analysis:** Apply the script to all 6,000 models. Use Euclidean distance to aggregate the 8 parameters into a single SNI score per sequence.
6. **Outlier Prioritization:** Cluster sequences by SNI score. Specifically, identify "Symmetry Breakers" ($\Delta z$ outliers) and "Non-Pore Helpers" ($\alpha_M$/$\rho_H$ outliers) for wet-lab validation (cell death assays).

#### Reasoning about Feasibility

The feasibility of testing the SNI is **moderately complex to resource-intensive** depending on the scale.

- **Computational Scalability:** The parameters themselves (angles, distances, Z-offsets) are mathematically simple and can be extracted in seconds using standard bioinformatics libraries. However, the requirement to run AF3 predictions for **6,000 sequences as homohexamers** is a massive computational bottleneck. Abstract [3] and [11] show that researchers are already doing this for smaller subsets (10–50 sequences), but scaling this by a factor of 100 requires significant GPU resources and time (estimated 2–3 months).
- **Metric Measurability:** Most parameters (1, 2, 3, 4, 7, 8) rely on standard atomic coordinates, making them highly feasible. Parameter 5 (Interface Shape Complementarity) and Parameter 6 (Pore Hydrophobic Density) are slightly more complex, requiring specialized scripts or third-party tools, but are still standard in structural biology.
- **Reference Alignment:** The availability of high-resolution PDBs for the NRC2, NRC3, and NRC4 resistosomes (Abstracts [1], [4], [7]) provides an excellent, well-defined baseline for "Canonical" states.
- **Reliability:** Abstract [3] explicitly notes that AF3 successfully models resistosome-like oligomers for helpers but not for sensors, suggesting the SNI will have a high signal-to-noise ratio.

While the "Pilot Study" to prove the index works is highly feasible (Score 7), the "Goal" of identifying unconventional NLRs across the **entire 6,000-sequence dataset** aligns with the quantitative benchmarks for a Score 5.

Answer: 5

**Impact potential:**

$\def\mathcal#1{\mathit{#1}}\def\mathscr#1{\mathit{#1}}$

#### 1. Related Article Abstracts

The following articles are most relevant for assessing the impact and feasibility of the Structural Novelty Index (SNI):

1. **[1] 9FP6: Structure of the NbNRC2 hexameric resistosome**: Provides the primary ground truth for a canonical activated hexamer, including domain coordinates essential for setting the "Canonical Origin" of the index.
2. **[8] A disease resistance protein triggers oligomerization... (Madhuprakash et al. 2024)**: This paper explicitly compares NbNRC2 (hexamer) to AtZAR1 (pentamer), identifying specific quantitative differences like a 10° increase in interdomain angles and pore diameter (17-19 Å). This validates the choice of angular and pore-based parameters.
3. **[11] AlphaFold 3 prediction of CC-NLR resistosomes...**: Confirms that AF3 can accurately model stoichiometry and N-terminal pores, and suggests that AF3 can be used to "triage" NLRs into functional categories—directly supporting the idea's methodology.
4. **[2] Activation of plant immunity through conversion of a helper NLR homodimer into a resistosome**: Describes the resting dimer interface. This is relevant for identifying "Symmetry Breakers" that might deviate from the dimer-to-hexamer transition.
5. **[10] An atypical NLR protein modulates the NRC immune receptor network...**: Describes NRCX, a real-world "unconventional" NRC that lacks canonical motifs. This serves as a primary biological target that the SNI should be able to identify.
6. **[3] A hierarchical immune receptor network in lettuce...**: Demonstrates that AF3 fails to form resistosome structures for sensors (NRC-S) but succeeds for helpers (NRC-H), validating the use of AF3 to detect "deviation" from canonical assembly.
7. **[9] A hydrophobic core in the coiled-coil domain is essential for NRC resistosome function**: Identifies structural features in the CC domain (α2-α4) necessary for function. This supports the inclusion of parameters focusing on CC-domain variations.
8. **[15] An N-terminal motif in NLR immune receptors is functionally conserved...**: Establishes the MADA motif as a functional baseline for helpers. Variations in this (captured by SNI parameter 3 and 6) are critical for functional prediction.
9. **[13] A disease resistance protein triggers... (PMC11540030)**: Discusses how AF3 can fill gaps in experimental structures, specifically for the N-terminal α1-helices, supporting the feasibility of high-confidence tilt angle measurements.
10. **[5] How to assess the quality of AlphaFold 3 predictions**: Provides the technical basis (pLDDT, ipTM, PAE) for filtering the AF3 models before they are processed by the SNI.

#### 2. Detailed Assumptions

To achieve its potential impact, the idea depends on the following assumptions:

1. **AF3 Structural Consistency:** AlphaFold 3 must produce consistent, high-confidence models when "forced" into a homohexameric state, even for sequences that might naturally be monomers or dimers.
2. **Topology over Sequence:** Structural divergence (e.g., a 10° shift in NB-ARC angle) must be a more reliable predictor of "unconventional" function than simple sequence identity (percent homology).
3. **Interpretability of "Failure":** When a sequence cannot form a hexamer, AF3 must produce a model where the SNI parameters (like Interface Shape Complementarity or Axial Displacement) diverge in a predictable way rather than resulting in random atomic clashes.
4. **Functional Correlation:** The 8 selected parameters must cover the minimal set of structural features that dictate the mechanism of action (pore formation vs. signaling scaffold).

#### 3. Feasibility and Reasoning

- **Feasibility of AF3 Screening:** Highly realistic. Abstract **[11]** and **[13]** explicitly state that AF3 successfully models NbNRC2 and can distinguish between hexameric and pentameric configurations. Using AF3 as a "platform for hypothesis generation and NLR classification" is already being explored in current research.
- **Feasibility of Quantitative Parameters:** The parameters chosen (Azimuthal rotation, Stair-step displacement, Tilt angles) are standard geometric metrics in structural biology. Abstract **[8]** specifically notes that a 10° difference in the NB-ARC interdomain angle is what distinguishes hexamers from pentamers, proving these metrics are biologically meaningful.
- **Effect on Impact:** The transition from sequence-based to structure-based triage is transformative. Simple RMSD (Root-Mean-Square Deviation) often fails to highlight specific functional changes (like the α4-helix bend mentioned in **[14]**). By using Z-score normalization against ground truth (**[1]**, **[4]**), the SNI provides a more nuanced way to identify proteins like NRCX (**[10]**) which might look like a helper by sequence but act as a regulator.
- **Limitation:** The "Pore Hydrophobic Density" and "MADA-Helix Tilt" assume that the N-terminal funnel is resolved. While Abstract **[13]** suggests AF3 is good at this, experimental structures often show high flexibility here, potentially introducing noise into the SNI.

#### 4. Suggested Improvements

1. **Confidence-Weighting:** Integrate AF3 confidence scores (pLDDT and PAE) into the SNI aggregation. If a specific interface has low ipTM (**[5]**), the Interface Shape Complementarity ($S_c$) score should be weighted less to avoid false positives.
2. **Dimer-Interface Check:** Add a 9th parameter that assesses the stability of the resting homodimer interface described in **[2]**. This would distinguish between NRCs that are "unconventional" because they never dimerize and those that dimerize but fail to hexamerize.
3. **Ligand/Ion Context:** Abstract **[5]** and **[13]** suggest that adding lipids (oleic acid) or nucleotides (ATP/ADP) improves AF3 accuracy for NLRs. The SNI process should mandate the inclusion of these small molecules in the AF3 input to ensure the models reflect the activated state.

#### 5. Overall Impact Potential

The Structural Novelty Index (SNI) has **High Impact** potential.

- **Feasibility:** The methodology leverages proven tools (AF3) and targets (Solanaceae NRCs). The parameters are grounded in specific structural differences already observed between pentameric (ZAR1) and hexameric (NRC2) resistosomes.
- **Scope:** Applying this to ~6,000 sequences allows for the discovery of entirely new classes of plant immune regulators. It moves the field beyond "helper vs. sensor" into a spectrum of structural archetypes.
- **Long-term Implications:** This framework could be adapted for any STAND superfamily proteins, not just plant NLRs. It establishes a workflow for "Structural Bioinformatics" where the 3D model, not the sequence, is the primary unit of search.
- **Addressing the Problem:** It solves the "triage problem" in NLR research. Instead of testing 6,000 proteins in planta, researchers can prioritize the top 1% of "Symmetry Breakers" or "Non-Pore Helpers," drastically accelerating the discovery of novel immune mechanisms like those of NRCX.

The idea is realistic, utilizes the ground truth data effectively, and provides significantly more nuanced insight than simple sequence alignments or global RMSD checks.

Answer: 8

References:

[1] [RCSB PDB - 9FP6: Structure of the NbNRC2 hexameric resistosome](https://www.rcsb.org/structure/9fp6)

[2] [Activation of plant immunity through conversion of a helper NLR homodimer into a resistosome - PMC](https://pmc.ncbi.nlm.nih.gov/articles/PMC11524475/)

[3] [A hierarchical immune receptor network in lettuce reveals contrasting patterns of evolution in sensor and helper NLRs](https://www.biorxiv.org/content/10.1101/2025.02.25.639832v1)

[4] [RCSB PDB - 9CC8: Hexameric state of the NRC4 resistosome](https://www.rcsb.org/structure/9cc8)

[5] [How to assess the quality of AlphaFold 3 predictions | AlphaFold](https://www.ebi.ac.uk/training/online/courses/alphafold/alphafold-3-and-alphafold-server/how-to-assess-the-quality-of-alphafold-3-predictions/)

[6] [A hierarchical immune receptor network in lettuce reveals contrasting patterns of evolution in sensor and helper NLRs | bioRxiv](https://www.biorxiv.org/content/10.1101/2025.02.25.639832v1.full-text)

[7] [A plant pathogen effector blocks stepwise assembly of a helper NLR resistosome](https://www.biorxiv.org/content/10.1101/2025.07.14.664264v1)

[8] [A disease resistance protein triggers oligomerization of its NLR helper into a hexameric resistosome to mediate innate immunity](https://www.ncbi.nlm.nih.gov/pmc/articles/PMC11540030/)

[9] [A hydrophobic core in the coiled-coil domain is essential for NRC resistosome function](https://www.biorxiv.org/content/10.1101/2025.01.21.634219v3)

[10] [An atypical NLR protein modulates the NRC immune receptor network in Nicotiana benthamiana | PLOS Genetics](https://journals.plos.org/plosgenetics/article?id=10.1371/journal.pgen.1010500)

[11] [A disease resistance protein triggers oligomerization of its NLR helper into a hexameric resistosome to mediate innate immunity](https://www.biorxiv.org/content/10.1101/2024.06.18.599586v1)

[12] [Frontiers | Tsw – A case study on structure-function puzzles in plant NLRs with unusually large LRR domains](https://www.frontiersin.org/journals/plant-science/articles/10.3389/fpls.2022.983693/full)

[13] [A disease resistance protein triggers oligomerization of its NLR helper into a hexameric resistosome to mediate innate immunity](https://www.ncbi.nlm.nih.gov/pmc/articles/PMC11540030/)

[14] [A disease resistance protein triggers oligomerization of its NLR helper into a hexameric resistosome to mediate innate immunity](https://www.biorxiv.org/content/10.1101/2024.06.18.599586v1)

[15] [An N-terminal motif in NLR immune receptors is functionally conserved across distantly related plant species | eLife](https://elifesciences.org/articles/49956)

[16] [AlphaFold 3: an unprecedent opportunity for fundamental research and drug development - PMC](https://pmc.ncbi.nlm.nih.gov/articles/PMC12342994/)

[17] [Structures of plant resistosome reveal how NLR immune receptors are activated - PMC](https://pmc.ncbi.nlm.nih.gov/articles/PMC9590527/)

[18] [Plant pathogens convergently evolved to counteract redundant nodes of an NLR immune receptor network](https://www.biorxiv.org/content/10.1101/2021.02.03.429184v1)

[19] [An atypical NLR protein modulates the NRC immune receptor network | bioRxiv](https://www.biorxiv.org/content/10.1101/2021.11.15.468391.full)

[20] [Oligomerised RIPK1 is the main core component of the CD95 necrosome](https://www.ncbi.nlm.nih.gov/pmc/articles/PMC12130296/)

[21] [Activation of plant immunity through conversion of a helper NLR homodimer into a resistosome](https://www.ncbi.nlm.nih.gov/pmc/articles/PMC11524475/)

[22] [What AlphaFold 3 struggles with | AlphaFold](https://www.ebi.ac.uk/training/online/courses/alphafold/alphafold-3-and-alphafold-server/introducing-alphafold-3/what-alphafold-3-struggles-with/)

[23] [A helper NLR targets organellar membranes to trigger immunity](https://www.biorxiv.org/content/10.1101/2024.09.19.613839v1)

[24] [A hydrophobic core in the coiled-coil domain is essential for NRC resistosome function](https://www.biorxiv.org/content/10.1101/2025.01.21.634219v2)

[25] [9cc8 - Hexameric state of the NRC4 resistosome - Summary - Protein Data Bank Japan](https://pdbj.org/mine/summary/9cc8)

[26] [NLRscape: an atlas of plant NLR proteins - PMC](https://pmc.ncbi.nlm.nih.gov/articles/PMC9825502/)

[27] [9fp6 - Structure of the NbNRC2 hexameric resistosome - Downloads - Protein Data Bank Japan](https://pdbj.org/mine/resources/9fp6)

[28] [From Prediction to Simulation: AlphaFold 3 as a Differentiable Framework for Structural Biology](https://arxiv.org/html/2508.18446v1)

[29] [Biochemical basis of activation and inhibition of an NLR immune receptor network](https://ueaeprints.uea.ac.uk/id/eprint/93477/1/2023ContrerasMPhD.pdf)

**Motivation:**

$\def\mathcal#1{\mathit{#1}}\def\mathscr#1{\mathit{#1}}$

**Coherence:**

$\def\mathcal#1{\mathit{#1}}\def\mathscr#1{\mathit{#1}}$

**Deep verification:**

$\def\mathcal#1{\mathit{#1}}\def\mathscr#1{\mathit{#1}}$
