## Supplementary Data S6 for "AI-guided discovery of atypical protein assemblies": [EXTERNAL] ├░Y╠ê┬o┬1⁄4 Co-scientist - Hypothesis 265226988747946 - A Multi-Parametric Structural Novelty Index (SNI) for Identifying Unconventional NRC-NLRs via AlphaFold 3 Screening - For research purposes only - Google confidential.docx

### 🧬 Co-scientist - A Multi-Parametric Structural Novelty Index (SNI) for Identifying Unconventional NRC-NLRs via AlphaFold 3 Screening

**About**: *This is an experimental system for generating novel and testable hypotheses. The hypotheses are generated by a model and may be wrong. For research purposes only.*

### Proposal

$\def\mathcal#1{\mathit{#1}}\def\mathscr#1{\mathit{#1}}$

#### The Structural Novelty Index (SNI) Framework

The Structural Novelty Index (SNI) is a multi-parametric score derived from AlphaFold 3 (AF3) models. This index quantifies the structural distance between a query NRC and the canonical templates provided by NbNRC2 (PDB IDs: 9FP6/9RI9) and SlNRC3 (PDB ID: 9CC8).

##### 1. The 9 Quantitative Parameters of the SNI

| # | Parameter Name | Unit | Structural & Biological Significance |
| --- | --- | --- | --- |
| **1** | **Protomer Angular Deviation ($\Delta \theta_{sym}$)** | Degrees ($^\circ$) | The angle between the centroids of three adjacent protomers in the multimer. In a hexamer, this is $60^\circ$. Deviations toward $72^\circ$ suggest pentameric ($C_5$) preference. |
| **2** | **$\alpha 1$-Pore Tilt Angle ($\Phi_{tilt}$)** | Degrees ($^\circ$) | The angle of the N-terminal $\alpha 1$ helix (Residues 1–15) relative to the central Z-axis of the pore. Values $> 25^\circ$ from the canonical "funnel" orientation indicate loss of pore-forming ability. |
| **3** | **NB-ARC Clamshell Aperture ($\Psi_{ARC}$)** | Degrees ($^\circ$) | The angle defined by the P-loop, the hinge region, and the MHD motif. It measures the "opening" of the NB-ARC. Canonical active helpers exhibit an aperture of $\sim 95$–$110^\circ$. |
| **4** | **Interface Shape Complementarity ($S_c$)** | Ratio (0–1) | Measures the geometric "fit" between the NB-ARC of protomer $n$ and the CC/NB-ARC of protomer $n+1$. Values $< 0.6$ suggest a weak or non-functional oligomerization interface. |
| **5** | **LRR Curvature Radius ($R_{LRR}$)** | Angstroms (Å) | Calculated by fitting a circle to the $\alpha$-carbons of the LRR internal solvent-exposed surface. A constricted radius suggests the LRR has evolved for specialized ligand binding. |
| **6** | **MADA Hydrophobic Alignment ($\Delta \mu_H$)** | Vector Angle ($^\circ$) | The alignment of the hydrophobic moment vector of the $\alpha 1$ helix toward the exterior of the funnel. Misalignment prevents stable membrane insertion. |
| **7** | **MHD-to-LRR Allosteric Gap ($\delta_{MHD}$)** | Angstroms (Å) | The Euclidean distance between the His residue of the MHD motif and the nearest LRR internal loop. Canonical helpers maintain a tight distance to ensure sensing translates to activation. |
| **8** | **Axial Stair-Stepping ($\Delta Z_{step}$)** | Angstroms (Å) | The vertical displacement between adjacent protomers. A non-zero $\Delta Z$ suggests a spiral or "broken" ring rather than a planar, pore-forming resistosome. |
| **9** | **Stoichiometry Delta ($\Delta \text{ipTM}$)** | Score (0–1) | The difference in AF3 interface confidence between a $C_6$ and a $C_5$ model ($\text{ipTM}_{C6} - \text{ipTM}_{C5}$). Negative values indicate the sequence "resists" hexameric folding. |

##### 2. Process for Parameter Extraction from AlphaFold 3

To ensure the SNI reflects biology rather than modeling artifacts, a standardized extraction pipeline is utilized:

1. **Generation:** Generate two AF3 models for each query: one as a homo-hexamer ($C_6$) and one as a homo-pentamer ($C_5$).
2. **Coordinate Mapping:** Using a Python-based script (utilizing Biopython and NumPy), anchor the model to conserved structural motifs:
   - **P-loop (GxxxxGK[S/T]):** Used as the origin for the NB-ARC domain.
   - **MHD motif:** Used to define the active-site state.
   - **$\alpha 1$ Hydrophobic Core:** (Residues 3, 6, 7, 10, 14) to calculate tilt and hydrophobic moment.
3. **Geometry Calculation:**
   - The **Central Pore Axis (Z)** is defined by the center of mass of the six NB-ARC domains.
   - $\Delta \theta_{sym}$ is calculated from the centroids of the ARC1 domains.
   - $\Phi_{tilt}$ is calculated by the vector formed by the $\alpha 1$ helix backbone relative to the Z-axis.

##### 3. Implementation: The Tiered Screening Pipeline

Given the computational cost of modeling 6,000 sequences, a phased approach is implemented:

- **Phase 1: Monomer Screening (mSNI):** Model all 6,000 sequences as monomers. Calculate Parameters **#3, #5, and #7**.
  - *Logic:* A sequence cannot form a canonical hexamer if its monomeric "clamshell" or LRR curvature is incompatible. Sequences with a Z-score $> 2.5$ or sequences that match canonical metrics proceed.
- **Phase 2: Multimer Frustration Analysis (cSNI):** Perform dual-stoichiometry modeling ($C_5$ vs $C_6$) for the top 500 candidates.
  - Calculate Parameter **#9 ($\Delta \text{ipTM}$)**. If the $C_5$ state yields a significantly higher confidence score than the $C_6$ state, the protein is flagged as an **Alternative Oligomer**.
- **Phase 3: Weighted SNI Integration:** Calculate the final SNI score for all Phase 2 candidates using the formula: $$SNI = \sum_{i=1}^{9} w_i \frac{|P_{query,i} - \mu_{ref,i}|}{\sigma_{ref,i}}$$
  - **$\mu_{ref}$ and $\sigma_{ref}$:** The mean and standard deviation of parameters derived from an **AF3-generated reference set** of 20 canonical helpers (NbNRC2, SlNRC3, NbNRC4 and close orthologs). This "AI-vs-AI" comparison removes systematic bias between AF3 models and experimental PDBs.
  - **Weights ($w$):** Parameters #1, #2, and #9 are given a weight of 2.0; others are 1.0.

##### 4. Differentiating Unconventional NLRs

The SNI score provides a categorization for the Solanaceae dataset:

- **Canonical Helpers (SNI < 15):** High structural fidelity to NRC2/4. These are functional redundants likely acting as nodes in the network.
- **Signaling Scaffolds (SNI 15–30; High $\Phi_{tilt}$, High $\delta_{MHD}$):** These proteins retain the NB-ARC architecture but have lost the pore-forming funnel or the sensor-to-activator coupling. They may act as decoys or bridge-proteins between sensors and helpers.
- **Specialized Sensors (SNI > 30; Low $R_{LRR}$, High $\Psi_{ARC}$):** NRC-family proteins that have evolved "clenched" LRRs for specific effector recognition and exhibit atypical NB-ARC apertures, suggesting they do not form resistosomes themselves but instead "guard" the network.
- **Non-Hexameric Divergents (SNI > 30; High $\Delta \theta_{sym}$, Negative $\Delta \text{ipTM}$):** Candidates that represent a shift in symmetry (e.g., $C_7$ or $C_8$). These are high-priority targets for structural biology, as they represent new paradigms of NLR execution in the Solanaceae.

### Reviews summary

$\def\mathcal#1{\mathit{#1}}\def\mathscr#1{\mathit{#1}}$

### Appendix:

**Motivation:**

$\def\mathcal#1{\mathit{#1}}\def\mathscr#1{\mathit{#1}}$

**Coherence:**

$\def\mathcal#1{\mathit{#1}}\def\mathscr#1{\mathit{#1}}$

**Deep verification:**

$\def\mathcal#1{\mathit{#1}}\def\mathscr#1{\mathit{#1}}$
