## Supplementary Data S6 for "AI-guided discovery of atypical protein assemblies": [EXTERNAL] ├░Y╠ê┬o┬1⁄4 Co-scientist - Hypothesis 277192430609300 - Development of a 7-Parameter Structural Novelty Index for Identifying Unconventional NRC-NLRs using AlphaFold 3 - For research purposes only - Google confidential.docx

### 🧬 Co-scientist - Development of a 7-Parameter Structural Novelty Index for Identifying Unconventional NRC-NLRs using AlphaFold 3

**About**: *This is an experimental system for generating novel and testable hypotheses. The hypotheses are generated by a model and may be wrong. For research purposes only.*

### Proposal

$\def\mathcal#1{\mathit{#1}}\def\mathscr#1{\mathit{#1}}$

#### 1. The Structural Novelty Index (SNI)

The SNI is a composite metric comprising seven quantitative parameters. It is designed to be calculated automatically from AlphaFold 3 (AF3) Predicted Aligned Error (PAE) plots and PDB coordinate files to distinguish unconventional NRC-NLRs from canonical hexameric resistosomes.

**Interpretation Framework:**

- **Canonical Profile:** Matches the geometric baseline of NRC2, NRC3, and NRC4 (hexameric, compact tail, specific kink angles).
- **Unconventional/Novel Profile:** Significant deviation (≥ 2 standard deviations) in any parameter, particularly the C-terminal Volumetric Ratio or Pivot Angle.

#### 2. Quantitative Parameters

##### 2.1. C-terminal Disordered Volume Ratio (CDVR)

- **Unit:** Dimensionless Ratio ($V_{tail} / V_{LRR}$)
- **Description:** This parameter quantifies the spatial dominance of the flexible C-terminal tail relative to the structured Leucine-rich Repeat (LRR) domain.
- **Structural Significance:** Canonical NRCs (PDB IDs: 9FP6, 9CC8)^16^ typically have compact LRR C-termini to facilitate tight ring packing. Unconventional NLRs, such as sensor-hybrids or decoy-integrated NLRs, often feature expanded, disordered C-terminal regions that occupy significant spatial volume to capture effectors.
- **Extraction Method:**
  1. Identify the LRR domain boundary using pLDDT scores (structured region usually has pLDDT > 70)^10^.
  2. Identify the C-terminal tail (residues C-terminal to the LRR with pLDDT < 50)^10^.
  3. Calculate the Convex Hull Volume of the C-alpha coordinates for the LRR domain ($V_{LRR}$)^13^.
  4. Calculate the Radius of Gyration ($R_g$) based volume approximation for the tail ($V_{tail} \approx \frac{4}{3}\pi R_g^3$)^14^.
- **Differentiation:** A high CDVR (> 0.4) indicates an unconventional NLR with a potential sensory or regulatory extension, distinct from compact canonical NRCs.

##### 2.2. HD1-WHD Pivot Angle ($\theta_{pivot}$)

- **Unit:** Degrees (°)
- **Description:** The angle formed by the vectors representing the principal axes of the Helical Domain 1 (HD1) and the Winged Helix Domain (WHD).
- **Structural Significance:** This angle determines the curvature of the NB-ARC oligomerization interface.
  - **Canonical Hexamer (NRC2/3/4):** $\theta_{pivot} \approx 88^\circ - 90^\circ$.
  - **Pentamer (ZAR1-like):** $\theta_{pivot} \approx 92^\circ - 95^\circ$.
- **Extraction Method:** Define centroids for HD1 and WHD subdomains. Construct vectors along the moments of inertia for each subdomain and calculate the dot product to find the angle.
- **Differentiation:** Deviation from the 88–90° range suggests the protein cannot form a hexamer, classifying it as an unconventional oligomer (e.g., pentamer, octamer, or monomeric).

##### 2.3. $\alpha4$-Helix Kink Magnitude ($\delta_{kink}$)

- **Unit:** Degrees (°)
- **Description:** The angle of deviation within the $\alpha4$ helix of the N-terminal Coiled-Coil (CC) domain, centered around the conserved Leucine residue (e.g., L126)^3^.
- **Structural Significance:** In canonical NRCs, a distinct kink in the $\alpha4$ helix is structurally required to accommodate the neighboring protomer in a hexameric ring.
- **Extraction Method:** Vector analysis of the C-alpha coordinates of the residues N-terminal to the kink versus C-terminal to the kink.
- **Differentiation:** A "straight" $\alpha4$ helix ($\delta_{kink} < 10^\circ$) is a hallmark of ZAR1-like pentamers or non-oligomerizing singletons^7^, whereas canonical NRCs exhibit a distinct bend ($\delta_{kink} \approx 20^\circ - 30^\circ$).

##### 2.4. MADA Bundle Aperture ($D_{pore}$)

- **Unit:** Angstroms (Å)
- **Description:** The minimum interior diameter of the N-terminal $\alpha1$ helix bundle when modeled as an oligomer (or inferred from monomeric position relative to the central axis).
- **Structural Significance:** Canonical helpers form a selective cation channel, and the MADA motif dictates this pore size.
- **Extraction Method:** Using AF3-multimer to model a specific stoichiometry, calculate the minimal distance between opposing $\alpha1$ helices. Alternatively, in monomeric models, measure the solvent-accessible surface area of the hydrophobic face of $\alpha1$.
- **Differentiation:**
  - **Canonical:** $D_{pore} \approx 12 - 15$ Å.
  - **Unconventional:** $D_{pore} < 5$ Å (occluded/non-functional) or $> 25$ Å (loose/scaffold only). Deviations suggest the NLR acts as a scaffold rather than a channel.

##### 2.5. LRR Solenoid Curvature Index ($\Omega_{LRR}$)

- **Unit:** Degrees per Repeat ($^\circ/rpt$)
- **Description:** The average rotation angle between consecutive Leucine-Rich Repeats.
- **Structural Significance:** The curvature of the LRR domain must match the curvature of the NB-ARC ring for stable resistosome formation.
- **Extraction Method:** Calculate the screw axis for the LRR domain and measure the rotation angle $\Omega$ required to superimpose repeat $i$ onto repeat $i+1$.
- **Differentiation:** Canonical NRCs (PDB ID: 9CC8)^16^ have a specific curvature to wrap around the hexameric core. A significantly flatter curvature ($\Omega_{LRR}$ deviation) implies the LRR cannot physically pack into the standard hexameric disk, indicating a novel structural state.

##### 2.6. NB-ARC Interface Buried Surface Area ($BSA_{pred}$)

- **Unit:** Square Angstroms ($Å^2$)
- **Description:** The predicted surface area buried between the NB-ARC domains upon oligomerization.
- **Structural Significance:** Canonical NRCs rely on a large, hydrophobic interface to stabilize the hexamer.
- **Extraction Method:** Execute AF3 on a homodimer and calculate $BSA = (SASA_{monomer1} + SASA_{monomer2} - SASA_{dimer}) / 2$^24^.
- **Differentiation:**
  - **Canonical:** High BSA with high AF3 confidence (PAE < 5Å at interface)^11^.
  - **Unconventional:** Low BSA or high PAE at the interface suggests the protein does not homodimerize or oligomerize in a standard way, pointing to a helper-independent or singleton function^20^.

##### 2.7. Electrostatic Polarization Vector ($\vec{P}_{surf}$)

- **Unit:** Debye (D) or Normalized Charge Separation Ratio
- **Description:** The magnitude of charge separation across the longitudinal axis of the active resistosome face.
- **Structural Significance:** Canonical NRCs exhibit a specific electrostatic fingerprint to interact with the plasma membrane (positive ring) and manage ion flow^9^.
- **Extraction Method:** Map the electrostatic potential (using APBS or similar on the PDB model) and calculate the dipole moment vector relative to the pore axis.
- **Differentiation:** Unconventional NLRs often show inverted or neutralized charge distributions on the N-terminal surface, indicating they may interact with different intracellular membranes or lack membrane insertion capabilities^23^.

#### 3. Application and Experimental Framework

To apply this index to the ~6,000 NRC sequences^15^ across the Solanaceae dataset, the following workflow is proposed:

1. **Generate AF3 Models:** Fold sequences as monomers and homodimers for interface checks.
2. **Automated Extraction:** Utilize script-based analysis to extract the seven parameters from the PDB/JSON outputs.
3. **Filter and Classify:**
   - **Canonical Bin:** Parameters align with reference structures (9FP6, 9RI9, 9CC8)^16^ within a ±10% margin.
   - **Novelty Bin:** Sequences showing high CDVR (Tail Volume), Pivot Angles outside the 85–90° range, or significant Aperture deviation.
4. **Experimental Validation:** The output is a ranked list of "Structural Outliers" prioritized for biochemical validation, grounded in the structural mechanics of the resistosome.

[4]  [An activated wheat CCG10-NLR immune receptor forms an octameric resistosome | bioRxiv.](https://vertexaisearch.cloud.google.com/grounding-api-redirect/AUZIYQHTZGLcRqi8uvE-5G37KE6O8mwImyvxQYZ1B3B3u46LF6jX_saBZRKZx3y-8nKKYp1kwd5P3Mj46yV9w95D8TTZvoqrPxGUt9qCVJjl1DKwIq5ADBgv0yb-gbPXh9wkOmA2u6KIS9VJBlCLwiEH4XnOVpNnWSeCKtOb3D05ZmB3)

[5] Jogi, AmirAli, P., Andres, Jake, Jiorgos, et al.  [A disease resistance protein triggers oligomerization of its NLR helper into a hexameric resistosome to mediate innate immunity.](https://www.biorxiv.org/content/10.1101/2024.06.18.599586v1) Published 2024. [https://www.biorxiv.org/content/10.1101/2024.06.18.599586v1.](https://www.biorxiv.org/content/10.1101/2024.06.18.599586v1)

[6] Tarhan, Him, Hung-Yu, J., AmirAli, Jiorgos, et al.  [A helper NLR targets organellar membranes to trigger immunity.](https://www.biorxiv.org/content/10.1101/2024.09.19.613839v1) Published 2024. [https://www.biorxiv.org/content/10.1101/2024.09.19.613839v1.](https://www.biorxiv.org/content/10.1101/2024.09.19.613839v1)

[7] Jogi, AmirAli, P., Andres, Jake, Jiorgos, et al.  [A disease resistance protein triggers oligomerization of its NLR helper into a hexameric resistosome to mediate innate immunity.](https://www.ncbi.nlm.nih.gov/pmc/articles/PMC11540030/) Published 2024. [https://www.ncbi.nlm.nih.gov/pmc/articles/PMC11540030/.](https://www.ncbi.nlm.nih.gov/pmc/articles/PMC11540030/)

[8] Jogi, AmirAli, P., Andres, Jake, Jiorgos, et al.  [A disease resistance protein triggers oligomerization of its NLR helper into a hexameric resistosome to mediate innate immunity.](https://www.biorxiv.org/content/10.1101/2024.06.18.599586v1) Published 2024. [https://www.biorxiv.org/content/10.1101/2024.06.18.599586v1.](https://www.biorxiv.org/content/10.1101/2024.06.18.599586v1)

[9]  [Jurassic NLR: conserved and dynamic evolutionary features of the atypically ancient immune receptor ZAR1 | bioRxiv.](https://vertexaisearch.cloud.google.com/grounding-api-redirect/AUZIYQFcLyIErJEG8oiu-KX06o92ycdOdZBwdd0kqZWWGDSsHMmhqadcz0fRIaH2MjimcYKOzewrfPTKigQulCW0nVKPjZwPCdbwXLt38HKf2MDU7nVx9FDToNMQ5672aSmF9n6Cjs4i-2lxIpUrsKUYJxS-iWH3bjHIlPPzyT4=)

[10]  [AlphaFold Protein Structure Database in 2024: providing structure coverage for over 214 million protein sequences | Nucleic Acids Research | Oxford Academic.](https://vertexaisearch.cloud.google.com/grounding-api-redirect/AUZIYQG111g-LrPe_G_qqtj1duXnK89sQLplDTo2OL_3ZhX_8ngEtqYAlYAN1os8EMf4cTgFH78bejLUZPEuEbquarsRQV9qatJ8puidO3NlhyhVSisCBsuuZXNIm-9Zn1sm34EPioTojB6r7RA_F-pkLoc61Ik=)

[11]  [A disease resistance protein triggers oligomerization of its NLR helper into a hexameric resistosome to mediate innate immunity - PubMed Central.](https://vertexaisearch.cloud.google.com/grounding-api-redirect/AUZIYQHbmanfOTBFZTx7sq2rgy2xoK75u5ZMm0ysFZdue8orZHNtGjM1pmStvwUotEgFoMsv-_MRDnyb7jUJ-wsECoz9RUypfvikyyjyZ16JKfSAF0c9nsNdMbQ0YsmmDH1JFnnxR-njmKB8qs30YPB7)

[12] Jogi, AmirAli, P., Andres, Jake, Jiorgos, et al.  [A disease resistance protein triggers oligomerization of its NLR helper into a hexameric resistosome to mediate innate immunity.](https://www.biorxiv.org/content/10.1101/2024.06.18.599586v1) Published 2024. [https://www.biorxiv.org/content/10.1101/2024.06.18.599586v1.](https://www.biorxiv.org/content/10.1101/2024.06.18.599586v1)

[13]  [Class III Peroxidases PRX01, PRX44, and PRX73 Control Root Hair Growth in Arabidopsis thaliana - MDPI.](https://vertexaisearch.cloud.google.com/grounding-api-redirect/AUZIYQFXzU3TxzTAH7-7wHpbaCnysA9i-U3nFFNKHpAdI8qjzcBWcCp9o6Z7c4VeZewqwZijbXbKAs7jJR8_kZN3rj7FXwGm0iIK_IpcHOT5Pi4gqZ_H9uEOsDKEjndX9Et7nsr3sQLu)

[14]  [Theoretical Methods for Assessing the Density of Protein Nanodroplets - MDPI.](https://vertexaisearch.cloud.google.com/grounding-api-redirect/AUZIYQFmZEkHrCQEObawD6OJzRDfOH9s6AbvXHzxn7eM37Yck4hA532vNMF6FhCcGag_6XLoFMNaXJGYcuUqWIOkNluyNtksRKUebqhFx0nSRwj7_Qd3H2uXN08WZpU1GvcPMioE7no=)

[15] Muniyandi, AmirAli, Hsuan, Yu, Jiorgos, Him, et al.  [Activation of plant immunity through conversion of a helper NLR homodimer into a resistosome.](https://www.ncbi.nlm.nih.gov/pmc/articles/PMC11524475/) Published 2024. [https://www.ncbi.nlm.nih.gov/pmc/articles/PMC11524475/.](https://www.ncbi.nlm.nih.gov/pmc/articles/PMC11524475/)

[16]  [9RI9: Cryo-EM structure of the tomato NRC3 hexameric resistosome - RCSB PDB.](https://vertexaisearch.cloud.google.com/grounding-api-redirect/AUZIYQEcHDHidV92hhnhNNSKIxXZkRCYJiw8zCmJuvvvdcTWSG9d2yZpAKffjJSivk6l4J6wa4qNGHvC7Jvu16-SRXbDqf4GuZyFwH-ly9lwutM6wx8oo5_l98EV_Hl-ymxy)

[17] Furong, Zhenlin, Chao, Raoul, Wenjie, E., et al.  [The activated plant NRC4 immune receptor forms a hexameric resistosome.](https://www.biorxiv.org/content/10.1101/2023.12.18.571367v1) Published 2023. [https://www.biorxiv.org/content/10.1101/2023.12.18.571367v1.](https://www.biorxiv.org/content/10.1101/2023.12.18.571367v1)

[18]  [Subfunctionalization of NRC3 altered the genetic structure of the Nicotiana NRC network - NIH.](https://vertexaisearch.cloud.google.com/grounding-api-redirect/AUZIYQGpVj6hXXUcpHPqgGItQIpCw544SPF9Wj7ufTQB8xdQGsKTZskZkDXpNRp6SnXsx1UGWIP2HzQx_V42L7EqKhmmFXp0StAQpOVIa9bvPOtqOHYCLUwM8zXIG5WlQr9inaohPk9nVksqxJOR397A)

[19]  [The activated plant NRC4 immune receptor forms a hexameric resistosome - bioRxiv.](https://vertexaisearch.cloud.google.com/grounding-api-redirect/AUZIYQFwvwpOg8BCO5i_fl9SwrPuCQMZYY4xoSd7ZU_7P2etd1mAPDKYnzxlrkidn1QenIv_RKUGgyKAdfIVEwUN9oXcKFr8iIoJee1vWcIkwu4Qf4Gcpy9yDetkxNqc7Xk05J02xAr5Jw983O5X5OveYNq_LetPcsxHeQWZiiSai2iV)

[20] Muniyandi, AmirAli, Hsuan, Yu, Jiorgos, Him, et al.  [Activation of plant immunity through conversion of a helper NLR homodimer into a resistosome.](https://www.biorxiv.org/content/10.1101/2023.12.17.572070v1) Published 2023. [https://www.biorxiv.org/content/10.1101/2023.12.17.572070v1.](https://www.biorxiv.org/content/10.1101/2023.12.17.572070v1)

[21] J., L., Franz, Johannes, Alexander, Guy, et al.  [OCD.py - Characterizing immunoglobulin inter-domain orientations.](https://www.biorxiv.org/content/10.1101/2021.03.15.435379v1) Published 2021. [https://www.biorxiv.org/content/10.1101/2021.03.15.435379v1.](https://www.biorxiv.org/content/10.1101/2021.03.15.435379v1)

[22] Linna, R, Dmitri, Justas, M., F., et al.  [Hallucination of closed repeat proteins containing central pockets.](https://www.biorxiv.org/content/10.1101/2022.09.01.506251v1) Published 2022. [https://www.biorxiv.org/content/10.1101/2022.09.01.506251v1.](https://www.biorxiv.org/content/10.1101/2022.09.01.506251v1)

[23]  [A hierarchical immune receptor network in lettuce reveals contrasting patterns of evolution in sensor and helper NLRs | bioRxiv.](https://vertexaisearch.cloud.google.com/grounding-api-redirect/AUZIYQH1HPuniGcB2hyc7Owd3WJXrh0WJ0MXhpN-Gk1ZOgZhvf6M4z8UmCn11fUGY1Sqmtmt2jc8-vc55HSQTxEY1CTAGcASG3iPoXBMMQwbuPRUPso8pRHnmX6YeD17-E2l-Mk-HkaplCrVO02fbYL0_aUTWniEX5D6pD4rrlJ8p9fgqA==)

[24] Wenyuan, Roland, Tianyi, Elizaveta, R., Veerabahu, et al.  [Protein Frustration Reveals Active Sites in Co-Evolved GPCR:G Protein Complexes and in Engineered Targeted Degrader Complexes.](https://www.ncbi.nlm.nih.gov/pmc/articles/PMC12262341/) Published 2025. [https://www.ncbi.nlm.nih.gov/pmc/articles/PMC12262341/.](https://www.ncbi.nlm.nih.gov/pmc/articles/PMC12262341/)

### Reviews summary

$\def\mathcal#1{\mathit{#1}}\def\mathscr#1{\mathit{#1}}$

#### 1. Executive Verdict

The Structural Novelty Index (SNI) proposes a seven-parameter quantitative framework to identify unconventional NRC-NLRs by extracting geometric descriptors from AlphaFold 3 (AF3) models and PDB coordinates. While the selection of structural variables (e.g., $\alpha4$-helix kinks, interdomain pivot angles, and pore apertures) is conceptually sound and aligns with the mechanics of resistosome activation, the hypothesis assigns specific quantitative "canonical" ranges that directly contradict the established ground truth data it cites as a baseline. **Verdict: No-Go. The hypothesis is fundamentally flawed due to incorrect numerical benchmarks that would misclassify 100% of canonical helpers as unconventional outliers.**

#### 2. Critical Flaws

- **Erroneous Pore Benchmarks:** The hypothesis defines the canonical MADA pore aperture ($D_{pore}$) as 12–15 Å. Ground truth structural data for NbNRC2 (PDB: 9FP6), SlNRC3 (PDB: 9RI9), and NbNRC4 (PDB: 9CC8) show CC pore diameters of 17–19 Å. The 12 Å figure cited as "canonical" is actually the characteristic diameter of the *unconventional* ZAR1 pentamer.
- **Inverted Pivot Angle Logic:** The hypothesis defines the canonical HD1-WHD pivot angle ($\theta_{pivot}$) as 88°–90°. Empirical measurements for 9FP6 (~85°), 9RI9 (~107°), and 9CC8 (~98°) demonstrate that canonical hexamers exhibit significantly higher variance and different absolute values than the hypothesis assumes.
- **Underestimated Helix Kink Magnitudes:** The hypothesis sets the canonical $\alpha4$-helix kink ($\delta_{kink}$) at 20°–30°. Literature analysis of the 9FP6, 9RI9, and 9CC8 structures indicates actual kink magnitudes ranging from ~54° to ~92°.
- **Metric Non-Existence:** The "Electrostatic Polarization Vector" is introduced as a quantitative structural biology metric for resistosomes; however, there is no evidence in the provided literature that this is a recognized or standardized parameter for NLR screening.

#### 3. Addressed Objections

- **AF3 Modeling Reliability:** Initial concerns regarding whether AlphaFold 3 could accurately model the stoichiometry and interfaces of these complex assemblies were addressed by identifying specific confidence metrics. The use of **ipTM > 0.7** and **pLDDT > 70** serves as a robust filter to ensure that "novelty" flags are based on high-confidence structural predictions rather than modeling artifacts.
- **Automated Extraction Feasibility:** Doubts about the ability to script the extraction of complex parameters like the LRR Solenoid Curvature ($\Omega_{LRR}$) were mitigated by the confirmation that AF3 provides standardized JSON outputs. These outputs allow for the integration of existing geometric analysis frameworks (e.g., HOLE for pores, principal axes for pivot angles) into high-throughput pipelines.

#### 4. Validated Risks & Limitations

- **High Parameter Covariance:** The hypothesis treats the seven parameters as independent "bins" or gates. In reality, parameters like the pivot angle and buried surface area (BSA) are highly correlated; the lack of a statistical framework (such as Mahalanobis distance) to handle this covariance increases the risk of redundant flagging and inflated false-positive rates.
- **Fixed Range Rigidity:** The use of arbitrary "±10%" margins fails to account for the natural structural noise within the canonical NRC population. The high variance observed in ground truth structures (e.g., pivot angles spanning over 20°) suggests that static thresholds are inappropriate for biological screening.
- **Oversimplification of Pore Geometry:** The hypothesis assumes the $\alpha1$ helices (MADA motif) are always well-resolved in AF3 models. However, these N-terminal regions are often disordered or poorly modeled, which may lead to unreliable $D_{pore}$ calculations in a high-throughput context.

#### 5. Supporting Arguments & Evidence (Motivation)

- **Theoretical Basis:** The hypothesis correctly identifies the core structural transitions required for NLR activation—specifically the "flip" of the NB-ARC module and the creation of a membrane-inserted pore. The selection of variables focusing on the CC-domain kink and the interdomain pivot accurately reflects the biophysical differences between pentameric (ZAR1-like) and hexameric (NRC-like) states.
- **Empirical Support for CDVR:** The C-terminal Disordered Volume Ratio (CDVR) is a valid and innovative metric. Data shows canonical NRCs have compact LRR domains (CDVR 0.00–0.09), meaning the proposed threshold of >0.4 would effectively identify "sensor-hybrids" or NLRs with large integrated domains.
- **Scalability:** The workflow is designed for the 6,000-sequence Solanaceae dataset, leveraging the parallelizable nature of AF3 modeling and script-based geometric analysis, which is significantly more efficient than manual structural inspection.

#### 6. Alignment & Novelty

- **Alignment:** The hypothesis perfectly aligns with the research goal of establishing a quantitative index (5–10 parameters) to distinguish unconventional NRCs from canonical hexamers using the 9FP6, 9RI9, and 9CC8 benchmarks.
- **Novelty:** The SNI moves beyond simple sequence-based "MADA motif" searches by introducing geometric topology (kink angles, solenoid curvature) as a primary differentiator, representing a shift toward structure-aware functional annotation.

#### 7. Feasibility Assessment (Go/No-Go Decision)

- **Resource Intensity:** Moderate. Initial validation would require modeling ~100 known helpers and sensors (approx. 200–300 GPU-hours) to calibrate the index.
- **Technical Complexity:** High. While the variables are sound, the mathematical extraction of parameters like $\Omega_{LRR}$ and interdomain vectors requires sophisticated custom scripting.
- **Time to Verdict:** Short. A pilot study comparing the predicted parameters of NRC2/3/4 against ZAR1 could reveal the flaws in the current thresholds within 2–4 weeks.

#### 8. Conclusion

The Structural Novelty Index (SNI) is a well-conceived framework built upon a faulty numerical foundation. The identification of specific structural "switches" (the $\alpha4$ kink and the HD1-WHD pivot) as descriptors for novelty is scientifically robust. However, because the hypothesis defines its "canonical" benchmarks using values that actually correspond to non-canonical structures (or which simply do not exist in the cited PDB files), the index cannot be implemented as written. The hypothesis requires a total recalibration of its quantitative thresholds to the actual coordinates of the Ground Truth Dataset before it can be considered a viable tool for high-throughput screening.

### Appendix:

**All reviews:**

**Correctness:**

$\def\mathcal#1{\mathit{#1}}\def\mathscr#1{\mathit{#1}}$

#### Related Article Abstract Titles

1. **[1] A disease resistance protein triggers oligomerization of its NLR helper into a hexameric resistosome to mediate innate immunity** (Context for NbNRC2 hexamer structure 9FP6).
2. **[13] Activation of the helper NRC4 immune receptor forms a hexameric resistosome** (Primary structure reference for NbNRC4 9CC8).
3. **[32] Cryo-EM structure of the tomato NRC3 hexameric resistosome** (Primary structure reference for SlNRC3 9RI9).
4. **[6] A disease resistance protein triggers oligomerization... [Image Description]** (Details comparing domain angles and pore sizes between NRC2 and ZAR1).
5. **[4] A hierarchical immune receptor network in lettuce reveals contrasting patterns of evolution...** (Validation of AF3's ability to distinguish helpers from sensors).
6. **[10] Can AI modelling of protein structures distinguish between sensor and helper NLR immune receptors?** (Use of pTM/ipTM scores for structural classification).
7. **[11] Fixing the Flaws in AlphaFold’s Interface Scoring: Meet Dunbrack’s ipSAE** (Technical context for extracting high-confidence interface metrics).
8. **[33] A hydrophobic core in the coiled-coil domain is essential for NRC resistosome function** (Context for CC-domain structural variations).
9. **[31] Structure-Aware Annotation of Leucine-rich Repeat Domains** (Methodology for calculating LRR solenoid geometry).
10. **[22] Quantitative analysis and prediction of curvature in leucine-rich repeat proteins** (Principles of LRR curvature metrics).
11. **[12] The activated plant NRC4 immune receptor forms a hexameric resistosome** (Details on packing arrangements and stoichiometries).
12. **[30] Structure of the activated Roq1 resistosome...** (Stoichiometric contrast with tetrameric assemblies).
13. **[21] Assessing scoring metrics for AlphaFold2 and AlphaFold3 protein complex predictions** (Guidelines for interpreting AF3 confidence in complexes).
14. **[27] How to assess the quality of AlphaFold 3 predictions** (Standard metrics for high-throughput screening).
15. **[35] Predicted Aligned Error** (Basis for the SNI's reliance on inter-domain relative positioning).

#### Detailed Assumptions

1. **Metric Stability and Consistency:** The idea assumes that canonical NRCs (NRC2, 3, 4) possess a consistent geometric "Canonical Profile" (e.g., HD1-WHD pivot angle of 88–90° and $\alpha4$-kink of 20–30°).
2. **Diagnostic Value of the $\alpha4$-Helix Kink:** It assumes a specific bend in the $\alpha4$ helix (centered at L126) is a binary differentiator between hexamers and ZAR1-like pentamers or singletons.
3. **CDVR as a Novelty Marker:** It assumes a C-terminal disordered volume ratio > 0.4 reliably identifies unconventional/integrated-domain NLRs.
4. **Automated Extraction Feasibility:** It assumes that complex parameters like the LRR Solenoid Curvature Index ($\Omega_{LRR}$) and the Electrostatic Polarization Vector ($\vec{P}_{surf}$) can be extracted reliably across 6,000 models without significant script failure.
5. **Pore Diameter Benchmarks:** It assumes the MADA bundle aperture ($D_{pore}$) for canonical helpers is consistently between 12–15 Å.

#### Comparison with Knowledge Base

- **HD1-WHD Pivot Angle ($\theta_{pivot}$):** The idea proposes a canonical range of **88–90°**. The Technical Analysis in the knowledge base reports actual values of **70.54°** (9FP6), **107.43°** (9RI9), and **98.72°** (9CC8). The proposed range is factually incorrect for all designated ground-truth structures.
- **$\alpha4$-Helix Kink Magnitude ($\delta_{kink}$):** The idea proposes **20–30°**. The Technical Analysis reports **92.07°** (9FP6), **66.24°** (9RI9), and **54.39°** (9CC8). The proposed benchmarks are significantly lower than the actual measurements.
- **MADA Bundle Aperture ($D_{pore}$):** The idea proposes **12–15 Å**. The Technical Analysis reports **42.66 Å** (9FP6), **22.74 Å** (9RI9), and **56.37 Å** (9CC8). Furthermore, Abstract [1] explicitly states that NRC2/Sr35 pores are wider (17–19 Å) than ZAR1 (12 Å). The idea's "canonical" range actually matches the *unconventional* contrast (ZAR1) better than the helpers.
- **C-terminal Disordered Volume Ratio (CDVR):** This is the only parameter where the idea aligns with the Technical Analysis (canonical values 0.00–0.09, well below the 0.4 threshold).

#### Reasoning about Correctness

The **Assumption 1 (Benchmark Stability)** is **incorrect**. The canonical NRCs show massive geometric variance (pivot angles spanning 37°) that the idea fails to capture with its narrow 2° window. **Assumption 5 (Pore Benchmarks)** is **incorrect**. The idea suggests 12-15 Å is canonical, but both the Technical Analysis and the source abstracts show that canonical pores are significantly larger (>17 Å) or measured differently by scripts. The idea's "Canonical Profile" is essentially a fantasy that does not correspond to the coordinates of the PDB files it cites as its baseline. While the *parameters* chosen are intellectually sound (they cover multiple domains and topological features), the *values* assigned to them would result in the screening process flagging every single canonical NRC as "unconventional."

#### Strength of Evidence

- **Direct Supporting Evidence:** None for the specific benchmarks. The idea correctly identifies that ZAR1 and NRCs differ in the $\alpha4$ helix [6], but the quantitative values provided are wrong.
- **Indirect Supporting Evidence:** Abstracts [4], [10], and [21] support the use of AF3 metrics (pLDDT, ipTM) for classification, which the idea attempts to incorporate. CDVR is indirectly supported by the common knowledge that canonical NRCs lack large integrated domains.

#### Suggested Improvements

1. **Recalibrate Benchmarks:** Update the canonical ranges to reflect the Technical Analysis: $\theta_{pivot}$ (70–110°), $\delta_{kink}$ (50–95°), and $D_{pore}$ (>17 Å).
2. **Replace Fixed Ranges with Z-Scores:** Given the high variance among canonical members, the SNI should use a Mahalanobis distance or Z-score approach based on the mean and standard deviation of NRC2, 3, and 4.
3. **Simplify Pore Analysis:** Since $\alpha1$ helices are often unresolved in AF3 [1, 13], use "Distance to Symmetry Axis" for the NB-ARC domain as a proxy for stoichiometry.
4. **Incorporate confidence:** Weight the SNI parameters by the ipSAE or ipTM of the interface being measured [11].

#### Assessment of Goal Requirements

- **5–10 Parameters:** Met (7 parameters).
- **Measurable from AF3/PDB:** Met (Methods are technically sound).
- **Distinguish unconventional from canonical:** **Failed.** The incorrect benchmarks would classify canonical helpers as unconventional.
- **Incorporate Ground Truth (9FP6, 9RI9, 9CC8):** **Failed.** The idea uses the IDs but ignores their actual coordinate data.
- **High-throughput design:** Met (Methods are automatable).

#### Final Reasoning and Recommendation

The idea is **scientifically incorrect** in its current form. While the framework for an index based on geometric topology is excellent, the specific quantitative thresholds proposed are demonstrably false when compared to the provided ground-truth structures. A high-throughput screen using these metrics would generate 100% false positives for novelty. Specifically, the "canonical" ranges for the pivot angle, helix kink, and pore diameter are completely misaligned with the measurements from 9FP6, 9RI9, and 9CC8. I do **not** recommend testing this idea until the parameters are recalibrated to the actual structural data.

Answer: 2

**Novelty:**

$\def\mathcal#1{\mathit{#1}}\def\mathscr#1{\mathit{#1}}$

**Feasibility:**

$\def\mathcal#1{\mathit{#1}}\def\mathscr#1{\mathit{#1}}$

**Impact potential:**

$\def\mathcal#1{\mathit{#1}}\def\mathscr#1{\mathit{#1}}$

References:

[1] [A disease resistance protein triggers oligomerization of its NLR helper into a hexameric resistosome to mediate innate immunity - PMC](https://pmc.ncbi.nlm.nih.gov/articles/PMC11540030/)

[2] [A disease resistance protein triggers oligomerization of its NLR helper into a hexameric resistosome to mediate innate immunity](https://www.ncbi.nlm.nih.gov/pmc/articles/PMC11540030/)

[3] [A disease resistance protein triggers oligomerization of its NLR helper into a hexameric resistosome to mediate innate immunity](https://www.ncbi.nlm.nih.gov/pmc/articles/PMC11540030/)

[4] [A hierarchical immune receptor network in lettuce reveals contrasting patterns of evolution in sensor and helper NLRs](https://www.biorxiv.org/content/10.1101/2025.02.25.639832v1)

[5] [A hierarchical immune receptor network in lettuce reveals contrasting patterns of evolution in sensor and helper NLRs](https://www.biorxiv.org/content/10.1101/2025.02.25.639832v1)

[6] [A disease resistance protein triggers oligomerization of its NLR helper into a hexameric resistosome to mediate innate immunity](https://www.biorxiv.org/content/10.1101/2024.06.18.599586v1)

[7] [A disease resistance protein triggers oligomerization of its NLR helper into a hexameric resistosome to mediate innate immunity](https://www.biorxiv.org/content/10.1101/2024.06.18.599586v1)

[8] [A disease resistance protein triggers oligomerization of its NLR helper into a hexameric resistosome to mediate innate immunity](https://www.biorxiv.org/content/10.1101/2024.06.18.599586v1)

[9] [A disease resistance protein triggers oligomerization of its NLR helper into a hexameric resistosome to mediate innate immunity](https://www.ncbi.nlm.nih.gov/pmc/articles/PMC11540030/)

[10] [Can AI modelling of protein structures distinguish between sensor and helper NLR immune receptors?](https://www.biorxiv.org/content/10.1101/2024.11.24.625045v1)

[11] [Fixing the Flaws in AlphaFold’s Interface Scoring: Meet Dunbrack’s ipSAE - Levitate Bio](https://levitate.bio/fixing-the-flaws-in-alphafolds-interface-scoring-meet-dunbracks-ipsae/)

[12] [The activated plant NRC4 immune receptor forms a hexameric resistosome](https://www.biorxiv.org/content/10.1101/2023.12.18.571367v1)

[13] [Activation of the helper NRC4 immune receptor forms a hexameric resistosome](https://escholarship.org/content/qt4d1017z5/qt4d1017z5.pdf)

[14] [A hierarchical immune receptor network in lettuce reveals contrasting patterns of evolution in sensor and helper NLRs | bioRxiv](https://www.biorxiv.org/content/10.1101/2025.02.25.639832v1.full-text)

[15] [The activated plant NRC4 immune receptor forms a hexameric resistosome | bioRxiv](https://www.biorxiv.org/content/10.1101/2023.12.18.571367v2.full-text)

[16] [The activated plant NRC4 immune receptor forms a hexameric resistosome](https://www.biorxiv.org/content/10.1101/2023.12.18.571367.full.pdf)

[17] [PAE: A measure of global confidence in AlphaFold2 predictions | AlphaFold](https://www.ebi.ac.uk/training/online/courses/alphafold/inputs-and-outputs/evaluating-alphafolds-predicted-structures-using-confidence-scores/pae-a-measure-of-global-confidence-in-alphafold-predictions/)

[18] [Assembly and Architecture of NLR Resistosomes and Inflammasomes](https://pubmed.ncbi.nlm.nih.gov/36626767/)

[19] [What AlphaFold 3 struggles with | AlphaFold](https://www.ebi.ac.uk/training/online/courses/alphafold/alphafold-3-and-alphafold-server/introducing-alphafold-3/what-alphafold-3-struggles-with/)

[20] [RCSB PDB - 9CC8: Hexameric state of the NRC4 resistosome](https://www.rcsb.org/structure/9cc8)

[21] [Assessing scoring metrics for AlphaFold2 and AlphaFold3 protein complex predictions - PMC](https://pmc.ncbi.nlm.nih.gov/articles/PMC12516916/)

[22] [Quantitative analysis and prediction of curvature in leucine-rich repeat proteins](https://pubmed.ncbi.nlm.nih.gov/19452560/)

[23] [Structures of plant resistosome reveal how NLR immune receptors are activated - PMC](https://pmc.ncbi.nlm.nih.gov/articles/PMC9590527/)

[24] [Tsw – A case study on structure-function puzzles in plant NLRs with unusually large LRR domains](https://www.ncbi.nlm.nih.gov/pmc/articles/PMC9585916/)

[25] [The nucleotide binding domain of NRC-dependent disease resistance proteins is sufficient to activate downstream helper NLR oligomerization and immune signaling](https://www.biorxiv.org/content/10.1101/2023.11.30.569466v1)

[26] [RCSB PDB - 9FP6: Structure of the NbNRC2 hexameric resistosome](https://www.rcsb.org/structure/9fp6)

[27] [How to assess the quality of AlphaFold 3 predictions | AlphaFold](https://www.ebi.ac.uk/training/online/courses/alphafold/alphafold-3-and-alphafold-server/how-to-assess-the-quality-of-alphafold-3-predictions/)

[28] [Activation of the helper NRC4 immune receptor forms a hexameric resistosome](https://pubmed.ncbi.nlm.nih.gov/39094568/)

[29] [AlphaFold 3 predictions for paired NLR proteins - Obsidian Vault - Obsidian v1.8.7](https://zenodo.org/records/15552925/files/AlphaFold%203%20predictions%20for%20paired%20NLR%20proteins.pdf?download=1)

[30] [Structure of the activated Roq1 resistosome directly recognizing the pathogen effector XopQ](https://www.biorxiv.org/content/10.1101/2020.08.13.246413v1)

[31] [Structure-Aware Annotation of Leucine-rich Repeat Domains](https://www.biorxiv.org/content/10.1101/2023.10.27.562987v1)

[32] [RCSB PDB - 9RI9: Cryo-EM structure of the tomato NRC3 hexameric resistosome](https://www.rcsb.org/structure/9ri9)

[33] [A hydrophobic core in the coiled-coil domain is essential for NRC resistosome function](https://www.biorxiv.org/content/10.1101/2025.01.21.634219v2)

[34] [GeomeTRe: accurate calculation of geometrical descriptors of tandem repeat proteins](https://www.biorxiv.org/content/10.1101/2025.04.24.650440v1)

[35] [Predicted Aligned Error - Wikipedia](https://en.wikipedia.org/wiki/Predicted_Aligned_Error)

[36] [NLR receptors in plant immunity: making sense of the alphabet soup - PMC](https://pmc.ncbi.nlm.nih.gov/articles/PMC10561179/)

[37] [Evolution of NLR resistance genes with non-canonical N-terminal domains in wild tomato species](https://www.biorxiv.org/content/10.1101/786194v1)

[38] [(PDF) Structural Transition of NRC4 to Hexameric Resistosome Activates Plant Immunity](https://www.researchgate.net/publication/386157649_Structural_Transition_of_NRC4_to_Hexameric_Resistosome_Activates_Plant_Immunity)

[39] [AlphaFold accurately predicts distinct conformations based on the oligomeric state of a de novo designed protein - PMC](https://pmc.ncbi.nlm.nih.gov/articles/PMC9207892/)

[40] [AlphaFlex: Accuracy modeling of protein multiple conformations via predicted flexible residues | bioRxiv](https://www.biorxiv.org/content/10.1101/2025.07.11.664327v1.full-text)

[41] [Inverse problems with experiment-guided AlphaFold](https://arxiv.org/html/2502.09372v1)

**Motivation:**

$\def\mathcal#1{\mathit{#1}}\def\mathscr#1{\mathit{#1}}$

#### Motivation and Evidence

The motivation for the **Structural Novelty Index (SNI)** is to move beyond sequence-based motifs—such as the MADA motif—toward a **structure-aware functional annotation** of NLRs. The SNI leverages AlphaFold 3 (AF3) to automate the identification of unconventional NRCs across large datasets (e.g., ~6,000 Solanaceae sequences).

**Supporting Evidence and Explanatory Power:**

- **Identification of Decoy/Sensor Domains:** While cryptic repeats and integrated domains are known qualitatively, the **C-terminal Disordered Volume Ratio (CDVR)** provides a "missing quantitative metric ($V_{tail} / V_{LRR}$) to automatically identify them in high-throughput AlphaFold 3 screens without requiring manual domain annotation" (**Observation 3**).
- **Quantifying Non-Functionality:** The index explains the "death switch" failure in atypical NLRs like NRCX. It provides a diagnostic score to determine "why NRCX fails (e.g., an occluded aperture or low predicted BSA)... providing a diagnostic score for non-functional 'novelty' in the NRC family" (**Observation 4**).
- **Structural Compatibility Checks:** The **LRR Solenoid Curvature Index ($\Omega_{LRR}$)** serves as a "mathematical way to check if a specific LRR sequence is physically capable of 'wrapping around' a hexameric core," explaining why certain sequences cannot form canonical resistosomes (**Observation 5**).
- **Functional Proxying:** The hypothesis utilizes AF3 confidence scores (ipTM/pLDDT) "as a quantitative parameter to differentiate 'canonical' stable helpers from 'unconventional' sensors or modulators that fail to form high-confidence interfaces" (**Observation 7**).

#### Differentiating Arguments: Why Choose This Hypothesis?

The SNI is unique because it codifies the **biophysical requirements of oligomerization** (e.g., the HD1-WHD pivot and $\alpha$4-helix kink) into a multi-parametric filter.

- **Automation:** It is designed for high-throughput "script-based analysis to extract the seven parameters from the PDB/JSON outputs," which is significantly more efficient than manual structural inspection.
- **Structural Mechanics:** Unlike sequence-only searches, it focuses on "core structural transitions required for NLR activation—specifically the 'flip' of the NB-ARC module and the creation of a membrane-inserted pore."

#### Counterarguments

- **Inaccurate Numerical Benchmarks:** A critical flaw is that the hypothesis’s defined "canonical" ranges contradict established data. For example, it defines the canonical pore ($D_{pore}$) as 12–15 Å, while ground truth for NRCs is 17–19 Å (the 12 Å figure actually describes the *unconventional* ZAR1 pentamer).
- **Empirical Discrepancies:** Actual kink magnitudes ($\delta_{kink}$) in PDB structures range from 54° to 92°, far exceeding the hypothesis’s assumed range of 20°–30°. Similarly, pivot angles ($\theta_{pivot}$) in canonical hexamers show much higher variance (85°–107°) than the proposed 88°–90°.
- **Theoretical Validity:** The "Electrostatic Polarization Vector" is cited by reviewers as a non-standardized metric with no evidence of recognition in current NLR literature.
- **Statistical Rigidity:** The use of "fixed range rigidity" and "arbitrary '±10%' margins" fails to account for natural structural noise and the high covariance between parameters like pivot angles and Buried Surface Area (BSA).

#### Summary for Selection Decision

**Choose this hypothesis if** the goal is to establish a high-throughput, automated pipeline for structural discovery that transcends simple motif searches. Its strength lies in its **conceptual framework** and metrics like **CDVR**. **Do not choose this hypothesis in its current form** because its numerical foundation is "fundamentally flawed," and it would currently "misclassify 100% of canonical helpers as unconventional outliers" due to incorrect baseline benchmarks.

**Coherence:**

$\def\mathcal#1{\mathit{#1}}\def\mathscr#1{\mathit{#1}}$

No incoherence found.

**Deep verification:**

$\def\mathcal#1{\mathit{#1}}\def\mathscr#1{\mathit{#1}}$

Based on the provided claims and assumptions, the **Structural Novelty Index (SNI)** hypothesis faces several critical challenges that may render its current formulation incorrect. The reasons can be summarized into three main categories:

#### 1. Factually Incorrect Structural Benchmarks

The SNI relies on specific numerical thresholds to define "canonical" NRCs, but these thresholds contradict established structural data (e.g., PDB 9FP6):

- **Incorrect Pivot Angle Baseline:** The hypothesis defines the canonical HD1-WHD pivot angle ($\theta_{pivot}$) as 88–90°. However, scientific literature confirms the actual angle for canonical hexameric NRCs (like NRC2) is 85°. This shift in the baseline would cause the SNI to misclassify standard NRCs as "novel."
- **Inaccurate Pore Diameter Requirements:** The SNI claims a 12–15 Å diameter ($D_{pore}$) is required for canonical function. In reality, canonical hexameric resistosomes like NRC2 and NRC4 possess a wider pore of approximately 19 Å. The range proposed in the hypothesis (12–15 Å) actually describes pentameric structures like ZAR1, meaning the metric would incorrectly flag canonical helpers as unconventional.

#### 2. Methodological and Technical Flaws

The implementation of specific parameters within the computational framework is technically problematic:

- **Misapplication of Radius of Gyration ($R_g$):** The **C-terminal Disordered Volume Ratio (CDVR)** uses $R_g$ to approximate the volume of disordered tails. Because AlphaFold 3 (AF3) provides static models rather than a representative conformational ensemble of disordered regions, the $R_g$ value is highly sensitive to random seeds. This leads to inconsistent data and a high probability of false positives or negatives regarding structural novelty.
- **Static vs. Dynamic Modeling:** The reliance on static AF3 outputs to measure inherently flexible or disordered regions (like the C-terminal tails) fails to capture the biological reality of how these "unconventional" NLRs capture effectors.

#### 3. Logical and Operational Limitations

The workflow of the SNI suffers from inherent logical inconsistencies that limit its predictive power:

- **Circular Logic in Stoichiometry:** To calculate key metrics like the Pivot Angle and Pore Aperture, the researcher must first assume and model a specific stoichiometry (e.g., n=5 or n=6). Consequently, the SNI cannot "discover" novelty from a single sequence; it acts only as a comparative screening tool that confirms the researcher’s initial assumptions about the protein's stoichiometry.
- **Requirement for Multi-model Testing:** Because the index is not a direct calculation from a monomeric sequence, its success depends on testing various stoichiometry configurations ($n=5, 6, 7, etc.$) and comparing confidence scores (pTM/ipTM). This significantly increases the complexity and potential for bias in the classification process.
