## Supplementary Data S6 for "AI-guided discovery of atypical protein assemblies": [EXTERNAL] ├░Y╠ê┬o┬1⁄4 Co-scientist - Research overview 230897543 - Quantitative Structural Novelty Index (SNI) for NRC-NLR Proteins - Google confidential - For research purposes only.docx

##### 🧬 Co-scientist - Quantitative Structural Novelty Index (SNI) for NRC-NLR Proteins

**Research Goal**: • Goal: Define a quantitative Structural Novelty Index (SNI) consisting of 5–10 parameters that distinguish unconventional NRC-NLRs from canonical hexameric NRC resistosomes. • Rationale: To identify unconventional NLRs among the ~6,000 sequences in the NRC family, we must first establish a robust mathematical and structural baseline for "deviation" from the standard paradigm. This ensures that our AlphaFold 3 (AF3) screening process is grounded in specific structural biology metrics rather than broad sequence similarity. • Preferences: * Focus on parameters that can be extracted from PDB or AlphaFold models, such as protomer interface angles, N-terminal CC-domain variations, and specific residue conservation patterns in the NB-ARC domain. o Leverage structural insights from the provided Ground Truth Dataset (PDB IDs: 9FP6, 9RI9, and 9CC8). • Attributes: o Output: A list of 5–10 quantitative parameters defining the SNI. o Reference Set: Canonical NRC2 (NbNRC2), NRC3 (SlNRC3), and NRC4 hexameric resistosomes. o Scope: Identifying structural and sequence-based divergence from the ~350 Solanaceae species dataset.

**About**: This is an experimental system for generating novel and testable hypotheses. The hypotheses are generated by a model and may be wrong. For research purposes only.

**Table of contents:**

- Summary of Main Research Directions
- Review guidelines
- Open questions
- Unexpected connections
- Research contacts
- Top ranking hypotheses

##### Main Directions of Possible Research:

$\def\mathcal#1{\mathit{#1}}\def\mathscr#1{\mathit{#1}}$

### Research Overview: Defining a Structural Novelty Index (SNI) for Unconventional NRC-NLRs

#### 1. Summary of Main Research Directions

The research goal is to move beyond sequence-based phylogenetics to define a structural-mechanical baseline for the NRC (NLR-required for cell death) family. The research is divided into five main directions:

1. **Quaternary Geometry and Oligomeric Symmetry:** Establishing the mathematical bounds of the canonical hexameric assembly using inter-protomer rotation and pitch.
2. **Activation Mechanodynamics (The NB-ARC Hinge):** Quantifying the "death-switch" transition by measuring the reorientation of the NB-ARC subdomains (NB, HD1, WHD).
3. **The MADA-Helix Pore Architecture:** Defining the structural requirements for membrane-insertion by measuring the tilt, protrusion, and amphipathicity of the N-terminal CC-domain.
4. **Interface Energetics and Solvation:** Assessing the "glue" that holds the resistosome together through Buried Surface Area (BSA) and hydrophobic density.
5. **Autoinhibitory Solenoid Curvature:** Mapping the LRR (Leucine-Rich Repeat) arc to understand how recognition surfaces deviate in unconventional variants.

#### 2. Detailed Description of Main Research Directions

##### I. Quaternary Geometry and Oligomeric Symmetry

This direction focuses on the global arrangement of the 6-unit NRC resistosome. Canonical models (9FP6, 9RI9, 9CC8) show a highly symmetric, wheel-like structure. We will define parameters like the **Azimuthal Protomer Turning Angle ($\theta_{turn}$)** and **Axial Stair-stepping ($\Delta z$)**. By modeling the ~6,000 sequences into AlphaFold 3 (AF3) multimers, we can calculate how much a sequence "strains" against the hexameric constraint.

##### II. Activation Mechanodynamics (The NB-ARC Hinge)

The NB-ARC domain acts as the molecular engine. In canonical NRCs, activation involves a massive rotation of the Winged-Helix Domain (WHD) relative to the Nucleotide-Binding (NB) slot. This direction utilizes the **HD1-WHD Pivot Angle ($\Phi_{hinge}$)** to distinguish between "flipping" NLRs (canonical) and "non-flipping" or "stiff" variants that may function as decoys or sensors rather than executors.

##### III. The MADA-Helix Pore Architecture

The N-terminal CC-domain contains the MADA motif, which forms the membrane-permeabilizing pore. We focus on the **MADA $\alpha1$ Protrusion Height ($H_{\alpha1}$)** and **Pore Constriction Radius ($R_{pore}$)**. Deviations here indicate NLRs that might signal through protein-protein interactions rather than direct membrane lysis.

##### IV. Interface Energetics and Solvation

The stability of the resistosome depends on specific contact points between the NB-ARC of one protomer and the LRR/CC of the next. We will quantify the **Interface Buried Surface Area (iBSA)** and **Hydrophobic Density ($I_H$)**. This direction seeks to identify "symmetry breakers" where the interface is chemically incompatible with hexamerization.

##### V. Autoinhibitory Solenoid Curvature

The LRR domain shields the NB-ARC in the inactive state. We will measure the **LRR Tangential Angle ($\Omega_{LRR}$)** and solenoid radius. Unconventional NRCs often have truncated or extended LRRs, which alters the "reach" of the domain and its ability to maintain the "off" state.

#### 3. Rationale for Research Areas

##### Why Research Quaternary Geometry?

Recent findings in cryo-EM (e.g., the NRC2/4 structures) show that the hexameric state is a strict requirement for calcium-channel activity. However, sequence diversity suggests some NRCs might form pentamers or octamers. Researching this area allows us to move from *assuming* a function to *predicting* a quaternary state based on spatial fit. If a sequence cannot mathematically close a hexameric ring without steric clashing, it is a high-priority "unconventional" candidate.

##### Why Research the NB-ARC Hinge?

The "death-switch" is the defining moment of plant immunity. Arguments for exploring this area include the discovery of "Integrated Decoys" where the NB-ARC is modified to stay "on" or "off" regardless of ATP binding. Understanding the mechanical limits of the HD1-WHD pivot helps us identify NRCs that have lost their catalytic power but gained a new signaling role.

#### 4. Specific Research Topics and Questions

##### Topic: The Isosteric Polarity Shift ($\chi_{iso}$)

- **Questions:** Do unconventional NRCs swap hydrophobic residues for polar ones at the NB-NB interface? Does this prevent the "locking" mechanism seen in PDB 9FP6?
- **What we could learn:** We can identify NRCs that are "monomeric-only" sensors that must heterodimerize with a helper rather than forming homohexamers.

##### Topic: MADA-Funnel Pitch and Depth

- **Questions:** Is the angle of the MADA helix relative to the pore axis conserved across all 350 Solanaceae species? Does a "shallower" funnel prevent membrane insertion?
- **What we could learn:** We could discover a sub-class of "signaling NRCs" that look like helpers but lack the physical "depth" to pierce the plasma membrane.

#### 5. Detailed Topics and Examples

##### Parameter 1: Azimuthal Protomer Turning Angle ($\theta_{turn}$)

- **Why Research:** This defines the "tightness" of the ring.
- **Idea/Example:** Compare the $\theta_{turn}$ of NbNRC2 (Ground Truth) to an outlier in the *Solanum lycopersicum* dataset. If the angle deviates by >5°, AF3 might predict a "broken ring," suggesting an unconventional stoichiometry like a heptamer.

##### Parameter 2: HD1-WHD Hinge Angle ($\Phi_{hinge}$)

- **Why Research:** It measures the displacement of the "latch" that releases the CC-domain.
- **Idea/Example:** Map the distance between the P-loop and the WHD. In 9CC8 (SlNRC3), this distance increases upon activation. An unconventional NLR might show a "pre-shifted" WHD, suggesting it is a "constitutively primed" sensor.

##### Parameter 3: MADA $\alpha1$ Protrusion Height ($H_{\alpha1}$)

- **Why Research:** The MADA helix must extend far enough to reach the membrane.
- **Idea/Example:** Measure the vertical displacement of the $\alpha1$ helix relative to the NB-ARC plane. NRCs with a "short" $H_{\alpha1}$ may have evolved to target internal organelles (like the chloroplast) rather than the plasma membrane.

##### Parameter 4: Interface Buried Surface Area (iBSA)

- **Why Research:** High iBSA correlates with stable, "death-inducing" resistosomes.
- **Idea/Example:** Calculate the iBSA for NRC4 (9RI9). Use this as a Z-score baseline. Sequences with an iBSA < 1500 Å² are likely transient interactors rather than stable pore-formers.

##### Parameter 5: LRR Solenoid Curvature ($\kappa_{LRR}$)

- **Why Research:** The curve of the LRR determines the size of the ligand-binding pocket.
- **Idea/Example:** Model the LRR as a geometric arc. Unconventional NRCs might show a "flattened" arc, indicating they have evolved to bind larger, multi-protein effector complexes instead of small small-molecule triggers.

#### 6. Summary of Parameters for the SNI (The Final List)

Based on the research above, the 10 quantitative parameters defining the **Structural Novelty Index (SNI)** are:

1. **$\theta_{turn}$:** The inter-protomer azimuthal rotation angle (Canonical: ~60°).
2. **$\Phi_{hinge}$:** The angular displacement between HD1 and WHD subdomains.
3. **$H_{\alpha1}$:** Vertical protrusion height of the MADA helix from the resistosome plane.
4. **$R_{pore}$:** The minimum constriction radius of the N-terminal funnel.
5. **iBSA:** Total buried surface area at the protomer-protomer interface.
6. **$\theta_{kink}$:** The magnitude of the kink in the $\alpha4$-helix of the NB-ARC.
7. **$d_{B-MHD}$:** Distance between the P-loop (BeF3/ATP site) and the MHD motif.
8. **$\chi_{iso}$:** A ratio of hydrophobic to polar residues at the core interface.
9. **$\Omega_{LRR}$:** The tangential angle of the LRR relative to the NB-ARC base.
10. **$\vec{A}_{MADA}$:** The magnitude of the amphipathic vector of the first 21 residues.

#### 7. Unexpected Areas for Research

- **Non-Pore-Forming "Decoy" NRCs:** Some sequences might have perfect NB-ARC geometry but "filled" pores. Researching the **Pore Cavity Occupancy Index** could reveal NRCs that act as "sponges" for pathogen effectors, neutralizing them by sequestering them inside a non-functional hexamer.
- **Asymmetric Resistosomes:** While canonical models are symmetric, some NRCs might form **"Gapped Hexamers"** or C-shaped structures. Investigating **N-to-C Terminal Displacement** could uncover NLRs that function as scaffolds for other immune proteins (like kinases) rather than acting as independent ion channels. This would redefine the NRC family from being just "helpers" to being "immune hubs."

##### Review guidelines:

$\def\mathcal#1{\mathit{#1}}\def\mathscr#1{\mathit{#1}}$

##### Review Guidelines for the Structural Novelty Index (SNI)

###### 1. Factual Accuracy of Ground Truth Baselines

- **PDB Consistency:** Reviewers must verify that the proposed numerical baselines for "canonical" structures match the actual data in PDB IDs 9FP6, 9RI9, and 9CC8. For example, 9RI9 is a resting-state dimer, not an active hexamer; using it as a reference for a "flat hexamer" metric is a fatal flaw.
- **The "Missing Residue" Problem:** Many experimental structures (like the ground truth set) lack coordinates for the N-terminal MADA motif (residues 1–15/20) because they are disordered. Ideas that propose measuring MADA "protrusion" or "solvent accessibility" against experimental PDBs as a baseline must explain how they account for these missing residues (e.g., using AF3-generated models of NRC2/3/4 as the true baseline).

###### 2. Technical Feasibility & Computational Scalability

- **Tiered Screening Strategy:** Modeling 6,000 homo-hexamers in AlphaFold 3 is computationally prohibitive (thousands of GPU hours). A "good" idea must incorporate a phased approach—such as using monomeric metrics (mSNI) to filter the dataset before proceeding to expensive multimeric simulations (cSNI).
- **Memory and Token Limits:** Reviewers should flag ideas that ignore the hardware constraints of modeling large complexes (~5,000+ residues). Proposals must address whether the full sequence or truncated domains (e.g., CC-NB-ARC) are being used for the high-throughput phase.

###### 3. Mitigation of AI Modeling Bias ("Forced Folding")

- **Detection of "Structural Frustration":** AlphaFold 3 may attempt to "force" a sequence into a hexameric shape even if the protein naturally prefers a different stoichiometry. An effective SNI must include metrics that detect this "strain," such as interface shape complementarity ($S_c$), inter-protomer angular deviation, or axial "stair-stepping."
- **Stoichiometric Contrast:** High-quality ideas should suggest "contrastive modeling"—comparing the confidence/fit of a sequence when modeled as a pentamer, hexamer, and heptamer—rather than assuming the hexameric model is biologically correct.

###### 4. Integration of Confidence Metrics

- **Biological Signal vs. Computational Noise:** Geometric deviations (e.g., a $15^\circ$ shift in NB-ARC) are only meaningful if the model confidence is high. Reviewers must ensure the SNI incorporates AF3 confidence scores like **ipTM** (interface predicted Template Modeling) and **pLDDT**.
- **Penalty for Low Confidence:** A "novel" structure with low ipTM/pLDDT should be flagged as a potential modeling failure rather than a "novel" discovery.

###### 5. Robustness of Automated Parameters

- **Anchor-Based vs. Index-Based Numbering:** Plant NLRs are highly diverse with frequent insertions/deletions. Reviewers should reject ideas that rely on absolute residue numbering (e.g., "distance between Res 10 and Res 150"). Instead, parameters should be anchored to conserved motifs (e.g., P-loop, MHD motif, or the $\alpha 1$ hydrophobic core).
- **Geometric Nuance over Global RMSD:** Simple RMSD is too blunt a tool. Look for parameters that capture specific mechanistic shifts, such as the **NB-ARC swivel angle** ($10^\circ$ shift), the **protomer rotation angle** ($60^\circ$ for hexamers vs $72^\circ$ for pentamers), and **pore aperture radius**.

###### 6. Functional Relevance of the CC-Domain

- **Pore-Forming Logic:** Since the NRC family consists of "helpers" (pore-formers) and "sensors" (signaling decoys), the SNI must include metrics specifically targeting the executioner function, such as the planarity of the $\alpha 4$ helix or the hydrophobic moment alignment of the MADA motif.

###### 7. Statistical Rigor

- **Reference Set Stability:** Calculating Z-scores or Mahalanobis distances based on only 3–4 ground truth structures is statistically sensitive. Reviewers should look for methods that account for the natural variance within the canonical NRC2/3/4 group to avoid over-sensitivity where minor biological fluctuations are flagged as "novelty."

##### Open questions:

$\def\mathcal#1{\mathit{#1}}\def\mathscr#1{\mathit{#1}}$

Based on the provided goal to define a **Structural Novelty Index (SNI)** and the supporting structural data (PDB IDs 9FP6, 9RI9, 9CC8), here is the analysis of patterns in claims and research directions.

##### 1. Broad and Relevant Open Questions for the Goal

1. **What specific quantitative parameters, such as protomer interface angles or distances between domains, most reliably differentiate unconventional NRC-NLRs from standard hexameric forms?** (Highest importance for defining the SNI).
2. **What structural determinants or inter-domain angles within NLR protomers dictate whether a resistosome will assemble as a pentamer or a hexamer?** (Crucial for establishing the "deviation" baseline).
3. **To what extent can the HD1-WHD pivot angle in structural models be utilized as a universal predictor for the oligomeric stoichiometry?** (A specific, measurable metric for the SNI).
4. **How can structural deformation metrics used in AlphaFold assessments (LDDT, ipTM, RMSD) be specifically calibrated to define an SNI for the NRC family?** (Addresses high-throughput screening needs).
5. **How can N-terminal CC-domain variations (e.g., the α4-helix kink) be represented as robust mathematical metrics for screening the NRC family?** (Focuses on a required structural preference).
6. **What specific residue conservation patterns (e.g., Shannon entropy in NB-ARC) correlate with structural divergence from the standard hexameric paradigm?** (Integrates sequence and structure).
7. **How does the identified kink in the α4-helix of the CC domain in specific NLRs like NbNRC2 influence the packing and stoichiometry of the resulting resistosome?** (Mechanical basis for hexamerization).
8. **Can the structural 'deviation' be mathematically defined by the relative orientation and tilt of the MADA motif residues in relation to the central axis?** (Specific N-terminal metric).
9. **What structural features explain why certain helper NLRs form dodecameric (dumbbell) states while others remain hexameric?** (Identifies higher-order structural novelty).
10. **How do variations in domain architecture (absence of CC, truncated LRR) impact the formation of the hexameric resistosome?** (Identifies gross structural outliers).

##### 2. Analysis of Key Questions and Research Directions

###### **Q1: Which parameters most reliably differentiate unconventional NRCs?**

- **Known:** Hexameric NbNRC2 has a wider NB-ARC interdomain angle (85°) than pentameric ZAR1 (75°) and wider pore diameters (14 Å vs 13 Å).
- **Unknown:** The specific numerical range of these angles across all 6,000 NRC sequences.
- **Research Direction 1 (Interdomain Angles):** Quantify the CC-NB-ARC interdomain angle across AF3 models. Claims show that a 10° widening is necessary to accommodate the sixth protomer in NRC2.
- **Research Direction 2 (Buried Surface Area):** Use PISA to calculate Interface Buried Surface Area (BSA). Claims state canonical NRC interfaces involve ~1,874 Å² for dimers and ~4,980 Å² for octameric unconventional variants.
- **Unexpected Direction:** Monitoring the "internal cavity volume." Claims suggest hexamers like NbNRC2 possess a significantly larger cavity between LRR/WHD/NB domains to allow deeper insertion of adjacent HD1 domains.

###### **Q2: Determinants of Pentamer vs. Hexamer Assembly?**

- **Known:** The HD1-WHD angle relative to the central helix is ~88° in hexamers (NRC2/4) but 92° in pentamers (ZAR1).
- **Unknown:** If minor amino acid substitutions in the HD1-WHD hinge can force a transition between these states.
- **Research Direction 1 (HD1-WHD Angle Mapping):** Screen the 6,000 sequences for the 88° vs 92° threshold. Claims identify this angle as a primary predictor of stoichiometry.
- **Research Direction 2 (Pivot Helix Analysis):** Analyze the amphipathic pivot helix (residues 479–491). Claims suggest side-chain size and packing density in this hydrophobic pocket modulate the hinge angle.
- **Unexpected Direction:** Investigating "Dodecameric Symmetry." Claims note NRC4 can form dumbbell-shaped dodecamers, a significant "novelty" deviation from the single-layer wheel.

###### **Q3: Representing CC-domain variations as mathematical metrics?**

- **Known:** NbNRC2 features a unique "kink" in the α4-helix near residue L126, which prevents steric overlap in the hexamer.
- **Unknown:** How to mathematically express "kinkiness" for automated screening.
- **Research Direction 1 (Helix Curvature):** Calculate the local bend angle of the α4-helix. Claims state this kink is absent in pentameric resistosomes, making it a binary novelty switch.
- **Research Direction 2 (Pore Constriction Radii):** Measure the radius at Gln126. Claims indicate NRC4 has a 4.3 Å radius, while unconventional octamers reach 12.7 Å.
- **Unexpected Direction:** Environmental Modeling. Claims suggest simulating AF3 models with oleic acid better resolves the CC-funnel geometry for more accurate metric extraction.

###### **Q4: Sequence patterns in NB-ARC indicating divergence?**

- **Known:** Specific residues (D239 in NRC2) mediate inter-protomer contacts. The MADA motif HMM score distinguishes helpers (HMM > 10) from sensors.
- **Unknown:** Which entropy shifts in the NB-ARC "dimerization stretches" predict a fail in hexamerization.
- **Research Direction 1 (Shannon Entropy Mapping):** Apply a threshold (e.g., 1.5 bits) to the eight dimerization stretches. Claims show these positions are highly variable across the superfamily but conserved within canonical clades.
- **Research Direction 2 (HMM Profiling):** Use the Hidden Markov Model score of the MADA motif. Claims show NRC4 (31.5) and NRC2 (24.5) have high scores, while unconventional NRCX (22.2) is functionally degenerated.
- **Unexpected Direction:** Monitoring "MHD-P-loop Distance." Claims suggest the spatial orientation between these motifs in AF3 models predicts the ability to cycle between ADP/ATP states.

##### 3. Patterns in Research Directions

- **Geometric Determinism:** A clear pattern exists where stoichiometry (5x vs. 6x vs. 8x) is dictated by specific, measurable angles: the HD1-WHD pivot angle (~88°-92°) and the CC-NB-ARC interdomain angle (75° vs. 85°).
- **Interface Energetics:** Buried Surface Area (BSA) and inter-protomer salt bridges (e.g., E283-K329 in NRC4) are recurring metrics for stability.
- **Functional "Kinks":** A repeated theme is the presence of structural "distortions" (the α4-helix kink, the MADA "flip-out") as prerequisites for canonical assembly.
- **Unexpected Pattern (Confidence as Metric):** Researchers are using AI confidence scores (ipTM/pTM) not just as validation, but as *quantitative parameters* themselves (e.g., ipTM > 0.7 for canonical hexamers).

##### 4. Reasoning on Most Promising Directions

The most promising approach for the SNI is a **multi-layered filter**. First, utilize high-throughput **sequence-based HMM scores** to filter the 6,000 sequences. Second, employ **AF3 modeling** to extract the **HD1-WHD interdomain angle**, as it appears to be the most "universal" predictor of stoichiometry across different NLR families. Finally, verify "canonicalness" by measuring the **α4-helix bend angle**, which provides a binary structural check for hexamer-compatibility.

##### 5. Summary of Promising Directions

**Summary of promising directions:**

1. **HD1-WHD Interdomain Angle Mapping:** This is the most promising metric because it distinguishes pentamers (92°), hexamers (88°), and octamers (80°) with high precision. It serves as a geometric "fingerprint" for stoichiometry, allowing researchers to predict the oligomeric state of the 6,000 NRC sequences directly from AF3 models.
2. **MADA Motif HMM Scoring:** Establishing an HMM score baseline (e.g., >24.5 for NRC2/4) allows for rapid sequence-level identification of "functional helpers." Claims show that even minor variations (like the Thr-4 polymorphism in NRCX) can drop the functionality while maintaining high sequence similarity, making HMM a sensitive novelty detector.
3. **CC-α4 Helix Curvature (Kink Detection):** Measuring the bend near residue L126 is critical. Since a straight α4-helix causes steric overlap in hexamers, the "kink" is a mandatory structural parameter for the canonical NRC2/3/4 paradigm, distinguishing it from ZAR1-like pentamers.
4. **Interface Predicted Template Modeling (ipTM) Thresholds:** Setting a mathematical cutoff (e.g., ipTM > 0.70) in AF3 models effectively separates stable canonical assemblies from "unconventional" or non-functional ones. Claims show that helper NLRs consistently yield higher ipTM scores than sensor NLRs when modeled as hexamers.
5. **Pore Constriction Radius Calculation:** Measuring the minimum radius at the CC-pore (e.g., 4.3 Å for NRC4 vs. 12.7 Å for octamers) provides a functional metric for ion-channel capacity. This distinguishes canonical "death switches" from wider, unconventional pore architectures.
6. **Shannon Entropy of Dimerization Stretches:** By calculating the entropy of the eight identified NB-LRR and LRR-LRR interface stretches, researchers can quantify "molecular insulation." Canonical NRCs show low entropy in these clades, while unconventional ones diverge to avoid cross-activation.
7. **NB-ARC Interdomain Module Widening:** The 10° widening (85° in NRC2 vs 75° in ZAR1) of the CC-NB-ARC module is a robust structural biology metric for "deviation." It explains the physical space required for the 6th protomer and can be extracted using UCSF Chimera.
8. **Internal Cavity Volume Measurement:** The volume formed between LRR, WHD, and NB domains is substantially larger in hexameric NRCs. This parameter determines the depth of adjacent HD1 domain insertion, providing a volumetric baseline for the hexameric resistosome.
9. **Buried Surface Area (BSA) Quantitation:** Utilizing PISA to measure the BSA (standardizing at ~1,874 Å² for resting dimers) provides a energetic baseline for autoinhibition. Unconventional NRCs that lack this specific buried area are likely "trigger-happy" or non-canonical.
10. **α1-Helix Orientation and Tilt:** Measuring the angle of the α1-helix relative to the central pore axis distinguishes the "funnel" geometry of activated resistosomes. Claims suggest that the inability to form this stable funnel is a primary indicator of a "sensor" or "unconventional" NLR.

##### Unexpected connections:

$\def\mathcal#1{\mathit{#1}}\def\mathscr#1{\mathit{#1}}$

Unexpected connections:

Unexpected connection 1:

- Claims: "The L126 "kink" in the α4-helix of NbNRC2 is essential for hexameric stoichiometry by preventing steric overlap." "NRC4 (9CC8) uses residue Q126 as the primary constriction point for the active pore." "The LRR tangent angle relative to the wheel tangent is smaller in NRC4 to allow dense packing."
- Reasoning: There is a multi-scalar geometric coupling between the internal torsion of the NB-ARC domain and the global stoichiometry of the resistosome. The "kink" at residue 126 is not merely a local structural feature; it acts as a stoichiometric gatekeeper. If the α4-helix is "straightened" (reducing the torsion angle), the steric overlap at the protomer interface forces a transition from a hexamer to a pentamer. Simultaneously, this same residue defines the pore diameter. This implies that the LRR's tangential pitch (which "pushes" the NB-ARC) and the 126-kink angle are co-dependent variables that can be used to mathematically predict stoichiometry.
- Novelty: While the 126-kink and LRR packing are known, the synthesis of these into a "Torsion-Tangent Ratio" to distinguish between pentameric (ZAR1-like), hexameric (NRC-like), and octameric (WAI3-like) states in high-throughput AF3 models is novel.
- Relevance: Parameter 1 of the SNI: **α4-Helix Torsion Angle ($\tau_{126}$)**. A deviation of $>15^\circ$ from the 9FP6 baseline indicates an unconventional (non-hexameric) assembly.

Unexpected connection 2:

- Claims: "NRCX has a MADA HMM score of 22.2 (higher than functional ZAR1), yet lacks cell death capacity." "NRCX is a functionally degenerated MADA-NLR." "The α1-helix is frequently invisible or disordered in inactive structural models."
- Reasoning: This identifies a "Sequence-Structure Decoupling" in unconventional NRCs. A high MADA HMM score (sequence identity) is insufficient for function if the structural "activation radius"—the ability of the α1-helix to fold into a stable, high-pLDDT funnel in AF3—is compromised. Unconventional modulators like NRCX mimic the helper sequence to maintain sensor-binding compatibility but are structurally "dead-switched" at the funnel-formation stage.
- Novelty: Unexpected. Traditional screening assumes sequence conservation in the MADA motif implies executor function. Using the ratio of HMM score to AF3 structural confidence (pLDDT) as a metric for "functional degeneration" is a new structural biology approach.
- Relevance: Parameter 2 of the SNI: **MADA Functional Divergence Ratio (FDR)**. Calculated as (HMM Score) / (α1-Funnel pLDDT). High FDR ($>0.8$) identifies unconventional modulators/sensors.

Unexpected connection 3:

- Claims: "HD1-1 region (residues 316-317) is a binding site for inhibitors like SS15." "The NB-HD1 domains undergo a 180-degree rotation relative to WHD-LRR during activation." "WAI3 octameric stoichiometry is defined by the HD1–WHD angle."
- Reasoning: The HD1-1 loop is a "functional nexus" where pathogen inhibition and stoichiometric determination overlap. The "Hinge Flex" (the angle between HD1 and WHD) determines if the resistosome can accommodate the wider curvature required for an octamer. Pathogen inhibitors (SS15) likely stabilize this hinge in a "closed" or "locked" configuration. Unconventional NRCs may evolve specific HD1-1 mutations that "stiffen" the hinge, not just to evade inhibitors, but to favor alternative stoichiometries.
- Novelty: Known in isolation, but the use of the HD1-WHD interdomain angle as a quantitative "Stoichiometry Dial" for AF3 screening is a novel application.
- Relevance: Parameter 3 of the SNI: **HD1-WHD Hinge Angle ($\theta_{Hinge}$)**. Canonical NRCs cluster at $88^\circ \pm 2^\circ$. Values $<82^\circ$ suggest octameric potential (WAI3-like).

Unexpected connection 4:

- Claims: "A 9-amino acid indel in the CC-NB-ARC linker of PinNRC3." "NRC4 forms a dumbbell-shaped dodecamer (210 Å height)." "The α1-helices associate internally to bridge two hexamers in the dodecamer."
- Reasoning: The "Linker-Tether" constraint. The length of the CC-NB-ARC linker defines the "displacement radius" of the CC domain. A specific linker length is required for the α1-helices to reach and bridge with a second hexamer. Indels in this region (like the 9-aa deletion) likely act as a "reach-switch," determining whether a sequence forms a single-layer hexamer or a double-layered dodecamer.
- Novelty: Unexpected. Indels are typically viewed as compatibility determinants for sensors, not as mechanical regulators of higher-order (dodecameric) stacking.
- Relevance: Parameter 4 of the SNI: **Linker Displacement Ratio ($L_{DR}$)**. (Linker Length / Protomer Height). Deviations from the 9FP6/9CC8 baseline suggest unconventional dodecameric stacking or "tethered" non-functional funnels.

Unexpected connection 5:

- Claims: "Residue Interaction Networks (RINs): Stress and Betweenness Centrality." "Shannon Entropy (measuring amino acid diversity per site)." "Unconventional NRCs show high sequence identity (80-100%) in sensor repeats."
- Reasoning: **Centrality-Entropy Decoupling (CED)**. In canonical NRCs, high-centrality residues (structural core) are evolutionarily "frozen" (low entropy). In unconventional NRC-sensors, we expect to see high-centrality nodes with high Shannon entropy, indicating residues that are structurally vital but under diversifying selection for novel pathogen-effector perception.
- Novelty: Not known. This synthesis of network topology and evolutionary dynamics is a novel method for identifying "subfunctionalized" NLRs.
- Relevance: Parameter 5 of the SNI: **CED Score**. Identifies residues where (Centrality Z-score $>2$) and (Shannon Entropy $>1.5$). High CED counts distinguish unconventional sensors from canonical helpers.

##### **Structural Novelty Index (SNI) for NRC-NLRs**

The following 8 parameters constitute the SNI, derived from the connections above and ground truth models (9FP6, 9RI9, 9CC8):

| # | Parameter | Metric | Canonical Baseline | Unconventional Deviation |
| --- | --- | --- | --- | --- |
| 1 | **$\alpha$4-Torsion ($\tau_{126}$)** | Angle ($^\circ$) at Res. 126 | $15^\circ$–$25^\circ$ (Kinked) | $<10^\circ$ (Straight/Pentameric) |
| 2 | **Hinge Angle ($\theta_{Hinge}$)** | HD1-WHD Angle ($^\circ$) | $88^\circ \pm 2^\circ$ | $<82^\circ$ (Octameric) or $>92^\circ$ |
| 3 | **FDR (MADA)** | HMM / $\alpha$1-pLDDT | Ratio $0.3 - 0.5$ | $>0.8$ (Sequence-Structure Mismatch) |
| 4 | **$\Delta$-ipTM** | AF3 6x vs. 5x/8x | Delta $>0.2$ | Delta $<0.1$ (Stoichiometric Plasticity) |
| 5 | **CED Score** | RIN Centrality vs. Entropy | Low Decoupling | High Decoupling (Sensor Specialty) |
| 6 | **Linker Ratio ($L_{DR}$)** | Linker Length / CC Height | $\sim 1.0$ | $\pm 0.3$ (Stacking/Tethering Novelty) |
| 7 | **Interface $\Delta$G** | MM/GBSA (NB-LRR) | $<-1.0$ kcal/mol | $>-0.2$ kcal/mol (Trigger-Happy) |
| 8 | **LRR Tangent ($\theta_{LRR}$)** | Angle relative to wheel | Compressed | Extended (Novel PPI Surfaces) |

##### Research contacts:

$\def\mathcal#1{\mathit{#1}}\def\mathscr#1{\mathit{#1}}$

##### **Research Direction 1: High-Throughput AF3 Modeling and Metric Extraction**

This direction focuses on the computational pipeline required to process ~6,000 sequences, using AlphaFold 3 (AF3) confidence scores and geometric outputs to populate the Structural Novelty Index (SNI).

- **Researchers:**
  - **AmirAli Toghani (The Sainsbury Laboratory):** A lead expert in using AF3 to distinguish between sensor and helper NLR configurations. He specializes in analyzing AF3 confidence metrics (ipTM, pTM) and "funnel-shaped" structural predictions.
  - **Sophien Kamoun (The Sainsbury Laboratory):** Pioneer of the NRC network framework. His group developed the methodology for triaging NLRs into functional categories using AF3 modeling in the presence of lipid proxies (oleic acid).
  - **Josh Abramson (Google DeepMind/AF3 Team):** While primarily a developer, his expertise is essential for extracting advanced metrics like Predicted Aligned Error (PAE) and inter-domain distances from AF3 outputs at scale.
- **Why they are best for this direction:** These researchers have already established the benchmark for using AF3 to categorize NLRs. They can define the "Isosteric Polarity Shift" ($\chi_{iso}$) and "ipTM variance" needed to flag sequences that fail to form a canonical hexameric funnel. They are uniquely positioned to bridge the gap between raw sequence data and the initial SNI score.
- **Example Hypothesis Titles:**
  - *Automated AF3-Driven Triaging of 6,000 NRC Sequences via ipTM and PAE Variance Analysis.*
  - *A Weighted Z-Score Model for NRC Heterogeneity Based on AF3 Interaction Confidence Scores.*

##### **Research Direction 2: Experimental Structural Validation and Stoichiometry**

This direction involves the physical validation of structural outliers identified by the SNI. These researchers can run experiments to determine if an unconventional SNI score correlates with non-hexameric assembly or altered activation.

- **Researchers:**
  - **Jogi Madhuprakash (The Sainsbury Laboratory):** The primary expert for PDB 9FP6 (NbNRC2). He has experience comparing hexameric NRCs to pentameric ZAR1 and can run the cryo-EM and biochemistry experiments needed to validate novel stoichiometries.
  - **Michael W. Webster (The Sainsbury Laboratory):** A senior structural biologist specializing in cryo-EM reconstructions and biochemical stabilization of NLR complexes. He is best suited to validate "NB-ARC vertical planarity" and "inter-protomer pitch angles."
  - **Furong Liu & Brian J. Staskawicz (UC Berkeley):** Lead researchers for PDB 9CC8 (NRC4). They are experts in interface mutagenesis and determining how specific residue patterns govern hexameric versus dodecameric (dumbbell) assemblies.
- **Why they are best for this direction:** These researchers provided the "Ground Truth" data (9FP6, 9CC8). They possess the established experimental pipelines (cryo-EM, mass photometry, AUC) to prove that a sequence with a high SNI score actually forms a unique structure (e.g., a pentamer, octamer, or decoy dimer).
- **Example Hypothesis Titles:**
  - *Cryo-EM Validation of Non-Canonical NB-ARC Swivel Angles in SNI-Flagged NRC Outliers.*
  - *Mapping the Correlation Between SNI Assembly Symmetry Metrics and In Vitro Oligomeric Stoichiometry.*

##### **Research Direction 3: N-terminal "Death-Switch" and Membrane Pore Mechanics**

This direction focuses on the executioner domain (CC-domain) and the MADA motif, specifically parameters related to the pore-forming funnel.

- **Researchers:**
  - **Hiroaki Adachi (The Sainsbury Laboratory):** Discovered the MADA motif. He is the best candidate to validate Parameter 1.2 ($\text{RMSD}_{MADA}$) and identify if unconventional N-terminal variations indicate a loss of pore-forming function (e.g., decoys like NRCX).
  - **Tolga O. Bozkurt (Imperial College London):** Expert in NLR-membrane interactions. He can run experiments (e.g., liposome leakage assays, confocal microscopy) to validate if SNI parameters like "Pore Constriction Aperture" ($D_{APEX}$) accurately predict membrane-disrupting activity.
  - **Hung-Yu Wang (The Sainsbury Laboratory):** Specialist in the hydrophobic core of the CC-domain. He is best suited to evaluate the "Interface Hydrophobic Density Score" ($I_{H}$) and how it affects resistosome stability.
- **Why they are best for this direction:** They focus on the *functional output* of the structure. If the SNI identifies a sequence with an unconventional $\alpha1$-helix tilt, these researchers can experimentally determine if that protein still triggers cell death or inserts into the plasma membrane.
- **Example Hypothesis Titles:**
  - *Functional Characterization of Non-Pore-Forming NRC Variants via N-terminal MADA Hydrophobicity Profiling.*
  - *Experimental Verification of SNI-Predicted Pore Constriction Radii using Liposome Permeability Assays.*

##### **Research Direction 4: Evolutionary and Genomic SNI Mapping**

This direction applies the SNI to the vast Solanaceae dataset to identify clades of unconventional NLRs and understand their evolutionary trajectory.

- **Researchers:**
  - **Chih-Hang Wu (Academia Sinica):** Expert in the phylogenomic analysis of the NRC family across thousands of Solanaceae sequences. He can integrate SNI scores with phylogenetic trees to identify clades where structural "deviations" became fixed.
  - **Mauricio P. Contreras (The Sainsbury Laboratory):** Expert in the evolution of the lettuce NRC network and the "activation-and-release" model. He can bridge structural SNI metrics with the divergence of sensor and helper NLR roles.
  - **Lida Derevnina (University of Cambridge):** Specialist in the hierarchical organization of NRC networks. She can investigate if SNI outliers represent specialized NLRs that have evolved to interact with specific pathogen effectors.
- **Why they are best for this direction:** These researchers understand the "6,000 sequences" context. They can validate if the SNI is robust enough to categorize entire subclades (e.g., NRC0, NRCX, or NRC-sensors) as unconventional versus canonical.
- **Example Hypothesis Titles:**
  - *Phylogenetic Mapping of the SNI Across 350 Solanaceae Species to Identify Ancestral Stoichiometric Transitions.*
  - *Evolutionary Divergence of NB-ARC Latch Distances in SNI-Categorized Sensor versus Helper NLRs.*

##### Top ranking hypotheses:

- **180708842503749: Developing an 8-Parameter Dual-Phase SNI for Unconventional NRC-NLR Identification via AlphaFold 3**

  $\def\mathcal#1{\mathit{#1}}\def\mathscr#1{\mathit{#1}}$
  The proposed Structural Novelty Index (SNI) is a two-phase quantitative framework designed to identify unconventional NRC-NLRs by measuring their structural deviation from canonical hexameric resistosomes (NRC2, NRC3, and NRC4). The index utilizes eight specific parameters: a monomeric phase (mSNI) filters candidates based on N-terminal helix propensity, NB-ARC swivel angles, "latch" proximity, and LRR expansion, while a complex-level phase (cSNI) validates novel assemblies through protomer dihedral angles, hydrophobicity gradients, interface burial efficiency, and stoichiometric preference scores. By applying a weighted Z-score formula to compare AlphaFold 3 (AF3) predictions against established ground truths, this workflow creates a scalable screening funnel that efficiently distinguishes non-canonical architectures—such as alternative stoichiometries or novel auto-regulatory states—from the broader NRC family.
  Details: [document](https://docs.google.com/document/d/1fNNInMSn-AOYfxuy9tYIaaEPaQPq_MlrAfJq7zYJ4co/edit).
- **114616009328010: An 8-Parameter Integrated Structural Novelty Index (SNI) for Identifying Unconventional NRC Resistosomes via AlphaFold 3**

  $\def\mathcal#1{\mathit{#1}}\def\mathscr#1{\mathit{#1}}$
  The Integrated Structural Novelty Index (SNI) is a quantitative bioinformatic framework designed to identify unconventional NRC-NLRs by measuring structural divergence from the canonical hexameric resistosome paradigm. Utilizing AlphaFold 3 (AF3) modeling, the SNI employs eight distinct parameters—including canonical compliance scores, MADA bundle convergence, protomer packing ratios, and inter-domain angles—to assess the feasibility of a sequence adopting the standard fold established by ground-truth structures like NbNRC2. By evaluating specific geometric and energetic metrics such as the $\alpha$4-helix curvature and pore electrostatic flux, the SNI enables a high-throughput screening of approximately 6,000 NRC sequences to classify outliers as potential octamers, pentamers, decoys, or novel ion channels for subsequent experimental validation.
  Details: [document](https://docs.google.com/document/d/1L8jDkLwYyp3vQlgW7H2Yu2arDMV70sH5QgDTQBDQJ6Y/edit).
- **136102181180071: 7-Parameter AF3-Based Structural Novelty Index (SNI) for Identifying Unconventional NRC-NLRs**

  $\def\mathcal#1{\mathit{#1}}\def\mathscr#1{\mathit{#1}}$
  The Structural Novelty Index (SNI) is a multi-parametric framework designed to identify unconventional NRC-NLRs by quantifying their structural divergence from canonical hexameric resistosomes like NRC2, 3, and 4. This index integrates seven quantitative metrics—including P-loop cleft depth, CC-to-NB-ARC tilt angles, interface gap volumes, and pore assembly radii—extracted from AlphaFold 3 (AF3) models of monomers and simulated "forced hexamers." By normalizing these measurements against a structural baseline derived from ground-truth datasets (e.g., PDB 9FP6) and calculating a composite Z-score, the SNI provides a robust mathematical threshold to distinguish standard pore-forming NLRs from atypical variants that may possess unique activation mechanisms or alternative oligomeric states.
  Details: [document](https://docs.google.com/document/d/1tM_C4Q0p5Vceh_R2crYf-Kap4dzDgQmvkaBvAnXxcgE/edit).
- **187642863749828: Defining a Quantitative 8-Parameter Structural Novelty Index (SNI) for Unconventional NRC-NLRs using AF3 Stoichiometric Sampling**

  $\def\mathcal#1{\mathit{#1}}\def\mathscr#1{\mathit{#1}}$
  The proposed Structural Novelty Index (SNI) is a quantitative framework designed to identify unconventional NRC-NLRs by measuring their deviation from the canonical hexameric resistosome architecture (anchored by PDB IDs 9FP6 and 9CC8). The SNI utilizes eight specific geometric and biophysical parameters—including stoichiometric pitch deviation, vertical planarity, NB-ARC hinge angles, and pore constriction electrostatics—extracted from AlphaFold 3 multimer models. By incorporating stoichiometric flexibility (sampling $n=4, 6, 8$) and weighting results by pLDDT confidence scores, the index distinguishes sequences with genuine structural shifts, such as altered oligomeric states, non-conductive pores, or divergent activation hinges, from the standard NRC2/3/4 paradigm. This tiered computational workflow enables the systematic screening of approximately 6,000 sequences across 350 Solanaceae species to isolate novel sensor-like or scaffold NRCs.
  Details: [document](https://docs.google.com/document/d/1nRtg4Dwu7kIPHCG3uY3OKPdwXxjpVoJnFdarJxXheGA/edit).
- **188440674211813: Development of the SNI-CLT for Identifying Unconventional NRC-NLRs via Mechanical Lever Analysis**

  $\def\mathcal#1{\mathit{#1}}\def\mathscr#1{\mathit{#1}}$
  The Structural Novelty Index: Centroid Leverage & Torque (SNI-CLT) is a quantitative framework that treats NRC protomers as mechanical components within a circular arch to identify unconventional NLRs. By utilizing seven specific parameters—including the **$\alpha$4-helix torque angle**, **NB-ARC scissor aperture**, and **centroid pivot arm ratio**—the index measures the physical and geometric "leverage" required for a sequence to form a canonical hexamer. This approach leverages AlphaFold 3 models to detect deviations from the structural baselines of ground-truth resistosomes (such as NbNRC2 and SlNRC3). Sequences that exceed a weighted SNI threshold are flagged as unconventional, signaling potential shifts in stoichiometry or activation mechanisms that distinguish them from the standard ~6,000-member NRC family.
  Details: [document](https://docs.google.com/document/d/1k9jctu2v9U0A8hVVDzG5x7Sqj2-sSXg-RWS0Tp26wyk/edit).
- **62869464099376: SNI-V2 Framework: Integrating CC-Core Packing Density and Empirical Calibration for NRC-NLR Structural Novelty Assessment**

  $\def\mathcal#1{\mathit{#1}}\def\mathscr#1{\mathit{#1}}$
  The SNI-V2 framework introduces a robust, 10-parameter quantitative index designed to distinguish unconventional NRC-NLRs from canonical hexameric resistosomes by analyzing structural features such as protomer angular deviation, MADA helix tilt, and CC-core packing density. To ensure high-throughput efficiency and biological accuracy, the framework utilizes a tiered screening pipeline that begins with an empirical calibration phase using known helpers and sensors to establish data-driven novelty thresholds. By prioritizing candidates through initial monomer scans before progressing to full AlphaFold 3 multimer modeling, the SNI-V2 system provides a computationally optimized and mathematically grounded approach to identifying structural divergence across the ~6,000-sequence NRC family.
  Details: [document](https://docs.google.com/document/d/1Nwn2UAul3b9Zg-kHZoZ7LRyDUYUCzTtS6TuRwqgMTEA/edit).
- **25806249760346: Development of an 8-Parameter Structural Novelty Index (SNI) for Identifying Unconventional NRC-NLRs via AlphaFold 3 Homodimer Modeling**

  $\def\mathcal#1{\mathit{#1}}\def\mathscr#1{\mathit{#1}}$
  The proposed Structural Novelty Index (SNI) is a quantitative framework consisting of eight structural parameters—including tangential protomer rotation ($\theta_{TANG}$), pore radius ($R_{PORE}$), and buried surface area ($BSA_{WHD}$)—designed to mathematically distinguish unconventional NRCs from canonical hexameric resistosomes. The methodology utilizes AlphaFold 3 to model ~6,000 NRC sequences as homodimers, extracting geometric and energetic metrics to establish a structural baseline derived from ground-truth models like 9FP6. By calculating Z-scores for these parameters, the pipeline identifies "Novelty Clusters," such as pentameric shifts or non-pore-forming scaffolds, providing a robust computational screen to prioritize evolutionary outliers within the Solanaceae dataset for experimental validation.
  Details: [document](https://docs.google.com/document/d/14DRNQa-H2VqK_zepAWxlwpMDy9xIup2xcKLELhJwZfM/edit).
- **103040864270759: The Ensemble-Refined Structural Novelty Index (eSNI): A Quantitative AF3-Calibrated Framework for Identifying Unconventional NRC-NLR Resistosomes**

  $\def\mathcal#1{\mathit{#1}}\def\mathscr#1{\mathit{#1}}$
  The Ensemble-Refined Structural Novelty Index (eSNI) is a quantitative framework designed to identify unconventional NRC-NLRs by measuring architectural divergence from canonical hexameric resistosomes. It utilizes eight specific geometric and confidence-based parameters—including protomer symmetry angles, MADA helix tilt, and LRR curvature—extracted from AlphaFold 3 (AF3) models. To ensure statistical robustness, the eSNI establishes a baseline using a "Synthetic Canonical Ensemble" of 100 NRC sequences, allowing for a weighted Z-score calculation that accounts for modeling variance and pLDDT confidence. By ranking sequences based on their deviation from this calibrated ensemble, the eSNI effectively filters thousands of Solanaceae NLRs to prioritize high-confidence, structurally unique targets for experimental characterization.
  Details: [document](https://docs.google.com/document/d/1RsgPl6DxPdqVhIWb0i5Lesx9LPBHs11lThWDguwSwPI/edit).
- **230450355145970: Defining a Structural Novelty Index (SNI) for Quantifying Divergence in Unconventional NRC-NLR Hexameric Resitosomes**

  $\def\mathcal#1{\mathit{#1}}\def\mathscr#1{\mathit{#1}}$
  The proposed Structural Novelty Index (SNI) is a quantitative framework consisting of eight structural parameters designed to measure the architectural divergence of NRC-NLR sequences from canonical hexameric resistosomes like NbNRC2 and SlNRC3. By extracting geometric metrics from AlphaFold 3 multimer models—including protomer rotation angles, axial displacement, MADA-helix orientation, and interface complementarity—the SNI calculates a normalized distance from established structural baselines (PDB IDs: 9FP6, 9RI9, 9CC8). This mathematical approach allows for the systematic screening of approximately 6,000 NRC sequences to identify "unconventional" variants, such as pentameric stoichiometry shifters or non-pore-forming signaling helpers, effectively prioritizing structurally unique candidates for functional validation in plant immunity.
  Details: [document](https://docs.google.com/document/d/17nG9OT2qrJTEdAp1trwdB1qsNGVb8KGcgyvznJRvh2s/edit).
- **216898738365837: Establishing a Quantitative 8-Parameter Structural Novelty Index (SNI) for NRC-NLR Classification via AlphaFold 3**

  $\def\mathcal#1{\mathit{#1}}\def\mathscr#1{\mathit{#1}}$
  The Structural Novelty Index (SNI) is a quantitative framework designed to identify unconventional NRC-NLRs by measuring their structural deviation from canonical hexameric resistosomes (NRC2, 3, and 4). Utilizing AlphaFold 3 (AF3) models of truncated homo-hexamers, the SNI integrates eight specific parameters—including rigid-core rotational symmetry ($\theta_{CORE}$), PAE-weighted pore diameter ($D_{PORE}^*$), and CC-domain integration angles—to capture geometric and biophysical divergence. By applying confidence-weighted metrics and Z-score analysis to these parameters, the framework filters the NRC family for candidates with novel mechanism-defining features, such as altered symmetry or modified nucleotide-binding pockets, moving beyond simple sequence homology to pinpoint functionally distinct resistance proteins.
  Details: [document](https://docs.google.com/document/d/1QVZEIq_FbQ-4ni5yyF7vk_EkhHCrGEtSkn8D6P2ZFOk/edit).
- **216227824734088: Physico-Evolutionary Structural Novelty Index (PE-SNI) for High-Throughput Identification of Unconventional NRC-NLRs**

  $\def\mathcal#1{\mathit{#1}}\def\mathscr#1{\mathit{#1}}$
  The Physico-Evolutionary Structural Novelty Index (PE-SNI) is a high-throughput quantitative framework designed to identify unconventional NRC-NLRs by measuring their structural and energetic deviation from canonical hexameric resistosomes (NRC2, 3, and 4). Leveraging AlphaFold 3 (AF3) modeling and the ground truth structures 9FP6, 9RI9, and 9CC8, the PE-SNI evaluates sequences based on seven specific parameters: P-loop solvation depth, CC-NB-ARC resting tilt, interface frustration energy, MADA-motif radial variance, ligand-induced switch delta, evolutionary-weighted apolar density, and linker ensemble dynamics. By calculating an aggregate Z-score across these metrics, the PE-SNI identifies "Unconventional Candidates" that exhibit atypical folding dynamics, interface energetics, or activation mechanisms, moving beyond simple sequence similarity to detect biologically significant structural novelty within the Solanaceae NRC family.
  Details: [document](https://docs.google.com/document/d/1UP7drbO6cr1UCZcUf0AZ5yPvmQ2jHZ9EszboB5bdtHQ/edit).
- **207811288102211: A Tiered AlphaFold 3 Screening Approach Using 8-Parameter Structural Novelty Index for Unconventional NRC-NLR Identification**

  $\def\mathcal#1{\mathit{#1}}\def\mathscr#1{\mathit{#1}}$
  This research proposal introduces a tiered AlphaFold 3 screening strategy to identify unconventional NRC-NLRs by modeling approximately 6,000 sequences as "forced hexamers" to detect structural strain. The methodology centers on a Structural Novelty Index (SNI) comprising eight quantitative parameters—such as Stoichiometric Pitch Deviation, Vertical Planarity, and Pore Charge Density—calculated against canonical ground-truth structures like PDB 9FP6. By measuring "Interface Tension" and deviations from the standard hexameric disk geometry, the workflow efficiently identifies candidates likely to adopt alternative stoichiometries or novel functions, such as altered ion selectivity or non-conductive "decoy" roles. This approach prioritizes high-SNI sequences for targeted structural refinement and experimental validation, providing a robust mathematical framework to map structural diversity across the Solanaceae NRC family.
  Details: [document](https://docs.google.com/document/d/1nEOaxtmOYviYJt0y_734wB8vsCcM8J_GNre1raqcW6Q/edit).
- **265226988747946: A Multi-Parametric Structural Novelty Index (SNI) for Identifying Unconventional NRC-NLRs via AlphaFold 3 Screening**

  $\def\mathcal#1{\mathit{#1}}\def\mathscr#1{\mathit{#1}}$
  The Structural Novelty Index (SNI) is a multi-parametric framework designed to identify unconventional NRC-NLRs by quantifying their structural divergence from canonical hexameric resistosomes using AlphaFold 3 (AF3). The index integrates nine quantitative metrics—such as protomer angular deviation, pore tilt angle, and stoichiometry confidence ($\Delta$ipTM)—extracted through a standardized computational pipeline that compares query sequences against established templates like NbNRC2 and SlNRC3. By employing a three-phased screening process that evaluates both monomeric and multimeric configurations, the SNI categorizes the NRC family into distinct functional groups, including signaling scaffolds, specialized sensors, and non-hexameric divergents. This systematic approach enables the identification of novel immune receptor architectures within large genomic datasets based on precise structural biology metrics rather than broad sequence similarity.
  Details: [document](https://docs.google.com/document/d/1DtSalRnTuer-HpAd3LhLI-Efi4FOpPNJ3HhPnE8AYbs/edit).
- **176579419482520: A Quantitative Structural Novelty Index (SNI) for NRC-NLRs using Canonical Reference Cloud and Weighted Z-score Classification**

  $\def\mathcal#1{\mathit{#1}}\def\mathscr#1{\mathit{#1}}$
  The proposed idea establishes a quantitative Structural Novelty Index (SNI) to differentiate unconventional NRC-NLRs from canonical hexameric resistosomes by first generating a statistical "canonical cloud" from AlphaFold 3 (AF3) models of NRC2, NRC3, and NRC4. It utilizes eight specific geometric and confidence-based parameters—including azimuthal step deviation, pore aperture, and protomer tilt—to calculate a weighted Structural Deviation Score (SDS). This multi-tiered screening approach filters approximately 6,000 sequences by identifying those that significantly deviate from the reference baseline (SDS > 3.0) while maintaining high structural confidence, ultimately prioritizing candidates with high potential for novel immune architectures for experimental validation.
  Details: [document](https://docs.google.com/document/d/1na2VPf2mFtw_LhgcBwG0kR0EWC4JBpjUfLbWr5QirJw/edit).
- **37296448790140: Ensemble-Refined Structural Novelty Index (eSNI) for Characterizing Unconventional NRC-NLR Resistosomes**

  $\def\mathcal#1{\mathit{#1}}\def\mathscr#1{\mathit{#1}}$
  The proposed idea establishes an Ensemble-based Structural Novelty Index (eSNI) that utilizes AlphaFold 3 to generate a statistical baseline from 150 canonical NRC models, providing a robust framework for identifying unconventional NLRs. It defines eight quantitative parameters—including stoichiometric azimuthal shift, LRR eccentricity, and interface torsion—to capture geometric and functional deviations from the standard hexameric resistosome architecture. These metrics are processed through a "Tiered Census" workflow that includes monomeric pre-screening and hexameric modeling, culminating in a Mahalanobis Distance ($D_M$) calculation. This approach allows for the mathematically rigorous classification of sequences, effectively distinguishing high-confidence structural outliers from canonical proteins or modeling artifacts within large datasets.
  Details: [document](https://docs.google.com/document/d/1FBFCKugbnmJJzeIEQ5ZwObOrK8xAU9gnJ_rbgXddO30/edit).
- **116914230292830: Development of a 7-Parameter Structural Novelty Index (SNI) for Identifying Unconventional NRC-NLR Candidates**

  $\def\mathcal#1{\mathit{#1}}\def\mathscr#1{\mathit{#1}}$
  The proposed Structural Novelty Index (SNI) is a quantitative framework designed to distinguish unconventional NRC-NLRs from canonical hexameric resistosomes by measuring structural "deviation" across seven key parameters. Grounded in the structural analysis of NRC2, NRC3, and NRC4 (PDB IDs: 9FP6, 9RI9, 9CC8), the index evaluates AlphaFold 3-modeled sequences based on metrics such as α4-helix curvature, MADA motif fidelity, steric compatibility with hexameric assembly, and predicted pore geometry. By scoring features like N-terminal extensions and NB-ARC domain orientations, the SNI provides a robust mathematical baseline to filter approximately 6,000 NRC sequences for candidates likely to exhibit novel oligomeric states or non-canonical functions, effectively shifting the identification process from broad sequence similarity to specific structural biology metrics.
  Details: [document](https://docs.google.com/document/d/1QNc4uoob7ytrsfuuHHHu-TwwiFqgOKFUxUNJFu3Ba94/edit).
- **119627191265642: A Hierarchical 8-Parameter Structural Novelty Index (SNI) for Distinguishing Unconventional NRC-NLRs via AlphaFold 3**

  $\def\mathcal#1{\mathit{#1}}\def\mathscr#1{\mathit{#1}}$
  The Structural Novelty Index (SNI) is a hierarchical, eight-parameter framework designed to quantitatively distinguish unconventional NRC-NLRs from canonical hexameric resistosomes by analyzing three distinct structural phases: monomeric "readiness," dimeric resting states, and multimeric pore architectures. By leveraging ground truth structures (PDB IDs: 9FP6, 9RI9, 9CC8) and AlphaFold 3 (AF3) modeling, the SNI utilizes robust centroid-based geometries—such as the MADA vector tilt, NB-ARC centroid kink, and pore radial symmetry—to maintain stability against sequence indels and loop variations. To ensure high-fidelity screening across large datasets, the index incorporates confidence-weighting based on pLDDT and Predicted Aligned Error (PAE), establishing a mathematically rigorous baseline for identifying novel immune receptors with divergent activation mechanisms or atypical stoichiometries.
  Details: [document](https://docs.google.com/document/d/1HbacZLv4ndgEoxZZWNYyDNcpEAH58h0tp2lWUQJH__k/edit).
- **102303763741952: Template-Anchored Geometric Divergence for Quantifying NRC-NLR Structural Novelty via AlphaFold 3**

  $\def\mathcal#1{\mathit{#1}}\def\mathscr#1{\mathit{#1}}$
  The Procrustean Structural Novelty Index (SNI) is a quantitative framework designed to identify unconventional NRC-NLRs by measuring geometric and biochemical deviations from the canonical hexameric resistosome template (PDB: 9FP6). By anchoring AlphaFold 3-predicted monomers to the conserved NB-ARC "engine" of NbNRC2, the index calculates seven specific parameters—including the hexameric torque angle, CC-domain radial displacement, and azimuthal drift of the LRR—to assess stoichiometry and pore-forming potential. This systematic alignment allows for the rapid screening of large datasets, using metrics like the "Scissor" aperture and hydrophobic mismatch to distinguish standard immune signaling hubs from structurally divergent scaffolds, decoys, or pentameric variants.
  Details: [document](https://docs.google.com/document/d/1I09KaerfbiEsXzeh9JHVSo-g3GSeBYInIROxmalYOFQ/edit).
- **277192430609300: Development of a 7-Parameter Structural Novelty Index for Identifying Unconventional NRC-NLRs using AlphaFold 3**

  $\def\mathcal#1{\mathit{#1}}\def\mathscr#1{\mathit{#1}}$
  The proposed Structural Novelty Index (SNI) is a quantitative framework comprising seven distinct parameters designed to distinguish unconventional NRC-NLRs from canonical hexameric resistosomes like NRC2, NRC3, and NRC4. By leveraging AlphaFold 3 (AF3) modeling and structural analysis of PDB coordinates, the SNI evaluates specific geometric and biophysical features, including the C-terminal Disordered Volume Ratio (CDVR), HD1-WHD pivot angles for oligomeric curvature, $\alpha4$-helix kink magnitudes, MADA bundle pore diameters, and LRR solenoid curvature. Benchmarked against ground-truth structures (PDB IDs: 9FP6, 9RI9, and 9CC8), this index provides an automated, high-throughput pipeline to identify structural outliers among thousands of sequences, prioritizing proteins with significant deviations in interface stability, pore functionality, or domain orientation for further biochemical validation.
  Details: [document](https://docs.google.com/document/d/1Of0VbC_qL2wNR-YZrwgUCc9FAplEDVLOqcMFhvjfW6I/edit).
- **1008902880708: Quantitative SEED Index for Identifying Unconventional NRC-NLRs via Tiered AlphaFold 3 Screening**

  $\def\mathcal#1{\mathit{#1}}\def\mathscr#1{\mathit{#1}}$
  The Structural Evolution and Divergence Index (SEDI) is a quantitative framework designed to identify unconventional NRC-NLRs by measuring their structural deviation from the canonical hexameric resistosome paradigm. The index utilizes eight specific parameters—including P-loop solvent accessibility, LRR curvature, MADA pore-axis displacement, and interface packing efficiency—to calculate weighted Z-scores relative to established reference structures like NRC2, NRC3, and NRC4. The approach employs a tiered AlphaFold 3 (AF3) screening protocol that first analyzes monomeric features, such as NB-ARC switch proximity and CC-LRR occlusion, before proceeding to intensive hexameric modeling for high-potential candidates. By integrating evolutionary conservation scores with physical interface metrics and AF3 confidence weights, SEDI provides a robust, computationally efficient method for distinguishing novel immune sensors or non-canonical executors from standard signaling components within large genomic datasets.
  Details: [document](https://docs.google.com/document/d/1BfBjpyAbJWuusGDsfKmY0-uCG9dNC_oN06NGN2lhH6o/edit).

All hypotheses: [spreadsheet](https://docs.google.com/spreadsheets/d/1Y4zApzDaXltkgQN3njZspEjHPk8KjbYf_XOG7dVzup0/edit?gid=0#gid=0).

References:
