## Supplementary Data S6 for "AI-guided discovery of atypical protein assemblies": [EXTERNAL] ├░Y╠ê┬o┬1⁄4 Co-scientist - Top ranking hypotheses 230897543 - Quantitative Structural Novelty Index (SNI) for NRC-NLR Proteins - Google confidential - For research purposes only.docx

$\def\mathcal#1{\mathit{#1}}\def\mathscr#1{\mathit{#1}}$

### Comparative Analysis of Structural Novelty Indices (SNI) for Unconventional NRC-NLR Identification

Prepared by 🧬 AI co-scientist on 2026-01-28. For research purposes only.

#### Research Goal

The primary objective of this research is to define a quantitative Structural Novelty Index (SNI) comprising 5–10 parameters that can robustly distinguish unconventional NRC-NLRs (Nucleotide-binding Leucine-rich Repeat receptors) from canonical hexameric NRC resistosomes. By leveraging high-throughput AlphaFold 3 (AF3) modeling and structural insights from ground-truth datasets (PDB IDs: 9FP6, 9RI9, and 9CC8), the SNI aims to establish a mathematical baseline for "deviation." This ensures the screening of approximately 6,000 sequences across ~350 Solanaceae species is grounded in structural biology metrics—such as protomer interface angles, N-terminal CC-domain variations, and NB-ARC conservation patterns—rather than broad sequence similarity.

#### Evaluation Criteria

To ensure the proposed SNI is both biologically relevant and computationally feasible, the following criteria are employed:

| **Criterion** | **Importance** |
| --- | --- |
| **Quantitative Extractability** | Parameters must be extractable from AF3 or PDB coordinates using automated scripts (e.g., Biopython) to facilitate high-throughput screening. |
| **Mechanistic Grounding** | Metrics must correlate with known functional transitions, such as the "death-switch" deployment or the NB-ARC molecular engine activation. |
| **Stoichiometric Sensitivity** | The index must detect subtle geometric shifts (e.g., inter-domain angles) that dictate whether a sequence forms a pentamer, hexamer, or octamer. |
| **Computational Scalability** | The methodology must balance the accuracy of multimeric modeling with the GPU resource requirements for screening 6,000 sequences. |
| **Statistical Robustness** | The index should use Z-score normalization against ground-truth structures to distinguish genuine biological novelty from modeling noise. |

#### Main Research Directions

Current research into NRC-NLR diversity is shifting from sequence-based phylogeny toward predictive structural mechanics. One major direction is the **"Industrialization" of Mechanical Determinants**, where qualitative cryo-EM observations (like the $\alpha4$-helix kink) are converted into quantitative benchmarks to predict stoichiometry. This is worth exploring because sequence identity often fails to distinguish helpers from sensors; however, a 10° shift in the NB-ARC inter-domain angle is a physical requirement for hexamerization.

Another critical direction is **Activation State Geometry Modeling**. By quantifying the "openness" of the catalytic pocket (MHD-to-P-loop distance), researchers can identify "broken" or "rewired" engines in unconventional variants. Furthermore, **N-terminal Funnel Biophysics** is an essential area, focusing on the hydrophobic moment and tilt of the MADA motif to predict membrane-piercing capability. Unexpectedly, recent synthesis suggests a **"Linker-Tether" Constraint**, where the length of the CC-NB-ARC linker may physically limit the "activation radius" of the signaling domain, effectively preventing pore formation even if the MADA motif is present.

#### Candidate Ideas

##### Idea 1: 8-Parameter SNI for AF3 Multimeric Models

This framework defines the SNI through eight specific geometric parameters extracted from AlphaFold 3 multimeric models. The motivation for this approach is to move beyond "black-box" AF3 confidence scores (ipTM) and provide a mechanistic explanation for structural divergence. The index measures the Inter-Protomer Rotation Angle ($\theta_{ROT}$), where 60° defines a canonical hexamer, and the Pore Constriction Aperture ($D_{APEX}$), which measures the "openness" of the death switch. Other parameters include the $\alpha1$-Helix Inclination Vector ($\phi_{TILT}$), the MHD-to-P-loop Spatial Gap ($D_{MHD-P}$), and the "Death-Switch" Amphipathic Moment ($\mu_H$). Additionally, it tracks the Winged-Helix Domain (WHD) interface buried surface area ($BSA_{WHD}$), core hydrophobicity ($H_{ABS}$), and the LRR-to-CC Proximity Index ($D_{INHIB}$).

The evidence supporting this idea lies in the remarkable consistency of these metrics across ground-truth structures; for instance, $D_{MHD-P}$ is strictly $14.5 \pm 0.3$ Å in active canonical states. By quantifying these, the SNI can identify "hidden" helpers that lack sequence motifs but maintain the necessary geometry for pore formation.

| **Category** | **Description** |
| --- | --- |
| **Structural Focus** | Full hexameric assembly and activation-state mechanics. |
| **Technical Requirement** | High (Requires AF3 multimer modeling of 6,000 homo-hexamers). |
| **Discriminatory Power** | High; distinguishes between non-pore-forming signaling NLRs and true helpers. |

**Judgment:** This idea addresses the research goal comprehensively by providing a robust, multi-domain metric set that is directly grounded in experimental PDB data.

##### Idea 2: 8-Parameter SNI for AF3 Homodimer Models

This approach establishes a quantitative framework by analyzing eight parameters extracted from AF3 homodimer models, which are computationally more efficient than full hexamers. Metrics include the Azimuthal Protomer Turning Angle ($\theta_{turn}$), Catalytic Core Expansion ($d_{B-MHD}$), and Interface Buried Surface Area (iBSA). The motivation is that dimer interface geometry and the resulting curvature are the primary physical determinants of higher-order stoichiometry. A 10° shift in the inter-domain angle is the established mechanism distinguishing pentameric ZAR1 from hexameric NRC2.

| **Category** | **Description** |
| --- | --- |
| **Structural Focus** | Dimer interface stability and curvature as a proxy for stoichiometry. |
| **Technical Requirement** | Moderate; homodimer modeling is an order of magnitude faster than hexamers. |
| **Discriminatory Power** | Moderate; prone to errors if structural decorations (LRRs) shift principal axes. |

**Judgment:** While computationally attractive, this idea is weakened by a factual error in its baseline calibration (misidentifying 9FP6 as octameric) and requires significant recalibration of its turning angle logic to be valid.

##### Idea 3: 10-Parameter SNI for Comprehensive Outlier Detection

This framework utilizes ten metrics to filter the Solanaceae dataset, focusing on "unconventional" flags when a sequence deviates by $>2.5$ standard deviations in at least three parameters. Unique parameters include the LRR Toroidal Radius ($R_{LRR}$), Catalytic Core Channel Connectivity ($C_{core}$), and the C-terminal Tail Flexibility Index ($pLDDT_{tail}$). The motivation is to shift the research paradigm from "canonical classification" to "outlier detection," identifying rare regulatory-only NLRs.

| **Category** | **Description** |
| --- | --- |
| **Structural Focus** | Global quaternary dimensions and activation core accessibility. |
| **Technical Requirement** | High; includes complex metrics like channel connectivity and solvation energy. |
| **Discriminatory Power** | High; identifies "Black Swan" NLRs with structured sensory extensions. |

**Judgment:** The idea is scientifically sophisticated but suffers from technical flaws in its geometric definitions (e.g., defining a dihedral with only three points) and physically contradictory modeling of activation states.

##### Idea 4: 6-Parameter SNI Using Interface and Domain Geometry

This index focuses on six parameters derived from both monomeric and dimeric AF3 models to assess domain geometry and oligomerization potential. Key metrics include the MADA-helix length ($L_{MADA}$), $\alpha4$-helix curvature ($\theta_{kink}$), and the CC–NBD "Wedge" Orientation ($\Omega_{W}$). The motivation is that stoichiometry is physically constrained by monomeric geometry; for example, the inter-domain angle between CC and NB-ARC is 85° in hexamers vs. 75° in pentamers.

| **Category** | **Description** |
| --- | --- |
| **Structural Focus** | Monomer-driven mechanical constraints on higher-order assembly. |
| **Technical Requirement** | Moderate; tiered screening (monomers then dimers) is highly efficient. |
| **Discriminatory Power** | High; effectively separates 5x, 6x, and 8x states using structural "switches." |

**Judgment:** This is a highly practical and technically sound approach that leverages the most recent (2024-2025) cryo-EM literature to define Stoichiometry Determinants.

##### Idea 5: 7-Parameter SNI Focusing on Volumetric Ratios

This proposal evaluates features like the C-terminal Disordered Volume Ratio (CDVR) and the LRR Solenoid Curvature Index ($\Omega_{LRR}$). The CDVR is a particularly innovative metric motivated by the need to identify "sensor-hybrids" that possess large, disordered integrated domains. However, the idea is hindered by incorrect numerical benchmarks for pore apertures ($D_{pore}$), which would misclassify canonical NRCs as unconventional.

| **Category** | **Description** |
| --- | --- |
| **Structural Focus** | Domain volumetric distribution and solenoid curvature. |
| **Technical Requirement** | Moderate; utilizes standard AF3 outputs and JSON parsing. |
| **Discriminatory Power** | Moderate; provides unique insights into sensors but has flawed thresholds. |

**Judgment:** The conceptual framework (CDVR) is strong, but the numerical foundation requires a total recalibration to be scientifically viable.

##### Idea 6: 8-Parameter SNI with Weighted Scoring

This idea utilizes a weighted Z-score system focusing on parameters like the NB-ARC Swivel Angle ($\phi_{swivel}$) and the Polarity Profile of the Pore Inner-Wall (PPIW). It leverages AF3 to model the N-terminal MADA motif, which is often unresolved in cryo-EM. The motivation is to capture a holistic "structural signature" rather than local mutations.

| **Category** | **Description** |
| --- | --- |
| **Structural Focus** | Holistic topological profiling of the CC, NB-ARC, and LRR domains. |
| **Technical Requirement** | High; requires complex 3D circle-fitting for LRR curvature. |
| **Discriminatory Power** | High; predicts activation thresholds and signaling mechanisms. |

**Judgment:** This is a mathematically rigorous framework, though it risks comparing "models against models" where experimental data for the N-terminus is lacking.

##### Idea 7: 8-Parameter SNI Using Weighted Z-Scores

Similar to Idea 6, this framework aggregates metrics such as the ARC1-ARC2 Clamshell Aperture ($\Psi_{ARC}$) and the MHD-Sensory Loop Proximity ($\delta_{MHD}$). It weights angular deviation and helix tilt most heavily. The motivation is to shift from confidence metrics to actual topological analysis.

| **Category** | **Description** |
| --- | --- |
| **Structural Focus** | Inter-domain distances and quaternary symmetry. |
| **Technical Requirement** | High; involves Euclidean span measurements and hydrophobic alignment vectors. |
| **Discriminatory Power** | High; identifies "cloaked" N-termini and altered ligand-binding roles. |

**Judgment:** This is a solid candidate that effectively translates qualitative structural biology into a high-throughput pipeline.

##### Idea 8: 7-Parameter SNI focusing on Vertical Planarity

This index proposes metrics like NB-ARC Domain Vertical Planarity ($\delta_{z}$) and N-terminal Radial Displacement ($\Delta R_{Met1}$). However, the idea is critically flawed: it assumes NRCs might form "lock-washer" structures without evidence and uses a resting-state dimer (9RI9) as a reference for a hexameric index.

| **Category** | **Description** |
| --- | --- |
| **Structural Focus** | 3D assembly shifts and pore vestibule electrostatics. |
| **Technical Requirement** | Moderate. |
| **Discriminatory Power** | Low; high susceptibility to modeling artifacts and "hallucinated" novelty. |

**Judgment:** This idea is not recommended due to fundamental flaws in its biological assumptions and baseline calibration.

##### Idea 9: 8-Parameter SNI with Isosteric Polarity Shift

This framework introduces the Isosteric Polarity Shift ($\chi_{iso}$), detecting "stealth" divergence where residue volume is preserved but chemistry is inverted. It also measures Signaling Funnel Pitch ($\rho_{SF}$). The motivation is to catch functional divergence that simple sequence-volume modeling would miss.

| **Category** | **Description** |
| --- | --- |
| **Structural Focus** | Chemical compatibility of interfaces and funnel mechanics. |
| **Technical Requirement** | Extremely High; modeling full hexamers ($>5,000$ residues) pushes AF3 limits. |
| **Discriminatory Power** | High; excellent at identifying non-hexameric NRCs that appear canonical by sequence. |

**Judgment:** The $\chi_{iso}$ parameter is highly insightful, but the computational requirements for full-scale 6,000-sequence screening are prohibitive.

##### Idea 10: 7-Parameter SNI for High-Throughput Screening

This proposal uses metrics like Assembly Symmetry Deviation ($\sigma_{sym}$) and $\alpha4$-helix kink angles ($\theta_{kink}$). It utilizes a $C_6$ symmetry operator to identify sequences that simply do not "fit" the hexameric mold.

| **Category** | **Description** |
| --- | --- |
| **Structural Focus** | Symmetry deviation and tangential LRR angles. |
| **Technical Requirement** | High; requires immense GPU resources for 6,000 hexamers. |
| **Discriminatory Power** | Moderate; establishes a strong baseline for Stoichiometric Novelty. |

**Judgment:** The use of symmetry operators is a clever way to detect "symmetry breakers," but the idea relies on fixed residue numbering, which is fragile in diverse datasets.

#### Comparison of Candidate Ideas

The candidate ideas diverge primarily in their **structural modeling depth** and **parameter specificity**. Ideas 1, 3, 6, 7, 9, and 10 rely on multimeric (hexameric) modeling, which provides the most accurate view of the "active" resistosome but faces severe computational bottlenecks. In contrast, Idea 4 (and Idea 2) focuses on monomeric and dimeric geometry, arguing that the "hinge" mechanics within a single protomer are sufficient to predict higher-order assembly.

Evidence points more strongly toward the parameters used in **Idea 1** and **Idea 4**. The $D_{MHD-P}$ distance and $\theta_{kink}$ angle are directly validated by high-resolution cryo-EM data (9FP6, 9CC8) as critical functional switches. Ideas that incorporate AF3 confidence metrics as a filter (e.g., Ideas 5 and 7) are more robust against modeling failures than those that rely solely on geometric coordinates (e.g., Idea 8).

##### Idea Comparison Table

| **Idea** | **Key Distinguishing Attribute** | **Computational Scalability** | **Supporting Evidence Basis** | **Primary Novelty Parameter** |
| --- | --- | --- | --- | --- |
| **1** | Multimeric Mechanistic Triage | Low (Full Hexamers) | $D_{MHD-P}$ (14.5Å) and $D_{APEX}$ (24Å) benchmarks | Inter-Protomer Rotation ($\theta_{ROT}$) |
| **2** | Homodimer-only Curvature | High | curved interface logic | Azimuthal Turning Angle ($\theta_{turn}$) |
| **3** | Outlier/Black Swan Focus | Low | Toroidal radius consequences | LRR Toroidal Radius ($R_{LRR}$) |
| **4** | Monomer "Hinge" Geometry | Moderate (Tiered) | $\alpha4$-helix kink at L126 | CC–NBD Wedge Angle ($\Omega_{W}$) |
| **5** | Volumetric ID Detection | Moderate | CDVR for sensor-hybrids | C-terminal Disordered Ratio (CDVR) |
| **9** | Interface Chemical Inversion | Very Low | Kyte-Doolittle interface shifts | Isosteric Polarity Shift ($\chi_{iso}$) |
| **10** | Symmetry Deviation | Low | $C_6$ operator RMSD | Assembly Symmetry ($\sigma_{sym}$) |

#### Comparison with Existing Solutions

Traditional solutions for identifying unconventional NLRs rely on **HMM-based motif detection** (e.g., searching for the MADA motif) or **sequence-based phylogeny**. While efficient, these methods fail to distinguish between functional helpers and non-functional decoys (like NRCX) that retain motifs but lack the required "hinge" geometry. The proposed SNIs represent a shift toward **Predictive Structural Bioinformatics**, where the physical mechanics of the protein are quantified.

| **Method** | **Approach** | **Sensitivity to Novelty** | **Scalability** |
| --- | --- | --- | --- |
| **Existing: BLAST/HMM** | Sequence Identity/Motifs | Low (misses decoys) | Very High |
| **Existing: AF3 ipTM** | Confidence of folding | Moderate (black-box) | Moderate |
| **Proposed: SNI** | Quantitative Geometry | High (mechanistic) | Moderate |

#### Unexpected Connections

A critical synthesis of the ideas reveals a **mechanical relationship between linker length and membrane insertion**. Specifically, a short CC-NB-ARC linker acts as a "tether," potentially preventing the 180-degree "flip-out" of the MADA motif required for pore formation. This suggests that the **Linker Displacement Ratio ($L_{DR}$)**—the ratio of linker length to CC-domain height—should be a prioritized parameter for the SNI.

Furthermore, the concept of **Stoichiometric Plasticity** emerged: unconventional NLRs may show high ipTM scores across *multiple* stoichiometries (5x, 6x, and 8x) in AF3, whereas canonical helpers show a sharp "confidence peak" at 6x. The **$\Delta$-ipTM** (difference between best and second-best fit) could thus be a powerful digital signature for structural specialization.

#### Relevant Research Contacts

The following researchers are essential for the validation and implementation of the SNI:

- **AmirAli Toghani & Sophien Kamoun (The Sainsbury Laboratory):** Experts in using AF3 to distinguish sensor/helper configurations and using lipid proxies to stabilize "death-switch" funnels.
- **Jogi Madhuprakash & Michael W. Webster (The Sainsbury Laboratory):** Primary experts for the 9FP6 (NbNRC2) ground truth; they provide the biochemical and cryo-EM pipelines to validate unconventional stoichiometries.
- **Hiroaki Adachi (The Sainsbury Laboratory):** The discoverer of the MADA motif, critical for validating if SNI-predicted $\alpha1$ deviations correlate with loss of function.
- **Furong Liu (UC Berkeley):** Expert on the NRC4 (9CC8) interface and the residue patterns governing hexameric vs. dodecameric assemblies.

#### Recommendation and Best Next Steps

It is recommended to proceed with **Idea 1 (8-Parameter SNI)** as the primary framework, but with a **tiered implementation strategy** inspired by Idea 4 to ensure computational feasibility.

##### Best Next Steps:

1. **Pilot Benchmarking (Weeks 1–2):** Run the 8-parameter SNI against a control set of known structures (NRC2/NRC4 hexamers vs. ZAR1 pentamer vs. WAI3 octamer). This will calibrate the "Canonical vs. Unconventional" Z-score thresholds.
2. **Tiered Screening (Weeks 3–8):**
   - **Phase A:** Screen all 6,000 sequences as AF3 monomers to calculate the $\alpha4$-helix kink ($\theta_{kink}$) and Wedge Angle ($\Omega_{W}$).
   - **Phase B:** Prioritize sequences with high monomeric SNI scores for AF3 hexamer modeling to extract the full 8-parameter index.
3. **Experimental Triage (Month 3+):** Select top "High-Interest" outliers for *in vitro* validation (e.g., mass photometry or cryo-EM) to confirm unconventional stoichiometry or activation mechanisms.

This tiered approach balances the mechanistic depth of Idea 1 with the high-throughput scalability required for a 6,000-sequence dataset.

#### References:

### Top ranking proposals

Development of a 7-Parameter Structural Novelty Index for Identifying Unconventional NRC-NLRs using AlphaFold 3
