## Supplementary Data S6 for "AI-guided discovery of atypical protein assemblies": [EXTERNAL] ├░Y╠ê┬o┬1⁄4 Co-scientist - Top ranking hypotheses - Quantitative Structural Novelty Index (SNI) for NRC-NLR Proteins - Google confidential - For research purposes only.pdf

**[11]** A disease resistance protein triggers oligomerization of its NLR helper into a hexameric resistosome to mediate innate immunity - PubMed Central.

**[12]** Jogi, AmirAli, P., Andres, Jake, Jiorgos, et al. A disease resistance protein triggers oligomerization of its NLR helper into a hexameric resistosome to mediate innate immunity. Published 2024. https://www.biorxiv.org/content/10.1101/2024.06.18.599586v1.

**[13]** Class III Peroxidases PRX01, PRX44, and PRX73 Control Root Hair Growth in Arabidopsis thaliana - MDPI.

**[14]** Theoretical Methods for Assessing the Density of Protein Nanodroplets - MDPI.

**[15]** Muniyandi, AmirAli, Hsuan, Yu, Jiorgos, Him, et al. Activation of plant immunity through conversion of a helper NLR homodimer into a resistosome. Published 2024. https://www.ncbi.nlm.nih.gov/pmc/articles/PMC11524475/.

**[16]** 9RI9: Cryo-EM structure of the tomato NRC3 hexameric resistosome - RCSB PDB.

**[17]** Furong, Zhenlin, Chao, Raoul, Wenjie, E., et al. The activated plant NRC4 immune receptor forms a hexameric resistosome. Published 2023. https://www.biorxiv.org/content/10.1101/2023.12.18.571367v1.

**[18]** Subfunctionalization of NRC3 altered the genetic structure of the Nicotiana NRC network - NIH.

**[19]** The activated plant NRC4 immune receptor forms a hexameric resistosome - bioRxiv.

**[20]** Muniyandi, AmirAli, Hsuan, Yu, Jiorgos, Him, et al. Activation of plant immunity through conversion of a helper NLR homodimer into a resistosome. Published 2023. https://www.biorxiv.org/content/10.1101/2023.12.17.572070v1.
