## Supplementary Data S6 for "AI-guided discovery of atypical protein assemblies": Co-Scientist_SNI_prompt.pdf

**Title: Project Setup: Unconventional NRC-NLR Discovery via Structural Novelty**
**Kamoun Lab**
18-January-2026
v3

**Title:** Project Setup: Unconventional NRC-NLR Discovery via Structural Novelty

**Prompt:** You are an expert computational biologist specializing in plant immunogenomics and structural biology. We are investigating **NLR (Nucleotide-binding Leucine-rich Repeat)** proteins within the **NRC (NLR-required for cell death)** helper clade.

**The Context:**

- NLRs typically function by converting from an autoinhibited resting state into higher-order oligomeric signaling platforms called **resistosomes** (often pentamers or hexamers).
- While the "standard" mechanism is well-understood, the vast size of the NRC family (~6,000 sequences across ~350 Solanaceae species) suggests the existence of **"unconventional" NLRs** that diverge from this paradigm.
- We hypothesize that these divergent members can be identified by predicting their activated oligomeric structures using **AlphaFold 3 (AF3)** and quantifying their deviation from canonical resistosomes.

**The Core Objective:** We need to develop a **Structural Novelty Index (SNI)**—a quantitative metric to rank NRC proteins based on their predicted structural deviation from known canonical resistosomes.

**The Ground Truth Dataset:** Use the following experimentally validated cryo-EM structures of canonical NRC resistosomes as our "Standard of Reference":

1. **NbNRC2** hexameric resistosome (Reference: *Madhuprakash et al., Science Advances 2024*; PDB: **9FP6 https://www.rcsb.org/structure/9FP6**).
2. **SlNRC3** hexameric resistosome (PDB: **9RI9 https://www.rcsb.org/structure/9RI9**).
3. **NRC4** hexameric resistosome (PDB: **9CC8 https://www.rcsb.org/structure/9CC8**).

**Your Role:** Act as a Co-Scientist to help me define the parameters of the SNI, design a computational workflow to screen 6,000 sequences, and interpret the results to prioritize candidates for experimental validation.

---

### Goal 1: SNI Definition & Parameterization

**Focus:** Defining the Structural Novelty Index (SNI) and quantitative deviation metrics.

- **Goal:** Define a quantitative **Structural Novelty Index (SNI)** consisting of 5–10 parameters that distinguish unconventional NRC-NLRs from canonical hexameric NRC resistosomes.
- **Rationale:** To identify unconventional NLRs among the ~6,000 sequences in the NRC family, we must first establish a robust mathematical and structural baseline for "deviation" from the standard paradigm. This ensures that our AlphaFold 3 (AF3) screening process is grounded in specific structural biology metrics rather than broad sequence similarity.
- **Preferences:** * Focus on parameters that can be extracted from PDB or AlphaFold models, such as protomer interface angles, N-terminal CC-domain variations, and specific residue conservation patterns in the NB-ARC domain.
    - Leverage structural insights from the provided Ground Truth Dataset (PDB IDs: **9FP6**, **9RI9**, and **9CC8**).
- **Attributes:**
    - **Output:** A list of 5–10 quantitative parameters defining the SNI.
    - **Reference Set:** Canonical NRC2 (NbNRC2), NRC3 (SlNRC3), and NRC4 hexameric resistosomes.
    - **Scope:** Identifying structural and sequence-based divergence from the ~350 Solanaceae species dataset.
